# Bottom-up investigation of spatiotemporal glycocalyx dynamics with interferometric scattering microscopy

**DOI:** 10.1101/2025.05.09.653034

**Authors:** Carla M. Brunner, Lorenz Pietsch, Ingo vom Sondern, Michael Röhrl, Cristian Popov, Marius F.W. Trollmann, Richard W. Taylor, Martin Blessing, Cornelia Holler, Karim Almahayni, Sven Ole Jaeschke, Vahid Sandoghdar, Rainer A. Böckmann, Thisbe K. Lindhorst, Leonhard Möckl

## Abstract

Over recent decades, the glycocalyx, an extracellular organelle comprised of a multitude of glycolipids, glycoproteins, proteoglycans and glycoRNA, has gained considerable interest in cellular biology. While research in this field has revealed its tremendous importance in evermore aspects of physiological and pathological cellular processes, many of the principles that govern the role of the glycocalyx in these processes on a molecular level are still unknown. In order to unravel the fundamental laws underlying glycocalyx function, new technologies are required that enable the distinction between individual subprocesses within the intricate environment of the glycocalyx. Here, we establish an experimental platform to investigate the dynamics of the glycocalyx at the nanometer and microsecond length and time scales in a bottom-up fashion. We synthesized defined model glycans and installed them on supported lipid bilayers, assembling glycocalyx model systems with tunable properties. By investigating these tunable model systems with interferometric scattering (iSCAT) microscopy, we gain access to the required spatiotemporal resolution. We found a strong correlation between the molecular structure of several investigated model glycans and global dynamics of the system. Our findings are corroborated by atomistic and coarsegrained molecular dynamics simulations. Our results provide the first direct experimental evidence on the relationship between glycan structure, organization, and dynamics, offering a robust and versatile basis for a quantitative understanding of glycocalyx biology and physics at the molecular level.

## Section 1: Introduction and overview

All cells in the human body are covered by a heavily glycosylated layer, the glycocalyx.^1-4^ It is comprised of glycolipids, glycoproteins, proteoglycans, and the recently discovered glycoRNA.^5, 6^ Depending on the cell type, the glycocalyx varies in thickness between several dozens to few hundreds of nanometers^3, 7, 8^ and plays a fundamental role in various cellular processes such as cell adhesion, cell recognition, immune cell trafficking, cancer development, and bacterial infection.^4, 9-11^ Due to the nanoscale size and considerable structural complexity of the glycocalyx, investigation of its properties still constitutes a substantial challenge.^8, 12, 13^

While innovative methods have fostered progress in all areas of biology over the past decades, glycobiology is an area that has particularly benefitted from methodological advances. Examples include the advent of metabolic oligosaccharide engineering,^14-16^ mass spectrometry methods tailored for the demanding task of glycomics, glycoproteomics and proteoglycomcis^17-20^ as well as advanced optical methods like super-resolution microscopy.^3, 12, 13, 21^ All these advances have opened new insights into glycobiology: Metabolic oligosaccharide engineering enables tracing of individual glycan building units, albeit with no context of the full glycan; mass spectrometry yielded superstructural information, but generally without the spatial context of the source cell or tissue specimen; and super-resolution microscopy revealed nanometer spatial organization, however, without temporal information and lacking the ability to identify the overall glycan structure.

Although previous advances promise to have a lasting impact, continuing progress in the field of functional glycobiology still highly depends on novel scientific methods. Particularly the investigation of fundamental principles of glycocalyx organization remains an elusive problem. This is mainly because studying these phenomena requires techniques that can trace both the location and movement of these molecules over time with exquisite resolution.

In this work, we describe a novel approach to study structure-function relationships in glycocalyx biology at the molecular level in a bottom-up fashion. We establish a framework that enables the inspection of nanometer and microsecond spatiotemporal properties of precisely defined glycocalyx model systems by integrating three core methodologies: organic synthesis of defined glycan species, supported lipid bilayer (SLB) formation,^22^ and iSCAT microscopy.^23, 24^

Our approach, addressing glycocalyx organization and its effect on diffusion is summarized in Figure 1. First, azide-functionalized and biotinylated SLBs (Figure 1a) were prepared according to an established protocol (SI). Additionally, selected propargylated glycans were synthesized (see section 2 for details) where the alkyne moiety of the propargyl aglycon was used to deco-rate the azido-functionalized SLBs with the respective glycan via copper-catalysed azide– alkyne cycloaddition (CuAAC), a reaction commonly referred to as click chemistry^25^ with the respective glycan (Figure 1b).^26, 27^ This approach allows us to employ a wide variety of glycans without the need to adapt the protocol for every individual carbohydrate, regardless of chain length, stereochemistry of the glycosidic linkage or branching of the employed oligosaccharide. Subsequently, we linked monovalent streptavidin-functionalized AuNPs (Figure S76) to biotinylated membrane lipids, which were added as tracers. Then, the diffusing AuNPs were tracked via iSCAT microscopy (Figure 1c and section 3). Due to their high scattering cross-section, gold nanoparticles (AuNPs) provide an excellent iSCAT signal, enabling high spatiotemporal resolution, which makes them an ideal choice for labeling. In the past, they have for instance been used to investigate GM1 diffusion in a SLB.^28, 29^ Employing image and data analysis, we extract the diffusion characteristics of the AuNPs, precisely reporting on the spatiotemporal dynamics of the system (Figure 1 and section 4). It should be stressed that in this approach, AuNPs only indirectly serve as tracking labels. Thus, their diffusion is associated with the diffusion of the lipid to which they are attached, not the diffusion of a glycan. Nevertheless, we also investigated direct tracking of glycans via labeling them with an AuNP. Several of the studied systems were computationally modeled with atomistic and coarse-grained molecular dynamics simulations, confirming our experimental results.

**Figure 1.**
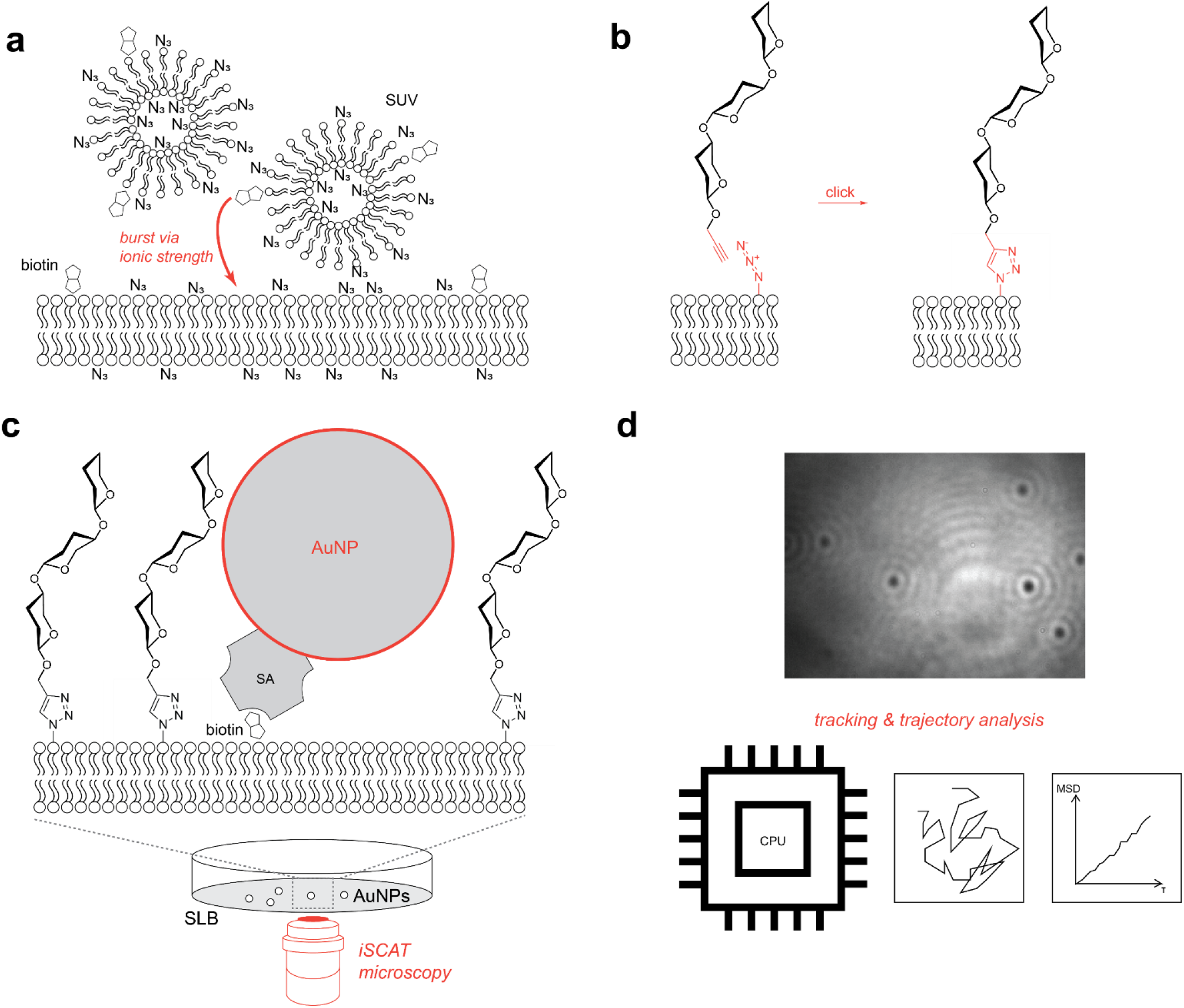
Overview of the experimental workflow. (a) SLBs with defined composition, bearing azido-functionalized and biotinylated lipid, are prepared via bursting of vesicles. (b) Functionalization of the SLBs with model glycans via copper-catalyzed azide-alkyne click chemistry. (c) iSCAT microscopy of diffusing AuNP. (d) A snapshot of a video recorded of diffusing AuNPs. Analysos of the video frames allow us to track individual particles and analyze their trajectories. SUV: small unilamellar vesicles; AuNP: gold nano particle; SA: streptavidin; SLB: supported lipid bilayer.

**Scheme 1.**
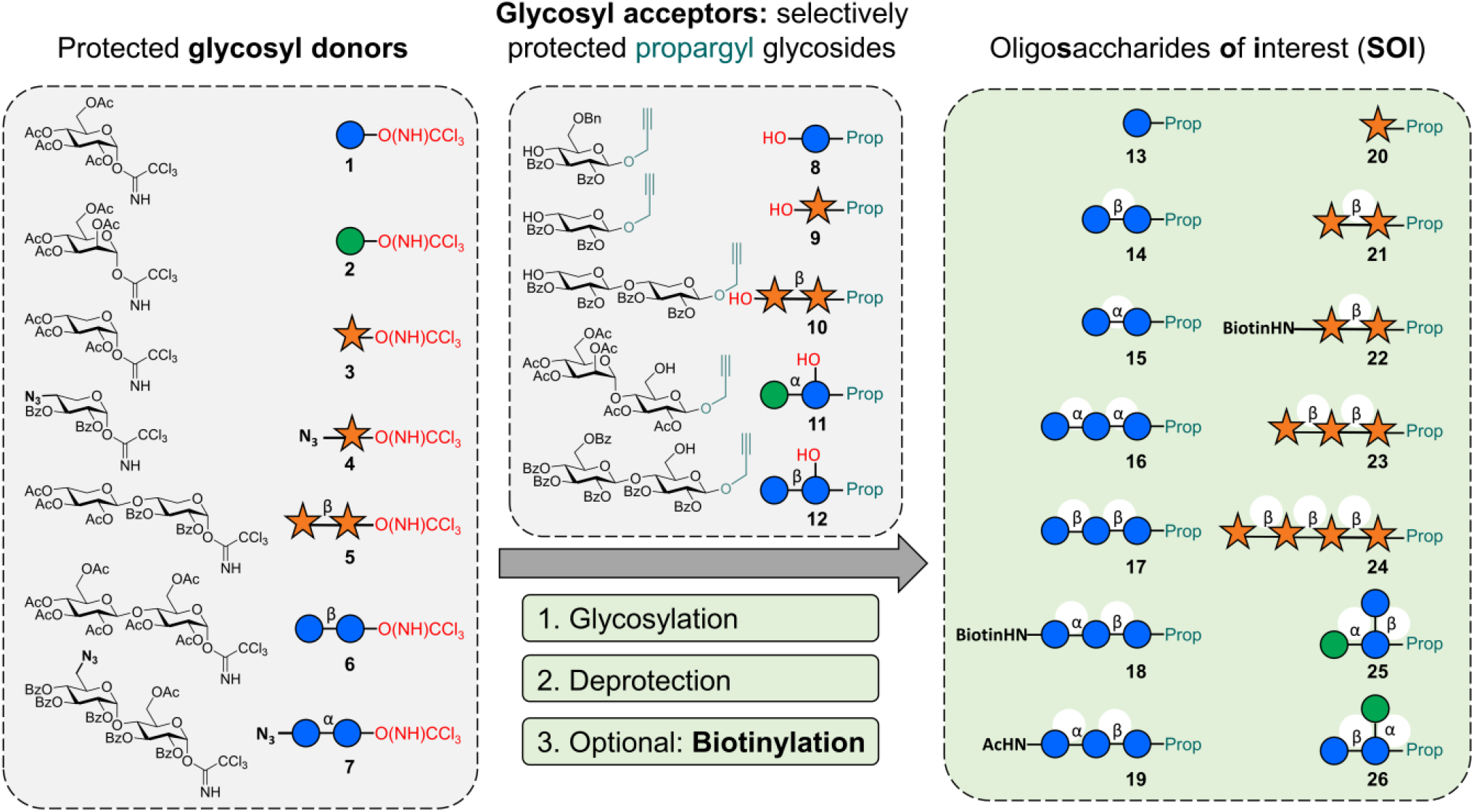
Overview of the oligosaccharide synthesis. Appropriately protected *O*-glycosyl trichloroacetimidates (**1-7**) were employed as glycosyl donors in glycosylation reactions with propargyl glycosides **8-12** as glycosyl acceptors to result in the depicted collection of oligosaccharides of interest (SOIs) **13-26** after deprotection. The SOIs contain 1,4-linked linear saccharides composed of either D-glucose or D-xylose units (**13-24**) or branched trisaccharides composed of D-glucose and D-mannose (**25-26**). They all carry a propargyl aglycon for later conjugation but differ in chain length and stereochemistry of the inter-glycosidic linkages. An optional biotinylation step is facilitated through an azido group present in glycosyl donors **4** and **7**. The symbol nomenclature for glycans (SNFG, blue circle: D-glucose; orange star: D-xylose; green circle: D-mannose) is applied to abbreviate carbohydrate structures.^30^ A complete summary on chemical structures and the synthesis performed is given in the SI. Ac: acetyl; Bz: benzoyl; Bn: benzyl; Prop: propargyl.

## Section 2: Glycan synthesis

A collection of oligosaccharides of interest (SOI, Scheme 1, right) was prepared according to classical glycoside synthesis (Scheme 1).^31^ They were assembled from D-glucose and D-xylose building blocks which were selected as aldohexose and aldopentose examples occurring in the glycocalyx. We relied on the respective *O*-glycosyl trichloroacetimidates^32^ as glycosyl donors (**1-7**) for glycoside synthesis. Propargyl glycosides served as glycosyl acceptors (**8-12**) to enable the later conjugation of the resulting SOIs with azido-functionalized SLBs via CuAAC.^25^ The glycosyl donors were activated by a Lewis acid to trigger the glycosylation of the free hydroxyl group of the selected glycosyl acceptors. For example, the reaction of disaccharide donor **5** with glycosyl acceptor **10** furnished the tetrasaccharide **24** after removal of the OH-protecting groups (deprotection). In order to enable an optional terminal biotinylation of the SOIs, the glycosyl donors can be prefunctionalized with an azido (N_3_) group (as demonstrated in glycosyl donors **4** and **7**, Scheme 1), which serves as a masked amino function in a subsequent peptide coupling step with biotin.

Following this scheme, the propargyl-functionalized saccharides **13-26** of different chain length were prepared as depicted in Scheme 1. Note that the stereochemistry of the glycosidic linkages is strictly controlled in every case, either by the appropriate choice of starting material from the chiral pool (maltobiose, maltotriose and cellobiose) or by stereospecific glycosylation chemistry utilizing neighboring group participation. A comprehensive description of all synthesis entailed can be found in the SI.

## Section 3: SLB glycosylation and iSCAT microscopy

The lipid mixtures used to prepare small unilamellar vesicles (SUVs) were derived from an initial composition consisting of 69% DOPC, 5% DOPS, 1% biotinyl-CAP-PE and 25% DOPE (see SI Chapter 4 for a list of abbreviations). The share of DOPE was partially substituted by its azido-functionalized analogue (DOPE-N_3_) in varying concentrations up to 25% to enable the subsequent introduction of glycans at varying densities (Table 1). We note that, owing to an identical lipid composition for all experimental conditions, all observed effects are caused by the introduced glycans.

**Table 1.** Overview of different conditions for SLB formation. The lipid mixture utilized in SLB formation consists of 69 or 70% DOPC, 0 or 1% biotinyl-CAP-PE, 5% DOPS, 25-n% DOPE-N_**3**_ and n% DOPE. The utilized glycan for SLB functionalization is represented by the corresponding abbreviation.

| Clicked Glycan | N<sub>3</sub>-DOPE (%) | DOPE (%) | biotinyl-CAP-PE (%) |
| --- | --- | --- | --- |
| Glc<sub>1</sub> **13** | 20 | 5 | 1 |
| βGlc<sub>2</sub> **14** | 20 | 5 | 1 |
| αGlc<sub>2</sub> **15** | 20 | 5 | 1 |
| αGlc<sub>3</sub> **16** | 20 | 5 | 1 |
| βGlc<sub>3</sub> **17** | 20 | 5 | 1 |
| Xyl<sub>1</sub> **20** | 20 | 5 | 1 |
| βXyl<sub>3</sub> **23** | 20 | 5 | 1 |
| βXyl<sub>4</sub> **24** | 20 | 5 | 1 |
| 4Man[6Glc]Glc **25** | 20 | 5 | 1 |
| 4Glc[6Man]Glc **26** | 20 | 5 | 1 |
| 1% βGlc<sub>3</sub> **17** | 1 | 24 | 1 |
| 8% βGlc<sub>3</sub> **17** | 8 | 17 | 1 |
| 16% βGlc<sub>3</sub> **17** | 16 | 9 | 1 |
| 25% βGlc<sub>3</sub> **17** | 25 | 0 | 1 |
| 19% αβGlc<sub>3</sub>-NHAc **19** +<br>1% αβGlc<sub>3</sub>-Biotin **18** | 20 | 5 | 0 |
| 19% Xyl<sub>2</sub> **21** +<br>1% Xyl<sub>2</sub>-Biotin **22** | 20 | 5 | 0 |
| 20% αβGlc<sub>3</sub>-Biotin **18** | 20 | 5 | 0 |
| 20% αβGlc<sub>3</sub>-NHAc **19** | 20 | 5 | 0 |
| No glycan | 20 | 5 | 1 |

A composition consisting of 69% DOPC, 5% DOPS, 1% biotinyl-CAP-PE, 5% DOPE and 20% DOPE-N_3_ was selected for investigation of the different glycans **13-17, 20, 23-26** at a constant surface density (Table 1). The same lipid composition was also utilized for functionalization of the formed SLB with only biotinylated glycan **18** and with only non-biotinylated glycan **19**. The latter showed almost no bound AuNPs, confirming the specificity of AuNP-biotin binding in the investigated model system (Figure S79).

In all cases, stable SUVs were obtained, which burst on a glass support via ionic strength, readily forming SLBs. The synthesized glycans were then ligated to the azido-functionalized SLBs via click chemistry. This modular approach enables the reproducible formation of well-defined SLBs that are functionalized with the specific glycan of interest. As the final step of sample preparation, monovalent AuNPs were added to the glycan-decorated SLBs, where they bind to the tracer biotin-functionalized lipids. The diffusing AuNPs thus indirectly report on the dynamics of the glycans on the SLB. Monovalent AuNPs were prepared by treating streptavidin-functionalized AuNPs with a biotin solution at an appropriate concentration to block excess binding sites, yielding monovalent streptavidin AuNPs. For details of SLB preparation and functionalization, see the SI (chapter 4).

Observation of the AuNP diffusion was facilitated by iSCAT microscopy. iSCAT is a super-resolution microscopy technique that relies on the interference of scattered light from nanoparticles with light reflected from the substrate.^33^ Contrary to fluorescence-based techniques, this method does not suffer from photobleaching and allows for tracking nanoparticles with exceptional spatial and temporal resolution^34^ which is required to analyze the motion of AuNPs attached to membrane lipids on the microsecond time scales. Here we use a setup as recently described by Taylor *et al*. (Figure 2).^35^

**Figure 2.**
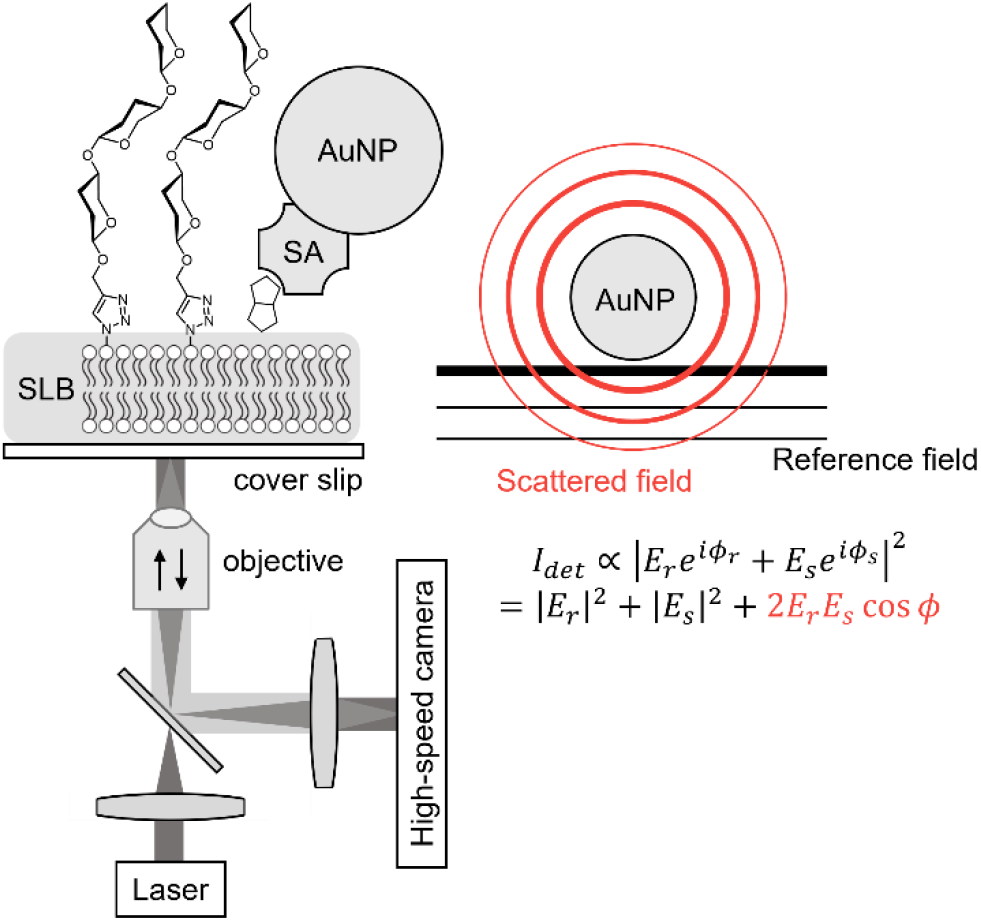
Optical setup for iSCAT microscopy. A collimated laser beam is focused into the back-focal plane of an oil-immersion objective to enable wide-field illumination of the sample. Scattered light from AuNPs is focused onto a camera, interfering with light reflected from the glass-medium interface.

A green laser is used for wide-field illumination of the sample. Scattered and reflected light interfere (complex electric fields denoted by 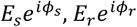 respectively) so that the intensity on the CMOS camera reads^23^

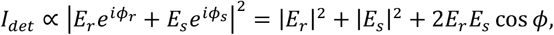

where the phase *ϕ* = *ϕ*_*r*_ − *ϕ*_*s*_ is the phase difference between reflected and scattered fields, including a Gouy phase term.^36^ Note that the pure scattering contribution |*E*_*s*_|^2^ scales with the square of the scattering cross-section and thus with the 6^th^ power of the scattering particle diameter while the interference term scales only with the 3^rd^ power of the diameter. It is therefore possible to retrieve a signal from extremely small particles.

## Section 4: Tracking and analysis

Videos of the AuNP diffusion of 1.1 s duration were recorded with an exposure time of 20 μs and a temporal resolution of 40 μs, corresponding to a frame rate of 25000 fps. This resulted in 27500 frames per video. We automatically tracked AuNPs with a minimum trajectory length of 110 ms using a radial symmetry value (RSV) optimization algorithm inspired by.^37^ A detailed description can be found in the SI (chapter 5). Across the 19 conditions (Table 1), we identified 42035 individual trajectories. Each condition was investigated in three independent technical replicates. The three independent technical replicates showed high consistency of diffusion constants for the same glycan composition (SI, Figure S81).

A global metric describing the diffusion characteristics of a trajectory is the mean squared displacement (MSD). For a time delay *τ*, the MSD returns the square of the distance covered by the particle within *τ*, averaged over the full trajectory:

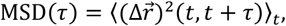

where

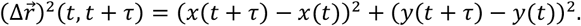

We determined the MSD of each trajectory of every condition for time delays *τ* between 0.04 ms and 27.6 ms. The results are compared between the SOIs. All investigated conditions and comparisons can be found in the SI (e.g., stereochemistry of the glycosidic bonds or glycan regiochemistry, S82, S83).

Here, we now focus on a particularly interesting and relevant case: Differences in SLB dynamics in relation to different glycan chain lengths. Specifically, chain lengths of one, two, and three glucose units as well as of one, three, and four xylose units are considered (Glc **13**, αGlc_2_ **15**, αGlc_3_ **16**; Xyl **20**, βXyl_3_ **23**, βXyl_4_ **24**).

Figure 3 shows the obtained MSD distributions for these structures alongside the negative control (no glycan) in the form of the non-glycosylated SLB for four increasing time delays (*τ* = 6.92, 13.80, 20.68, 27.56 ms). We first note that for every condition the distribution continuously shifts to the right, i.e. to larger MSDs. This is expected since particles undergo, on average, a larger displacement for longer time delays. Interestingly, in both (a) and (b) we also observe a clear difference between the MSD distribution obtained for the monosaccharides (Glc **13**, Xyl **20**) and the corresponding oligosaccharides of longer chain length. Namely, the peaks of the distribution are at lower MSDs in case of the monosaccharides and closer to the peak of the negative control. Thus, for the considered time delays, varying between 6.92 ms and 27.56 ms, AuNPs on SLBs glycosylated with structures of longer chain length exhibit higher diffusivity in our experiments. This holds for both SOIs based on glucose and xylose. Differences in distribution height is due to different total numbers of trajectories (see the SI, Figure S79 for details on selection of trajectories).

**Figure 3.**
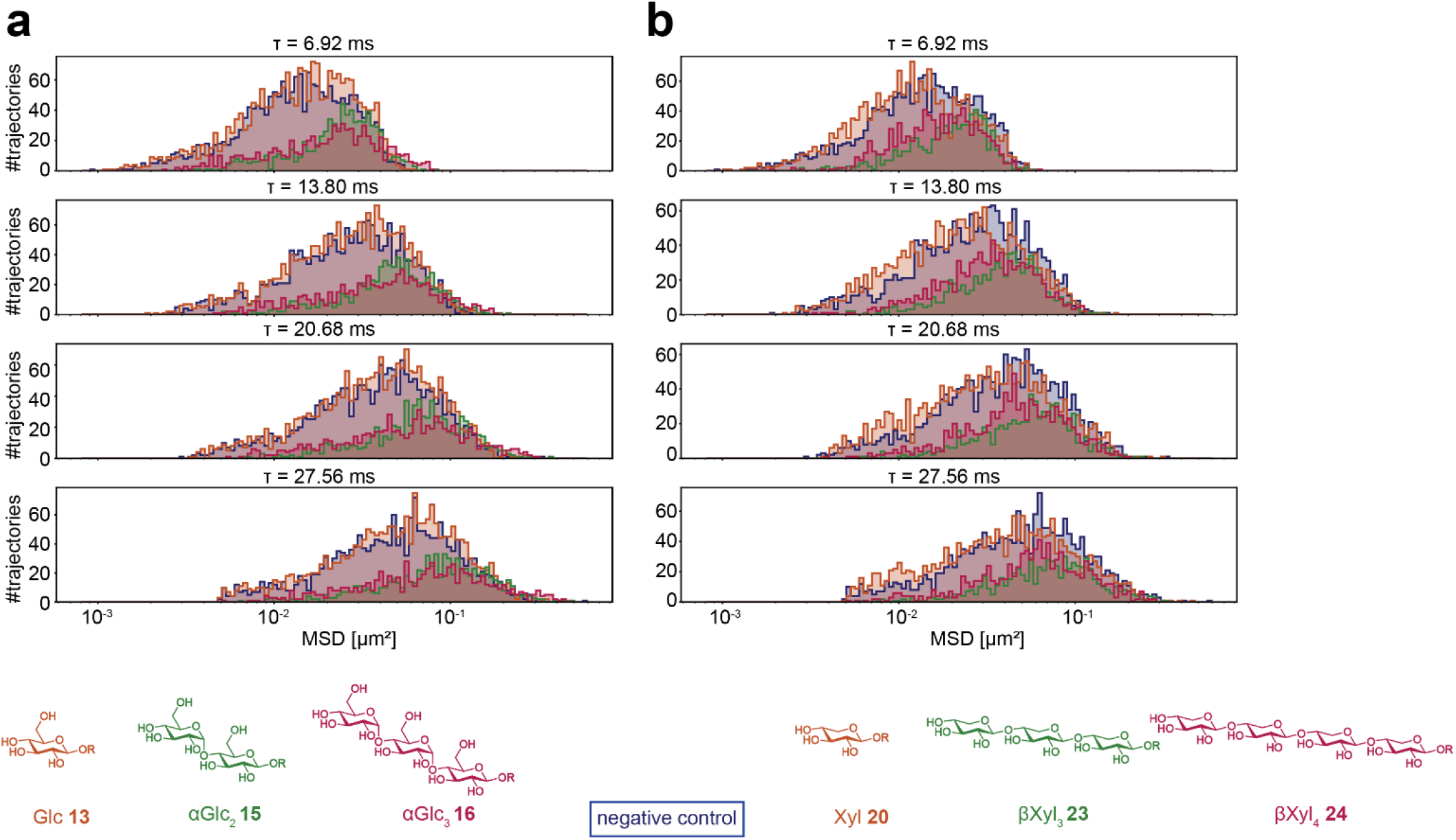
Effect of chain length on the dynamics of the system studied. For glucosides as well as xylosides, histograms of MSDs for two sets of three model glycans alongside the negative control (no glycans on SLB) are shown. (a) MSD histograms for the analysis of glycocalyx model systems containing mono-, di-, and tri-glucosides (orange, green, red, respectively; negative control: blue) (b) MSD histograms of glycocalyx model systems containing mono-, tri-, and tetra-xylosides (orange, green, red, respectively; negative control: blue). R denotes the site where the glycan is connected to the azido-functionalized lipid.

In order to further understand the observed global effects, we turned to local metrics that do not average over the full trajectory. To this end, we first randomly chose 36 full representative trajectories with a duration of 1.1 s from the conditions of interest, equaling 12 from each of the three SLBs prepared for each condition. This way, we obtained the same number of trajectories for all conditions. For each trajectory and at each point in time, we then determined a local anomalous diffusion exponent *α*_*i*_. This is accomplished by fitting a linear function to ln (*SD*) vs ln(*τ*) from *τ*_1_ → *τ*_2_.^38^ Here, SD is the non-averaged squared displacement,

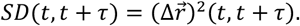

Generally, this can be done for different time scales, here we chose *τ*_1_, *τ*_2_ as 0.8 ms, 1.6 ms, resulting in a favorable tradeoff between time resolution and accuracy. A more detailed analysis of different time scales can be found in the SI (Figure S85 and Table S1). The local anomalous diffusion exponent is a universal metric characterizing (non-)Brownian diffusion. Values larger than 1 are attributed to directed motion while values smaller than 1 indicate confined motion.

Additionally, we calculate a local directional correlation coefficient *C*_*i*_ and deflection angle *φ*_*i*_ at each point in time,

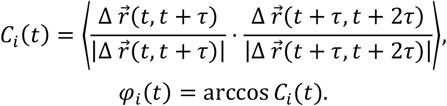

Here, we chose *τ* =0.4 ms. A deflection angle of 0° implies continuous motion in the same direction whereas a deflection angle of 180° signifies deflection in the opposite direction. Again, we compare these metrics for the trajectories obtained on the SLBs glycosylated with SOIs of different chain length (Figure 4): (a) Glc **13**, αGlc_2_ **15**, αGlc_3_ **16** and (b) Xyl **20**, βXyl_3_ **23**, βXyl_4_ **24** alongside the negative control.

**Figure 4.**
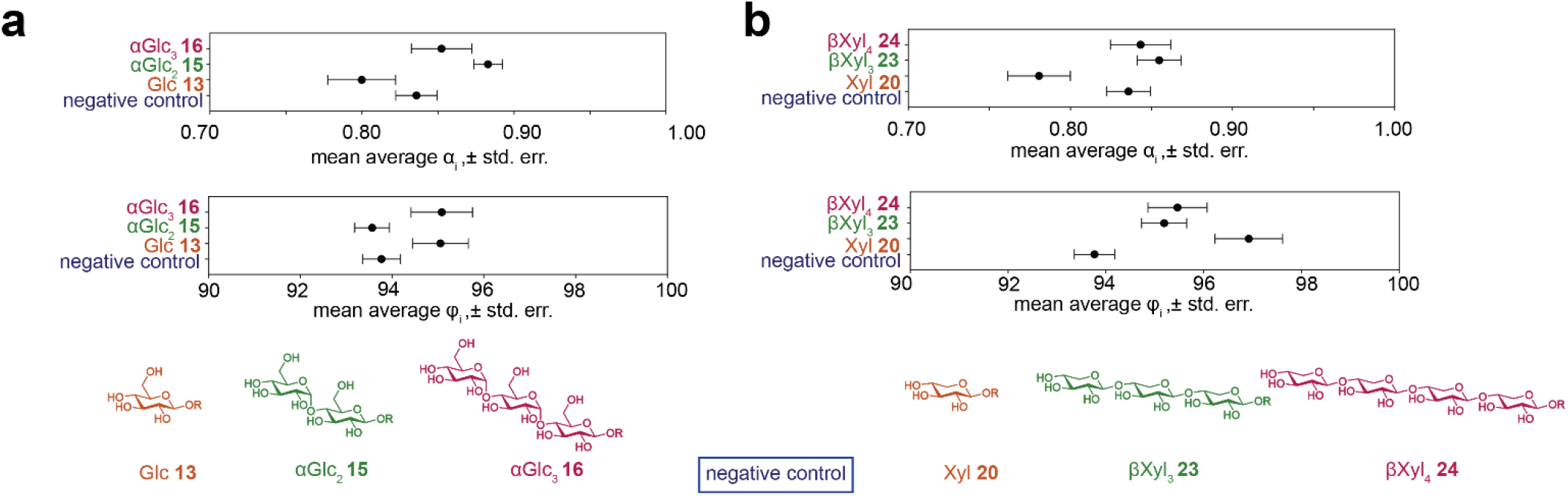
Local anomalous diffusion exponents and local deflection angles for glycocalyx model systems containing (a) mono-, di-, and triglucoside; (b) mono-, tri-, and tetra-xyloside (orange, green, red, respectively; negative control: blue). R denotes the site where the glycan is connected to the azido-functionalized lipid. Error bars: standard error of the mean.

For both the local anomalous diffusion exponent *α*_*i*_ and the deflection angle *φ*_*i*_, we first take the temporal average for each representative trajectory. For glucosides as well as xy-losides, the local *α*_*i*_ are significantly lower for the monosaccharide condition than for the di-, tri- or tetrasaccharides. This is related to more confined diffusion. Similarly, in (a), the deflection angles *φ*_*i*_ are larger for the monosaccharide Glc **13** than for the negative control and the disaccharide αGlc_2_ **15**, but do not significantly differ from those obtained for the trisaccharide αGlc_3_ **16**. In (b), the average values of *φ*_*i*_ for the monosaccharide Xyl **20** are larger than for either the negative control or the tri- and tetrasaccharide βXyl_3_ **23**, βXyl_4_ **24**.

We can conclude that a lower global diffusivity for the monosaccharide conditions, as observed in the previous global MSD analysis, manifests itself in more confined diffusion at time scales below 1 ms and vice versa. The pattern shown here can be identified across all considered time scales.

Taken together, these results indicate less confined diffusion of AuNP attached lipids in the lipid membrane if glycans on the SLB are of longer chain length. This observation can be explained by the assumption that for longer chain lengths, the glycans start to interact with each other, yielding “islands” of associated glycans, which leads to less confined diffusion of the AuNPs outside the glycan islands.

Interestingly, our results indicate that this behavior strongly depends on whether oligosaccharides are linear or branched. This is evident from studying the dynamics of the branched trisaccharides 4Man[6Glc]Glc **25** and 4Glc[6Man]Glc **26** (Figure 5(a)). On a global scale, quantified by the MSD distributions for multiple time delays as introduced in section 4, we obtained a slightly higher average diffusivity in negative control samples compared to both samples with glycans present. This indicates that for these two species, diffusion of the AuNPs on the lipid bilayer is more hindered than in the negative control, which suggests that these two species do not exhibit sufficient self-interactions to form islands. This observation corresponds to a significantly lower average anomalous diffusion exponent and larger average deflection angles for 4Man[6Glc]Glc **25** and 4Glc[6Man]Glc **26** than in case of the negative control on shorter time scales below 1 ms (Figure 5(b),(c)).

**Figure 5.**
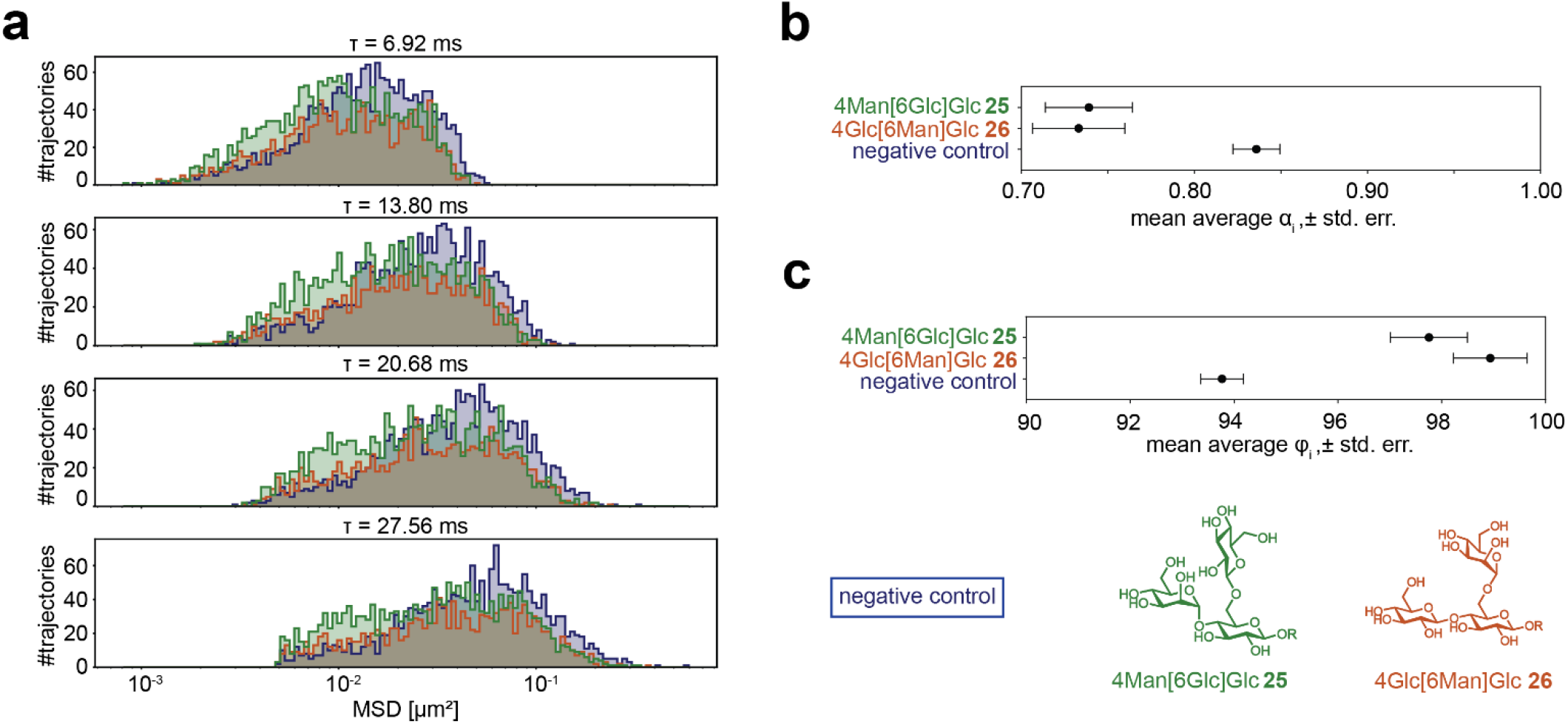
Effect of regiochemistry on system dynamics. Two branched model glycans were compared with the negative control on both a global and local level. (a) MSD histograms for 4Man[6Glc]Glc **25** and 4Glc[6Man]Glc **26** (green, orange, respectively; negative control: blue) (b) Local anomalous diffusion exponents. (c) Local deflection angles.

**Figure 6.**
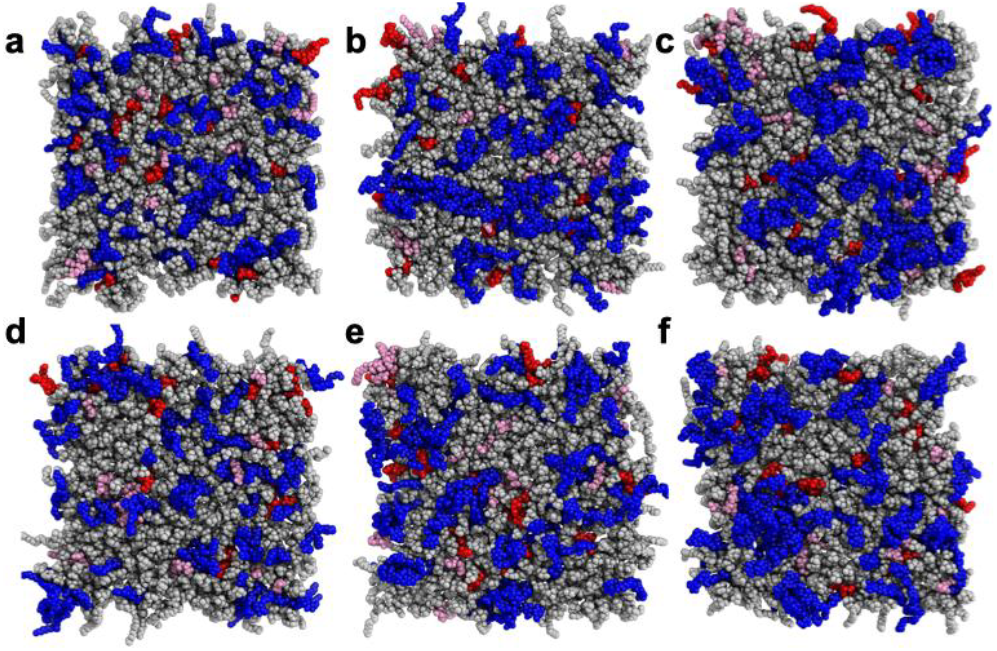
Top view of membrane leaflet after 25 μs of MD simulation. The top row (a-c) represents systems with Na+ counterions, while the bottom row (d-f) corresponds to systems with Mg2+ counterions. Panels (a) and (d) include Glc 13 glycolipids, (b) and (e) feature αGlc2 15 glycolipids, and (c) and (f) show αGlc3 16 glycolipids. Lipids are colorcoded as follows: DOPC in gray, DOPS in pink, DOPE in red, and glycolipids in blue. Visualization created using PyMOL.^39^

## Section 5: Computational modeling

To complement these analyses, we implemented all-atom molecular dynamics (MD) simulations corresponding to the systems studied in the iSCAT-based experiments. These MD simulations covered time windows of 25 μs (Figure 10). The systems simulated included glycolipids with azido-functionalized headgroups of varying sizes – Glc **13**, αGlc_2_ **15**, or αGlc_3_ **16** – and were solvated with either monovalent (Na+) or divalent (Mg^2+^) counterions (details on parameterization are provided in the SI chapter 6).

The simulations revealed glycolipid clustering, which increased with the size of the sugar headgroup (Figure 7). This trend was consistent across systems with both Na+ (Figure 7, upper panel) and Mg^2+^ counterions (Figure 7 lower panel).

**Figure 7.**
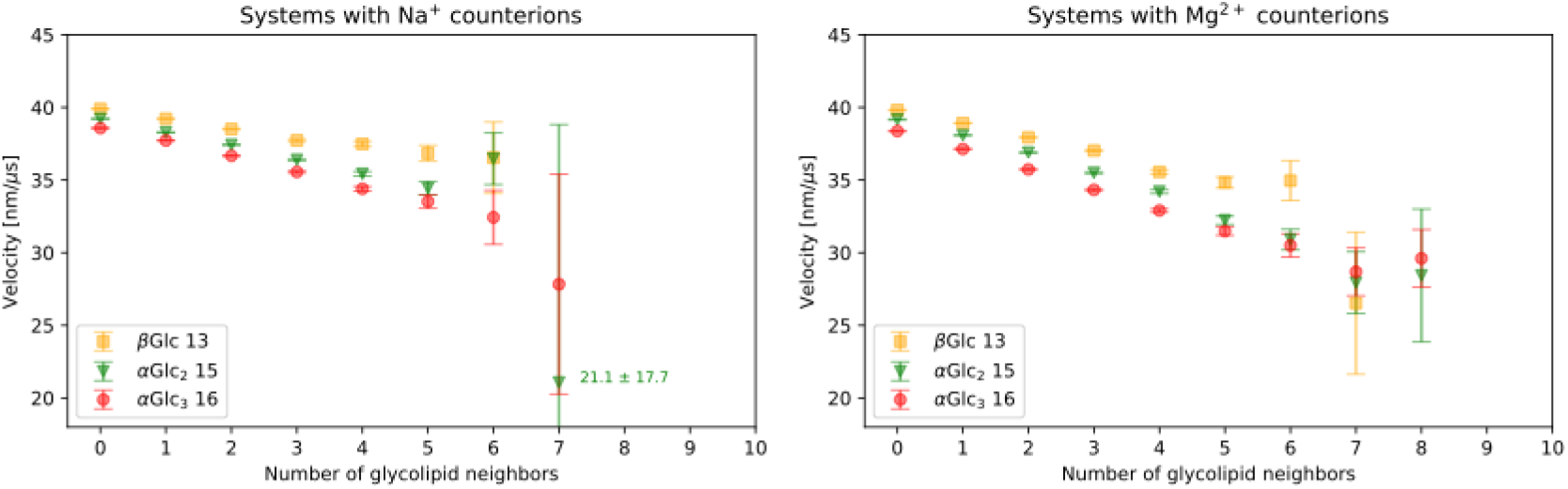
Diffusion velocity of DOPC lipids as a function of the number of neighboring glycolipids, calculated using a time step of *τ* = 20 ns. The number of glycolipid neighbors is defined as the number of glycolipids within a 1.2 nm cutoff distance from the phosphate group of a DOPC lipid. The glycolipid type and counterion for each system are indicated in the legend. Error bars represent the standard error of the mean.

Clustering of glycolipids significantly reduced the diffusion velocity of neighboring (DOPC) phospholipids in a manner dependent on the glycolipid headgroup. In particular, systems containing triglucosides **16** and **23** exhibited a more substantial reduction in the diffusion velocity of phospholipids embedded within clusters compared to systems glucoside **13** headgroups (Figure 7). This effect intensified with the number of neighboring glycolipids, as the increased drag exerted by these clusters further impeded lipid mobility.

The effect of sugar headgroup-driven glycolipid clustering on the lipid dynamics was further addressed in Brownian dynamics (BD) simulations on the ms-timescale. Our results for 2-dimensional lipid systems containing domains of different size (SI, Figure S93) suggest that larger domains result in MSD distributions shifted to larger values for the MSD (SI, Figure S94).

## Section 6: Conclusion and outlook

Studying dynamics of complex molecular system, such as the glycocalyx, is a nascent topic extending well past the limit of current technology. Here, we leverage a combination of several fields, in particular organic synthesis, advanced microscopy, statistical analysis and computational modeling, to address these challenges. With this integrated approach, we establish new insights into the complex functional relationships that govern glycocalyx organization. Due to the high spatiotemporal resolution, iSCAT microscopy presents a suitable tool for tracking and studying the diffusion dynamics of glycosylated membranes. Relying on a large number of trajectories, measuring different glycosylation conditions in triplicates, and developing an efficient data processing pipeline, we have found significant differences of characteristic diffusion metrics such as mean squared displacement or local directional correlation coefficients. In selected cases, strong glycan-specific correlation between the systems structure and its dynamics has been observed. The performed computational simulations of the system show the clustering of surface bound glycans, substantiating our experimental findings.

Fundamentally, our method of a bottom-up construction of defined glycosylated lipid bilayers lends itself to a more extensive investigation of other glycans and their mutual interactions. Moreover, due to the possibility of tracking in complex environments based on iSCAT, controlled addition of AuNPs^40^ and the biocompatibility of copper-free click chemistry,^41^ live-cell measurements are expected to be feasible in the near future. Further research in this regard could recreate specific glycocalyces found in physiological and diseased cell states as well as study glycan-protein interactions via external addition of known or putative glycan binders.

In summary, our results provide first answers to one of the oldest questions in the field of glycobiology: How does glycan structure functionally affect glycan behavior in the supramolecular environment of the lipid membrane? Our approach has the potential to reveal novel insights into the roles of glycans in biological processes, and we anticipate that these findings will have significant implications for the development of new clinical tools, for the establishment of impactful biotechnological applications, and for a deeper understanding of fundamental cell biology.

## ASSOCIATED CONTENT

Supporting information, detailing experimental and analysis procedures. This material is available free of charge via the Internet at http://pubs.acs.org.

## AUTHOR INFORMATION

### Author Contributions

CMB performed data evaluation with assistance by LM. LP, IvS and SOJ performed organic synthesis with assistance by TKL. IvS and MR performed iSCAT sample preparation and measurements with assistance from RT, MB, CH, KA, and VS. CP, MT, and RAB performed computational simulations. RAB, TKL, and LM supervised the project. TKL and LM conceived the project. CMB, LP, IvS, MR, RAB, TKL, and LM wrote the paper. All authors read and approved the manuscript. ‡ Co-first authors CMB, LP and IvS (listed alphabetically) contributed equally to this manuscript, and each has the right to list themselves first in author order on their CVs.

