## Supplemental Information for "Bottom-up investigation of spatiotemporal glycocalyx dynamics with interferometric scattering microscopy"

### Electronic Supplementary Information (ESI)

for

#### 1. General Synthetic Methods

Moisture-sensitive reactions were carried out in oven-dried glassware under a positive pressure of nitrogen. Analytical thin layer chromatography (TLC) was performed on silica gel plates (Alugram® Xtra Sil G/$UV_{254}$). Visualization was achieved by UV light and/or with 10 % sulfuric acid in ethanol or vanillin stain (900 mg vanillin, 12 mL $H_2SO_4$, 90 mL $H_2O$, 90 mL EtOH) or ninhydrin stain (200 mg ninhydrin, 1 mL acidic acid, 200 mL acetone), followed by heat treatment at approx. 200 °C. The products were purified by flash chromatography on silica gel columns (Merck, 230–400 mesh, particle size 0.040–0.063 mm) or by automated flash chromatography using a puriFlash450 or puriFlash5.020 device from Interchim®. For automated flash chromatography prepacked columns (particle size 30 µm) were used manufactured by Interchim®. For automated reversed phase flash chromatography prepacked columns (C18AQ, particle size 15 µm) were used manufactured by Interchim®. MeOH and DMF were dried over molecular sieves (3Å) under a nitrogen atmosphere. $CH_2Cl_2$ was dried over aluminum oxide columns in a PureSolv MD5 solvent purification system from Inert Corporation®. Dry 1,2-dichloroethane was purchased from Acros. Optical rotations were measured on an Anton Paar MCP5100 polarimeter at 20 °C with a sodium D-line (589 nm) and a cuvette of 10 cm path length in the solvents indicated. Proton ($^{1}H$) nuclear magnetic resonance spectra and carbon ($^{13}C$) nuclear magnetic resonance spectra were recorded on a Bruker AvanceNeo 500 and Bruker Avance 600 spectrometer at 298 K. Chemical shifts are referenced to the internal standard tetramethylsilane (TMS) or to the residual proton of the NMR solvent. Multiplets (multiplicity s=singlet, d=doublet, t=triplet, q=quartet, m=multiplet, br=broad) are listed according to chemical shift, coupling constants are given in Hertz (Hz). Full assignment of the signals was achieved by using 2D NMR techniques ($^{1}H$-$^{1}H$ COSY, $^{1}H$-$^{1}H$ TOCSY, $^{1}H$-$^{1}H$ NOESY, $^{1}H$-$^{13}C$ HMBC and $^{1}H$-$^{13}C$ HSQC). The first carbohydrate unit at the downstream end of an oligosaccharide is named Glc or Xyl respectively, the second carbohydrate unit Glc' or Xyl', the third Glc'' or Xyl''. Infrared (IR) spectra were measured on a PerkinElmer FT-IR Paragon 1000 (ATR) spectrometer and are reported in $cm^{-1}$. HR-ESI mass spectra were recorded on a ThermoFischer Orbitrap using MeCN, MeOH, $CH_2Cl_2$ or a solvent mixture of MeCN/$H_2O$-80/20 with 1.3 mM ammonium formate additive as a solvent.

#### 2. Synthesis of Oligosaccharides

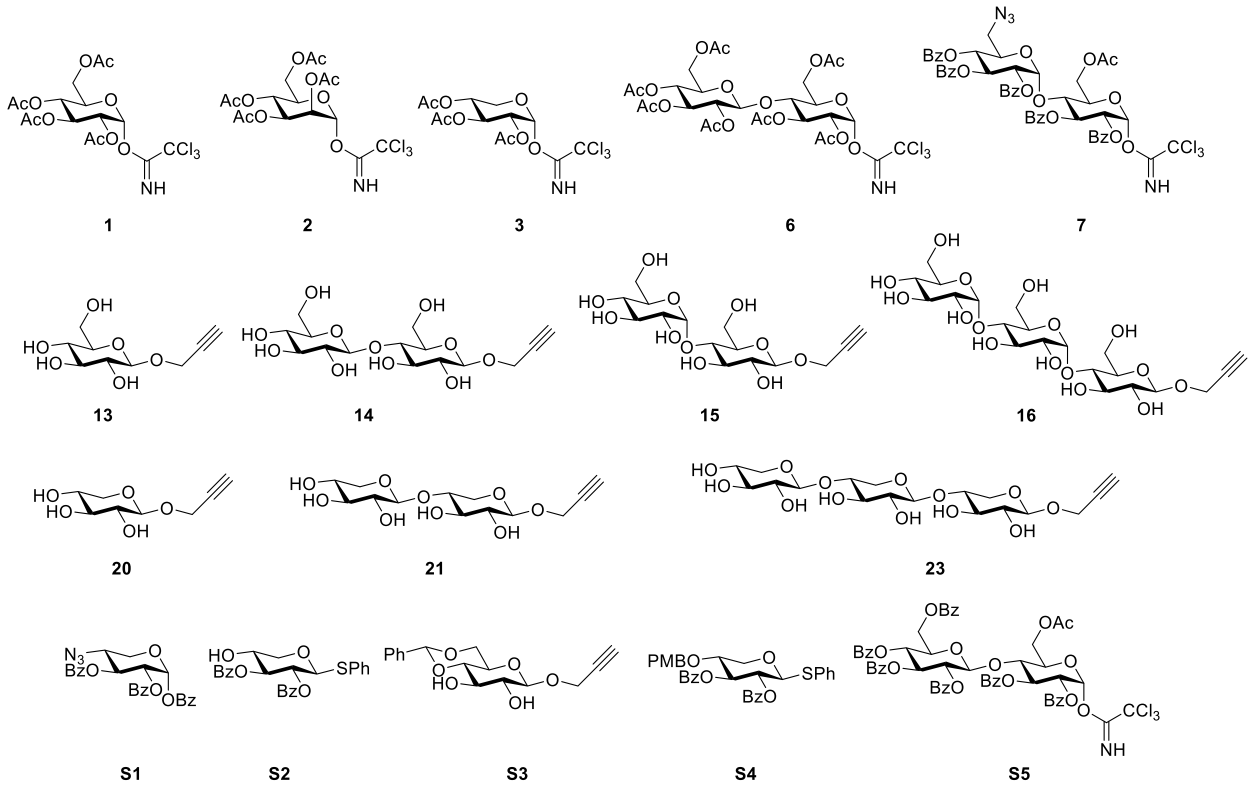

**Figure S1.** Literature-known saccharides. The saccharides **1**,[1] **2**,[2] **3**,[3] **6**,[4] **7**,[5] **13**,[6] **14**,[7] **15**,[8] **16**,[7] **20**[9] (cf. Scheme 1), **S1**,[10] **S2**,[11] **S3**,[12,13] **S4**,[14] **S5**[15] were synthesized according to literature procedures. For the synthesis of the xylosides **21** and **23**, an enzymatic route is published.[16] Here, we report the chemical synthesis of **21** and **23** (vide infra). Other than reported in the literature, the propargyl glycosides **13-16**, **20**, **21**, and **23** were additionally purified on reversed phase silica gel and obtained as lyophilisates. **20** was used as an anomeric mixture of α:β, 9:91. The analytical data of all literature-known saccharides are in full agreement with those reported.

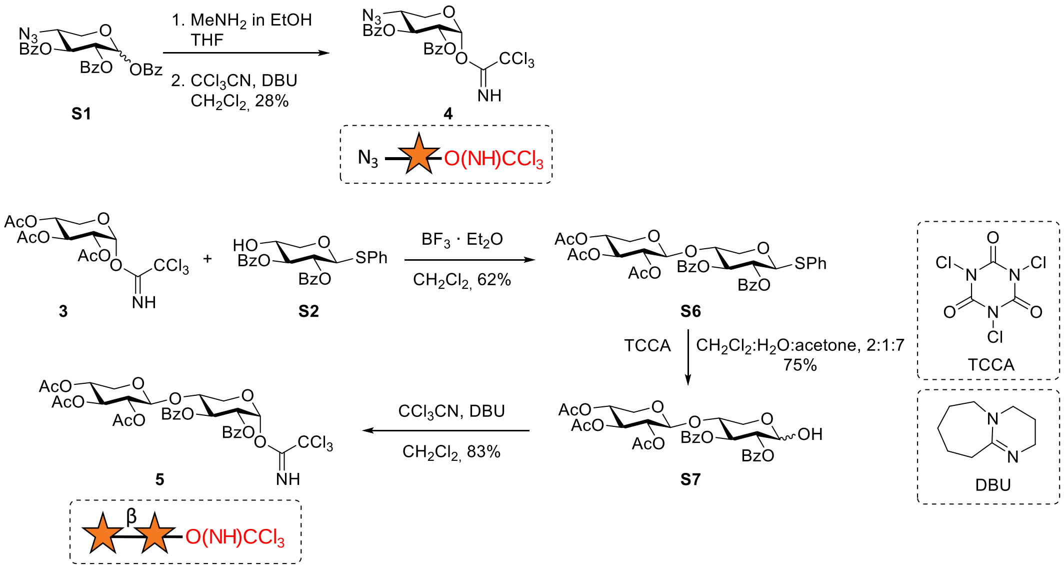

**Scheme S1.** Synthesis of trichloroacetimidates **4** and **5** (cf. Scheme 1). TCCA: trichloroisocyanuric acid; DBU: 1,8-diazabicyclo[5.4.0]undec-7-en.

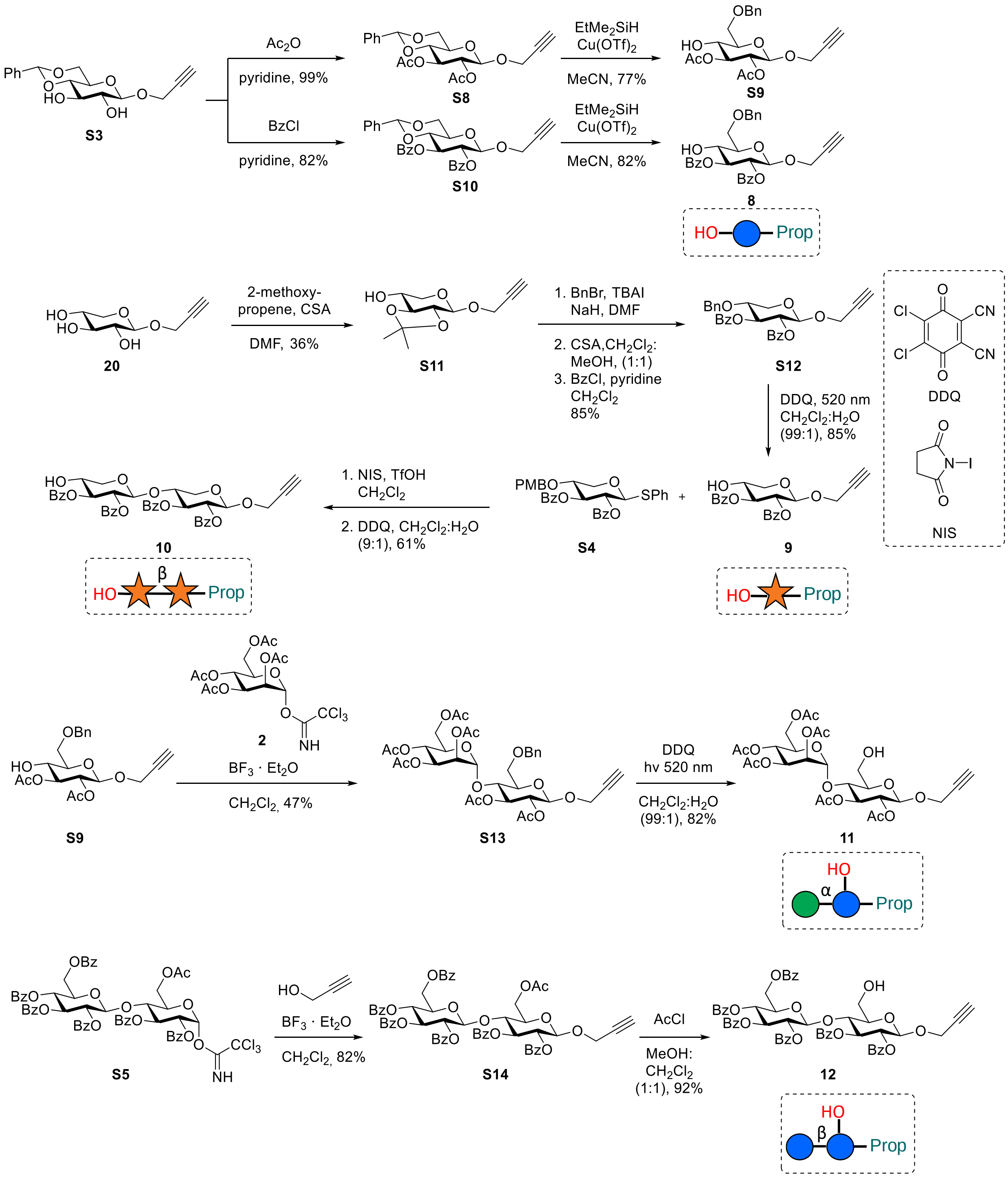

**Scheme S2.** Synthesis of acceptor propargyl glycosides **S9**, **8**, **10**, **11** and **12** (cf. Scheme 1). CSA: (±)-camphorsulfonic acid; TBAI: tetra-*n*-butylammonium iodide; DDQ: 2,3-dichloro-5,6-dicyano-1,4-benzoquinone; NIS: *N*-iodosuccinimide.

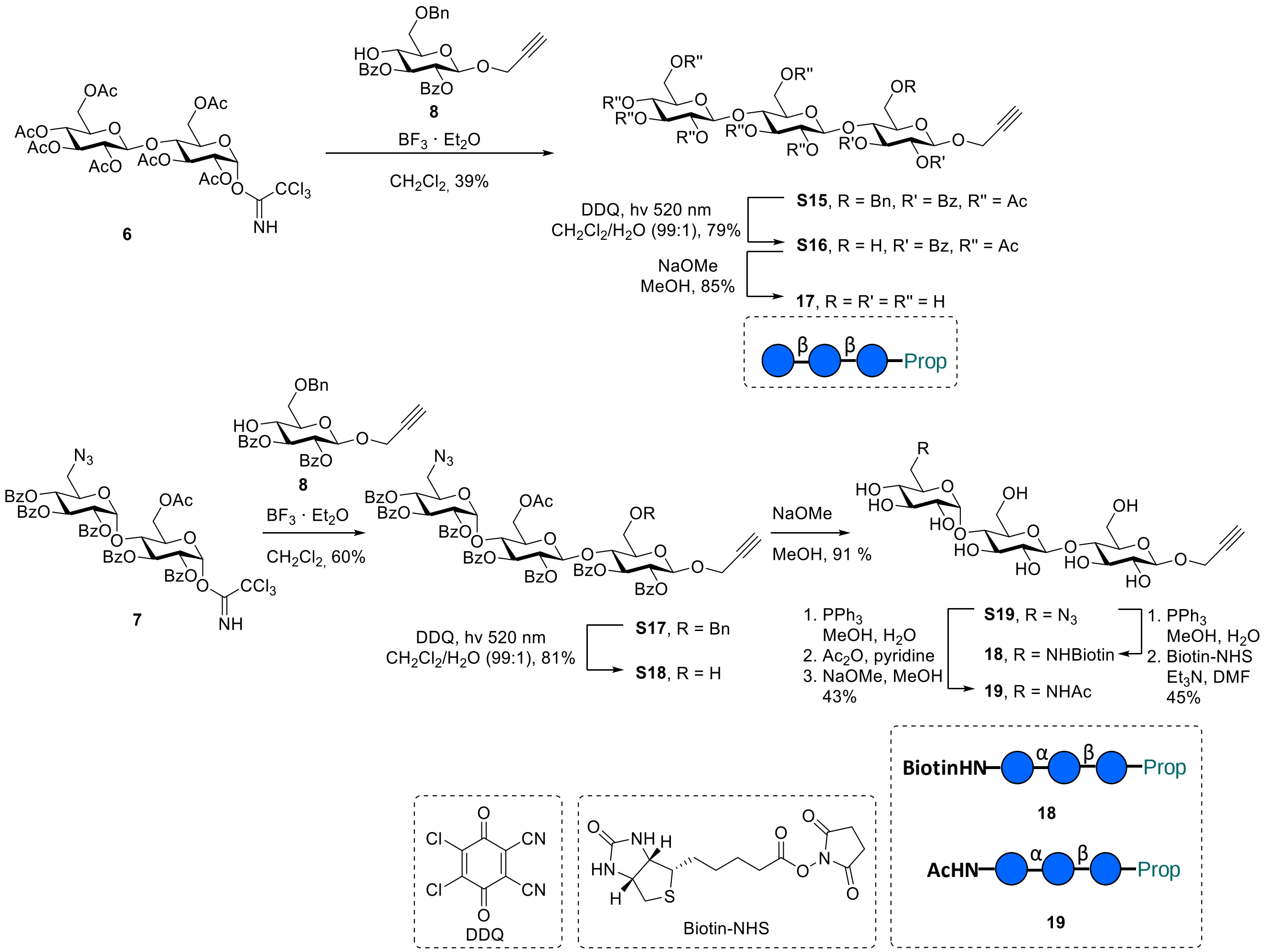

**Scheme S3.** Synthesis of propargyl glycosides **17**, **18** and **19** (cf. Scheme 1). DDQ: 2,3-dichloro-5,6-dicyano-1,4-benzoquinone; Biotin-NHS: biotin-*N*-hydroxysuccinimide ester.

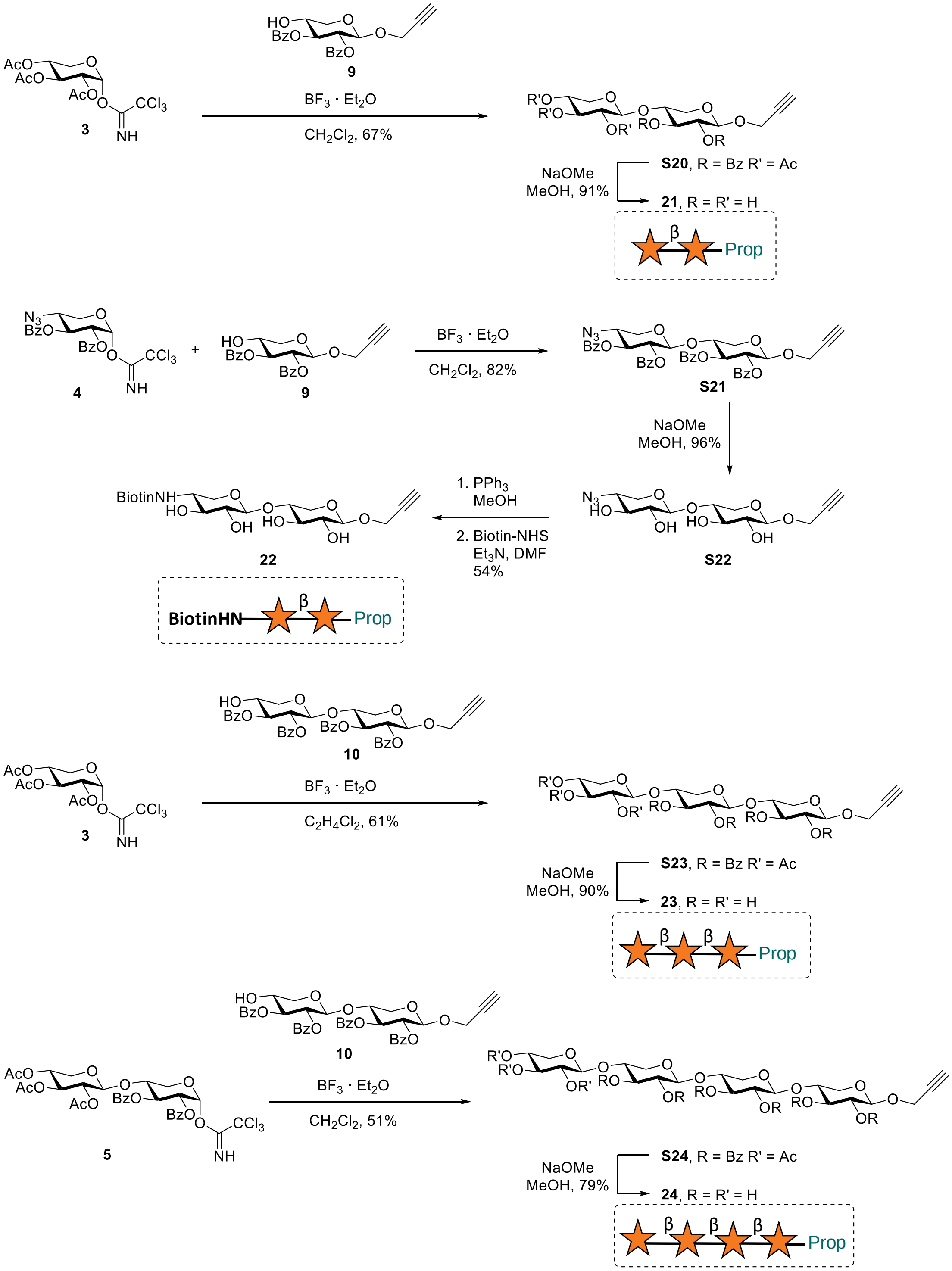

**Scheme S4.** Synthesis of propargyl glycosides **21**, **22**, **23** and **24** (cf. Scheme 1). Biotin-NHS: biotin-*N*-hydroxysuccinimide ester.

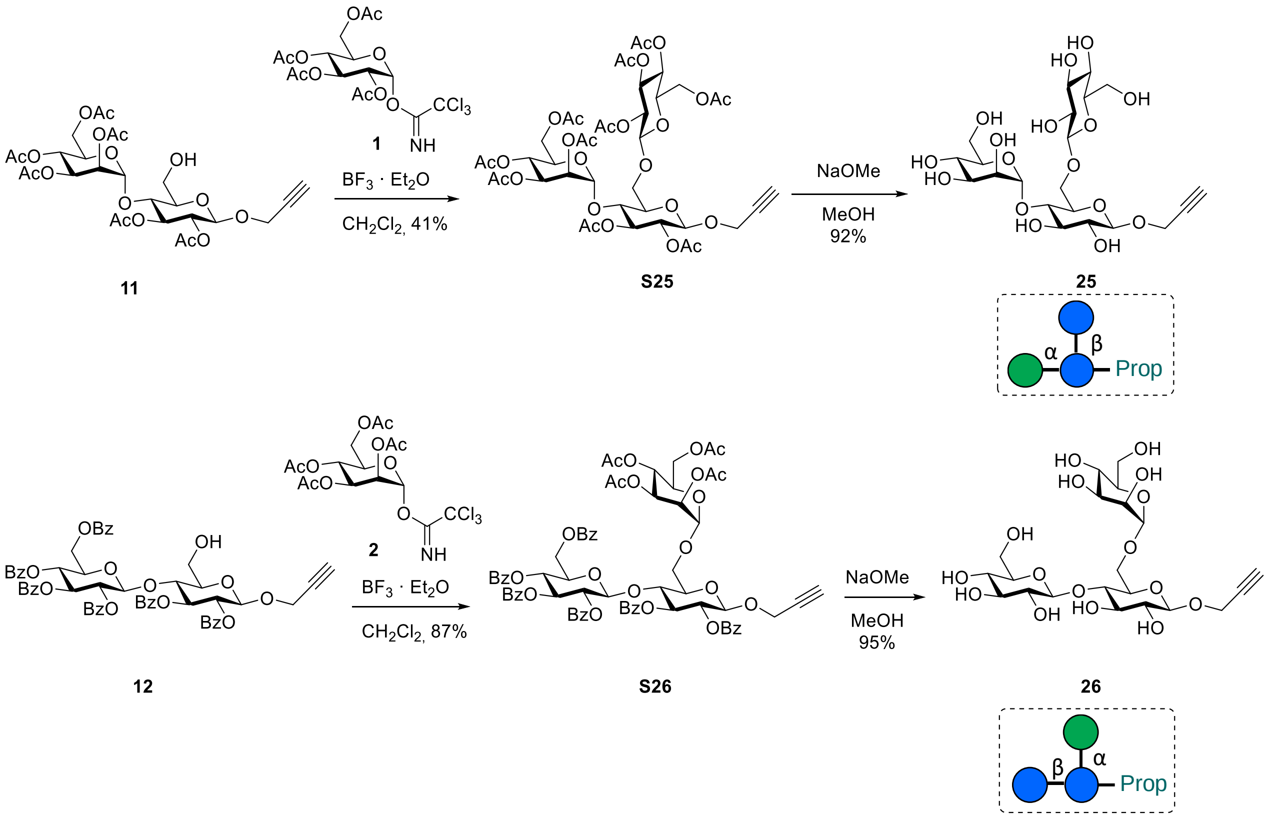

**Scheme S5.** Synthesis of propargyl glycosides **25** and **26** (cf. Scheme 1).

#### 2.1. *O*-(4-Azido-2,3-di-*O*-benzoyl-4-deoxy-α-D-xylopyranosyl) trichloroacetimidate (4)

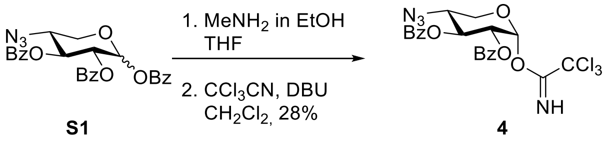

The xylose derivative **S1**[10] (600 mg, 1.23 mmol) was dissolved in dry THF (8.6 mL) at room temperature and methylamine in ethanol (33%, 1.03 mL) was added dropwise under a nitrogen atmosphere. After 45 min at room temperature the mixture was diluted with ethyl acetate. The organic phase (80 mL) was washed with brine (4 x 10 mL). The organic phase was dried over $MgSO_4$, filtered and the solvent removed under reduced pressure to a volume of 3 mL, then toluene (5 mL) was added and it was further concentrated. The crude product was purified on silica gel via automated flash chromatography (cyclohexane:ethyl acetate, 90:10→60:40) to obtain the reducing sugar intermediate (cyclohexane:ethyl acetate, 3:2, $R_F$ = 0.59). In the next reaction step, the hemiacetal was dissolved in dry $CH_2Cl_2$ (4.0 mL) under a nitrogen atmosphere, it was cooled to 0°C and DBU (16 µL, 107 µmol) was added dropwise followed by trichloroacetonitrile (290 µL, 2.89 mmol). The reaction mixture was stirred for 1 h at 0 °C min until TLC indicated full conversion of the starting material. The mixture was diluted with $CH_2Cl_2$ (40 mL) and washed with brine (10 mL). The phases were separated and the aq. phase was extracted with $CH_2Cl_2$ (10 mL). All organic phases were combined and dried over $MgSO_4$, filtrated and the solvent was removed under reduced pressure. The crude product was purified on silica gel via automated flash chromatography (cyclohexane:ethyl acetate, 95:5→60:40) to obtain the title compound as a colorless foam (181 mg, 343 µmol, 28%).

***R*<sub></sub>**$_F$ (cyclohexane:ethyl acetate, 3:2) = 0.73.

$[\alpha]_D^{20}$ = +135.5 ($c$ = 0.60, $CHCl_3$).

**IR (ATR):** $\tilde{\nu}$ = 3340, 3063, 2952, 2108, 1727, 1675, 1256, 705 $cm^{-1}$.

**$^1$H NMR** (500 MHz, $CDCl_3$, 298 K): δ = 8.61 (s, 1H, N<u>H</u>), 8.02-7.98 (m, 2H, $OBz_{ortho}$), 7.57-7.48 (m, 2H, 2 x $OBz_{para}$), 7.43-7.38 (m, 2H, $OBz_{meta}$), 7.38-7.33 (m, 2H, $OBz_{meta}$), 6.65 (d, $^3J_{1,2}$ = 3.6 Hz, 1H, H-1), 5.98 (t, $^3J_{2,3}$ = $^3J_{3,4}$ =9.9 Hz, 1H, H-3), 5.43 (dd, $^3J_{2,3}$ = 10.1 Hz, $^3J_{1,2}$ = 3.6 Hz, 1H, H-2), 4.08 (dd, $^2J_{5eq,5ax}$ = 10.8 Hz, $^3J_{4,5eq}$ = 5.4 Hz, 1H, H-$5_{eq}$), 4.05-3.99 (m, 1H, H-4), 3.95-3.90 (m, 1H, H-$5_{ax}$) ppm.

**$^{13}$C NMR** (126 MHz, $CDCl_3$, 298 K): δ = 165.5 (2 x $C{=}O_{OBz}$), 160.7 (<u>C</u>=NH), 133.61 ($OBz_{para}$), 133.56 ($OBz_{para}$), 129.9 ($OBz_{ortho}$), 129.8 ($OBz_{ortho}$), 128.9 ($OBz_{ipso}$), 128.52 ($OBz_{meta}$), 128.46 ($OBz_{ipso}$, $OBz_{meta}$), 93.5 (C-1), 90.7 (<u>C</u>$Cl_3$, assigned by HMBC), 70.7 (C-3), 70.6 (C-2), 62.1 (C-5), 59.3 (C-4) ppm.

**ESI-HRMS:** $m/z$ = 549.01017 $[M+Na]^+$ (calculated $m/z$ = 549.01059 for $[M+Na]^+$).

##### 2.2. *O*-(2,3,4-Tri-*O*-acetyl-β-D-xylopyranosyl-(1→4)-2,3-di-*O*-benzoyl-α-D-xylopyranosyl) trichloroacetimidate (5)

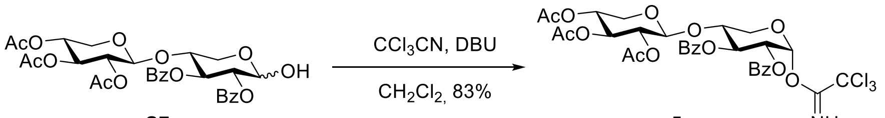

S7 5

The xylobiose **S7** (280 mg, 454 µmol) was dissolved in dry $CH_2Cl_2$ (4.0 mL) under a nitrogen atmosphere, it was cooled to 0°C and DBU (15 µL, 100 µmol) was added dropwise followed by trichloroacetonitrile (250 µL, 2.49 mmol). The reaction mixture was stirred for 1 h at 0 °C until TLC indicated full conversion of the starting material. The mixture was diluted with $CH_2Cl_2$ (40 mL) and washed with brine (10 mL). The phases were separated and the aq. phase was extracted with $CH_2Cl_2$ (10 mL). All organic phases were combined and dried over $MgSO_4$, filtrated and the solvent was removed under reduced pressure. The crude product was purified on silica gel via automated flash chromatography (cyclohexane:ethyl acetate, 9:1→6:4) to obtain the title compound as a colorless foam (288 mg, 378 µmol, 83%).

***R*<sub></sub>**$_F$ (cyclohexane:ethyl acetate, 1:1) = 0.56.

$[\alpha]_D^{20}$ = +33.8 ($c$ = 0.47, $CHCl_3$).

**IR (ATR):** $\tilde{\nu}$ = 3320, 2953, 1731, 1676, 1248, 1217, 1028, 708 $cm^{-1}$.

**$^1$H NMR** (500 MHz, Acetone-$d_6$, 298 K): δ = 9.36 (s, 1H, N<u>H</u>), 8.06-8.01 (m, 2H, $OBz_{ortho}$), 7.96-7.91 (m, 2H, $OBz_{ortho}$), 7.61-7.55 (m, 2H, 2 x $OBz_{para}$), 7.49-7.40 (m, 4H, 2 x $OBz_{meta}$), 6.70 (d, $^3J_{1,2}$ = 3.6 Hz, 1H, H-1), 5.97 (t, $^3J_{2,3}$ = $^3J_{3,4}$ = 9.8 Hz, 1H, H-3), 5.43 (dd, $^3J_{2,3}$ = 10.2 Hz, $^3J_{1,2}$ = 3.6 Hz, 1H, H-2), 5.08 (t, $^3J_{2,3}$ = $^3J_{3,4}$ = 8.5 Hz, 1H, H-3'), 4.91 (d, $^3J_{1,2}$ = 6.7 Hz, 1H, H-1'), 4.77 (dd, $^3J_{2,3}$ = 8.6 Hz, $^3J_{1,2}$ = 6.8 Hz, 1H, H-2'), 4.72-4.65 (m, H-4'), 4.47-4.38 (m, 1H, H-4), 4.14 (dd, $^2J_{5ax,5eq}$ =

11.4 Hz, $^3J_{4,5eq}$ = 5.9 Hz, 1H, H-5$_{eq}$), 3.90 (t, $^2J_{5ax,5eq}$ = $^3J_{4,5ax}$ = 11.2 Hz, 1H, H-5$_{ax}$), 3.67 (dd, $^2J_{5ax,5eq}$ = 11.8 Hz, $^3J_{4,5eq}$ = 5.1 Hz, 1H, H-5$_{eq}$'), 3.29 (dd, $^2J_{5ax,5eq}$ = 11.9 Hz, $^3J_{4,5ax}$ = 8.7 Hz, 1H, H-5$_{ax}$'), 2.00 (s, 3H, CH3$_{\text{OAc at C-2'}}$), 1.93 (s, 3H, CH3$_{\text{OAc at C-3'}}$), 1.92 (s, 3H, CH3$_{\text{OAc at C-4'}}$) ppm.

**$^{13}$C NMR** (126 MHz, Acetone-*d*$_6$, 298 K): δ = 170.2 (C=O$_{\text{OAc at C-3'}}$), 170.1 (C=O$_{\text{OAc at C-4'}}$), 169.7 (C=O$_{\text{OAc at C-2'}}$), 166.0 (C=O$_{\text{OBz at C-3}}$), 165.9 (C=O$_{\text{OBz at C-2}}$), 160.5 (C=NH), 134.5 (OBz$_{para}$), 134.0 (OBz$_{para}$), 131.0 (OBz$_{ipso}$), 130.5 (OBz$_{ortho}$), 130.4 (OBz$_{ortho}$), 129.8 (OBz$_{ipso}$), 129.5 (OBz$_{meta}$), 129.3 (OBz$_{meta}$), 101.4 (C-1'), 94.1 (C-1), 91.6 (CCl$_3$, assigned by HMBC),76.1 (C-4), 71.9 (C-2), 71.8 (C-3'), 71.6 (C-3), 71.4 (C-2'), 69.3 (C-4'), 62.7 (C-5), 62.4 (C-5'), 20.6 (CH3$_{OAc}$), 20.5 (2 x CH3$_{OAc}$) ppm.

**ESI-HRMS:** *m/z* = 794.05919 [M+Cl]$^-$ (calculated *m/z* = 794.05714 for [M+Cl]$^-$).

##### 2.3. Propargyl 2,3-di-*O*-benzoyl-6-*O*-benzyl-β-D-glucopyranoside (8)

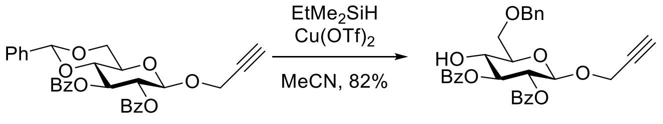

**S10** **8**

The benzylidene-protected glucoside **S10** (2.04 g, 3.96 mmol, 1 equiv) was dissolved in dry acetonitrile (40 mL) under nitrogen atmosphere. Ethyldimethylsilane (1.05 mL, 7.93 mmol, 2 equiv) was added and the reaction mixture was cooled to 0°C. A solution of freshly activated (vacuum, ~200 °C, 15 min) copper(II)triflate (14.4 mg, 39.7 µmol, 0.01 equiv) in acetonitrile (5 mL) was added and stirred at 0°C for 2 h. Ethyl acetate (25 mL) was added and the reaction mixture was vigorously stirred under air atmosphere for 2h. The reaction mixture was diluted with ethyl acetate (300 mL) and washed with satd. aq. $NaHCO_3$ solution (50 mL), then washed with brine (50 mL), dried over $MgSO_4$, filtrated and the solvent was removed under reduced pressure. The crude was purified on silica gel via automated flash chromatography (cyclohexane:ethyl acetate, 8:2) to yield the product as a colorless foam (1.67 g, 3.23 mmol, 82%).

***R*$_F$** (cyclohexane:ethyl acetate, 8:2) = 0.22.

$[\alpha]_D^{20}$ = +46.6 (*c* = 1.21 in $CHCl_3$).

**IR (ATR):** $\tilde{\nu}$ = 3448 (br, w), 3296 (w), 2871 (w), 1724 (s), 1602 (w), 1451 (m), 1264 (s), 1093 (s), 1067 (s), 989 (m), 707 (s) cm$^{-1}$.

**$^1$H NMR** (600 MHz, $CDCl_3$, 298 K) δ = 8.00 – 7.95 (m, 4H, 2 x OBz$_{ortho}$), 7.56 – 7.47 (m, 2H, 2 x OBz$_{para}$), 7.41 – 7.28 (m, 9H, 2 x OBz$_{meta}$, OBn), 5.50 – 5.41 (m, 2H, H-2, H-3), 4.98 (d, $^3J_{1,2}$ = 7.6 Hz, 1H, H-1), 4.69 – 4.58 (m, 2H, PhC<u>H</u>$_2$), 4.46 – 4.33 (m, 2H, C<u>H</u>$_2$C≡CH), 3.99 (td, $^3J_{3,4}$ = $^3J_{4,5}$ = 9.2 Hz, $^3J_{4,OH}$ = 3.4 Hz, 1H, H-4), 3.86 (d, $^3J_{5,6}$ = 4.5 Hz, 2H, H-6a, H-6b), 3.72 (dt, $^3J_{4,5}$ = 9.3 Hz, $^3J_{5,6}$ = 4.5 Hz, 1H, H-5), 3.23 (d, $^3J_{4,OH}$ = 3.4 Hz, 1H, OH-4), 2.38 (t, $^4J_{C≡CH,CH2}$ = 2.4 Hz, 1H, C≡C<u>H</u>) ppm.

**$^{13}$C NMR** (151 MHz, $CDCl_3$, 298 K) δ = 167.3 (Ph<u>C</u>=O), 165.4 (Ph<u>C</u>=O), 137.6 (OBn$_{ipso}$), 133.5 (OBz$_{para}$), 133.1 (OBz$_{para}$), 130.0 (OBz), 129.9 (OBz), 129.5 (OBz$_{ipso}$), 129.0 (OBz$_{ipso}$), 128.5 (OBz), 128.4 (OBz), 128.3 (OBn), 127.9 (OBn$_{para}$), 127.8 (OBn), 98.4 (C-1), 78.4 (CH$_2$<u>C</u>≡CH), 76.7 (C-2/C-3), 75.3 (CH$_2$C≡<u>C</u>H), 74.8 (C-5), 73.8 (Ph<u>C</u>H$_2$), 71.1 (C-2/C-3), 70.9 (C-4), 69.8 (C-6), 55.9 (<u>C</u>H$_2$C≡CH) ppm.

**ESI-HRMS:** *m/z* = 534.21143 $[M+NH_4]^+$ (calculated *m/z* = 534.21224).

##### 2.4. Propargyl 2,3-di-*O*-benzoyl-β-D-xylopyranoside (9)

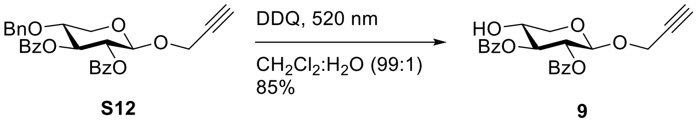

**S12** **9**

Following an adapted procedure from Cavedon[17] et al., the xyloside **S12** (1.46 g, 3.00 mmol) and 2,3-dichloro-5,6-dicyano-1,4-benzoquinone (1.02 g, 4.49 mmol) were dissolved in dry $CH_2Cl_2$ (150 mL) and $H_2O$ (1.5 mL) under a nitrogen atmosphere. The reaction mixture was stirred at room temperature and irradiated with 520 nm. The reaction mixture was cooled with a CPU fan adjusted next to the reaction flask. After 4 h thin layer chromatography indicated full conversion of the starting material and the reaction was diluted with $CH_2Cl_2$ (100 mL) and the organic phase was washed with a satd. aq. $NaHCO_3$ solution (4 x 50 mL). The combined aq. phases were extracted with $CH_2Cl_2$ (50 mL). The combined organic phases were dried over $MgSO_4$, it was filtered and the solvent was removed under reduced pressure. The crude product was loaded onto celite® and purified via automated flash chromatography on silica (cyclohexane:ethyl acetate, 90:10→65:35) to obtain the product as a colorless amorphous solid (1.01 g, 2.55 mmol, 85%).

$\boldsymbol{R}_{\mathrm{F}}$ (cyclohexane:ethyl acetate, 3:2) = 0.48.

$[\boldsymbol{\alpha}]_{\mathbf{D}}^{\mathbf{20}}$ = +64.3 (*c* = 1.07, $CHCl_3$).

**IR (ATR):** $\tilde{\nu}$ = 3453, 3293, 2869, 2121, 1721, 1262, 1068, 1026, 707 $cm^{-1}$.

**$^1$H NMR** (500 MHz, $CDCl_3$, 298 K): δ = 8.03-7.98 (m, 4H, 2 x $OBz_{ortho}$), 7.57-7.51 (m, 2H, 2 x $OBz_{para}$), 7.43-7.36 (m, 4H, 2 x $OBz_{meta}$), 5.38 (dd, $^3J_{2,3}$ = $^3J_{3,4}$ = 7.9 Hz, $^3J_{1,2}$ = 6.1 Hz, 1H, H-2), 5.32-5.28 (m, 1H, H-3), 4.98 (d, $^3J_{1,2}$ = 6.0 Hz, 1H, H-1), 4.43-4.34 (m, 2H, C<u>H</u>$_2$C≡CH), 4.24 (dd, $^2J_{5ax,5eq}$ = 12.0 Hz, $^3J_{4,5eq}$ = 4.5 Hz, 1H, H-$5_{eq}$), 4.05-3.99 (m, 1H, H-4), 3.58 (dd, $^2J_{5ax,5eq}$ = 12.1 Hz, $^3J_{4,5ax}$ = 7.8 Hz, 1H, H-$5_{ax}$), 3.10 (d, $^3J_{4,OH}$ = 5.8 Hz, 1H, O<u>H</u>), 2.43 (t, $^4J_{CH2,C≡CH}$ = 2.4 Hz, 1H, C≡C<u>H</u>) ppm.

**$^{13}$C NMR** (126 MHz, $CDCl_3$, 298 K): δ = 167.2 (C=$O_{OBz\ at\ C\text{-}3}$), 165.2 (C=$O_{OBz\ at\ C\text{-}2}$), 133.7 ($OBz_{para}$),133.4 ($OBz_{para}$), 130.1 ($OBz_{ortho}$), 129.9 ($OBz_{ortho}$), 129.3 ($OBz_{ipso}$), 128.9 ($OBz_{ipso}$), 128.5 ($OBz_{meta}$), 128.4 ($OBz_{meta}$), 98.2 (C-1), 78.4 (<u>C</u>≡CH), 75.5 (C-3), 75.2 (C≡<u>C</u>H), 70.2 (C-2), 68.6 (C-4), 64.6 (C-5), 55.7 (<u>C</u>$H_2$C≡CH) ppm.

**ESI-HRMS:** *m/z* = 419.10936 $[M+Na]^+$ (calculated *m/z* = 419.11012 for $[M+Na]^+$).

##### 2.5. Propargyl 2,3-di-*O*-benzoyl-β-D-xylopyranosyl-(1→4)-2,3-di-*O*-benzoyl-β-D-xylopyranoside (10)

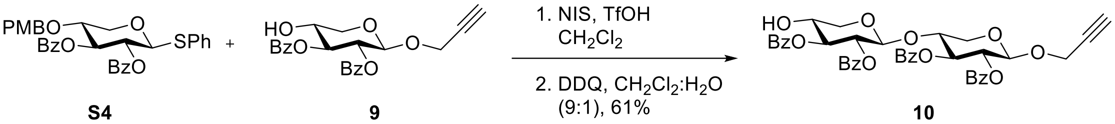

**S4** **9** **10**

Following an adapted procedure from Underlin[18] et al., the thioglycoside-donor **S4**[14] (110 mg, 193 µmol) and acceptor xyloside **9** (91.6 mg, 231 µmol) were dissolved in dry $CH_2Cl_2$ (3.3 mL) under a nitrogen atmosphere, freshly activated powdered 4Å molecular sieves was added and the reaction mixture stirred at room temperature for 15 min. Then, it was cooled to -40 °C and *N*-iodosuccinimide (51.7 mg, 230 µmol) and trifluoromethanesulfonic acid (5 µL, 57.0 µmol) were consecutively added under stirring. The mixture turned red and after 5 min at -40 °C thin layer chromatography analysis indicated full conversion of the donor. The reaction mixture was quenched with triethylamine (80 µL, 574 µmol) and a concentrated aq. sodium thiosulfate solution (3 mL) was added. The yellow mixture was stirred at room temperature for 10 min. The colorless mixture was diluted with $CH_2Cl_2$, filtered and the organic phase was separated and washed with brine. Afterwards the organic phase was dried over $MgSO_4$, filtered and concentrated in vacuo. After purification via automated flash chromatography on silica (cyclohexane:ethyl acetate, 9:1→6:4) the protected disaccharide intermediate (124 mg) was obtained. The intermediate (121 mg) was dissolved in $CH_2Cl_2$ (6.0 mL) and $H_2O$ (0.6 mL) and 2,3-dichloro-5,6-dicyano-1,4-benzoquinone (48.0 mg, 212 µmol) was added. The mixture was stirred in the dark at room temperature for 14 h. Then, the reaction mixture was diluted with $CH_2Cl_2$ and quenched with a satd. aq. $NaHCO_3$ solution (1 mL). The organic phase was washed with a satd. aq. $NaHCO_3$ solution (2 x 20 mL). The organic phase was dried over $MgSO_4$, filtered and the solvent was removed under reduced pressure. The crude product was purified via automated flash chromatography on silica (cyclohexane:ethyl acetate, 9:1→1:1) to obtain the product as a colorless amorphous solid (87.1 mg, 118 µmol, 61%).

***R*$_F$** (cyclohexane:ethyl acetate, 3:2) = 0.37.

$[\alpha]_D^{20}$ = +24.5 (*c* = 0.32, $CHCl_3$).

**IR (ATR):** $\tilde{\nu}$ = 3452, 3294, 2921, 2852, 2083, 1720,1451, 1259,1069 $cm^{-1}$.

**$^1$H NMR** (500 MHz, $CDCl_3$, 298 K): δ = 8.03-7.93 (m, 8H, 4 x $OBz_{ortho}$), 7.58-7.48 (m, 4H, 4 x $OBz_{para}$), 7.45-7.35 (m, 8H, 4 x $OBz_{meta}$), 5.65 (t, $^3J_{2,3}$ = $^3J_{3,4}$ = 8.1 Hz, 1H, H-3), 5.31-5.24 (m, 2H, H-2, H-2'), 4.90 (d, $^3J_{1,2}$ = 6.6 Hz, 1H, H-1), 4.79 (d, $^3J_{1,2}$ = 6.0 Hz, 1H, H-1'), 4.35-4.27 (m, 2H, C<u>H</u>$_2$C≡CH), 4.09-3.98 (m, 2H, H-4, H-$5_{eq}$), 3.78-3.68 (m, 2H, H-4', H-$5_{eq}$'), 3.48 (dd, $^2J_{5ax,5eq}$ = 12.0 Hz, $^3J_{4,5ax}$ = 8.5 Hz, 1H, H-$5_{ax}$), 3.19 (dd, $^2J_{5ax,5eq}$ = 11.8 Hz, $^3J_{4,5ax}$ = 7.4 Hz, 1H, H-$5_{ax}$'), 2.91 (d, $^3J_{4,OH}$ = 5.5 Hz, 1H, O<u>H</u>), 2.37 (t, $^4J_{CH2,C≡CH}$ = 2.4 Hz, 1H, C≡C<u>H</u>) ppm.

**$^{13}$C NMR** (126 MHz, $CDCl_3$, 298 K): δ = 167.3 (C=$O_{OBz\ at\ C-3'}$), 165.6 (C=$O_{OBz\ at\ C-3}$), 165.5 (C=$O_{OBz\ at\ C-2}$), 165.1 (C=$O_{OBz\ at\ C-2'}$), 133.8 ($OBz_{para}$), 133.7 ($OBz_{para}$), 133.3 (2 x $OBz_{para}$), 130.2 ($OBz_{ortho}$), 130.1 ($OBz_{ortho}$), 129.9 ($OBz_{ortho}$), 129.8 ($OBz_{ortho}$, $OBz_{ipso}$), 129.6 ($OBz_{ipso}$), 129.2 ($OBz_{ipso}$),128.9 ($OBz_{ipso}$), 128.73 ($OBz_{meta}$), 128.68 ($OBz_{meta}$), 128.5 ($OBz_{meta}$), 128.4 ($OBz_{meta}$), 101.1 (C-1'), 98.7 (C-1), 78.4 (<u>C</u>≡CH), 76.2 (C-4), 75.6 (C-3'), 75.4 (C≡<u>C</u>H), 72.3 (C-3), 71.0 (C-2), 70.8 (C-2'), 68.5 (C-4'), 64.5 (C-5'), 62.9 (C-5), 55.9 (<u>C</u>H$_2$C≡CH) ppm.

**ESI-HRMS:** *m/z* = 754.24780 $[M+NH_4]^+$ (calculated *m/z* = 754.24942 for $[M+NH_4]^+$).

#### 2.6. Propargyl 2,3,4,6-tetra-*O*-acetyl-α-D-mannopyranosyl-(1→4)-2,3-di-*O*-acetyl-β-D-glucopyranoside (11)

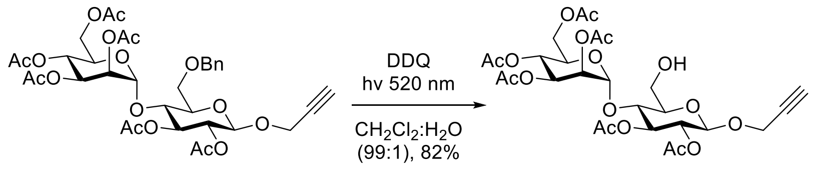

**S13** **11**

Following the procedure from Cavedon[17] et al., the disaccharide **S13** (821 mg, 1.14 mmol, 1 equiv) and 2,3-dichloro-5,6-dicyano-1,4-benzoquinone (387 mg, 1.70 mmol, 1.5 equiv) were dissolved in dry $CH_2Cl_2$ (50 mL) under nitrogen atmosphere and water (500 µL) was added. During strong stirring the reaction mixture was irradiated with 520 nm green light for 6.5 h at room temperature. The reaction mixture was diluted with $CH_2Cl_2$ (200 mL) and washed with satd. aq. $NaHCO_3$ solution (3 x 100 mL). The aq. phases were extracted with $CH_2Cl_2$ (50 mL). All organic phases were combined and dried over $MgSO_4$, filtrated and the solvent was removed under reduced pressure. The crude was purified on silica gel via automated flash chromatography (cyclohexane:ethyl acetate, 1:1) to yield the product as a colorless foam (588 mg, 930 µmol, 82%).

$\boldsymbol{R}_\mathbf{F}$ (cyclohexane:ethyl acetate, 1:1) = 0.24.

$[\boldsymbol{\alpha}]_\mathbf{D}^{\mathbf{20}}$ = -17.8 ($c$ = 0.52 in $CHCl_3$).

**IR (ATR):** $\tilde{\nu}$ = 3531 (w), 3278 (w), 2943 (w), 1741 (s), 1431 (w), 1368 (m), 1214 (s), 1135 (m), 1037 (s), 899 (m), 600 (m) $cm^{-1}$.

**$^1$H NMR** (600 MHz, $CDCl_3$, 298 K) δ = 5.30 (t, $^3J_{2,3}$ = $^3J_{3,4}$ = 9.5 Hz, 1H, H-$3_{Glc}$), 5.27 – 5.19 (m, 2H, H-$3_{Man}$, H-$4_{Man}$), 5.07 – 5.01 (m, 2H, H-$1_{Man}$, H-$2_{Man}$), 4.88 (dd, $^3J_{2,3}$ = 9.6 Hz, $^3J_{1,2}$ = 8.0 Hz, 1H, H-$2_{Glc}$), 4.79 (d, $^3J_{1,2}$ = 8.0 Hz, 1H, H-$1_{Glc}$), 4.42 – 4.32 (m, 2H, $C\underline{H}_2C{\equiv}CH$), 4.23 (dd, $^2J$ = 12.2 Hz, $^3J_{5,6a}$ = 6.5 Hz, 1H, H-$6a_{Man}$), 4.14 (dd, $^2J$ = 12.2 Hz, $^3J_{5,6b}$ = 2.5 Hz, 1H, H-$6b_{Man}$), 4.05 (ddd, $^3J_{4,5}$ = 9.1 Hz, $^3J_{5,6a}$ = 6.5 Hz, $^3J_{5,6b}$ = 2.5 Hz, 1H, H-$5_{Man}$), 3.98 (t, $^3J_{3,4}$ = $^3J_{4,5}$ = 9.5 Hz, 1H, H-$4_{Glc}$), 3.96 – 3.92 (m, 1H, H-$6a_{Glc}$), 3.89 – 3.83 (m, 1H, H-$6b_{Glc}$), 3.52 (ddd, $^3J_{4,5}$ = 9.7 Hz, $^3J_{5,6}$ = 3.9 Hz, $^3J_{5,6}$ = 2.1 Hz, 1H, H-$5_{Glc}$), 2.48 (t, $^4J_{C{\equiv}CH,CH2}$ = 2.4 Hz, 1H, $C{\equiv}C\underline{H}$), 2.13 (s, 3H, OAc), 2.12 (s, 3H, OAc), 2.08 (s, 3H, OAc), 2.05 (s, 6H, 2xOAc), 2.00 (s, 3H, OAc) ppm.

**$^{13}$C NMR** (151 MHz, $CDCl_3$, 298 K) δ = 170.6 ($CH_3\underline{C}O$), 170.3 ($CH_3\underline{C}O$), 169.8 ($CH_3\underline{C}O$), 169.7 ($CH_3\underline{C}O$), 169.7 ($CH_3\underline{C}O$), 169.6 ($CH_3\underline{C}O$), 99.2 (C-$1_{Man}$), 98.3 (C-$1_{Glc}$), 78.3 ($CH_2\underline{C}{\equiv}CH$), 75.7 (C-$4_{Glc}$), 75.5 ($CH_2C{\equiv}\underline{C}H$), 74.8 (C-$5_{Glc}$), 74.3 (C-$3_{Glc}$), 71.6 (C-$2_{Glc}$), 69.8 (C-$5_{Man}$), 69.7 (C-$2_{Man}$), 68.3 (C-$3_{Man}$/C-$4_{Man}$), 66.2 (C-$3_{Man}$/C-$4_{Man}$), 62.8 (C-$6_{Man}$), 61.2 (C-$6_{Glc}$), 56.2 ($\underline{C}H_2C{\equiv}CH$), 20.8 (2 x $CH_3$), 20.68 (2 x $CH_3$), 20.65 ($CH_3$), 20.62 ($CH_3$) ppm.

**ESI-HRMS:** $m/z$ = 650.22839 $[M+NH_4]^+$ (calculated $m/z$ = 650.22908).

##### 2.7. Propargyl 2,3,4,6-tetra-*O*-benzoyl-β-D-glucopyranosyl-(1→4)-2,3-di-*O*-benzoyl-β-D-glucopyranoside (12)

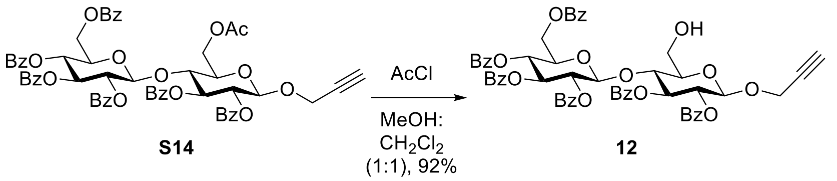

The protected disaccharide **S14** (1.16 g, 1.11 mmol, 1 equiv) was dissolved in a mixture of $CH_2Cl_2$:MeOH (1:1, 23 mL) and cooled to 0 °C. Acetyl chloride (712 µL, 9.97 mmol, 9 equiv) was added over a period of 5 min while stirring. The mixture was stirred for 1 h at 0 °C and for 17 h at room temperature, followed by addition of triethylamine (3 mL) at 0 °C. The solvent was removed under reduced pressure and the solid was suspended in ethyl acetate (50 mL) and filtrated. The solvent was removed under reduced pressure and the crude was purified on silica gel via automated flash chromatography (toluene:ethyl acetate, 4:1) to yield the product as a colorless foam (1.02 g, 1.02 mmol, 92%).

***R*<sub></sub>**$\boldsymbol{R_F}$ (toluene:ethyl acetate, 8:2) = 0.39.

$[\alpha]_D^{20}$ = -4.6 (*c* = 1.08 in $CHCl_3$).

**IR (ATR):** $\tilde{\nu}$ = 3294 (w), 3063 (w), 1719 (s), 1602 (m), 1451 (m), 1256 (s), 1091 (s), 1067 (s), 1026 (s), 705 (s), 687 (s) $cm^{-1}$.

**$^1$H NMR** (500 MHz, $CDCl_3$, 298 K) δ = 8.01 – 7.88 (m, 8H, OBz), 7.81 – 7.72 (m, 4H, OBz), 7.58 – 7.14 (m, 18H, OBz), 5.81 (t, $^3J_{2,3}$ = $^3J_{3,4}$ = 9.7 Hz, 1H, H-$3_{Glc'}$), 5.75 (t, $^3J_{2,3}$ = $^3J_{3,4}$ = 9.6 Hz, 1H, H-$3_{Glc}$), 5.50 (dd, $^3J_{2,3}$ = 9.9 Hz, $^3J_{1,2}$ = 8.0 Hz, 1H, H-$2_{Glc'}$), 5.41 (t, $^3J_{3,4}$ = $^3J_{4,5}$ = 9.7 Hz, 1H, H-$4_{Glc'}$), 5.37 (dd, $^3J_{2,3}$ = 9.8 Hz, $^3J_{1,2}$ = 8.0 Hz, 1H, H-$2_{Glc}$), 5.03 (d, $^3J_{1,2}$ = 8.0 Hz, 1H, H-$1_{Glc'}$), 4.94 (d, $^3J_{1,2}$ = 8.0 Hz, 1H, H-$1_{Glc}$), 4.37 – 4.23 (m, 3H, H-$4_{Glc}$, C$\underline{H}_2$C≡CH), 4.03 (dd, $^2J$ = 11.8 Hz, $^3J_{5,6a}$ = 3.3 Hz, 1H, H-$6a_{Glc'}$), 4.00 – 3.95 (m, 1H, H-$5_{Glc'}$), 3.83 (dd, $^2J$ = 11.8 Hz, $^3J_{5,6b}$ = 5.1 Hz, 1H, H-$6b_{Glc'}$), 3.81 – 3.78 (m, 2H, H-$6a_{Glc}$, H-$6b_{Glc}$), 3.50 (dt, $^3J_{4,5}$ = 9.8 Hz, $^3J_{5,6a}$ = $^3J_{5,6b}$ = 2.4 Hz, 1H, H-$5_{Glc}$), 2.35 (t, $^4J_{C≡CH,CH2}$ = 2.4 Hz, 1H, C≡C$\underline{H}$), 1.94 (t, $^3J_{OH,6}$ = 6.9 Hz, 1H, 6-OH) ppm.

**$^{13}$C NMR** (126 MHz, $CDCl_3$, 298 K) δ = 165.8 (Ph$\underline{C}$O), 165.7 (Ph$\underline{C}$O), 165.5 (Ph$\underline{C}$O), 165.3 (Ph$\underline{C}$O), 165.0 (Ph$\underline{C}$O), 164.7 (Ph$\underline{C}$O), 133.4 ($OBz_{para}$), 133.3 ($OBz_{para}$), 133.2 ($OBz_{para}$), 133.1 ($OBz_{para}$), 133.1 (2 x $OBz_{para}$), 129.8 (OBz), 129.8 (OBz), 129.7 (OBz), 129.7 (OBz), 129.7 (OBz), 129.6 (OBz), 129.5 (OBz), 129.5 (OBz), 129.3 (OBz), 129.0 (OBz), 128.9 (OBz), 128.7 (OBz), 128.7 (OBz), 128.5 (OBz), 128.3 (OBz), 128.3 (2 x OBz), 128.2 (OBz), 100.9 (C-$1_{Glc'}$), 98.6 (C-$1_{Glc}$), 78.1 ($CH_2\underline{C}$≡CH), 75.5 ($CH_2$C≡$\underline{C}$H), 75.2 (C-$4_{Glc}$), 75.1 (C-$5_{Glc}$), 72.9 (C-$3_{Glc'}$), 72.7 (C-$3_{Glc}$), 72.2 (C-$5_{Glc'}$), 72.0 (C-$2_{Glc'}$), 71.6 (C-$2_{Glc}$), 69.5 (C-$4_{Glc'}$), 62.7 (C-$6_{Glc'}$), 60.2 (C-$6_{Glc}$), 56.3 ($\underline{C}H_2$C≡CH) ppm.

**ESI-HRMS:** *m/z* = 1027.27851 $[M+Na]^+$ (calculated *m/z* = 1027.27837).

#### 2.8. Propargyl β-D-glucopyranosyl-(1→4)-β-D-glucopyranosyl-(1→4)-β-D-glucopyranoside (17)

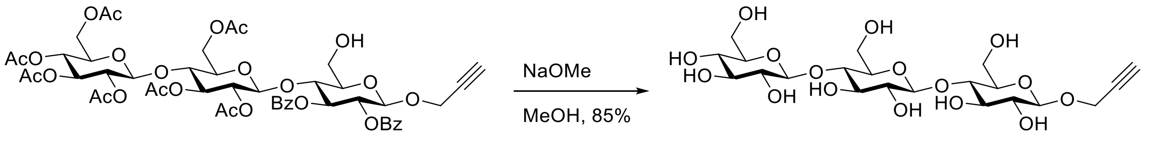

**S16** **17**

The protected trisaccharide **S16** (25.2 mg, 24.1 μmol, 1 equiv) was dissolved in methanol (2 mL) and a solution of sodium methoxide in methanol (10 μL, 50.0 μmol, 5 M, 2.07 equiv) was added at room temperature. The reaction mixture was stirred at room temperature for 18 h. It was neutralized by addition of Amberlite®-IRC 120 $H^+$ resin and stirring for 15 min. The resin was filtered off and the filtrate was concentrated under reduced pressure. The crude was purified on reversed phase silica gel via automated flash chromatography ($H_2O$:MeCN, 100:0 1 CV, 100:0 → 0:100 over 16 CV) and lyophilized to obtain the product as a lyophilizate (11.1 mg, 20.5 μmol, 85%).

$[\alpha]_D^{20}$= -43.1 (*c* = 0.23 in MeOH).

**IR (ATR):** $\tilde{\nu}$ = 3339 (m, br), 2889 (w), 1364 (w), 1157 (m), 1027 (s), 897 (w) $cm^{-1}$.

**$^1$H NMR** (600 MHz, methanol-d$_4$, 298 K) δ = 4.49 (d, $^3J_{1,2}$ = 7.9 Hz, 1H, H-1$_{Glc}$), 4.45 (d, $^3J_{1,2}$ = 7.9 Hz, 1H, H-1$_{Glc'}$), 4.46 – 4.36 (m, 3H, H-1$_{Glc''}$, C$\underline{H}_2$C≡CH), 3.93 – 3.83 (m, 5H, H-6a$_{Glc}$, H-6b$_{Glc}$, H-6a$_{Glc'}$, H-6b$_{Glc'}$, H-6a$_{Glc''}$), 3.66 (dd, $^2J_{6b,6a}$ = 11.9 Hz, $^3J_{5,6b}$ = 5.7 Hz, 1H, H-6b$_{Glc''}$), 3.61 – 3.56 (m, 2H, H-4$_{Glc}$, H-4$_{Glc'}$), 3.56 – 3.49 (m, 2H, H-3$_{Glc}$, H-3$_{Glc'}$), 3.47 (ddd, $^3J_{4,5}$ = 9.7 Hz, $^3J_{5,6a}$ = 4.2 Hz, $^3J_{5,6b}$ = 2.4 Hz, 1H, H-5$_{Glc/Glc'}$), 3.40 (ddd, $^3J_{4,5}$ = 9.5 Hz, $^3J_{5,6a}$ = 4.0 Hz, $^3J_{5,6b}$ = 2.6 Hz, 1H, H-5$_{Glc/Glc'}$), 3.38 – 3.31 (m, 3H, H-3$_{Glc''}$, H-4$_{Glc''}$, H-5$_{Glc''}$), 3.29 – 3.19 (m, 3H, H-2$_{Glc}$, H-2$_{Glc'}$, H-2$_{Glc''}$), 2.86 (t, $^4J_{C≡CH,CH2}$ = 2.4 Hz, 1H, C≡C$\underline{H}$) ppm.

**$^{13}$C NMR** (151 MHz, methanol-d$_4$, 298 K) δ = 104.6 (C-1$_{Glc''}$), 104.4 (C-1$_{Glc'}$), 101.9 (C-1$_{Glc}$), 80.4 (C-4$_{Glc/Glc'}$), 80.2 (C-4$_{Glc/Glc'}$), 79.9 (CH$_2\underline{C}$≡CH), 78.1 (C-5$_{Glc''}$), 77.9 (C-3$_{Glc''}$), 76.7 (C-5$_{Glc/Glc'}$), 76.6 (C-5$_{Glc/Glc'}$), 76.3 (C-3$_{Glc/Glc'}$), 76.3 (C-3$_{Glc/Glc'}$), 76.2 (CH$_2$C≡$\underline{C}$H), 74.9 (C-2$_{Glc''}$), 74.6 (C-2$_{Glc/Glc'}$), 74.6 (C-2$_{Glc/Glc'}$), 71.4 (C-4$_{Glc''}$), 62.4 (C-6$_{Glc''}$), 61.7 (C-6$_{Glc/Glc'}$), 61.5 (C-6$_{Glc/Glc'}$), 56.6 ($\underline{C}$H$_2$C≡CH) ppm.

**ESI-HRMS:** *m/z* = 543.19194 [M+H]$^+$ (calculated *m/z* = 543.19196).

##### 2.9. Propargyl 6-*N*-biotinamide-6-deoxy-α-D-glucopyranosyl-(1→4)-β-D-glucopyranosyl-(1→4)-β-D-glucopyranoside (18)

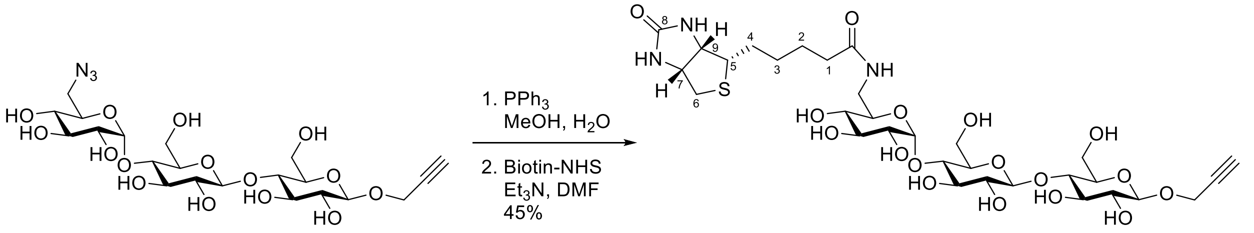

**S19** **18**

The 6''-azido-functionalized trisaccharide **S19** (34.0 mg, 60.0 µmol, 1 equiv) was dissolved in methanol (3 mL), triphenylphosphine (17.3 mg, 66.0 µmol, 1.1 equiv) was added and the reaction mixture stirred at room temperature for 18 h. Water (25 mL) was added and the mixture was filtered, followed by lyophilization of the filtrate. The crude and biotin-*N*-hydroxysuccinimide ester (24.6 mg, 72.0 µmol, 1.2 equiv) were dissolved in dry DMF (1 mL) under nitrogen atmosphere. At room temperature triethylamine (12 µL, 90 µmol, 1.5 equiv) was added and stirred for 18 h. The reaction mixture was purified on reversed phase silica gel via automated flash chromatography ($H_2O$:MeCN, 100:0 1 CV, 100:0 → 0:100 over 16 CV) and lyophilized to obtain the product as lyophilizate (20.5 mg, 26.7 µmol, 45%).

$[\alpha]_D^{20}$= +13.9 (*c* = 0.21 in MeOH).

**IR (ATR):** $\tilde{\nu}$ = 3299 (br, m), 2923 (w), 1674 (m), 1430 (m), 1366 (m), 1028 (s), 897 (w) cm$^{-1}$.

**$^1$H NMR** (600 MHz, $D_2O$, 298 K) δ = 5.45 (d, $^3J_{1,2}$ = 3.9 Hz, 1H, H-1$_{Glc''}$), 4.66 (d, $^3J_{1,2}$ = 8.0 Hz, 1H, H-1$_{Glc}$), 4.61 (dd, $^3J_{7,6b}$ = 7.9 Hz, $^3J_{7,6a}$ = 4.8 Hz, 1H, H-7$_{Biotin}$), 4.53 – 4.39 (m, 4H, H-1$_{Glc'}$, H-9$_{Biotin}$, C$\underline{H}_2$C≡CH), 3.97 (dd, $^2J$ = 12.3 Hz, $^3J_{5,6a}$ = 1.6 Hz, 1H, H-6a$_{Glc}$), 3.91 (dd, $^2J$ = 12.2 Hz, $^3J_{5,6a}$ = 2.2 Hz, 1H, H-6a$_{Glc'}$), 3.83 – 3.76 (m, 2H, H-3$_{Glc'}$, H-6b$_{Glc}$), 3.74 – 3.55 (m, 10H, H-2$_{Glc''}$, H-3$_{Glc}$, H-3$_{Glc''}$, H-4$_{Glc}$, H-4$_{Glc'}$, H-5$_{Glc}$, H-5$_{Glc'}$, H-5$_{Glc''}$, H-6b$_{Glc'}$, H-6a$_{Glc''}$), 3.37 – 3.29 (m, 4H, H-2$_{Glc}$, H-2$_{Glc'}$, H-5$_{Biotin}$, H-6b$_{Glc''}$), 3.26 (t, $^3J_{3,4}$ = $^3J_{4,5}$ = 9.4 Hz, 1H, H-4$_{Glc''}$), 3.00 (dd, $^2J_{6a,6b}$ = 13.1 Hz, $^3J_{7,6a}$ = 5.0 Hz, 1H, H-6a$_{Biotin}$), 2.91 (t, $^4J_{C≡CH,CH2}$ = 2.4 Hz, 1H, C≡C$\underline{H}$), 2.79 (d, $^2J_{6b,6a}$ = 13.0 Hz, 1H, H-6b$_{Biotin}$), 2.30 (t, $^3J_{1,2}$ = 7.4 Hz, 2H, H-1$_{Biotin}$), 1.74 (dq, $^2J_{4a,4b}$ = 13.8 Hz, $^3J_{3,4a}$ = 7.3 Hz, 1H, H-4a$_{Biotin}$), 1.69 – 1.56 (m, 3H, H-2$_{Biotin}$, H-4b$_{Biotin}$), 1.43 (p, $^3J_{2,3}$ = $^3J_{3,4}$ = 7.7 Hz, 2H, H-3$_{Biotin}$) ppm.

**$^{13}$C NMR** (151 MHz, $D_2O$, 298 K) δ = 177.0 ($CH_2\underline{C}$=ONH), 165.4 (C-$8_{Biotin}$), 102.4 (C-$1_{Glc'}$), 100.3 (C-$1_{Glc}$), 98.9 (C-$1_{Glc''}$), 78.8*, 78.7 ($CH_2\underline{C}$≡CH), 76.3 ($CH_2C$≡$\underline{C}$H), 76.0 (C-$3_{Glc'}$), 75.4*, 74.8*, 74.5*, 74.3*, 73.2 (C-$2_{Glc/Glc'}$), 72.6 (C-$2_{Glc/Glc'}$), 72.5*, 71.6 (C-$2_{Glc''}$), 71.14*, 71.08 (C-$4_{Glc''}$), 62.0 (C-$9_{Biotin}$), 60.6 (C-$6_{Glc'}$), 60.3 (C-$7_{Biotin}$), 59.9 (C-$6_{Glc}$), 56.5 ($\underline{C}H_2C$≡CH), 55.3 (C-$5_{Biotin}$), 40.0 (C-$6_{Glc''}$), 39.6 (C-$6_{Biotin}$), 35.5 (C-$1_{Biotin}$), 28.0 (C-$3_{Biotin}$), 27.7 (C-$4_{Biotin}$), 25.3 (C-$2_{Biotin}$) ppm.

*Assignment is ambiguous due to signal overlap (C-$3_{Glc}$, C-$3_{Glc''}$, C-$4_{Glc}$, C-$4_{Glc'}$, C-$5_{Glc}$, C-$5_{Glc'}$, C-$5_{Glc''}$).

**ESI-HRMS:** *m/z* = 790.26650 $[M+Na]^+$ (calculated *m/z* = 790.26749).

##### 2.10. Propargyl 6-*N*-acetylamino-6-deoxy-α-D-glucopyranosyl-(1→4)-β-D-glucopyranosyl-(1→4)-β-D-glucopyranoside (19)

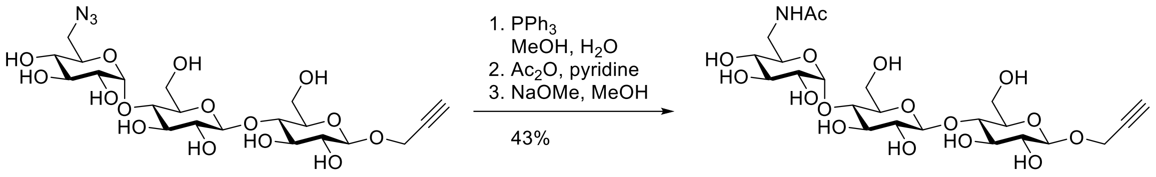

**S19** **18**

The 6''-azido-functionalied trisaccharide **S19** (14.0 mg, 24.7 µmol, 1.00 equiv) was dissolved in methanol (1.5 mL) and triphenylphosphine (7.1 mg, 27.1 µmol, 1.1 equiv) was added. The reaction mixture was stirred at room temperature for 16 h. Water (500 µL) was added and stirred for 2 h at room temperature. The solvent was removed under reduced pressure and the crude was dissolved in pyridine (1 mL), followed by addition of acetic anhydride (500 µL, 5.29 mmol, 214 equiv) at room temperature and stirred for 18 h. The solution was diluted with $CH_2Cl_2$ (10 mL), washed with 1 M HCl (2 x 10 mL) and the combined aq. phases were extracted with $CH_2Cl_2$ (10 mL). The combined organic phases were dried over $MgSO_4$, filtrated and the solvent was removed under reduced pressure. The crude was purified on silica gel via automated flash chromatography ($CH_2Cl_2$:MeOH, 100:0 → 95:5) to yield a mixture of peracetylated α-Glc(2,3,4-OAc)(6-NHAc)-(1→4)-β-Glc(2,3,6-OAc)-(1→4)-β-Glc(2,3,6-OAc)-Prop and partially acetylated α-Glc(2,3-OAc)(6-NHAc)-(1→4)-β-Glc(2,3,6-OAc)-(1→4)-β-Glc(2,3,6-OAc)-Prop. The crude mixture (22.2 mg) was dissolved in methanol (1 mL) and a solution of sodium methoxide in methanol (25.0 µL, 125 µmol, 5 M) was added at room temperature. The reaction mixture was stirred at room temperature for 21 h. It was neutralized by addition of Amberlite®-IRC 120 $H^+$ and stirring for 20 min. The resin was filtered off and the filtrate was concentrated under reduced pressure. The crude was purified on reversed phase silica gel via automated flash chromatography ($H_2O$:MeCN, 100:0 1 CV, 100:0 → 0:100 over 16 CV) and lyophilized to obtain the product as lyophilizate (6.1 mg, 10.5 µmol, 43%).

$[\boldsymbol{\alpha}]_{\mathbf{D}}^{\mathbf{20}}$= +22.2 (*c* = 0.25 in MeOH).

**IR (ATR):** $\tilde{\nu}$ = 3285 (br, m), 2886 (w), 1631 (m), 1561 (w), 1431 (m), 1372 (m), 1028 (s), 901 (m) $cm^{-1}$.

**$^{1}$H NMR** (500 MHz, methanol-$d_4$, 298 K) δ = 5.20 (d, $^3J_{1,2}$ = 3.8 Hz, 1H, H-$1_{Glc''}$), 4.50 (d, $^3J_{1,2}$ = 7.8 Hz, 1H, H-$1_{Glc}$), 4.47 – 4.36 (m, 3H, H-$1_{Glc''}$, $C\underline{H}_2C$≡CH), 3.94 – 3.83 (m, 3H, H-$6a_{Glc}$, H-$6b_{Glc}$, H-$6a_{Glc'}$), 3.75 (dd, $^2J_{6b,6a}$ = 12.2 Hz, $^3J_{5,6b}$ = 4.4 Hz, 1H, H-$6b_{Glc'}$), 3.72 – 3.50 (m, 7H, H-$3_{Glc}$, H-$3_{Glc'}$, H-$3_{Glc''}$, H-$4_{Glc}$, H-$4_{Glc'}$, H-$5_{Glc''}$, H-$6a_{Glc''}$), 3.47 – 3.37 (m, 3H, H-$2_{Glc''}$, H-$5_{Glc}$, H-$5_{Glc'}$), 3.33 – 3.23 (m,

3H, H-$2_{Glc}$, H-$2_{Glc'}$, H-$6b_{Glc''}$), 3.09 (t, $^3J_{3,4}$ = $^3J_{4,5}$ = 9.4 Hz, 1H, H-$4_{Glc''}$), 2.87 (t, $^4J_{C≡CH,CH2}$ = 2.4 Hz, 1H, C≡C$\underline{H}$), 1.98 (s, 3H, $CH_3$) ppm.

**$^{13}$C NMR** (126 MHz, methanol-$d_4$, 298 K) δ = 173.8 (C=O), 104.6 (C-$1_{Glc'}$), 102.7 (C-$1_{Glc''}$), 102.0 (C-$1_{Glc}$), 80.5*, 80.4*, 80.0 ($CH_2\underline{C}$≡CH), 77.7*, 76.6 (C-$5_{Glc}$, C-$5_{Glc'}$), 76.4*, 76.3 ($CH_2$C≡$\underline{C}$H), 74.7 (C-$2_{Glc}$, C-$2_{Glc'}$), 74.5*, 74.1 (C-$2_{Glc''}$), 73.1 (C-$4_{Glc''}$), 72.9*, 61.8 (C-$6_{Glc/Glc'}$), 61.7 (C-$6_{Glc/Glc'}$), 56.6 ($\underline{C}H_2$C≡CH), 41.9 (C-$6_{Glc''}$), 22.5 ($CH_3$).

*Assignment is ambiguous due to signal overlap (C-$3_{Glc}$, C-$3_{Glc'}$, C-$3_{Glc''}$, C-$4_{Glc}$, C-$4_{Glc'}$, C-$5_{Glc''}$).

**ESI-HRMS:** *m/z* = 606.19984 $[M+Na]^+$ (calculated *m/z* = 606.20046).

#### 2.11. Propargyl β-D-xylopyranosyl-(1→4)-β-D-xylopyranoside (21)

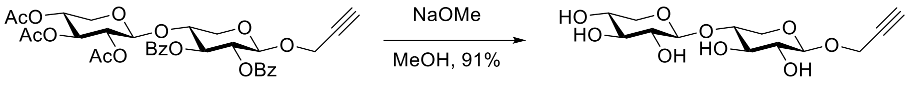

**S20** **21**

The protected disaccharide **S20** (220 mg, 336 µmol) was suspended in methanol (18 mL), sodium methoxide (5.4 M in methanol, 95 µL) was added dropwise at room temperature and the reaction mixture stirred for 18 h at room temperature. It was neutralized by addition of Amberlite®-IRC 120 $H^+$ resin. The resin was filtered off and the filtrate was concentrated under reduced pressure. The crude product was purified on reversed phase silica gel via automated flash chromatography ($H_2O$:MeCN, 100:0 1 CV, 100:0 → 0:100 over 16 CV) and lyophilized to obtain the product as lyophilizate (98.0 mg, 306 µmol, 91%).

$[\boldsymbol{\alpha}]_{\mathbf{D}}^{\mathbf{20}}$ = -96.7 (*c* = 0.99, MeOH).

**IR (ATR)**: $\tilde{\nu}$ = 3359, 3280, 2875, 2121, 1635, 1035, 980 $cm^{-1}$.

**$^1$H NMR** (600 MHz, methanol-$d_4$, 298 K): δ = 4.44 (d, $^3J_{1,2}$ = 7.4 Hz, 1H, H-1), 4.40-4.30 (m, 3H, C$\underline{H}_2$C≡CH, H-1'), 4.02 (dd, $^2J_{5ax,5eq}$ = 11.7 Hz, $^3J_{4,5eq}$ = 5.2 Hz, 1H, H-$5_{eq}$), 3.89 (dd, $^2J_{5ax,5eq}$ = 11.4 Hz, $^3J_{4,5eq}$ = 5.4 Hz, 1H, H-$5_{eq}$'), 3.68-3.62 (m, 1H, H-4), 3.53-3.44 (m, 2H, H-3, H-4'), 3.34-3.27 (m, 2H, H-3', H-$5_{ax}$), 3.26-3.17 (m, 3H, H-2, H-2', H-$5_{ax}$'), 2.87 (t, $^4J_{CH2,C≡CH}$ = 2.4 Hz, 1H, C≡C$\underline{H}$).

**$^{13}$C NMR** (151 MHz, methanol-$d_4$, 298 K): δ = 104.0 (C-1'), 102.6 (C-1), 79.8 ($\underline{C}$≡CH), 78.1 (C-4), 77.6 (C-3'), 76.3 (C≡$\underline{C}$H), 75.7 (C-3), 74.4 (C-2), 74.3 (C-2'), 71.1 (C-4'), 67.1 (C-5'), 64.5 (C-5), 56.5 ($\underline{C}H_2$C≡CH).

**ESI-HRMS:** *m/z* = 343.09944 $[M+Na]^+$ (calculated *m/z* = 343.09995 for $[M+Na]^+$).

#### 2.12. Propargyl 4-*N*-biotinamide-4-deoxy-β-D-xylopyranosyl-(1→4)-β-D-xylopyranoside (22)

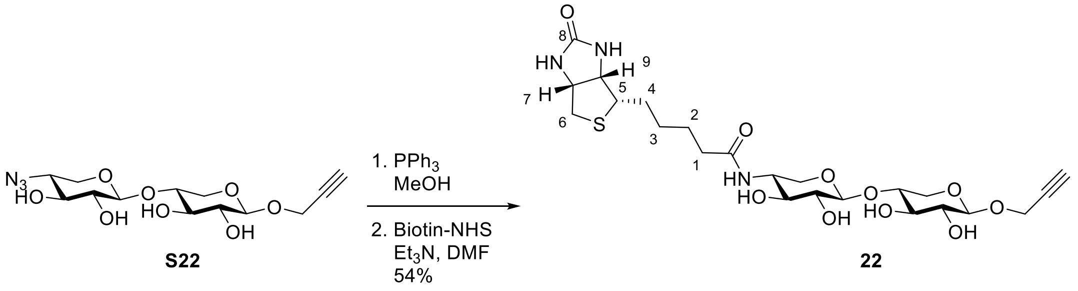

The 4''-azido-functionalized disaccharide **S22** (94.4 mg, 273 µmol) was dissolved in methanol (24 mL) and triphenylphosphine (108 mg, 410 µmol) was added. The reaction mixture was stirred at room temperature for 14 h, then it was diluted with water and dichloromethane. The aq. phase was washed with dichloromethane (5 x 30 mL) and finally with diethyl ether (30 mL). The aq. phase was concentrated. The resulting white solid was dissolved in dry DMF (10 mL) under a nitrogen atmosphere. Biotin-*N*-hydroxysuccinimide ester (127 mg, 371 µmol) and triethylamine (62 µL, 445 µmol) were added. The mixture was stirred at room temperature for 15 h. The reaction mixture was concentrated and the crude product was purified on reversed phase silica gel via automated flash chromatography ($H_2O$:MeCN, 100:0 1 CV, 100:0 → 0:100 over 16 CV) and lyophilized to obtain the title compound (80.0 mg, 147 µmol, 54%) with minor impurities (<10% according to $^1$H NMR).

$[\alpha]_D^{20}$ = -45.8 (*c* = 0.23, $H_2O$).

**IR (ATR):** $\tilde{\nu}$ = 3255, 2933, 1690, 1649, 1545, 1084, 990 cm$^{-1}$.

**$^1$H NMR** (600 MHz, $D_2O$, 298 K): δ = 4.56-4.51 (m, 2H, H-$7_{Biotin}$, H-1), 4.40-4.33 (m, 4H, C<u>H</u>$_2$C≡CH, H-1', H-$9_b$), 4.03 (dd, $^2J_{5ax,5eq}$ = 11.9 Hz, $^3J_{4,5eq}$ = 5.3 Hz, 1H, H-$5_{eq}$), 3.89-3.80 (m, 2H, H-$5_{eq}$', H-4'), 3.74-3.68 (m, 1H, H-4), 3.52 (t, $^3J_{2,3}$ = $^3J_{3,4}$ = 9.1 Hz, 1H, H-3), 3.46 (t, $^3J_{2,3}$ = $^3J_{3,4}$ = 9.5 Hz, 1H, H-3'), 3.34 (dd, $^2J_{5ax,5eq}$ = 11.9 Hz, $^3J_{4,5ax}$ = 10.2 Hz, 1H, H-$5_{ax}$), 3.29-3.19 (m, 4H, H-2', H-2, H-$5_{Biotin}$, H-$5_{ax}$'), 2.93 (dd, $^2J_{H6aBiotin,H6bBiotin}$ = 13.1 Hz, $^3J_{H6aBiotin,H7Biotin}$ = 5.0 Hz, 1H, H-$6a_{Biotin}$), 2.71 (d, $^2J_{H6aBiotin,H6bBiotin}$ = 13.1 Hz, 1H, H-$6b_{Biotin}$), 2.24-2.19 (m, 2H, H-$1_{Biotin}$), 1.69-1.47 (m, 4H, H-$2_{Biotin}$, H-$4_{Biotin}$), 1.41-1.29 (m, 2H, H-$3_{Biotin}$) ppm.

**$^{13}$C NMR** (151 MHz, $D_2O$, 298 K): δ = 177.0 ($CH_2$<u>C</u>=ONH), 165.0 (C-$8_{Biotin}$), 101.4 (C-1'), 100.9 (C-1), 77.9 (<u>C</u>≡CH), 75.9 (C-4), 73.2 (C-3), 73.0 (C-2'), 72.5 (C-3'), 72.2 (C-2), 63.1 (C-5'), 62.5 (C-5), 61.6 (C-$9_{Biotin}$), 59.8 (C-$7_{Biotin}$), 56.3 (<u>C</u>$H_2$C≡CH), 54.9 (C-$5_{Biotin}$), 50.5 (C-4'), 39.3 (C-$6_{Biotin}$), 35.1 (C-$1_{Biotin}$), 27.4 (C-$3_{Biotin}$), 27.2 (C-$4_{Biotin}$), 24.6 (C-$2_{Biotin}$) ppm.

Propargyl C≡<u>C</u>H cannot be assigned.

**ESI-HRMS:** *m/z* = 568.19332 $[M+Na]^+$ (calculated *m/z* = 568.19354 for $[M+Na]^+$).

#### 2.13. Propargyl β-D-xylopyranosyl-(1→4)-β-D-xylopyranosyl-(1→4)-β-D-xylopyranoside (23)

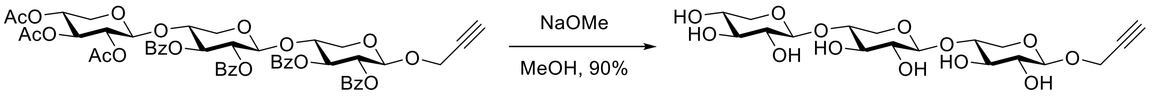

S23 23

The protected trisaccharide **S23** (54.7 mg, 55.0 µmol) was suspended in dry methanol (4 mL) under a nitrogen atmosphere and sodium methoxide (5 M in methanol, 22 µL, 110 µmol) was added dropwise. The reaction mixture was stirred at room temperature for 18 h. It was neutralized by addition of Amberlite®-IRC 120 $H^+$ resin. The resin was filtered off and the filtrate was concentrated under reduced pressure. The crude product was purified on reversed phase silica gel by automated flash chromatography ($H_2O$:MeCN, 100:0 1 CV, 100:0 → 0:100 over 16 CV) to obtain the title compound after lyophilization (22.3 mg, 49.3 µmol, 90%).

$[\alpha]_D^{20}$ = -99.2 (*c* = 0.16, MeOH).

**IR (ATR)**: $\tilde{\nu}$ = 3378, 2873, 1637, 1039, 981, 895 $cm^{-1}$.

**$^1$H NMR** (600 MHz, methanol-$d_4$, 298 K): δ = 4.43 (d, $^3J_{1,2}$ = 7.4 Hz, 1H, H-1), 4.39-4.29 (m, 4H, H-1', H-1'', C$\underline{H}_2$C≡CH), 4.04 (dd, $^2J_{5ax,5eq}$ = 11.7 Hz, $^3J_{4,5eq}$ = 5.3 Hz, 1H, H-$5_{eq}$'), 4.01 (dd, $^2J_{5ax,5eq}$ = 11.8 Hz, $^3J_{4,5eq}$ = 5.2 Hz, 1H, H-$5_{eq}$), 3.89 (dd, $^2J_{5ax,5eq}$ = 11.4 Hz, $^3J_{4,5eq}$ = 5.4 Hz, 1H, H-$5_{eq}$''), 3.70-3.62 (m, 2H, H-4, H-4'), 3.52-3.42 (m, 1H, H-4''), 3.47 (t, $^3J_{2,3}$ = $^3J_{3,4}$ = 8.6 Hz, 1H, H-3), 3.45 (t, $^3J_{2,3}$ = $^3J_{3,4}$ = 8.7 Hz, 1H, H-3'), 3.36-3.17 (m, 7H, H-2, H-2', H-2'', H-3'', H-$5_{ax}$, H-$5_{ax}$', H-$5_{ax}$''), 2.86 (t, $^4J_{CH2,C≡CH}$ = 2.5 Hz, 1H, C≡C$\underline{H}$) ppm.

**$^{13}$C NMR** (151 MHz, methanol-$d_4$, 298 K): δ = 104.0 (C-1''), 103.7 (C-1'), 102.6 (C-1), 79.8 ($\underline{C}$≡CH assigned by HMBC), 78.0 (C-4, C-4'), 77.6 (C-3''), 76.3 (C≡$\underline{C}$H), 75.7 (C-3, C-3'), 74.4 (C-2), 74.3 (C-2''), 74.1 (C-2'), 71.1 (C-4''), 67.1 (C-5''), 64.7 (C-5'), 64.4 (C-5), 56.5 ($\underline{C}H_2$C≡CH) ppm.

**ESI-HRMS:** *m/z* = 475.14149 $[M+Na]^+$ (calculated *m/z* = 475.14221 for $[M+Na]^+$).

#### 2.14. Propargyl β-D-xylopyranosyl-(1→4)-β-D-xylopyranosyl-(1→4)-β-D-xylopyranosyl-(1→4)-β-D-xylopyranoside (24)

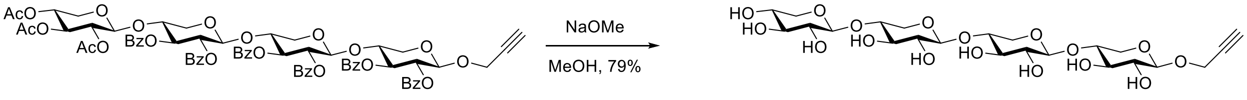

S24 24

The protected tetrasaccharide **S24** (42.0 mg, 31.5 µmol) was suspended in methanol (7 mL) and sodium methoxide (5.4 M in methanol, 40 µL, 216 µmol) was added dropwise. The reaction mixture was stirred at room temperature for 18 h. It was neutralized by addition of Amberlite®-IRC 120 $H^+$ resin. The resin was filtered off and the filtrate was concentrated under reduced pressure. The crude product was purified on reversed phase silica gel via automated flash chromatography ($H_2O$:MeCN, 100:0 1 CV, 100:0 → 0:100 over 16 CV) to obtain the title compound after lyophilization (14.6 mg, 25.0 µmol, 79%).

$[\alpha]_D^{20}$ = -108.1 (*c* = 0.20, MeOH).

**IR (ATR):** $\tilde{\nu}$ = 3366, 2918, 2118, 1163, 1035 $cm^{-1}$.

**$^{1}$H NMR** (500 MHz, methanol-d$_4$, 298 K): δ = 4.43 (d, $^{3}J_{1,2}$ = 7.4 Hz, 1H, H-1), 4.40-4.29 (m, 5H, H-1’, H-1’’, H-1’’’, C<u>H</u>$_2$C≡CH), 4.04 (dd, $^{2}J_{5ax,5eq}$ = 11.6 Hz, $^{3}J_{4,5eq}$ = 5.3 Hz, 2H, H-5$_{eq}$’, H-5$_{eq}$’’), 4.01 (dd, $^{2}J_{5ax,5eq}$ = 11.8 Hz, $^{3}J_{4,5eq}$ = 5.2 Hz, 1H, H-5$_{eq}$), 3.89 (dd, $^{2}J_{5ax,5eq}$ = 11.5 Hz, $^{3}J_{4,5eq}$ = 5.4 Hz, 1H, H-5$_{eq}$’’’), 3.70-3.61 (m, 3H, H-4, H-4’, H-4’’), 3.52-3.42 (m, 4H, H-4’’’, H-3, H-3’, H-3’’), 3.38-3.17 (m, 9H, H-5$_{ax}$, H-5$_{ax}$’, H-5$_{ax}$’’, H-5$_{ax}$’’’, H-3’’’, H-2, H-2’, H-2’’, H-2’’’), 2.87 (t, $^{4}J_{CH2,C≡CH}$ = 2.4 Hz, 1H, C≡C<u>H</u>) ppm.

**$^{13}$C NMR** (151 MHz, methanol-d$_4$, 298 K): δ = 104.0 (C-1’’’), 103.7 (C-1’, C-1’’), 102.6 (C-1), 79.8 (<u>C</u>≡CH), 78.0/77.9 (C-4, C-4’, C-4’’), 77.6 (C-3’’’), 76.3 (C≡<u>C</u>H), 75.7/75.6 (C-3, C-3’, C-3’’), 74.4 (C-2), 74.3 (C-2’’’), 74.1 (C-2’, C-2’’), 71.0 (C-4’’’), 67.1 (C-5’’’), 64.7/64.6 (C-5’, C-5’’), 64.4 (C-5), 56.5 (<u>C</u>H$_2$C≡CH) ppm.

**ESI-HRMS:** *m/z* = 607.18431 [M+Na]$^+$ (calculated *m/z* = 607.18447 for [M+Na]$^+$).

#### 2.15. Propargyl α-D-mannopyranosyl-(1→4)-[β-D-glucopyranosyl-(1→6)]-β-D-glucopyranoside (25)

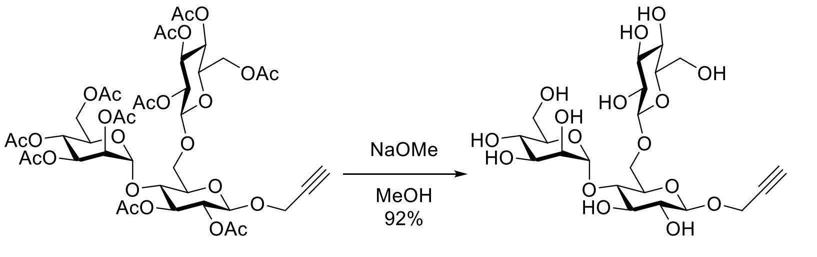

**S25** **25**

The protected trisaccharide **S25** (328 mg, 340 μmol, 1 equiv) was dissolved in methanol (20 mL) and a solution of sodium methoxide in methanol (50 μL, 250 μmol, 5 M, 0.74 equiv) was added at room temperature. The reaction mixture was stirred at room temperature for 5 h. It was neutralized by addition of Amberlite®-IRC 120 H$^+$ resin and stirring for 15 min. The resin was filtered off and the filtrate was concentrated under reduced pressure. The crude was purified on reversed phase silica gel via automated flash chromatography ($H_2O$:MeCN, 100:0 1 CV, 100:0 → 0:100 over 16 CV) to obtain the title compound after lyophilization (170 mg, 313 μmol, 92%).

$[\alpha]_{D}^{20}$ = -11.7 (*c* = 0.90 in MeOH).

**IR (ATR):** $\tilde{\nu}$ = 3278 (br, m), 2922 (w), 1647 (w), 1361 (m), 1011 (s), 812 (m) cm$^{-1}$.

**$^{1}$H NMR** (500 MHz, methanol-d$_4$, 298 K) δ = 5.30 (d, $^{3}J_{1,2}$ = 1.9 Hz, 1H, H-1$_{Man}$), 4.47 – 4.34 (m, 4H, H-1$_{Glc}$, H-1$_{Glc'}$, C<u>H</u>$_2$C≡CH), 4.20 (dd, $^{2}J$ = 11.4 Hz, $^{3}J_{5,6a}$ = 1.6 Hz, 1H, H-6a$_{Glc}$), 3.95 (dd, $^{3}J_{2,3}$ = 3.2 Hz, $^{3}J_{1,2}$ = 1.9 Hz, 1H, H-2$_{Man}$), 3.90 – 3.84 (m, 2H, H-6a$_{Glc'}$, H-6a$_{Man}$), 3.81 (dd, $^{2}J$ = 11.4 Hz, $^{3}J_{5,6b}$ = 5.3 Hz, 1H, H-6b$_{Glc}$), 3.72 – 3.62 (m, 4H, H-3$_{Man}$, H-5$_{Glc'}$, H-6b$_{Glc'}$, H-6b$_{Man}$), 3.61 – 3.53 (m, 2H, H-4$_{Glc}$, H-5$_{Glc}$, H-5$_{Man}$), 3.50 (t, $^{3}J_{2,3}$ = $^{3}J_{3,4}$ = 8.7 Hz, 1H, H-3$_{Glc}$), 3.40 – 3.33 (m, 1H, H-3$_{Glc'}$), 3.30 – 3.26 (m, 2H, H-4$_{Man}$, H-4$_{Glc'}$), 3.23 – 3.17 (m, 2H, H-2$_{Glc}$, H-2$_{Glc'}$), 2.86 (t, $^{4}J_{C≡CH,CH2}$ = 2.4 Hz, 1H, C≡C<u>H</u>) ppm.

**$^{13}$C NMR** (126 MHz, methanol-d$_4$, 298 K) δ = 104.8 (C-1$_{Glc'}$), 103.0 (C-1$_{Man}$), 102.2 (C-1$_{Glc}$), 80.2 (CH$_2$<u>C</u>≡CH), 78.5 (C-3$_{Glc}$), 78.2*, 78.0 (C-3$_{Glc'}$, C-4$_{Man}$/C-4$_{Glc'}$), 76.3 (CH$_2$C≡<u>C</u>H), 75.9*, 75.7*, 75.2 (C-2$_{Glc}$/C-2$_{Glc'}$), 75.1 (C-2$_{Glc}$/C-2$_{Glc'}$), 72.4*, 72.3 (C-2$_{Man}$), 71.6 (C-4$_{Man}$/C-4$_{Glc'}$), 70.2 (C-6$_{Glc}$), 68.7*, 63.3 (C-6$_{Glc'}$/C-6$_{Man}$), 62.8 (C-6$_{Glc'}$/C-6$_{Man}$), 56.9 (<u>C</u>H$_2$C≡CH) ppm.

*Assignment is ambiguous due to signal overlap (C-3$_{Man}$, C-4$_{Glc}$, C-5$_{Glc}$, C-5$_{Glc'}$, C-5$_{Man}$).

**ESI-HRMS:** *m/z* = 565.17277 $[M+Na]^+$ (calculated *m/z* = 565.17391).

#### 2.16. Propargyl β-D-glucopyranosyl-(1→4)-[α-D-mannopyranosyl-(1→6)]-β-D-glucopyranoside (26)

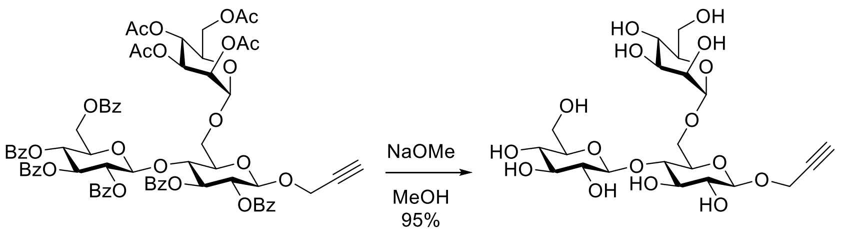

**S26** **26**

The protected trisaccharide **S26** (434 mg, 325 μmol, 1 equiv) was dissolved in methanol (20 mL) and a solution of sodium methoxide in methanol (100 μL, 500 μmol, 5 M, 1.5 equiv) was added at room temperature. The reaction mixture was stirred at room temperature for 18 h. It was neutralized by addition of Amberlite®-IRC 120 $H^+$ resin and stirring for 30 min. The resin was filtered off and the filtrate was concentrated under reduced pressure. The crude was purified on reversed phase silica gel via automated flash chromatography ($H_2O$:MeCN, 100:0 1 CV, 100:0 → 0:100 over 16 CV) to obtain the title compound after lyophilization (168 mg, 310 μmol, 95%).

$[\alpha]_D^{20}$ = -9.1 (*c* = 0.51 in MeOH).

**IR (ATR):** $\tilde{\nu}$ = 3324 (br, m), 2922 (w), 1637 (w), 1364 (m), 1020 (s), 895 (m), 811 (m) $cm^{-1}$.

**$^1$H NMR** (500 MHz, methanol-d$_4$, 298 K) δ = 4.86* (m, 1H, H-1$_{Man}$), 4.48 (d, $^3J_{1,2}$ = 7.8 Hz, 1H, H-1$_{Glc}$), 4.44 – 4.34 (m, 3H, H-1$_{Glc'}$, C$\underline{H}_2$C≡CH), 4.03 (dd, $^2J$ = 11.6 Hz, $^3J_{5,6a}$ = 3.7 Hz, 1H, H-6a$_{Glc}$), 3.93 – 3.81 (m, 4H, H-2$_{Man}$, H-6b$_{Glc}$, H-6a$_{Glc'}$, H-6a$_{Man}$), 3.75 – 3.61 (m, 5H, H-3$_{Man}$, H-5$_{Glc'}$, H-5$_{Man}$, H-6b$_{Glc'}$, H-6b$_{man}$), 3.57 – 3.49 (m, 3H, H-3$_{Glc}$, H-4$_{Glc}$, H-5$_{Glc}$), 3.39 (dd, $^3J$ = 9.3 Hz, $^3J$ = 8.8 Hz, 1H, H-3$_{Glc'}$), 3.37 – 3.18 (m, 4H, H-2$_{Glc}$, H-2$_{Glc'}$, H-4$_{Glc'}$, H-4$_{Man}$), 2.88 (t, $^4J_{C≡CH,CH2}$ = 2.4 Hz, 1H, C≡C$\underline{H}$) ppm.

*Chemical shift was deduced from HSQC-NMR due to signal overlap with residual HDO.

**$^{13}$C NMR** (126 MHz, methanol-d$_4$, 298 K) δ = 104.6 (C-1$_{Glc'}$), 102.1 (C-1$_{Glc}$/C-1$_{Man}$), 102.0 (C-1$_{Glc}$/C-1$_{Man}$), 80.6 (C-4$_{Glc}$/C-5$_{Glc}$), 79.9 ($CH_2\underline{C}$≡CH), 78.2 (C-4$_{Man}$/C-4$_{Glc'}$), 77.8 (C-3$_{Glc'}$), 76.4 ($CH_2$C≡$\underline{C}$H), 76.3 (C-3$_{Glc}$), 75.5 (C-4$_{Glc}$/C-5$_{Glc}$), 75.1 (C-2$_{Glc}$/C-2$_{Glc'}$), 74.7 (C-2$_{Glc}$/C-2$_{Glc'}$), 74.6 (C-5$_{Man}$/C-5$_{Glc}$), 72.7 (C-3$_{Man}$), 72.0 (C-2$_{Man}$), 71.5 (C-4$_{Man}$/C-4$_{Glc'}$), 68.6 (C-5$_{Man}$/C-5$_{Glc}$), 66.7 (C-6$_{Glc}$), 63.1 (C-6$_{Glc'}$/ C-6$_{Man}$), 62.5 (C-6$_{Glc'}$/C-6$_{Man}$), 56.8 ($\underline{C}H_2$C≡CH) ppm.

**ESI-HRMS:** *m/z* = 560.21771 $[M+NH_4]^+$ (calculated *m/z* = 560.21851).

#### 2.17. Phenyl 1-thio-2,3,4-tri-*O*-acetyl-β-D-xylopyranosyl-(1→4)-2,3-di-*O*-benzoyl-β-D-xylopyranoside (S6)

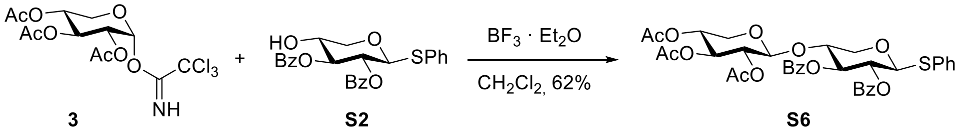

**3** **S2** **S6**

The xylosyl donor **3**[3] (207 mg, 492 µmol) and acceptor xyloside **S2**[11] (148 mg, 329 µmol) were dissolved in dry $CH_2Cl_2$ (6.6 mL) under a nitrogen atmosphere, freshly activated powdered molecular sieves (4Å, ~170 mg) was added and the reaction mixture was stirred at room temperature for 15 min. At 0°C boron trifluoride diethyl etherate (10 µL, 79 µmol) was added. The reaction mixture was stirred at 0 °C for 2 h followed by 12 h at room temperature. The reaction mixture was quenched with a satd. aq. $NaHCO_3$ solution (1 mL) and diluted with $CH_2Cl_2$ (40 mL). After filtration over celite® the organic phase was washed with a satd. aq. $NaHCO_3$ solution (10 mL) and the aq. phase was extracted with $CH_2Cl_2$ (10 mL). All organic phases were combined and dried over $MgSO_4$, filtrated and the solvent was removed under reduced pressure. The crude product was purified on silica gel via automated flash chromatography (cyclohexane:ethyl acetate, 9:1→6:4) to obtain the product as a colorless amorphous solid (145 mg, 205 µmol, 62%).

***R*<sub></sub>** 

$\boldsymbol{R}_{\mathrm{F}}$ (cyclohexane:ethyl acetate, 3:2) = 0.36.

$[\boldsymbol{\alpha}]_{\mathbf{D}}^{\mathbf{20}}$ = -25.4 (*c* = 0.24, $CHCl_3$).

**IR (ATR):** $\tilde{\nu}$ = 2959, 1752, 1732, 1718, 1249, 1219, 708 $cm^{-1}$.

**$^1$H NMR** (500 MHz, $CDCl_3$, 298 K): δ = 8.02-7.96 (m, 4H, 2 x $OBz_{ortho}$), 7.57-7.50 (m, 2H, 2 x $OBz_{para}$), 7.49-7.45 (m, 2H, $SPh_{ortho}$), 7.44-7.36 (m, 4H, 2 x $OBz_{meta}$), 7.32-7.28 (m, 3H, $SPh_{para}$, $SPh_{meta}$), 5.61 (t, $^3J_{2,3}$ = $^3J_{3,4}$ =7.8 Hz, 1H, H-3), 5.34 (t, $^3J_{1,2}$ = $^3J_{2,3}$ = 7.9 Hz, 1H, H-2), 5.04 (d, $^3J_{1,2}$ = 7.9 Hz, 1H, H-1), 5.00 (t, $^3J_{2,3}$ = $^3J_{3,4}$ = 7.4 Hz, 1H, H-3'), 4.79 (dd, $^3J_{2,3}$ = 7.5 Hz, $^3J_{1,2}$ = 5.7 Hz, 1H, H-2'), 4.67-4.60 (m, 2H, H-1', H-4'), 4.31 (dd, $^2J_{5ax,5eq}$ = 12.0 Hz, $^3J_{4,5eq}$ = 4.6 Hz, 1H, H-$5_{eq}$), 4.05-3.98 (m, 1H, H-4), 3.73 (dd, $^2J_{5ax,5eq}$ = 12.2 Hz, $^3J_{4,5eq}$ = 4.4 Hz, 1H, H-$5_{eq}$'), 3.58 (dd, $^2J_{5ax,5eq}$ = 12.1 Hz, $^3J_{4,5ax}$ = 8.4 Hz, 1H, H-$5_{ax}$), 3.15 (dd, $^2J_{5ax,5eq}$ = 12.1 Hz, $^3J_{4,5ax}$ = 7.3 Hz, 1H, H-$5_{ax}$'), 2.03 (s, 3H, $CH3_{OAc\ at\ C\text{-}2'}$), 1.99 (s, 3H, $CH3_{OAc\ at\ C\text{-}3'}$), 1.98 (s, 3H, $CH3_{OAc\ at\ C\text{-}4'}$) ppm.

**$^{13}$C NMR** (126 MHz, $CDCl_3$, 298 K): δ = 170.0 ($C{=}O_{OAc\ at\ C\text{-}3'}$), 169.7 ($C{=}O_{OAc\ at\ C\text{-}4'}$), 169.2 ($C{=}O_{OAc\ at\ C\text{-}2'}$), 165.3 ($C{=}O_{OBz\ at\ C\text{-}3}$), 165.2 ($C{=}O_{OBz\ at\ C\text{-}2}$), 133.33 ($OBz_{para}$), 133.28 ($OBz_{para}$), 132.8 ($SPh_{ipso}$), 132.6 ($SPh_{ortho}$), 129.9 ($OBz_{ortho}$), 129.7 ($OBz_{ortho}$), 129.4 ($OBz_{ipso}$), 129.3 ($OBz_{ipso}$), 129.0 ($SPh_{meta}$), 128.44 ($OBz_{meta}$), 128.41 ($OBz_{meta}$), 128.1 ($SPh_{para}$), 99.4 (C-1'), 86.7 (C-1), 74.3 (C-4), 72.7 (C-3), 70.3 (C-2), 70.1 (C-3'), 70.0 (C-2'), 68.0 (C-4'), 65.5 (C-5), 61.0 (C-5'), 20.72 ($CH3_{OAc}$), 20.67 ($CH3_{OAc}$), 20.60 ($CH3_{OAc}$) ppm.

**ESI-HRMS:** *m/z* = 731.17665 $[M+Na]^+$ (calculated *m/z* = 731.17688 for $[M+Na]^+$).

##### 2.18. 2,3,4-Tri-*O*-acetyl-β-D-xylopyranosyl-(1→4)-2,3-di-*O*-benzoyl-D-xylopyranose (S7)

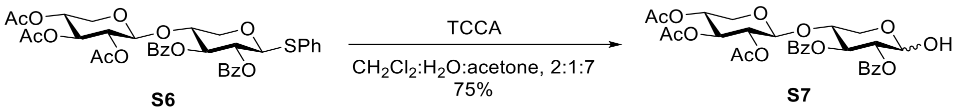

The thioglycoside **S6** (218 mg, 308 µmol) was dissolved in $CH_2Cl_2$ (2 mL) and a mixture of acetone (7 mL) and water (1 mL) was added. The mixture was stirred and cooled to 0°C and trichloroisocyanuric acid (71.6 mg, 308 µmol) was added slowly. The reaction mixture was stirred at 0 °C. After 1 h the organic solvent was removed in vacuo and the crude product was diluted with dichloromethane (75 mL) and washed with satd. aq. $NaHCO_3$ (25 mL) solution. The organic phase was separated and the aq. phase was extracted with dichloromethane (25 mL). The combined organic phases were washed with brine (25 mL). The organic phase was separated and the aq. phase was extracted with dichloromethane (25 mL). The combined organic phases were dried over $MgSO_4$, filtered and the solvent was removed under reduced pressure. The crude product was purified on silica gel via automated flash chromatography (cyclohexane:ethyl acetate, 9:1→1:1) to obtain the product as a colorless amorphous solid (142 mg, 230 µmol, 75%).

***R*F** (cyclohexane:ethyl acetate, 3:2) = 0.19.

$[\alpha]_D^{20}$ = +19.5 (*c* = 0.17, $CHCl_3$).

**IR (ATR):** $\tilde{\nu}$ = 3410, 1743, 1721, 1705, 1238, 1054, 707 $cm^{-1}$.

Alpha (major 79%)

**$^1$H NMR** (500 MHz, $CDCl_3$, 298 K): 8.01-7.94 (m, 4H, 2 x $OBz_{ortho}$), 7.55-7.49 (m, 2H, 2 x $OBz_{para}$), 7.42-7.35 (m, 4H, 2 x $OBz_{meta}$), 5.94 (dd, $^3J$ = 9.9 Hz, $^3J$ = 8.8 Hz, 1H, H-3), 5.59 (t, $^3J_{1,2}$ = $^3J_{1,OH}$ =3.8 Hz, 1H, H-1), 5.13-5.09 (m, 1H, H-2), 4.98 (t, $^3J_{2,3}$ = $^3J_{3,4}$ = 7.0 Hz, 1H, H-3'), 4.78 (dd, $^3J_{2,3}$ = 7.0 Hz, $^3J_{1,2}$ = 5.3 Hz, 1H, H-2'), 4.67-4.61 (m, 1H, H-4'), 4.65 (d, $^3J_{1,2}$ = 5.3 Hz, 1H, H-1'), 4.16-3.98 (m, 2H, H-4, H-5$_{eq}$), 3.85-3.80 (m, 1H, H-5$_{ax}$), 3.77-3.72 (m, 1H, H-5$_{eq}$'), 3.15 (dd, $^2J_{5ax,5eq}$ = 12.3 Hz, $^3J_{4,5ax}$ = 6.6 Hz, 1H, H-5$_{ax}$'), 2.84-2.82 (m, 1H, OH), 2.04 (s, 3H, $CH3_{OAc\ at\ C\text{-}2'}$), 2.01 (s, 3H, $CH3_{OAc\ at\ C\text{-}3'}$), 2.00 (s, 3H, $CH3_{OAc\ at\ C\text{-}4'}$) ppm.

**$^{13}$C NMR** (126 MHz, $CDCl_3$, 298 K): δ = 170.0 ($C{=}O_{OAc\ at\ C\text{-}3'}$), 169.7 ($C{=}O_{OAc\ at\ C\text{-}4'}$), 169.2 ($C{=}O_{OAc\ at\ C\text{-}2'}$), 165.9 ($C{=}O_{OBz\ at\ C\text{-}2}$), 165.4 ($C{=}O_{OBz\ at\ C\text{-}3}$), 133.4 ($OBz_{para}$), 133.1 ($OBz_{para}$), 129.9 ($OBz_{ortho}$), 129.8 ($OBz_{ipso}$), 129.6 ($OBz_{ortho}$), 129.0 ($OBz_{ipso}$), 128.5 ($OBz_{meta}$), 128.4 ($OBz_{meta}$), 98.9 (C-1'), 90.5 (C-1), 75.0 (C-4), 72.3 (C-2), 70.3 (C-3), 69.8 (C-2'), 69.7 (C-3'), 67.9 (C-4'), 60.7 (C-5'), 59.4 (C-5), 20.8 ($CH3_{OAc}$), 20.7 ($CH3_{OAc}$), 20.6 ($CH3_{OAc}$) ppm.

Beta (minor 21%)

**$^1$H NMR** (500 MHz, $CDCl_3$, 298 K): 8.01-7.94 (m, 4H, 2 x $OBz_{ortho}$), 7.55-7.49 (m, 2H, 2 x $OBz_{para}$), 7.42-7.35 (m, 4H, 2 x $OBz_{meta}$), 5.70-5.66 (m, 1H, H-3), 5.13-5.09 (m, 1H, H-2), 4.99 (t, $^3J_{2,3}$ = $^3J_{3,4}$ =7.2 Hz, 1H, H-3'), 4.87-4.83 (m, 1H, H-1), 4.80-4.75 (m, 1H, H-2'), 4.67-4.61 (m, 2H, H-1', H-4'), 4.16-3.98 (m, 2H, H-4, H-5$_{eq}$), 3.77-3.72 (m, 1H, OH), 3.70 (dd, $^2J_{5ax,5eq}$ = 12.2 Hz, $^3J_{4,5eq}$ = 4.3 Hz, 1H, H-5$_{eq}$'), 3.53-3.47 (m, 1H, H-5$_{ax}$), 3.17-3.11 (m, 1H, H-5$_{ax}$'), 2.04 (s, 3H, $CH3_{OAc}$), 2.00 (s, 3H, $CH3_{OAc}$), 1.996 (s, 3H, $CH3_{OAc}$) ppm.

**$^{13}$C NMR** (126 MHz, $CDCl_3$, 298 K): δ = 170.0 ($C{=}O_{OAc\ at\ C\text{-}3'}$), 169.7 ($C{=}O_{OAc\ at\ C\text{-}4'}$), 169.2 ($C{=}O_{OAc\ at\ C\text{-}2'}$), 167.2 ($C{=}O_{OBz\ at\ C\text{-}2}$), 165.4 ($C{=}O_{OBz\ at\ C\text{-}3}$), 133.7 ($OBz_{para}$), 133.3 ($OBz_{para}$), 130.0 ($OBz_{ortho}$), 129.6 ($OBz_{ortho}$), 129.4 ($OBz_{ipso}$), 128.7 ($OBz_{ipso}$), 128.5 ($OBz_{meta}$), 128.4 ($OBz_{meta}$), 99.2 (C-1'), 96.4 (C-1), 75.0 (C-4), 74.3 (C-2), 72.3 (C-3), 69.9 (C-2'), 69.8 (C-3'), 67.9 (C-4'), 63.5 (C-5), 60.9 (C-5'), 20.8 ($CH3_{OAc}$), 20.7 ($CH3_{OAc}$), 20.6 ($CH3_{OAc}$) ppm.

**ESI-HRMS:** *m/z* = 639.16804 $[M+Na]^+$ (calculated *m/z* = 639.16843).

##### 2.19. Propargyl 2,3-di-*O*-acetyl-4,6-*O*-benzylidene-β-D-glucopyranoside (S8)

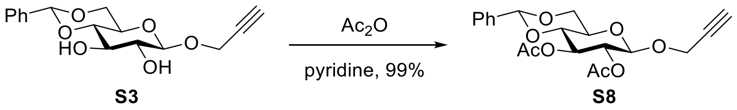

The benzylidene-protected glucoside **S3**[12,13] (1.59 g, 5.17 mmol) was dissolved in pyridine (15 mL), acetic anhydride (7.5 mL, 79.3 mmol) was added at room temperature and the reaction mixture was stirred for 2.5 h at room temperature. Then, the solvent was removed under reduced pressure and the residue co-evaporated with toluene (3 x 50 mL) and then dichloromethane (50 mL) to yield a colorless amorphous solid (2.02 g, 5.17 mmol, 99%).

***R*F** (cyclohexane:ethyl acetate, 8:2) = 0.32.

$[\alpha]_D^{20}$ = +104.7 (*c* = 0.92 in $CHCl_3$).

**IR (ATR):** $\tilde{\nu}$ = 3268 (w), 2891 (w), 1748 (s), 1459 (w), 1374 (m), 1235 (s), 1218 (s), 1045 (s), 1029 (s), 992 (s); 971 (s), 905 (m), 763 (m), 697 (s), 657 (m) $cm^{-1}$.

**$^1$H NMR** (600 MHz, $CDCl_3$, 298 K) δ = 7.46 – 7.41 (m, 2H, <u>Ph</u>$CH_{ortho}$), 7.39 – 7.33 (m, 3H, <u>Ph</u>$CH_{meta+para}$), 5.51 (s, 1H, PhC<u>H</u>), 5.35 (t, $^3J_{2,3}$ = $^3J_{3,4}$ = 9.5 Hz, 1H, H-3), 5.02 (dd, $^3J_{2,3}$ = 9.2 Hz, $^3J_{1,2}$ = 7.9 Hz, 1H, H-2), 4.85 (d, $^3J_{1,2}$ = 7.9 Hz, 1H, H-1), 4.40 – 4.36 (m, 3H, C<u>H</u>$_2$C≡CH, H-6a), 3.80 (t, $^2J$ = $^3J_{5,6a}$ = 10.3 Hz, 1H, H-6b), 3.71 (t, $^3J_{3,4}$ = $^3J_{4,5}$ = 9.6 Hz, 1H, H-4), 3.56 (td, $^3J_{4,5}$ = $^3J_{5,6b}$ = 9.7 Hz, $^3J_{5,6a}$ = 5.0 Hz, 1H, H-5), 2.48 (t, $^4J_{C≡CH,CH2}$ = 2.4 Hz, 1H, C≡C<u>H</u>), 2.08 (s, 3H, OAc), 2.05 (s, 3H, OAc) ppm.

**$^{13}$C NMR** (151 MHz, $CDCl_3$, 298 K) δ = 170.1 ($CH_3$<u>C</u>O), 169.7 ($CH_3$<u>C</u>O), 136.8 (<u>Ph</u>$_{ipso}$CH), 129.2 (<u>Ph</u>$_{para}$CH), 128.3 (<u>Ph</u>$_{meta}$CH), 126.2 (<u>Ph</u>$_{ortho}$CH), 101.6 (Ph<u>C</u>H), 98.9 (C-1), 78.3 (C-4), 78.1 ($CH_2$<u>C</u>≡CH), 75.5 ($CH_2$C≡<u>C</u>H), 71.9 (C-2), 71.7 (C-3), 68.5 (C-6), 66.4 (C-5), 56.2 (<u>C</u>$H_2$C≡CH), 20.8 ($CH_3$), 20.7 ($CH_3$) ppm.

**ESI-HRMS:** *m/z* = 408.16452 $[M+NH_4]^+$ (calculated *m/z* = 408.16529).

##### 2.20. Propargyl 2,3-di-*O*-acetyl-6-*O*-benzyl-β-D-glucopyranoside (S9)

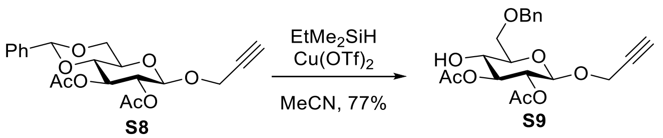

The benzylidene-protected glucoside **S8** (1.94 g, 4.97 mmol, 1 equiv) was dissolved in dry acetonitrile (45 mL) under nitrogen atmosphere. Ethyldimethylsilane (1.31 mL, 9.94 mmol, 2 equiv) was added and the reaction mixture was cooled to 0 °C. A solution of freshly activated (vacuum, ~200 °C, 15 min) copper(II)triflate (18 mg, 49.7 µmol, 0.01 equiv) in acetonitrile (5 mL) was added and stirred at 0 °C for 1 h, followed by stirring at room temperature for 1 h. Ethyl acetate (20 mL) was added and the reaction mixture was stirred strongly under air atmosphere for 15 min. The reaction mixture was diluted with ethyl acetate (300 mL) and washed with satd. aq. $NaHCO_3$ solution (100 mL), dried over $MgSO_4$, filtrated and the solvent was removed under reduced pressure. The crude was purified on silica gel via automated flash chromatography (cyclohexane:ethyl acetate, 2:1) to yield the product as a colorless oil (1.50 g, 3.82 mmol, 77%).

***R*F** (cyclohexane:ethyl acetate, 2:1) = 0.23.

$[\alpha]_D^{20}$ = -58.3 (*c* = 1.26 in $CHCl_3$).

**IR (ATR):** $\tilde{\nu}$ = 3449 (br, w), 3276 (w), 2871 (w), 1747 (s), 1453 (w), 1365 (m), 1238 (s), 1218 (s), 1051 (s), 1032 (s), 903 (m), 698 (m) $cm^{-1}$.

**$^1$H NMR** (500 MHz, $CDCl_3$, 298 K) δ = 7.39 – 7.28 (m, 5H, OBn), 5.08 (dd, $^3J_{2,3}$ = 9.7 Hz, $^3J_{3,4}$ = 9.2 Hz, 1H, H-3), 4.94 (dd, $^3J_{2,3}$ = 9.7 Hz, $^3J_{1,2}$ = 7.9 Hz, 1H, H-2), 4.73 (d, $^3J_{1,2}$ = 7.9 Hz, 1H, H-1), 4.65 – 4.53 (m, 2H, Bn-$C\underline{H}_2$), 4.35 (dd, $^4J_{C≡CH,CH2}$ = 2.4 Hz, $J$ = 0.7 Hz, 2H, $C\underline{H}_2C≡CH$), 3.84 – 3.71 (m, 3H, H-4, H-6a, H-6b), 3.56 (dt, $^3J_{4,5}$ = 9.5 Hz, $^3J_{5,6a}$ = $^3J_{5,6b}$ = 4.7 Hz, 1H, H-5), 2.99 (d, $^3J_{OH,4}$ = 3.6 Hz, 1H, OH-4), 2.45 (t, $^4J_{C≡CH,CH2}$ = 2.4 Hz, 1H, $C≡C\underline{H}$), 2.08 (s, 3H, OAc), 2.06 (s, 3H, OAc) ppm.

**$^{13}$C NMR** (126 MHz, $CDCl_3$, 298 K) δ = 171.3 ($CH_3\underline{C}O$), 169.7 ($CH_3\underline{C}O$), 137.5 ($OBn_{ipso}$), 128.5 ($OBn_{ortho/meta}$), 128.0 ($OBn_{para}$), 127.8 ($OBn_{ortho/meta}$), 98.2 (C-1), 78.4 ($CH_2\underline{C}≡CH$), 75.6 (C-3), 75.3 ($CH_2C≡\underline{C}H$), 74.2 (C-5), 73.8 ($Ph\underline{C}H_2$), 71.0 (C-2), 70.7 (C-4), 69.9 (C-6), 55.8 ($\underline{C}H_2C≡CH$), 20.9 ($CH_3$), 20.7 ($CH_3$) ppm.

**ESI-HRMS:** *m/z* = 410.18030 $[M+NH_4]^+$ (calculated *m/z* = 410.18094).

#### 2.21. Propargyl 2,3-di-*O*-benzoyl-4,6-*O*-benzylidene-β-D-glucopyranoside (S10)

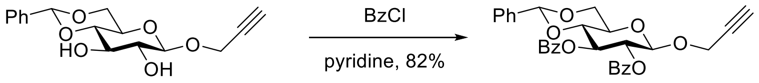

**S3** **S10**

The benzylidene-protected glucoside **S3**[13] (1.99 g, 6.50 mmol, 1 equiv) was dissolved in pyridine (14 mL) and cooled to 0 °C. Benzoyl chloride (2.41 mL, 20.7 mmol, 3.2 equiv) was added and the reaction mixture was stirred at room temperature overnight. The reaction was quenched by addition of methanol (5 mL) and stirring for 15 min at room temperature. The reaction mixture was diluted with $CH_2Cl_2$ (150 mL) and washed with water (50 mL). The aq. phase was extracted with $CH_2Cl_2$ (2 x 50 mL) and the combined organic phases were dried over $MgSO_4$, filtrated and the solvent was removed under reduced pressure. The crude was purified on silica gel via automated flash chromatography (cyclohexane:ethyl acetate, 100:0 → 75:25 over 12 CV) to yield the product as a colorless foam (2.73 g, 5.31 mmol, 82%).

$\boldsymbol{R}_\mathbf{F}$ (cyclohexane:ethyl acetate, 8:2) = 0.46.

$[\boldsymbol{\alpha}]_\mathbf{D}^\mathbf{20}$ = +4.8 ($c$ = 1.17 in $CHCl_3$).

**IR (ATR):** $\tilde{\nu}$ = 3287 (w), 2976 (w), 2958 (w), 2878 (w), 1729 (s), 1721 (s), 1602 (w), 1451 (m), 1266 (s), 1179 (m), 1070 (s), 976 (s), 699 (s) $cm^{-1}$.

**$^1$H NMR** (600 MHz, $CDCl_3$, 298 K) δ = 8.01 – 7.94 (m, 4H, 2 x $OBz_{ortho}$), 7.54 – 7.29 (m, 11H, 2 x $OBz_{meta+para}$, $\underline{Ph}CH$), 5.80 (t, $^3J_{2,3}$ = $^3J_{3,4}$ = 9.5 Hz, 1H, H-3), 5.55 (s, 1H, $PhC\underline{H}$), 5.49 (dd, $^3J_{2,3}$ = 9.4 Hz, $^3J_{1,2}$ = 7.9 Hz, 1H, H-2), 5.09 (d, $^3J_{1,2}$ = 7.8 Hz, 1H, H-1), 4.47 – 4.35 (m, 3H, H-6a, $C\underline{H}_2C≡CH$), 3.94 (t, $^3J_{3,4}$ = $^3J_{4,5}$ = 9.5 Hz, 1H, H-4), 3.89 (t, $^2J$ = $^3J_{5,6b}$ = 10.3 Hz, 1H, H-6b), 3.74 (ddd, $^3J_{5,6b}$ = 9.8 Hz, $^3J_{4,5}$ = 9.7 Hz, $^3J_{5,6a}$ = 4.9 Hz, 1H, H-5), 2.40 (t, $^4J_{C≡CH,CH2}$ = 2.4 Hz, 1H, $C≡C\underline{H}$) ppm.

**$^{13}$C NMR** (151 MHz, $CDCl_3$, 298 K) δ = 165.6 ($Ph\underline{C}=O$), 165.3 ($Ph\underline{C}=O$), 136.7 ($\underline{Ph}_{ipso}CH$), 133.2 ($OBz_{para}$), 133.1 ($OBz_{para}$), 129.9 ($OBz_{ortho}$), 129.8 ($OBz_{ortho}$), 129.4 ($OBz_{ipso}$), 129.3 ($OBz_{ipso}$), 129.1 ($Ph_{para}CH$), 128.3 ($Ph_{meta}CH$), 128.3 ($OBz_{meta}$), 128.2 ($OBz_{meta}$), 126.1 ($Ph_{ortho}CH$), 101.5 ($Ph\underline{C}H$), 99.1 (C-1), 78.8 (C-4), 78.1 ($CH_2\underline{C}≡CH$), 75.6 ($CH_2C≡\underline{C}H$), 72.15 (C-2/C-3), 72.07 (C-2/C-3), 68.6 (C-6), 66.7 (C-5), 56.3 ($\underline{C}H_2C≡CH$) ppm.

**ESI-HRMS:** *m/z* = 1051.31337 $[2M+Na]^+$ (calculated *m/z* = 1051.31476).

#### 2.22. Propargyl 2,3-*O*-isopropylidene-β-D-xylopyranoside (S11)

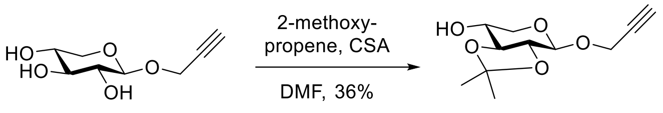

Xyloside **20**[9] (2.00 g, 10.6 mmol) and (±)-camphorsulfonic acid (247 mg, 1.06 mmol) were dissolved in dry DMF (18 mL) under a nitrogen atmosphere. The mixture was heated at 60 °C and 2-methoxypropene (3.05 mL, 31.8 mmol) was added dropwise. After 1 h the mixture was allowed to reach room temperature and was quenched with triethylamine (2.7 mL, 19.4 mmol). The crude mixture was concentrated in vacuo and the residue was co-evaporated with toluene (2 x 40 mL). The yellow crude syrup was loaded onto celite® and purified on silica gel via automated flash chromatography (cyclohexane:ethyl acetate, 9:1→6:4) to obtain the product as a colorless syrup (867 mg, 3.80 mmol, 36%).

$R_F$ (cyclohexane:ethyl acetate, 3:2) = 0.28.

$[\alpha]_D^{20}$ = -67.2 (*c* = 0.27, $CHCl_3$).

**IR (ATR):** $\tilde{\nu}$ = 3439, 3275, 2987, 2934, 2887, 2124, 1374, 1228, 1054 $cm^{-1}$.

**$^1$H NMR** (500 MHz, Acetone-$d_6$, 298 K): δ = 4.84 (d, $^3J_{1,2}$ = 7.4 Hz, 1H, H-1), 4.54 (d, $^3J_{4,OH}$ = 4.8 Hz, 1H, O<u>H</u>), 4.40-4.29 (m, 2H, C<u>H</u>$_2$C≡CH), 3.96 (dd, $^2J_{5ax,5eq}$ = 11.6 Hz, $^3J_{4,5eq}$ = 5.2 Hz, 1H, H-5$_{eq}$), 3.93-3.87 (m, 1H, H-4), 3.54 (dd, $^3J_{2,3}$ = 9.6 Hz, $^3J_{3,4}$ = 8.9 Hz, 1H, H-3), 3.28 (dd, $^3J_{2,3}$ = 9.6 Hz, $^3J_{1,2}$ = 7.4 Hz, 1H, H-2), 3.25 (dd, $^2J_{5ax,5eq}$ = 11.6 Hz, $^3J_{4,5ax}$ = 7.4 Hz, 1H, H-5$_{ax}$), 2.99 (t, $^4J_{CH2,C≡CH}$ = 2.4 Hz, 1H, C≡C<u>H</u>), 1.37 (s, 3H, C(C<u>H</u>$_3$), 1.36 (s, 3H, C(C<u>H</u>$_3$) ppm.

**$^{13}$C NMR** (126 MHz, Acetone-$d_6$, 298 K): δ = 111.3 (<u>C</u>(CH$_3$)$_2$), 100.5 (C-1), 81.9 (C-3), 79.9 (<u>C</u>≡CH), 77.4 (C-2), 76.3 (C≡<u>C</u>H), 69.8 (C-4), 68.6 (C-5), 55.4 (<u>C</u>H$_2$C≡CH), 27.0 (C(<u>C</u>H$_3$), 26.8 (C(<u>C</u>H$_3$) ppm.

**ESI-HRMS:** *m/z* = 229.10673 $[M+H]^+$ (calculated *m/z* = 229.10705 for $[M+H]^+$).

#### 2.23. Propargyl 2,3-di-*O*-benzoyl-4-*O*-benzyl-β-D-xylopyranoside (S12)

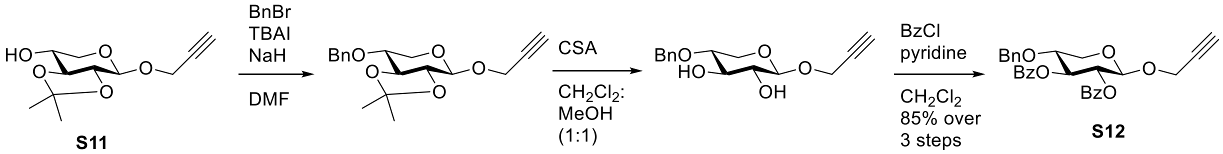

Xyloside **S11** (851 mg, 3.73 mmol) was dissolved in dry DMF (15 mL) and cooled to 0 °C under a nitrogen atmosphere. NaH (60% dispersion in paraffin, 810 mg, 20.2 mmol) was added slowly followed by tetra-*n*-butylammonium iodide (138 mg, 374 µmol). After stirring of the mixture for 10 min at 0 °C benzyl bromide (1.10 mL, 9.26 mmol) was added dropwise. The mixture was further stirred at 0 °C for 2 h. Thin layer chromatography indicated full conversion (cyclohexane:ethyl acetate, 3:2, $R_F$ = 0.81) and the reaction mixture was diluted with ethyl acetate (100 mL) and dropwise quenched with water (20 mL). The aq. phase was separated and the organic phase was washed with brine (3 x 25 mL). Afterwards the organic phase was dried over $MgSO_4$, filtered and the solvent was removed under reduced pressure to obtain the crude intermediate as a yellow syrup. The yellow syrup was dissolved in $CH_2Cl_2$:MeOH (7 mL:7 mL). At 0 °C (±)-camphorsulfonic acid (433 mg, 1.86 mmol) was slowly added. After stirring for 30 min at room temperature thin layer chromatography indicated full consumption of the starting material and the reaction mixture was quenched with triethylamine (3 mL). The reaction mixture was concentrated in vacuo and co-evaporated with toluene. Afterwards, propargyl 4-*O*-benzyl-β-D-xylopyranoside was dissolved in a mixture of dry pyridine and dry $CH_2Cl_2$ (8 mL:2.5 mL) under a nitrogen atmosphere. At 0 °C benzoyl chloride (1.95 mL, 16.8 mmol) was added dropwise. After stirring at room temperature for 8 h thin layer chromatography indicated full conversion to the target molecule. The reaction mixture was cooled to 0°C and quenched with methanol (5 mL). The reaction mixture was then diluted with ethyl acetate (200 mL). The organic phase was washed with 1 M HCl (4 x 25 mL), satd. aq. $NaHCO_3$ solution (2 x 25 mL) and brine (2 x 25 mL). The organic phase was dried over $MgSO_4$, filtered and the solvent was removed under reduced pressure. The crude product was loaded onto celite® and purified on silica gel via automated flash chromatography (cyclohexane:ethyl acetate, 100:0→7:1) to obtain the product as a colorless syrup (1.55 g, 3.19 mmol, 85% over three steps).

$\boldsymbol{R}_{\mathbf{F}}$ (cyclohexane:ethyl acetate, 3:2) = 0.78.

$[\boldsymbol{\alpha}]_{\mathbf{D}}^{\mathbf{20}}$ = +54.9 (*c* = 0.97, $CHCl_3$).

**IR (ATR):** $\tilde{\nu}$ = 3290, 3033, 2871, 2122, 1722, 1451, 1252, 1088, 1069, 707 $cm^{-1}$.

**$^1$H NMR** (500 MHz, $CDCl_3$, 298 K): δ = 8.00-7.94 (m, 4H, 2 x $OBz_{ortho}$), 7.57-7.46 (m, 2H, 2 x $OBz_{para}$), 7.43-7.37 (m, 2H, $OBz_{meta}$), 7.37-7.32 (m, 2H, $OBz_{meta}$), 7.24-7.18 (m, 5H, OBn), 5.62 (t, $^3J_{2,3}$ = $^3J_{3,4}$ = 8.0 Hz, 1H, H-3), 5.28 (dd, $^3J_{2,3}$ = 8.3 Hz, $^3J_{1,2}$ = 6.4 Hz, 1H, H-2), 4.94 (d, $^3J_{1,2}$ = 6.4 Hz, 1H, H-1), 4.65-4.58 (m, 2H, PhC<u>H</u>$_2$), 4.40-4.30 (m, 2H, C<u>H</u>$_2$C≡CH), 4.14 (dd, $^2J_{5ax,5eq}$ = 12.0 Hz, $^3J_{4,5eq}$ = 4.6 Hz, 1H, H-$5_{eq}$), 3.79 (td, $^3J_{3,4}$ = $^3J_{4,5ax}$ = 8.1 Hz, $^3J_{4,5eq}$ = 4.6 Hz, 1H, H-4), 3.57 (dd, $^2J_{5ax,5eq}$ = 12.0 Hz, $^3J_{4,5ax}$ = 8.0 Hz, 1H, H-$5_{ax}$), 2.40 (t, $^4J_{CH2,C≡CH}$ = 2.4 Hz, 1H, C≡C<u>H</u>) ppm.

**$^{13}$C NMR** (126 MHz, $CDCl_3$, 298 K): δ = 165.6 (C=$O_{OBz\ at\ C\text{-}3}$), 165.4 (C=$O_{OBz\ at\ C\text{-}2}$), 137.5 ($OBn_{ipso}$), 133.2 ($OBz_{para}$), 133.1 ($OBz_{para}$), 129.94 ($OBz_{ortho}$), 129.89 ($OBz_{ortho}$), 129.5 ($OBz_{ipso}$), 129.4 ($OBz_{ipso}$), 128.42 (OBn), 128.36 ($OBz_{meta}$), 128.3 ($OBz_{meta}$), 127.92 (OBn), 127.86 (OBn), 98.5 (C-1), 78.5 (<u>C</u>≡CH), 75.2 (C≡<u>C</u>H), 74.3 (C-4), 72.7 (Ph<u>C</u>H$_2$), 72.4 (C-3), 70.8 (C-2), 63.0 (C-5), 55.7 (<u>C</u>H$_2$C≡CH) ppm.

**ESI-HRMS:** *m/z* = 509.15670 $[M+Na]^+$ (calculated *m/z* = 509.15707 for $[M+Na]^+$).

#### 2.24. Propargyl 2,3,4,6-tetra-*O*-acetyl-α-D-mannopyranosyl-(1→4)-2,3-di-*O*-acetyl-6-*O*-benzyl-β-D-glucopyranoside (S13)

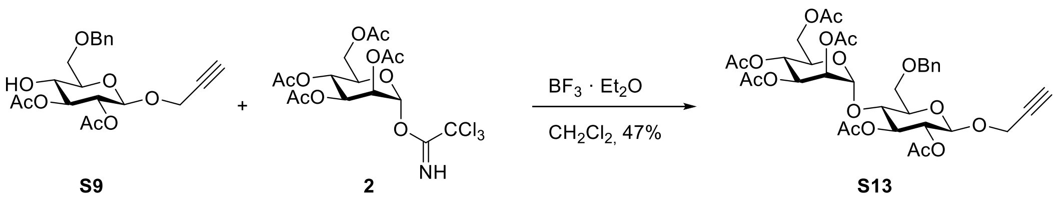

**S9** **2** **S13**

The acceptor glucoside **S9** (1.13 g, 2.87 mmol, 1 equiv) and the mannosyl donor **2**[2] (1.70 g, 3.45 mmol, 1.2 equiv) were dissolved in dry $CH_2Cl_2$ (12 mL) under nitrogen atmosphere, freshly activated molecular sieve (4Å, ~400 mg) was added and the reaction mixture stirred at room temperature for 30 min. At 0 °C boron trifluoride diethyl etherate (400 µL, 3.16 mmol, 1.1 equiv) was added and the reaction mixture was stirred at 0 °C for 2 h, followed by stirring at room temperature for 3 h. Additional solution of donor **2**[2] (327 mg, 664 µmol, 0.23 equiv) in dry $CH_2Cl_2$ (5 mL) was added at 0 °C and stirred for further 2.5 h. The reaction mixture was diluted with $CH_2Cl_2$ (20 mL) and satd. aq. $NaHCO_3$ solution (20 mL) and strongly stirred for 10 min. After filtration over celite® the phases were separated and the aq. phase was extracted with ethyl acetate (150 mL). The combined organic phases were washed with brine (50 mL), dried over $MgSO_4$, it was filtrated and the solvent was removed under reduced pressure. The crude product was purified on silica gel via automated flash chromatography (cyclohexane:ethyl acetate, 6:4) to yield the product as a colorless foam (972 mg, 1.33 mmol, 47%).

$\boldsymbol{R}_F$ (cyclohexane:ethyl acetate, 6:4) = 0.30.

$[\boldsymbol{\alpha}]_{\mathbf{D}}^{\mathbf{20}}$ = -17.7 ($c$ = 0.31 in $CHCl_3$).

**IR (ATR):** $\tilde{\nu}$ = 3277 (w), 2942 (w), 1743 (s), 1432 (w), 1367 (m), 1214 (s), 1134 (m), 1037 (s), 983 (m), 900 (m), 699 (m) $cm^{-1}$.

**$^1$H NMR** (600 MHz, $CDCl_3$, 298 K) δ = 7.37 – 7.27 (m, 5H, OBn), 5.27 (t, $^3J_{2,3}$ = $^3J_{3,4}$ = 9.3 Hz, 1H, H-$3_{Glc}$), 5.24 (t, $^3J_{3,4}$ = $^3J_{4,5}$ = 9.6 Hz, 1H, H-$4_{Man}$), 5.21 (dd, $^3J_{3,4}$ = 9.9 Hz, $^3J_{2,3}$ = 3.1 Hz, 1H, H-$3_{Man}$), 5.05 (dd, $^3J_{2,3}$ = 3.1 Hz, $^3J_{1,2}$ = 2.1 Hz, 1H, H-$2_{Man}$), 5.02 (d, $^3J_{1,2}$ = 2.1 Hz, 1H, H-$1_{Man}$), 4.90 (dd, $^3J_{2,3}$ = 9.5 Hz, $^3J_{1,2}$ = 7.9 Hz, 1H, H-$2_{Glc}$), 4.75 (d, $^3J_{1,2}$ = 7.9 Hz, 1H, H-$1_{Glc}$), 4.66 – 4.56 (m, 2H, PhC$\underline{H}_2$), 4.41 – 4.32 (m, 2H, C$\underline{H}_2$C≡CH), 4.17 (dd, $^2J$ = 12.2 Hz, $^3J_{5,6a}$ = 5.1 Hz, 1H, H-$6a_{Man}$), 4.01 – 3.92 (m, 3H, H-$4_{Glc}$, H-$5_{Man}$, H-$6b_{Man}$), 3.85 – 3.77 (m, 2H, H-$6a_{Glc}$, H-$6b_{Glc}$), 3.61 (ddd, $^3J_{4,5}$ = 9.6 Hz, $^3J_{5,6}$ = 3.9 Hz, $^3J_{5,6}$ = 2.3 Hz, 1H, H-$5_{Glc}$), 2.46 (t, $^4J_{C≡CH,CH2}$ = 2.4 Hz, 1H, C≡C$\underline{H}$), 2.12 (s, 3H, OAc), 2.08 (s, 3H, OAc), 2.05 (s, 6H, 2xOAc), 2.03 (s, 3H, OAc), 1.98 (s, 3H, OAc) ppm.

**$^{13}$C NMR** (151 MHz, $CDCl_3$, 298 K) δ = 170.5 ($CH_3\underline{C}O$), 170.3 ($CH_3\underline{C}O$), 169.8 ($CH_3\underline{C}O$), 169.8 ($CH_3\underline{C}O$), 169.6 (2x$CH_3\underline{C}O$), 137.9 ($OBn_{ipso}$), 128.4 (OBn), 127.8 ($OBn_{para}$), 127.6 (OBn), 99.1 (C-$1_{Man}$), 97.9 (C-$1_{Glc}$), 78.3 ($CH_2\underline{C}$≡CH), 76.3 (C-$4_{Glc}$), 75.4 ($CH_2$C≡$\underline{C}$H), 74.7 (C-$5_{Glc}$), 74.3 (C-$3_{Glc}$), 73.5 (Ph$\underline{C}H_2$), 71.7 (C-$2_{Glc}$), 69.7 (C-$2_{Man}$), 69.6 (C-$5_{Man}$), 68.5 (C-$3_{Man}$), 68.4 (C-$6_{Glc}$), 65.9 (C-$4_{Man}$), 62.4 (C-$6_{Man}$), 55.8 ($\underline{C}H_2$C≡CH), 20.8 (2 x $CH_3$), 20.74 ($CH_3$), 20.71 ($CH_3$), 20.69 ($CH_3$), 20.62 ($CH_3$) ppm.

**ESI-HRMS:** $m/z$ = 740.27484 $[M+NH_4]^+$ (calculated $m/z$ = 740.27603).

#### 2.25. Propargyl 2,3,4,6-tetra-*O*-benzoyl-β-D-glucopyranosyl-(1→4)-6-*O*-acetyl-2,3-di-*O*-benzoyl-β-D-glucopyranoside (S14)

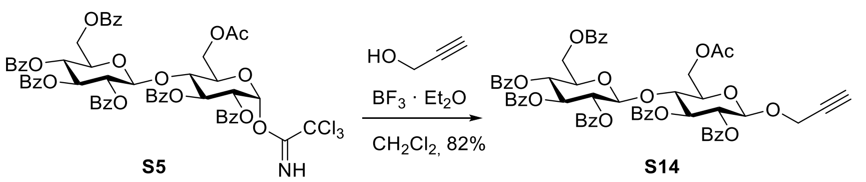

The disaccharide donor **S5**[15] (1.63 g, 1.41 mmol, 1 equiv) was dissolved in dry $CH_2Cl_2$ (3 mL) under nitrogen atmosphere, freshly activated molecular sieve (3Å, ~1 g) and propargyl alcohol (91.7 µL, 1.55 mmol, 1.1 equiv) were added. After stirring at room temperature for 30 min the mixture was cooled to 0 °C. Boron trifluoride diethyl etherate (179 µL, 1.41 mmol, 1 equiv) was added and the reaction mixture was stirred at 0 °C for 2 h, followed by stirring at room temperature for 20 h. The reaction mixture was diluted with $CH_2Cl_2$ (20 mL) and satd. aq. $NaHCO_3$ solution (20 mL) and strongly stirred for 20 min. After filtration over celite® the phases were separated and the aq. phase was extracted with $CH_2Cl_2$ (2 x 20 mL). All organic phases were combined and dried over $MgSO_4$, filtrated and the solvent was removed under reduced pressure. The crude was purified on silica gel via automated flash chromatography (cyclohexane:ethyl acetate, 3:1 → 2:1) to yield the product as a colorless foam (1.22 g, 1.16 mmol, 82%).

***R*F** (toluene:ethyl acetate, 8:2) = 0.65.

$[\alpha]_D^{20}$ = +11.0 (*c* = 0.59 in $CHCl_3$).

**IR (ATR):** $\tilde{\nu}$ = 3278 (w), 3063 (w), 2962 (w), 1720 (s), 1602 (w), 1451 (m), 1315 (m), 1248 (s), 1177 (m), 1090 (s), 1067 (s), 1026 (s), 853 (m), 705 (s) $cm^{-1}$.

**$^1$H NMR** (600 MHz, $CDCl_3$, 298 K) δ = 8.03 – 7.87 (m, 8H, OBz), 7.80 – 7.72 (m, 4H, OBz), 7.60 – 7.32 (m, 12H, OBz), 7.31 – 7.19 (m, 6H, OBz), 5.82 – 5.72 (m, 2H, H-3$_{Glc}$, H-3$_{Glc'}$), 5.50 (dd, $^3J_{2,3}$ = 9.8 Hz, $^3J_{1,2}$ = 7.9 Hz, 1H, H-2$_{Glc'}$), 5.41 (t, $^3J_{3,4}$ = $^3J_{4,5}$ = 9.7 Hz, 1H, H-4$_{Glc'}$), 5.37 (dd, $^3J_{2,3}$ = 9.7 Hz, $^3J_{1,2}$ = 7.9 Hz, 1H, H-2$_{Glc}$), 4.92 (d, $^3J_{1,2}$ = 7.9 Hz, 1H, H-1$_{Glc}$), 4.89 (d, $^3J_{1,2}$ = 7.9 Hz, 1H, H-1$_{Glc'}$), 4.41 (dd, $^2J$ =12.1 Hz, $^3J_{5,6a}$ = 2.0 Hz, 1H, H-6a$_{Glc}$), 4.35 – 4.26 (m, 2H, C<u>H</u>$_2$C≡CH), 4.16 (dd, $^2J$ = 12.1 Hz, $^3J_{5,6b}$ = 4.3 Hz, 1H, H-6b$_{Glc}$), 4.13 – 4.09 (m, 1H, H-4$_{Glc,}$), 4.00 (dd, $^2J$ = 11.8 Hz, $^3J_{5,6a}$ = 3.2 Hz, 1H, H-6a$_{Glc'}$), 3.87 (ddd, $^3J_{4,5}$ = 9.7 Hz, $^3J_{5,6b}$ = 5.1 Hz, $^3J_{5,6a}$ = 3.3 Hz, 1H, H-5$_{Glc'}$), 3.83 (dd, $^2J$ = 11.8 Hz, $^3J_{5,6b}$ = 5.1 Hz, 1H, H-6b$_{Glc'}$), 3.71 (ddd, $^3J_{4,5}$ = 9.9 Hz, $^3J_{5,6b}$ = 4.3 Hz, $^3J_{5,6a}$ = 2.1 Hz, 1H, H-5$_{Glc}$), 2.34 (t, $^4J_{C≡CH,CH2}$ = 2.4 Hz, 1H, C≡C<u>H</u>), 1.97 (s, 3H, OAc) ppm.

**$^{13}$C NMR** (151 MHz, $CDCl_3$, 298 K) δ = 170.3 (CH$_3$<u>C</u>O), 165.7 (Ph<u>C</u>O), 165.7 (Ph<u>C</u>O), 165.3 (Ph<u>C</u>O), 165.3 (Ph<u>C</u>O), 165.0 (Ph<u>C</u>O), 164.7 (Ph<u>C</u>O), 133.5 (OBz$_{para}$), 133.4 (OBz$_{para}$), 133.2 (2 x OBz$_{para}$), 133.2 (OBz$_{para}$), 133.1 (OBz$_{para}$), 129.9 (2 x OBz), 129.7 (OBz), 129.7 (OBz), 129.7 (OBz), 129.61 (OBz), 129.56 (OBz), 129.4 (OBz), 129.4 (OBz), 129.3 (OBz), 128.7 (OBz), 128.6 (OBz), 128.5 (OBz), 128.4 (OBz), 128.3 (2 x OBz), 128.3 (2 x OBz), 101.0 (C-1$_{Glc'}$), 98.1 (C-1$_{Glc}$), 78.0 (CH$_2$<u>C</u>≡CH), 76.4 (C-4$_{Glc}$), 75.5 (CH$_2$C≡<u>C</u>H), 72.9 (C-5$_{Glc}$), 72.85 (C-3$_{Glc}$/C-3$_{Glc'}$), 72.79 (C-3$_{Glc}$/ C-3$_{Glc'}$), 72.3 (C-5$_{Glc'}$), 72.0 (C-2$_{Glc'}$), 71.5 (C-2$_{Glc}$), 69.4 (C-4$_{Glc'}$), 62.6 (C-6$_{Glc'}$), 61.6 (C-6$_{Glc}$), 56.0 (<u>C</u>H$_2$C≡CH), 20.7 ($CH_3$) ppm.

**ESI-HRMS:** *m/z* = 1064.33179 $[M+NH_4]^+$ (calculated *m/z* = 1064.33354).

##### 2.26. Propargyl 2,3,4,6-tetra-*O*-acetyl-β-D-glucopyranosyl-(1→4)-2,3,6-tri-*O*-acetyl-β-D-gluco-pyranosyl-(1→4)-2,3-di-*O*-benzoyl-6-*O*-benzyl-β-D-glucopyranoside (S15)

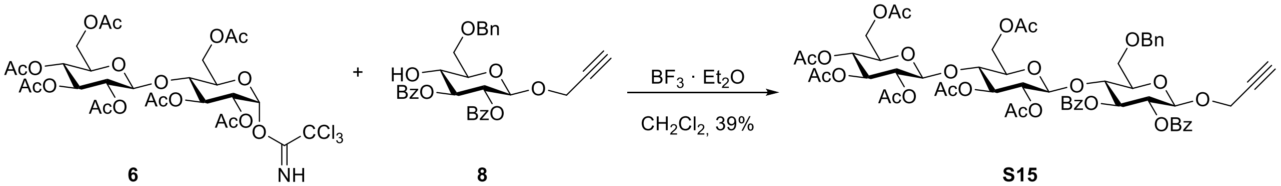

6 8 S15

The disaccharide donor **6**[4] (75 mg, 96.0 µmol, 1.2 equiv) and the acceptor glucoside **8** (42 mg, 81.3 µmol, 1 equiv) were dissolved in dry $CH_2Cl_2$ (5 mL) under nitrogen atmosphere, freshly activated molecular sieve (4Å, ~100 mg) was added and the reaction mixture was stirred at room temperature for 30 min. Then, it was cooled to 0 °C and boron trifluoride diethyl etherate (5 µL, 39.5 µmol, 0.5 equiv) was added. The reaction mixture was stirred at 0 °C for 2 h and further for

17 h at room temperature. It was diluted with $CH_2Cl_2$ (20 mL) and satd. aq. $NaHCO_3$ solution (20 mL) and strongly stirred for 10 min. After filtration over celite® the phases were separated and the aq. phase was extracted with $CH_2Cl_2$ (2 x 20 mL). All organic phases were combined and dried over $MgSO_4$, it was filtrated and the solvent was removed under reduced pressure. The crude was purified on silica gel via automated flash chromatography (cyclohexane:ethyl acetate, 80:20 → 50:50 over 12 CV, then 50:50 over 5 CV) to yield the product as a colorless foam (35.7 mg, 31.5 µmol, 39%).

***R*<sub></sub>**

$\boldsymbol{R_F}$ (cyclohexane:ethyl acetate, 1:1) = 0.39.

$[\alpha]_D^{20}$ = -6.5 ($c$ = 0.43 in $CHCl_3$).

**IR (ATR):** $\tilde{\nu}$ = 3254 (w), 2940 (w), 1736 (m), 1452 (w), 1366 (w), 1217 (s), 1038 (s), 711 (m) $cm^{-1}$.

**$^1$H NMR** (600 MHz, $CDCl_3$, 298 K) δ = 7.95 – 7.89 (m, 4H, 2 x $OBz_{ortho}$), 7.51 – 7.46 (m, 2H, 2 x $OBz_{para}$), 7.44 – 7.29 (m, 9H, OBn, 2 x $OBz_{meta}$), 5.63 (t, $^3J_{2,3}$ = $^3J_{3,4}$ = 9.5 Hz, 1H, H-$3_{Glc}$), 5.37 (dd, $^3J_{2,3}$ = 9.7 Hz, $^3J_{1,2}$ = 7.9 Hz, 1H, H-$2_{Glc}$), 5.08 – 4.98 (m, 2H, H-$3_{Glc''}$, H-$4_{Glc''}$), 4.96 – 4.89 (m, 2H, H-$1_{Glc}$, H-$3_{Glc'}$), 4.86 (dd, $^3J_{2,3}$ = 9.0 Hz, $^3J_{1,2}$ = 7.9 Hz, 1H, H-$2_{Glc''}$), 4.81 (d, $^2J$ = 12.0 Hz, 1H, Ar-C$\underline{H}_A$$H_B$), 4.75 (dd, $^3J_{2,3}$ = 9.6 Hz, $^3J_{1,2}$ = 8.0 Hz, 1H, H-$2_{Glc'}$), 4.52 (d, $^2J$ = 12.0 Hz, 1H, Ar-C$H_A$$\underline{H}_B$), 4.43 (d, $^3J_{1,2}$ = 8.0 Hz, 1H, H-$1_{Glc'}$) 4.45 – 4.33 (m, 2H, C$\underline{H}_2$C≡CH), 4.31 (dd, $^2J$ = 12.4 Hz, $^3J_{5,6a}$ = 4.4 Hz, 1H, H-$6a_{Glc''}$), 4.24 (d, $^3J_{1,2}$ = 7.9 Hz, 1H, H-$1_{Glc''}$), 4.18 (t, $^3J_{3,4}$ = $^3J_{4,5}$ = 9.5 Hz, 1H, H-$4_{Glc}$), 3.96 (m, 2H, H-$6a_{Glc'}$, H-$6b_{Glc''}$), 3.82 (m, 2H, H-$6a_{Glc}$, H-$6b_{Glc}$), 3.67 – 3.60 (m, 1H, H-$5_{Glc}$), 3.53 (ddd, $^3J_{4,5}$ = 9.7 Hz, $^3J_{5,6a}$ = 4.4 Hz, $^3J_{5,6b}$ = 2.2 Hz, 1H, H-$5_{Glc''}$), 3.47 – 3.41 (m, 2H, H-$4_{Glc'}$, H-$6b_{Glc'}$), 3.14 (ddd, $^3J_{4,5}$ = 10.0 Hz, $^3J_{5,6a}$ = 5.8 Hz, $^3J_{5,6b}$ = 1.8 Hz, 1H, H-$5_{Glc'}$), 2.37 (t, $^4J_{C≡CH,CH2}$ = 2.4 Hz, 1H, C≡C$\underline{H}$), 2.06 (s, 3H, OAc), 1.98 (s, 3H, OAc), 1.98 (s, 3H, OAc), 1.96 (s, 3H, OAc), 1.95 (s, 3H, OAc), 1.93 (s, 3H, OAc), 1.91 (s, 3H, OAc) ppm.

**$^{13}$C NMR** (151 MHz, $CDCl_3$, 298 K) δ = 170.6 ($CH_3\underline{C}O$), 170.3 ($CH_3\underline{C}O$), 170.2 ($CH_3\underline{C}O$), 169.9 ($CH_3\underline{C}O$), 169.4 ($CH_3\underline{C}O$), 169.3 ($CH_3\underline{C}O$), 169.2 ($CH_3\underline{C}O$), 165.4 (Ph$\underline{C}$=O), 165.3 (Ph$\underline{C}$=O), 137.7 ($OBn_{ipso}$), 133.2 (2 x $OBz_{para}$), 130.0 ($OBz_{ortho}$), 129.9 ($OBz_{ipso}$), 129.8 ($OBz_{ortho}$), 129.5 ($OBz_{ipso}$), 128.9 ($OBn_{para}$), 128.5 ($OBn_{ortho}$, $OBn_{meta}$), 128.4 ($OBz_{meta}$), 128.4 ($OBz_{meta}$), 100.9 (C-$1_{Glc''}$), 99.8 (C-$1_{Glc'}$), 98.4 (C-$1_{Glc}$), 78.4 ($CH_2\underline{C}$≡CH), 76.3 (C-$4_{Glc'}$), 75.6 ($CH_2$C≡$\underline{C}$H), 75.1 (C-$4_{Glc}$/C-$5_{Glc}$), 75.0 (C-$4_{Glc}$/C-$5_{Glc}$), 74.0 (Ar-$\underline{C}H_2$), 73.2 (C-$3_{Glc}$), 73.0 (C-$3_{Glc'}$/C-$3_{Glc''}$), 72.9 (C-$3_{Glc'}$/C-$3_{Glc''}$), 72.6 (C-$5_{Glc'}$), 72.0 (C-$5_{Glc''}$), 71.93 (C-$2_{Glc'}$), 71.86 (C-$2_{Glc}$), 71.5 (C-$2_{Glc''}$), 67.9 (C-$4_{Glc''}$), 67.2 (C-$6_{Glc}$), 62.3 (C-$6_{Glc'}$), 61.6 (C-$6_{Glc''}$), 56.0 ($\underline{C}H_2$C≡CH), 20.9 ($\underline{C}H_3$CO), 20.8 ($\underline{C}H_3$CO), 20.8 ($\underline{C}H_3$CO), 20.68 (2 x $\underline{C}H_3$CO), 20.62 ($\underline{C}H_3$CO), 20.60 ($\underline{C}H_3$CO) ppm.

**ESI-HRMS:** $m/z$ = 1157.34275 $[M+Na]^+$ (calculated $m/z$ = 1157.34724).

#### 2.27. Propargyl 2,3,4,6-tetra-*O*-acetyl-β-D-glucopyranosyl-(1→4)-2,3,6-tri-*O*-acetyl-β-D-gluco-pyranosyl-(1→4)-2,3-di-*O*-benzoyl-β-D-glucopyranoside (S16)

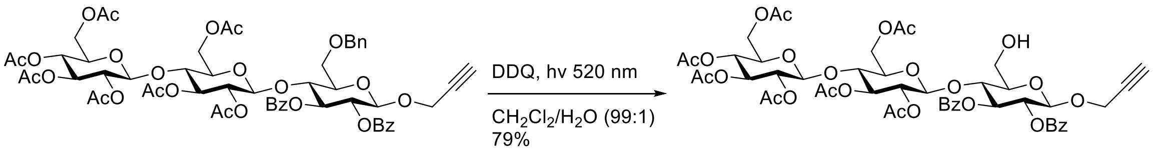

**S15** **S16**

Following the procedure from Cavedon[17] et al. trisaccharide **S15** (32.6 mg, 28.7 µmol, 1 equiv) and 2,3-dichloro-5,6-dicyano-1,4-benzoquinone (9.8 mg, 43.1 µmol, 1.5 equiv) were dissolved in dry $CH_2Cl_2$ (1.5 mL) under nitrogen atmosphere and water (15 µL) was added. During strong stirring the reaction mixture was irradiated with 520 nm green light for 3.5 h at room temperature. The reaction mixture was diluted with $CH_2Cl_2$ (30 mL) and washed with satd. aq. $NaHCO_3$ solution (20 mL). The aq. phase was extracted with $CH_2Cl_2$ (20 mL). All organic phases were combined and dried over $MgSO_4$, filtrated and the solvent was removed under reduced pressure. The crude was purified on silica gel via automated flash chromatography (cyclohexane:ethyl acetate, 65:35 → 40:60 over 12 CV) to yield the product as a colorless foam (23.7 mg, 22.7 µmol, 79%).

$\boldsymbol{R}_F$ (cyclohexane:ethyl acetate, 1:1) = 0.17.

$[\alpha]_D^{20}$= +6.3 ($c$ = 0.28 in $CHCl_3$).

**IR (ATR):** $\tilde{\nu}$ = 3489 (w, br), 3269 (w), 2944 (w), 1732 (s), 1367 (m), 1215 (s), 1032 (s), 710 (m) $cm^{-1}$.

**$^1$H NMR** (600 MHz, $CDCl_3$, 298 K) δ = 8.00 – 7.88 (m, 4H, 2 x $OBz_{ortho}$), 7.57 – 7.46 (m, 2H, 2 x $OBz_{para}$), 7.41 – 7.32 (m, 4H, 2 x $OBz_{meta}$), 5.70 (t, $^3J_{2,3}$ = $^3J_{3,4}$ = 9.5 Hz, 1H, H-$3_{Glc}$), 5.32 (dd, $^3J_{2,3}$ = 9.7 Hz, $^3J_{1,2}$ = 7.9 Hz, 1H, H-$2_{Glc}$), 5.07 (t, $^3J_{2,3}$ = $^3J_{3,4}$ = 9.4 Hz, 1H, H-$3_{Glc'}$), 5.04 (t, $^3J_{2,3}$ = $^3J_{3,4}$ = 9.2 Hz, 1H, H-$3_{Glc''}$), 5.00 (t, $^3J_{3,4}$ = $^3J_{4,5}$ = 9.5 Hz, 1H, H-$4_{Glc''}$), 4.98 (d, $^3J_{1,2}$ = 8.0 Hz, 1H, H-$1_{Glc}$), 4.87 – 4.79 (m, 2H, H-$2_{Glc'}$, H-$2_{Glc''}$), 4.66 (d, $^3J_{1,2}$ = 8.0 Hz, 1H, H-$1_{Glc'}$), 4.40 (dd, $^2J$ = 16.0 Hz, $^4J_{CH2,C≡CH}$ = 2.4 Hz, 1H, C$\underline{H}_A$$H_B$C≡CH), 4.34 (dd, $^2J$ = 16.0 Hz, $^4J_{CH2,C≡CH}$ = 2.3 Hz, 1H, $CH_A$$\underline{H}_B$C≡CH), 4.32 – 4.30 (m, 1H, H-$6a_{Glc''}$), 4.29 (d, $^3J_{1,2}$ = 8.1 Hz, 1H, H-$1_{Glc''}$), 4.16 (t, $^3J_{3,4}$ = $^3J_{4,5}$ = 9.5 Hz, 1H, H-$4_{Glc}$), 4.00 – 3.94 (m, 3H, H-$6a_{Glc}$, H-$6a_{Glc'}$, H-$6b_{Glc''}$), 3.82 (ddd, $^2J_{6b,6a}$ = 12.1 Hz, $^3J_{6b,OH}$ = 8.8 Hz, $^3J_{6b,5}$ = 3.1 Hz, 1H, H-$6b_{Glc}$), 3.62 – 3.51 (m, 4H, H-$4_{Glc'}$, H-$5_{Glc}$, H-$5_{Glc''}$, H-$6b_{Glc'}$), 3.31 (ddd, $^3J_{4,5}$ = 9.9 Hz, $^3J_{5,6a}$ = 4.5 Hz, $^3J_{5,6b}$ = 1.5 Hz, 1H, H-$5_{Glc'}$), 2.39 (t, $^4J_{C≡CH,CH2}$ = 2.4 Hz, 1H, C≡C$\underline{H}$), 2.06 (s, 3H, OAc), 2.01 (s, 3H, OAc), 1.98 (s, 3H, OAc), 1.96 (s, 3H, OAc), 1.95 (s, 6H, 2 x OAc), 1.92 (s, 3H, OAc) ppm.

**$^{13}$C NMR** (151 MHz, $CDCl_3$, 298 K) δ = 170.5 ($CH_3\underline{C}O$), 170.2 ($CH_3\underline{C}O$), 170.0 ($CH_3\underline{C}O$), 169.8 ($CH_3\underline{C}O$), 169.4 ($CH_3\underline{C}O$), 169.3 ($CH_3\underline{C}O$), 168.9 ($CH_3\underline{C}O$), 165.3 (Ph$\underline{C}$O), 165.0 (Ph$\underline{C}$O), 133.3 ($OBz_{para}$), 133.2 ($OBz_{para}$), 129.9 ($OBz_{ortho}$), 129.6 ($OBz_{ortho}$), 129.5 ($OBz_{ipso}$), 129.3 ($OBz_{ipso}$), 128.4 ($OBz_{meta}$), 128.3 ($OBz_{meta}$), 100.6 (C-$1_{Glc''}$), 100.3 (C-$1_{Glc'}$), 98.6 (C-$1_{Glc}$), 78.2 ($CH_2\underline{C}$≡CH), 75.8 (C-$4_{Glc'}$), 75.6 ($CH_2$C≡$\underline{C}$H), 75.2 (C-$5_{Glc}$), 75.1 (C-$4_{Glc}$), 73.1 (C-$3_{Glc}$), 72.9 (C-$3_{Glc''}$), 72.7 (C-$5_{Glc'}$), 72.6 (C-$3_{Glc'}$), 71.9 (C-$5_{Glc''}$), 71.9 (C-$2_{Glc'}$), 71.8 (C-$2_{Glc}$), 71.4 (C-$2_{Glc''}$), 67.8 (C-$4_{Glc''}$), 61.8 (C-$6_{Glc'}$), 61.5 (C-$6_{Glc''}$), 60.4 (C-$6_{Glc}$), 56.3 ($\underline{C}H_2$C≡CH), 20.8 ($\underline{C}H_3CO$), 20.7 ($\underline{C}H_3CO$), 20.6 ($\underline{C}H_3CO$), 20.53 ($\underline{C}H_3CO$), 20.53 ($\underline{C}H_3CO$), 20.47 ($\underline{C}H_3CO$), 20.4 ($\underline{C}H_3CO$) ppm.

**ESI-HRMS:** $m/z$ = 1062.34426 $[M+NH_4]^+$ (calculated $m/z$ = 1062.34489).

##### 2.28. Propargyl 6-azido-2,3,4-tri-*O*-benzoyl-6-deoxy-α-D-glucopyranosyl-(1→4)-6-*O*-acetyl-2,3-di-*O*-benzoyl-β-D-glucopyranosyl-(1→4)-2,3-di-*O*-benzoyl-6-*O*-benzyl-β-D-glucopyranoside (S17)

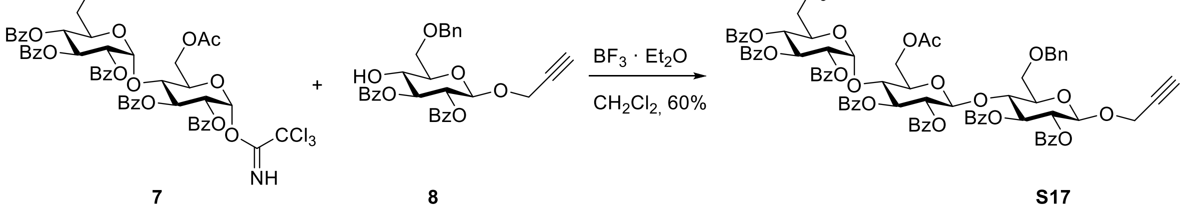

**7** **8** **S17**

The disaccharide donor **7**[5] (160 mg, 149 µmol, 1 equiv) and the acceptor glucoside **8** (78.5 mg, 152 µmol, 1.02 equiv) were dissolved in dry $CH_2Cl_2$ (5 mL) under nitrogen atmosphere, freshly activated molecular sieve (4Å, ~100 mg) was added and the reaction mixture was stirred at room temperature for 30 min. At 0 °C boron trifluoride diethyl etherate (5 µL, 39.5 µmol, 0.26 equiv) was added and the reaction mixture was stirred at 0 °C for 2 h, followed by stirring at room temperature for 14.5 h. The reaction mixture was diluted with $CH_2Cl_2$ (20 mL) and satd. aq. $NaHCO_3$ solution (20 mL) and strongly stirred for 10 min. After filtration over celite® the phases were separated and the aq. phase was extracted with $CH_2Cl_2$ (2 x 20 mL). All organic phases were combined and dried over $MgSO_4$, filtrated and the solvent was removed under reduced pressure. The crude was purified on silica gel via automated flash chromatography (cyclohexane:ethyl acetate, 85:15 → 70:30 over 18 CV) to yield the product as a colorless foam (128 mg, 89.6 µmol, 60%).

***R*$_F$** (toluene:ethyl acetate, 9:1) = 0.47.

$[\boldsymbol{\alpha}]_{\mathbf{D}}^{\mathbf{20}}$= +9.7 (*c* = 0.40 in $CHCl_3$).

**IR (ATR):** $\tilde{\nu}$ = 3297 (w), 3066 (w), 2927 (w), 2105 (m), 1727 (s), 1602 (m), 1452 (m), 1255 (s), 1090 (s), 1067 (s), 1026 (s), 705 (s) $cm^{-1}$.

**$^1$H NMR** (500 MHz, $CDCl_3$, 298 K) δ = 8.02 – 7.97 (m, 2H, ArC$\underline{H}$), 7.93 – 7.89 (m, 4H, ArC$\underline{H}$), 7.78 – 7.67 (m, 6H, ArC$\underline{H}$), 7.58 – 7.28 (m, 23H, ArC$\underline{H}$), 7.25 – 7.16 (m, 5H, ArC$\underline{H}$), 5.95 (dd, $^3J$ = 10.5 Hz, $^3J$ = 9.5 Hz, 1H, H-3$_{Glc''}$), 5.63 (t, $^3J_{2,3}$ = $^3J_{3,4}$ = 9.5 Hz, 1H, H-3$_{Glc}$), 5.51 (d, $^3J_{1,2}$ = 3.9 Hz, 1H, H-1$_{Glc''}$), 5.49 – 5.42 (m, 2H, H-3$_{Glc'}$, H-4$_{Glc''}$), 5.38 (dd, $^3J_{2,3}$ = 9.7 Hz, $^3J_{1,2}$ = 8.0 Hz, 1H, H-2$_{Glc}$), 5.17 – 5.10 (m, 2H, H-2$_{Glc'}$, H-2$_{Glc''}$), 4.86 (d, $^3J_{1,2}$ = 8.0 Hz, 1H, H-1$_{Glc}$), 4.76 (d, $^2J_{PhC\underline{H}H,PhCH\underline{H}}$ = 12.1 Hz, 1H, PhC$\underline{H}_A$H$_B$), 4.63 (d, $^3J_{1,2}$ = 7.9 Hz, 1H, H-1$_{Glc'}$), 4.38 (d, $^2J_{PhCH\underline{H},PhC\underline{H}H}$ = 12.1 Hz, 1H, PhCH$_A\underline{H}_B$), 4.34 (dd, $^2J$ = 16.0 Hz, $^4J_{CH2,C≡CH}$ = 2.4 Hz, 1H, C$\underline{H}_A$H$_B$C≡CH), 4.30 (dd, $^2J$ = 16.0 Hz, $^4J_{CH2,C≡CH}$ = 2.5 Hz, 1H, CH$_A\underline{H}_B$C≡CH), 4.25 (t, $^3J_{3,4}$ = $^3J_{4,5}$ = 9.5 Hz, 1H, H-4$_{Glc}$), 4.12 – 4.03 (m, 2H, H-5$_{Glc''}$, H-6a$_{Glc'}$), 3.97 (t, $^3J_{3,4}$ = $^3J_{4,5}$ = 9.2 Hz, 1H, H-4$_{Glc'}$), 3.73 (dd, $^2J_{6b,6a}$ = 11.9 Hz, $^3J_{5,6b}$ = 5.1 Hz, 1H, H-6b$_{Glc'}$), 3.66 (dd, $^2J_{6a,6b}$ = 11.0 Hz, $^3J_{5,6a}$ = 3.1 Hz, 1H, H-6a$_{Glc}$), 3.57 – 3.53 (m, 1H, H-6b$_{Glc}$), 3.51 – 3.39 (m, 3H, H-5$_{Glc}$, H-5$_{Glc'}$, H-6a$_{Glc''}$), 3.34 (dd, $^2J_{6b,6a}$ = 13.5 Hz, $^3J_{5,6b}$ = 5.2 Hz, 1H, H-6b$_{Glc''}$), 2.31 (t, $^4J_{C≡CH,CH2}$ =2.4, 1H, C≡C$\underline{H}$), 2.01 (s, 3H, OAc) ppm.

**$^{13}$C NMR** (126 MHz, $CDCl_3$, 298 K) δ = 170.5 ($CH_3\underline{C}O$), 165.6 (Ph$\underline{C}$O), 165.4 (Ph$\underline{C}$O), 165.3 (Ph$\underline{C}$O), 165.2 (2 x Ph$\underline{C}$O), 164.8 (Ph$\underline{C}$O), 164.6 (Ph$\underline{C}$O), 137.7 (Bn$_{ipso}$), 133.6 (OBz$_{para}$), 133.2 (OBz$_{para}$), 133.11 (OBz$_{para}$), 133.09 (2 x OBz$_{para}$), 133.06 (OBz$_{para}$), 133.0 (OBz$_{para}$), 129.93, 129.90, 129.8, 129.7, 129.6, 129.5, 129.4, 129.0, 128.9, 128.8, 128.7, 128.6, 128.49, 128.46, 128.3, 128.2, 128.17, 128.1, 128.0 (40 C, OBz, OBn), 99.8 (C-1$_{Glc'}$), 98.2 (C-1$_{Glc}$), 96.6 (C-1$_{Glc''}$), 78.2 ($CH_2\underline{C}$≡CH), 75.3 ($CH_2$C≡$\underline{C}$H), 74.8 (C-3$_{Glc'}$, C-4$_{Glc}$), 74.7 (C-5$_{Glc}$), 74.0 (C-4$_{Glc'}$), 73.7 (Ph$\underline{C}H_2$), 73.1 (C-3$_{Glc}$), 72.3 (C-5$_{Glc'}$), 72.2 (C-2$_{Glc'/Glc''}$), 71.7 (C-2$_{Glc}$), 70.8 (C-2$_{Glc'/Glc''}$), 70.2 (C-5$_{Glc''}$), 69.5 (C-4$_{Glc''}$/C-3$_{Glc''}$), 69.4 (C-4$_{Glc''}$/C-3$_{Glc''}$), 67.0 (C-6$_{Glc}$), 63.3 (C-6$_{Glc'}$), 55.8 ($\underline{C}H_2$C≡CH), 50.9 (C-6$_{Glc''}$), 20.9 ($\underline{C}H_3$CO) ppm.

**ESI-HRMS:** *m/z* = 1445.46423 $[M+NH_4]^+$ (calculated *m/z* = 1445.46601).

#### 2.29. Propargyl 6-azido-2,3,4-tri-*O*-benzoyl-6-deoxy-α-D-glucopyranosyl-(1→4)-6-*O*-acetyl-2,3-di-*O*-benzoyl-β-D-glucopyranosyl-(1→4)-2,3-di-*O*-benzoyl-β-D-glucopyranoside (S18)

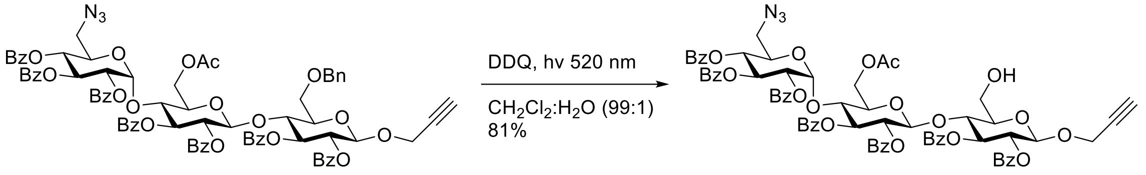

**S17** **S18**

Following the procedure from Cavedon[17] et al. trisaccharide **S17** (128 mg, 89.6 µmol, 1 equiv) and 2,3-dichloro-5,6-dicyano-1,4-benzoquinone (30.5 mg, 134 µmol, 1.50 equiv) were dissolved in dry $CH_2Cl_2$ (4.5 mL) under nitrogen atmosphere and water (45 µL) was added. During strong stirring the reaction mixture was irradiated with 520 nm green light for 6 h at room temperature. The reaction mixture was diluted with $CH_2Cl_2$ (60 mL) and washed with satd. aq. $NaHCO_3$ solution (50 mL). The aq. phase was extracted with $CH_2Cl_2$ (2 x 50 mL). All organic phases were combined, washed with brine and dried over $MgSO_4$, filtrated and the solvent was removed under reduced pressure. The crude was purified on silica gel via automated flash chromatography (cyclohexane:ethyl acetate, 70:30 → 55:45 over 11 CV) to yield the product as a colorless foam (97.6 mg, 72.9 µmol, 81%).

***R*$_F$** (toluene:ethyl acetate, 8:2) = 0.35.

$[\boldsymbol{\alpha}]_{\mathbf{D}}^{\mathbf{20}}$= +12.4 (*c* = 0.47 in $CHCl_3$).

**IR (ATR):** $\tilde{\nu}$ = 3449 (w), 3298 (w), 2956 (w), 2105 (m), 1725 (s), 1602 (m), 1451 (m), 1250 (s), 1090 (s), 1067 (s), 1025 (s), 705 (s) $cm^{-1}$.

**$^1$H NMR** (500 MHz, $CDCl_3$, 298 K) δ = 8.03 – 7.98 (m, 2H, OBz), 7.94 – 7.88 (m, 4H, OBz), 7.87 – 7.81 (m, 2H, OBz), 7.70 – 7.66 (m, 2H, OBz), 7.66 – 7.60 (m, 2H, OBz), 7.55 – 7.45 (m, 6H, OBz), 7.44 – 7.30 (m, 11H, OBz), 7.25 – 7.14 (m, 6H, OBz), 5.90 (dd, $^3J_{2,3}$ = 10.5 Hz, $^3J_{3,4}$ = 9.5 Hz, 1H, H-3$_{Glc''}$), 5.69 (t, $^3J_{2,3}$ = $^3J_{3,4}$ = 9.5 Hz, 1H, H-3$_{Glc}$), 5.62 (dd, $^3J_{2,3}$ = 9.5 Hz, $^3J_{3,4}$ = 8.7 Hz, 1H, H-3$_{Glc'}$), 5.53 (d, $^3J_{1,2}$ = 3.9 Hz, 1H, H-1$_{Glc''}$), 5.43 (t, $^3J_{3,4}$ = $^3J_{4,5}$ = 9.8 Hz, 1H, H-4$_{Glc''}$), 5.35 (dd, $^3J_{2,3}$ = 9.8 Hz, $^3J_{1,2}$ = 8.0 Hz, 1H, H-2$_{Glc}$), 5.25 (dd, $^3J_{2,3}$ = 9.5 Hz, $^3J_{1,2}$ = 7.8 Hz, 1H, H-2$_{Glc'}$), 5.17 (dd, $^3J_{2,3}$ = 10.5 Hz, $^3J_{1,2}$ = 3.9 Hz, 1H, H-2$_{Glc''}$), 4.93 (d, $^3J_{1,2}$ = 7.8 Hz, 1H, H-1$_{Glc'}$), 4.92 (d, $^3J_{1,2}$ = 7.9 Hz, 1H, H-1$_{Glc}$), 4.35 – 4.26 (m, 2H, C$\underline{H}_2$C≡CH), 4.23 (t, $^3J_{3,4}$ = $^3J_{4,5}$ = 9.5 Hz, 1H, H-4$_{Glc}$), 4.14 – 4.01 (m, 3H, H-4$_{Glc'}$, H-5$_{Glc''}$, H-6a$_{Glc'}$), 3.86 – 3.69 (m, 4H, H-5$_{Glc'}$, H-6a$_{Glc}$, H-6b$_{Glc}$, H-6b$_{Glc'}$), 3.46 – 3.39 (m, 2H, H-5$_{Glc}$, H-6a$_{Glc''}$), 3.34 (dd, $^2J$ = 13.5 Hz, $^3J_{5,6b}$ = 5.2 Hz, 1H, H-6b$_{Glc''}$), 2.34 (t, $^4J_{C≡CH,CH2}$ = 2.4 Hz, 1H, C≡C$\underline{H}$), 2.01 (s, 1H, OAc), 1.94 – 1.88 (m, 1H, OH) ppm.

**$^{13}$C NMR** (126 MHz, $CDCl_3$, 298 K) δ = 170.5 ($CH_3\underline{C}O$), 165.5 (Ph$\underline{C}$O), 165.35 (Ph$\underline{C}$O), 165.28 (2 x Ph$\underline{C}$O), 165.18 (Ph$\underline{C}$O), 164.82 (Ph$\underline{C}$O), 164.77 (Ph$\underline{C}$O), 133.6 (OBz$_{para}$), 133.21 (OBz$_{para}$), 133.18 (2 x OBz$_{para}$), 133.13 (OBz$_{para}$), 133.11 (OBz$_{para}$), 133.06 (OBz$_{para}$), 129.9, 129.8, 129.70, 129.67, 129.6, 129.5, 129.3, 129.0, 128.8, 128.55, 128.47, 128.4, 128.3, 128.24, 128.22, 128.1, 128.0 (35 C, OBz), 100.4 (C-1$_{Glc'}$), 98.5 (C-1$_{Glc}$), 96.8 (C-1$_{Glc''}$), 78.1 ($CH_2\underline{C}$≡CH), 75.5 ($CH_2$C≡$\underline{C}$H), 75.14 (C-4$_{Glc}$/C-5$_{Glc}$), 75.09 (C-4$_{Glc}$/C-5$_{Glc}$), 74.8 (C-3$_{Glc'}$), 74.1 (C-4$_{Glc'}$), 72.9 (C-3$_{Glc}$), 72.7 (C-5$_{Glc'}$), 72.3 (C-2$_{Glc'}$), 71.7 (C-2$_{Glc}$), 70.6 (C-2$_{Glc''}$), 70.2 (C-5$_{Glc''}$), 69.6 (C-3$_{Glc''}$/C-4$_{Glc''}$), 69.5 (C-3$_{Glc''}$/C-4$_{Glc''}$), 63.1 (C-6$_{Glc'}$), 60.2 (C-6$_{Glc}$), 56.2 ($\underline{C}H_2$C≡CH), 50.9 (C-6$_{Glc''}$), 20.9 ($\underline{C}H_3$CO) ppm.

**ESI-HRMS:** *m/z* = 1360.37328 $[M+Na]^+$ (calculated *m/z* = 1360.37446).

#### 2.30. Propargyl 6-azido-6-deoxy-α-D-glucopyranosyl-(1→4)-β-D-glucopyranosyl-(1→4)-β-D-glucopyranoside (S19)

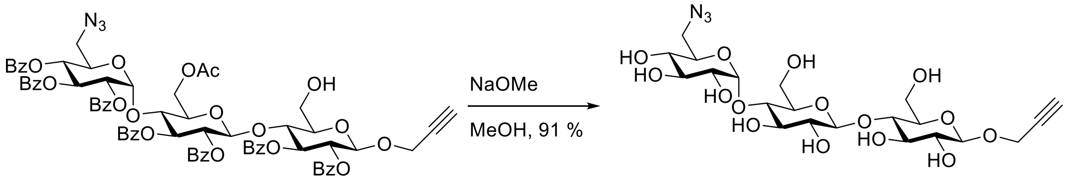

**S18** **S19**

The protected trisaccharide **S18** (96.6 mg, 72.2 µmol, 1 equiv) was dissolved in methanol (5 mL), a solution of sodium methoxide in methanol (25 µL, 125 µmol, 5 M, 1.7 equiv) was added and the reaction mixture stirred at room temperature for 21 h. It was neutralized by addition of Amberlite®-IRC 120 $H^+$ resin and stirring for 15 min. The resin was filtered off and the filtrate was concentrated under reduced pressure. The crude was purified on reversed phase silica gel via automated flash chromatography ($H_2O$:MeCN, 100:0 1 CV, 100:0 → 0:100 over 16 CV) to obtain the title compound after lyophilization (37.4 mg, 65.9 µmol, 91%).

$[\alpha]_D^{20}$= +9.3 (*c* = 0.03 in $H_2O$).

**IR (ATR):** $\tilde{\nu}$ = 3349 (br, m), 2925 (w), 2110 (w), 1644 (w), 1366 (m), 1048 (s) $cm^{-1}$.

**$^1$H NMR** (600 MHz, methanol-d$_4$, 298 K) δ = 5.18 (d, $^3J_{1,2}$ = 3.8 Hz, 1H, H-1$_{Glc''}$), 4.49 (d, $^3J_{1,2}$ = 7.8 Hz, 1H, H-1$_{Glc}$), 4.43 (d, $^3J_{1,2}$ = 7.9 Hz, 1H, H-1$_{Glc'}$), 4.45 – 4.37 (m, 2H, C<u>H</u>$_2$C≡CH), 3.95 (dd, $^2J$ = 12.0 Hz, $^3J_{5,6a}$ = 2.2 Hz, 1H, H-6a$_{Glc'}$), 3.90 (dd, $^2J$ = 12.2 Hz, $^3J_{5,6a}$ = 2.5 Hz, 1H, H-6a$_{Glc}$), 3.86 (dd, $^2J$ = 12.2 Hz, $^3J_{5,6b}$ = 4.1 Hz, 1H, H-6b$_{Glc}$), 3.82 – 3.75 (m, 2H, H-5$_{Glc''}$, H-6b$_{Glc'}$), 3.63 (t, $^3J_{2,3}$ = $^3J_{3,4}$ = 9.1 Hz, 1H, H-3$_{Glc'}$), 3.61 – 3.50 (m, 5H, H-3$_{Glc}$, H-3$_{Glc''}$, H-4$_{Glc}$, H-4$_{Glc'}$, H-6a$_{Glc''}$), 3.48 – 3.43 (m, 2H, H-2$_{Glc''}$, H-5$_{Glc'}$), 3.43 – 3.38 (m, 2H, H-5$_{Glc}$, H-6b$_{Glc''}$), 3.29 – 3.24 (m, 3H, H-2$_{Glc}$, H-2$_{Glc'}$, H-4$_{Glc''}$) ppm.

**$^1$H NMR*** (200 MHz, methanol-d$_4$, 298 K) δ = 2.86 (t, $^4J_{C≡CH,CH2}$ = 2.4 Hz, 1H, C≡C<u>H</u>) ppm.

*Signal for $CH_2$C≡C<u>H</u> was only observed directly after dissolving in methanol-d$_4$, due to complete H/D exchange over a period of two days.

**$^{13}$C NMR** (151 MHz, methanol-d$_4$, 298 K) δ = 104.4 (C-1$_{Glc'}$), 102.9 (C-1$_{Glc''}$), 101.9 (C-1$_{Glc}$), 81.3**, 80.4**, 79.5*** ($CH_2$<u>C</u>≡CH), 77.6 (C-3$_{Glc'}$), 76.7 (C-5$_{Glc'}$), 76.6 (C-5$_{Glc}$), 76.6*** ($CH_2$C≡<u>C</u>H), 76.4**, 74.8**, 74.6 (C-2$_{Glc/Glc'}$), 74.5 (C-2$_{Glc/Glc'}$), 74.1 (C-2$_{Glc''}$), 73.8 (C-5$_{Glc''}$), 72.2 (C-4$_{Glc''}$), 62.1 (C-6$_{Glc'}$), 61.7 (C-6$_{Glc}$), 56.6 (<u>C</u>H$_2$C≡CH), 53.0 (C-6$_{Glc''}$) ppm.

**Assignment is ambiguous due to signal overlap (C-3$_{Glc}$, C-3$_{Glc''}$, C-4$_{Glc}$, C-4$_{Glc'}$).

***Chemical shifts were deduced from HMBC-NMR due to complete H/D exchange of $CH_2$C≡C<u>H</u>.

**ESI-HRMS:** *m/z* = 590.17989 $[M+Na]^+$ (calculated *m/z* = 590.18038).

#### 2.31. Propargyl 2,3,4-tri-*O*-acetyl-β-D-xylopyranosyl-(1→4)-2,3-di-*O*-benzoyl-β-D-xylopyranoside (S20)

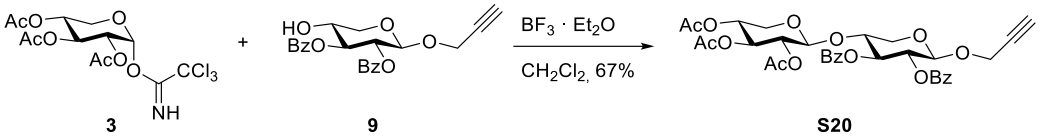

**3** **9** **S20**

The xylosyl donor **3**[3] (167 mg, 397 µmol) and acceptor xyloside **9** (119 mg, 300 µmol) were dissolved in dry $CH_2Cl_2$ (4 mL) under a nitrogen atmosphere, freshly activated powdered molecular sieves 4Å was added and the reaction mixture was stirred at room temperature for 15 min. At 0°C boron trifluoride diethyl etherate (10 µL, 79 µmol) was added. The reaction mixture was stirred at 0 °C for 2 h followed by 12 h at room temperature. The reaction mixture was quenched with a satd. aq. $NaHCO_3$ solution (1 mL) and diluted with $CH_2Cl_2$. After filtration over celite® the organic phase (100 mL) was washed with a satd. aq. $NaHCO_3$ solution (15 mL) and the aq. phase was extracted with $CH_2Cl_2$ (15 mL). All organic phases were combined and dried over $MgSO_4$, filtrated and the solvent was removed under reduced pressure. The crude product was purified on silica gel via automated flash chromatography (cyclohexane:ethyl acetate, 85:15→60:40) to obtain the product as a colorless foam (131 mg, 200 µmol, 67%).

$\boldsymbol{R}_F$ (cyclohexane:ethyl acetate, 1:1) = 0.51.

$[\alpha]_D^{20}$ = -23.5 (*c* = 0.50, $CHCl_3$).

**IR (ATR):** $\tilde{\nu}$ = 3276, 2944, 2872, 1727, 1247, 1216, 1028, 708 $cm^{-1}$.

**$^1$H NMR** (500 MHz, $CDCl_3$, 298 K): δ = 8.01-7.96 (m, 4H, 2 x $OBz_{ortho}$), 7.55-7.49 (m, 2H, 2 x $OBz_{para}$), 7.42-7.37 (m, 4H, 2 x $OBz_{meta}$), 5.61 (t, $^3J_{2,3}$ = $^3J_{3,4}$ = 8.1 Hz, 1H, H-3), 5.29 (dd, $^3J_{2,3}$ = 8.3 Hz, $^3J_{1,2}$ = 6.4 Hz, 1H, H-2), 5.01 (t, $^3J_{2,3}$ = $^3J_{3,4}$ = 7.4 Hz, 1H, H-3'), 4.95 (d, $^3J_{1,2}$ = 6.4 Hz, 1H, H-1), 4.80 (dd, $^3J_{2,3}$ = 7.5 Hz, $^3J_{1,2}$ = 5.7 Hz, 1H, H-2'), 4.67-4.61 (m, 2H, H-1', H-4'), 4.41-4.31 (m, 2H, C<u>H</u>$_2$C≡CH), 4.14 (dd, $^2J_{5ax,5eq}$ = 12.0 Hz, $^3J_{4,5eq}$ = 4.7 Hz, 1H, H-$5_{eq}$), 4.07-4.01 (m, 1H, H-4), 3.73 (dd, $^2J_{5ax,5eq}$ = 12.2 Hz, $^3J_{4,5eq}$ = 4.5 Hz, 1H, H-$5_{eq}$'), 3.56 (dd, $^2J_{5ax,5eq}$ = 12.1 Hz, $^3J_{4,5ax}$ = 8.3 Hz, 1H, H-$5_{ax}$), 3.15 (dd, $^2J_{5ax,5eq}$ = 12.2 Hz, $^3J_{4,5ax}$ = 7.3 Hz, 1H, H-$5_{ax}$'), 2.41 (t, $^4J_{CH2,C≡CH}$ = 2.4 Hz, 1H, C≡C<u>H</u>), 2.04 (s, 3H, $CH3_{OAc\ at\ C\text{-}2'}$), 2.00 (s, 6H, $CH3_{OAc\ at\ C\text{-}3'}$, $CH3_{OAc\ at\ C\text{-}4'}$) ppm.

**$^{13}$C NMR** (126 MHz, $CDCl_3$, 298 K): δ = 170.0 (C=$O_{OAc\ at\ C\text{-}3'}$), 169.7 (C=$O_{OAc\ at\ C\text{-}4'}$), 169.2 (C=$O_{OAc\ at\ C\text{-}2'}$), 165.3 (C=$O_{OBz\ at\ C\text{-}3}$, C=$O_{OBz\ at\ C\text{-}2}$), 133.21 ($OBz_{para}$), 133.18 ($OBz_{para}$), 129.9 ($OBz_{ortho}$), 129.7 ($OBz_{ortho}$), 129.5 ($OBz_{ipso}$), 129.4 ($OBz_{ipso}$), 128.4 ($OBz_{meta}$), 128.3 ($OBz_{meta}$), 99.4 (C-1'), 98.5 (C-1), 78.3(<u>C</u>≡CH), 75.3 (C≡<u>C</u>H), 74.5 (C-4), 71.8 (C-3), 70.6 (C-2), 70.1 (C-3'), 70.0 (C-2'), 68.0 (C-4'), 62.4 (C-5), 61.0 (C-5'), 55.7 (<u>C</u>H$_2$C≡CH), 20.71 ($CH3_{OAc}$), 20.66 ($CH3_{OAc}$), 20.6 ($CH3_{OAc}$) ppm.

**ESI-HRMS:** *m/z* = 672.22812 $[M+NH_4]^+$ (calculated *m/z* = 672.22868 for $[M+NH_4]^+$).

#### 2.32. Propargyl 4-azido-2,3-di-*O*-benzoyl-4-deoxy-xylopyranosyl-(1→4)-2,3-di-*O*-benzoyl-β-D-xylopyranoside (S21)

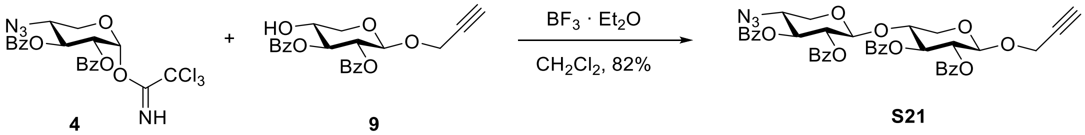

**4** **9** **S21**

The xylosyl donor **4** (145 mg, 275 µmol) and acceptor xyloside **9** (100 mg, 252 µmol) were dissolved in dry $CH_2Cl_2$ (2.0 mL) under a nitrogen atmosphere, freshly activated powdered molecular sieves 4Å was added and the reaction mixture was stirred at room temperature for 15 min. At 0°C boron trifluoride diethyl etherate (8 µL, 63.2 µmol) was added. The reaction mixture was stirred at 0 °C for 1 h followed by 90 min at room temperature. The reaction mixture was quenched with a satd. aq. $NaHCO_3$ solution (150 µL) and diluted with $CH_2Cl_2$ (50 mL). The mixture was dried over $MgSO_4$, filtrated and the solvent was removed under reduced pressure. The crude product was purified on silica gel via automated flash chromatography (toluene:ethyl acetate, 100:0→9:1) to obtain the product as a colorless foam (158 mg, 207 µmol, 82%).

***R*$_F$** (toluene:ethyl acetate, 9:1) = 0.46.

$[\alpha]_D^{20}$ = +16.6 (*c* = 0.38, $CHCl_3$).

**IR (ATR):** $\tilde{\nu}$ = 3290, 2949, 2108, 1721, 1451, 1251, 1068, 706 $cm^{-1}$.

**$^1$H NMR** (600 MHz, $CDCl_3$, 298 K): δ = 8.02-7.97 (m, 4H, 2 x $OBz_{ortho}$), 7.96-7.92 (m, 4H, 2 x $OBz_{ortho}$), 7.57-7.50 (m, 4H, 4 x $OBz_{para}$), 7.43-7.37 (m, 8H, 4 x $OBz_{meta}$), 5.64 (dd, $^3J_{2,3}$ = 8.4 Hz, $^3J_{3,4}$ = 7.7 Hz, 1H, H-3), 5.40 (t, $^3J_{2,3}$ = $^3J_{3,4}$ = 8.5 Hz, 1H, H-3'), 5.27 (dd, $^3J_{2,3}$ = 8.4 Hz, $^3J_{1,2}$ = 6.6 Hz, 1H, H-2), 5.20 (dd, $^3J_{2,3}$ = 8.4 Hz, $^3J_{1,2}$ = 6.4 Hz, 1H, H-2'), 4.89 (d, $^3J_{1,2}$ = 6.5 Hz, 1H, H-1), 4.75 (d, $^3J_{1,2}$ = 6.4 Hz, 1H, H-1'), 4.35-4.27 (m, 2H, C<u>H</u>$_2$C≡CH), 4.06-3.95 (m, 2H, H-4, H-5$_{eq}$), 3.65 (dd, $^2J_{5ax,5eq}$ = 12.2 Hz, $^3J_{4,5eq}$ = 4.9 Hz, 1H, H-5$_{eq}$'), 3.60-3.54 (m, 1H, H-4'), 3.46 (dd, $^2J_{5ax,5eq}$ = 12.1 Hz, $^3J_{4,5ax}$ = 8.4 Hz, 1H, H-5$_{ax}$), 3.15 (dd, $^2J_{5ax,5eq}$ = 12.2 Hz, $^3J_{4,5ax}$ = 9.1 Hz, 1H, H-5$_{ax}$'), 2.37 (t, $^4J_{CH2,C≡CH}$ = 2.4 Hz, 1H, C≡C<u>H</u>) ppm.

**$^{13}$C NMR** (151 MHz, $CDCl_3$, 298 K): δ = 165.6 (C=O$_{OBz\ at\ C\text{-}3'}$), 165.5 (C=O$_{OBz\ at\ C\text{-}3}$), 165.4 (C=O$_{OBz\ at\ C\text{-}2}$), 165.2 (C=O$_{OBz\ at\ C\text{-}2'}$), 133.7 ($OBz_{para}$), 133.6 ($OBz_{para}$), 133.34 ($OBz_{para}$), 133.29 ($OBz_{para}$), 130.1 ($OBz_{ortho}$), 130.0 ($OBz_{ortho}$),129.9 (2 x $OBz_{ortho}$), 129.8 ($OBz_{ipso}$), 129.6 ($OBz_{ipso}$), 129.1 ($OBz_{ipso}$), 128.9 ($OBz_{ipso}$), 128.7 ($OBz_{meta}$), 128.6 ($OBz_{meta}$), 128.5 ($OBz_{meta}$), 128.4 ($OBz_{meta}$), 101.5 (C-1'), 98.6 (C-1), 78.4 (<u>C</u>≡CH), 76.6 (C-4), 75.4 (C≡<u>C</u>H), 72.4 (C-3'), 72.2 (C-3), 71.4 (C-2'), 70.9 (C-2), 62.8 (C-5), 62.7 (C-5'), 58.5 (C-4'), 55.9 (<u>C</u>H$_2$C≡CH) ppm.

**ESI-HRMS:** *m/z* = 779.25496 $[M+NH_4]^+$ (calculated *m/z* = 779.25590 for $[M+NH_4]^+$).

#### 2.33. Propargyl 4-azido-4-deoxy-xylopyranosyl-(1→4)-β-D-xylopyranoside (S22)

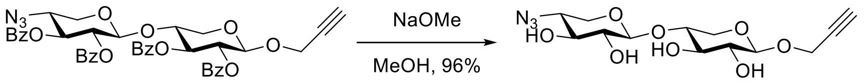

The protected disaccharide **S21** (116 mg, 152 µmol) was dissolved in dry $CH_2Cl_2$ (3 mL) and dry methanol (6 mL), sodium methoxide (5.4 M in methanol, 14 µL, 76 µmol) was added dropwise and the reaction mixture was stirred at room temperature for 15 h. It was neutralized by addition of Amberlite®-IRC 120 $H^+$ resin. The reaction mixture was filtered and the solvent was removed under reduced pressure. The crude product was purified on reversed phase silica gel by automated flash chromatography ($H_2O$:MeCN, 100:0 1 CV, 100:0 → 0:100 over 16 CV) to obtain the title compound after lyophilization (50.3 mg, 146 µmol, 96%).

$[\alpha]_D^{20}$ = -70.1 (*c* = 0.41, $H_2O$).

**IR (ATR)**: $\tilde{\nu}$ = 3381, 3287, 2871, 2108, 1082, 1041, 980, 897 $cm^{-1}$.

**$^1$H NMR** (500 MHz, methanol-d$_4$, 298 K): δ = 4.43 (d, $^3J_{1,2}$ = 7.4 Hz, 1H, H-1), 4.40-4.30 (m, 2H, C<u>H</u>$_2$C≡CH), 4.30 (d, $^3J_{1,2}$ = 7.7 Hz, 1H, H-1'), 4.01 (dd, $^2J_{5ax,5eq}$ = 11.7 Hz, $^3J_{4,5eq}$ = 5.2 Hz, 1H, H-5$_{eq}$), 3.94 (dd, $^2J_{5ax,5eq}$ = 11.6 Hz, $^3J_{4,5eq}$ = 5.0 Hz, 1H, H-5$_{eq}$'), 3.68-3.61 (m, 1H, H-4), 3.50-3.38 (m, 3H, H-3, H-3', H-4'), 3.32-3.29 (m, 1H, H-5$_{ax}$), 3.29-3.16 (m, 3H, H-2', H-2, H-5$_{ax}$'), 2.87 (t, $^4J_{CH2,C≡CH}$ = 2.4 Hz, 1H, C≡C<u>H</u>) ppm.

**$^{13}$C NMR** (151 MHz, methanol-d$_4$, 298 K): δ = 103.8 (C-1'), 102.6 (C-1), 79.8 (<u>C</u>≡CH), 78.0 (C-4), 76.9 (C-3'), 76.3 (C≡<u>C</u>H), 75.6 (C-3), 74.7 (C-2'), 74.4 (C-2), 65.1 (C-5'), 64.4 (C-5), 63.0 (C-4'), 56.5 (<u>C</u>H$_2$C≡CH) ppm.

**ESI-HRMS:** *m/z* = 368.10652 $[M+Na]^+$ (calculated *m/z* = 368.10644 for $[M+Na]^+$).

#### 2.34. Propargyl 2,3,4-tri-*O*-acetyl-β-D-xylopyranosyl-(1→4)-2,3-di-*O*-benzoyl-β-D-xylopyranosyl-(1→4)-2,3-di-*O*-benzoyl-β-D-xylopyranoside (S23)

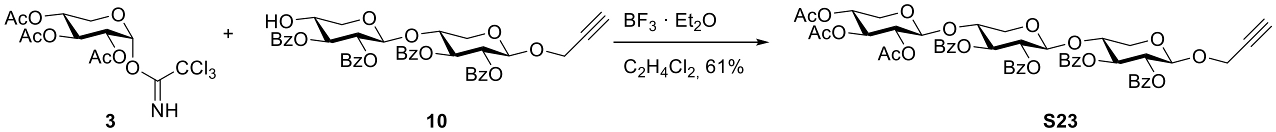

The xylosyl donor **3**[3] (62.3 mg, 148 µmol) and acceptor disaccharide **10** (83.0 mg, 113 µmol) were dissolved in dry $C_2H_4Cl_2$ (3.4 mL) under a nitrogen atmosphere, freshly activated powdered molecular sieves 4Å was added and the reaction mixture was stirred at room temperature for 15 min. At 0°C boron trifluoride diethyl etherate (10 µL, 79 µmol) was added. The reaction mixture was stirred at 0 °C for 2 h followed by 12 h at room temperature. The reaction mixture was quenched with a satd. aq. $NaHCO_3$ solution (1 mL) and diluted with $CH_2Cl_2$. After filtration over celite® the organic phase (50 mL) was washed with a satd. aq. $NaHCO_3$ solution (10 mL) and the aq. phase was extracted with $CH_2Cl_2$ (10 mL). All organic phases were combined and dried over $MgSO_4$, filtrated and the solvent was removed under reduced pressure. The crude product was purified on silica gel via automated flash chromatography (toluene:ethyl acetate, 9:1→8:2) to obtain the product as a colorless amorphous solid (69.0 mg, 69.4 µmol, 61%).

***R*$_F$** (toluene:ethyl acetate, 4:1) = 0.26.

$[\alpha]_D^{20}$ = -22.1 (*c* = 0.31, $CHCl_3$).

**IR (ATR):** $\tilde{\nu}$ = 3287, 2936, 1725, 1249, 707 $cm^{-1}$.

**$^1$H NMR** (600 MHz, $CDCl_3$, 298 K): δ = 8.02-7.90 (m, 8H, 4 x OBz$_{ortho}$), 7.57-7.49 (m, 4H, 4 x OBz$_{para}$), 7.44-7.35 (m, 8H, 4 x OBz$_{meta}$), 5.64 (t, $^3J_{2,3}$ = $^3J_{3,4}$ = 8.0 Hz 1H, H-3), 5.47 (t, $^3J_{2,3}$ = $^3J_{3,4}$ = 8.1 Hz, 1H, H-3'), 5.27 (dd, $^3J_{2,3}$ = 8.4 Hz, $^3J_{1,2}$ = 6.5 Hz, 1H, H-2), 5.17 (dd, $^3J_{2,3}$ = 8.3 Hz, $^3J_{1,2}$ = 6.3 Hz, 1H, H-2'), 4.95 (t, $^3J_{2,3}$ = $^3J_{3,4}$ = 7.7 Hz, 1H, H-3''), 4.89 (d, $^3J_{1,2}$ = 6.5 Hz, 1H, H-1), 4.76 (d, $^3J_{1,2}$ = 6.3 Hz, 1H, H-1'), 4.71 (dd, $^3J_{2,3}$ = 7.9 Hz, $^3J_{1,2}$ = 6.0 Hz, 1H, H-2''), 4.65-4.59 (m, 1H, H-4''), 4.38 (d, $^3J_{1,2}$ = 6.0 Hz, 1H, H-1''), 4.35-4.27 (m, 2H, C<u>H</u>$_2$C≡CH), 4.05-3.96 (m, 2H, H-4, H-5$_{eq}$), 3.78-3.71 (m, 1H, H-4'), 3.64 (dd, $^2J_{5ax,5eq}$ = 12.1 Hz, $^3J_{4,5eq}$ = 4.6 Hz, 1H, H-5$_{eq}$''), 3.58 (dd, $^2J_{5ax,5eq}$ = 12.2 Hz, $^3J_{4,5eq}$ = 4.8 Hz, 1H, H-5$_{eq}$'), 3.46 (dd, $^2J_{5ax,5eq}$ = 11.9 Hz, $^3J_{4,5eq}$ = 8.2 Hz, 1H, H-5$_{ax}$), 3.18 (dd, $^2J_{5ax,5eq}$ = 12.2 Hz, $^3J_{4,5ax}$ = 8.4 Hz, 1H, H-5$_{ax}$'), 3.05 (dd, $^2J_{5ax,5eq}$ = 12.1 Hz, $^3J_{4,5ax}$

= 7.7 Hz, 1H, H-5$_{ax}$''), 2.37 (t, $^{4}J_{CH2,C≡CH}$ = 2.4 Hz, 1H, C≡C<u>H</u>), 1.98 (s, 3H, $CH3_{OAc\ at\ C\text{-}4''}$), 1.97 (s, 3H, $CH3_{OAc\ at\ C\text{-}3''}$), 1.93 (s, 3H, $CH3_{OAc\ at\ C\text{-}2''}$) ppm.

**$^{13}$C NMR** (151 MHz, $CDCl_3$, 298 K): δ = 170.1 ($C{=}O_{OAc\ at\ C\text{-}3''}$), 169.8 ($C{=}O_{OAc\ at\ C\text{-}4''}$), 169.1 ($C{=}O_{OAc\ at\ C\text{-}2''}$), 165.5 ($C{=}O_{OBz\ at\ C\text{-}3}$), 165.4 ($C{=}O_{OBz\ at\ C\text{-}3'}$), 165.4 ($C{=}O_{OBz\ at\ C\text{-}2}$), 165.2 ($C{=}O_{OBz\ at\ C\text{-}2'}$), 133.5 ($OBz_{para}$), 133.34 ($OBz_{para}$), 133.25 (2 x $OBz_{para}$), 130.1 ($OBz_{ortho}$), 129.90 (2 x $OBz_{ortho}$), 129.86 ($OBz_{ipso}$), 129.81 ($OBz_{ortho}$), 129.6 ($OBz_{ipso}$), 129.5 ($OBz_{ipso}$), 129.3 ($OBz_{ipso}$), 128.64 ($OBz_{meta}$), 128.55 ($OBz_{meta}$), 128.48 ($OBz_{meta}$), 128.42 ($OBz_{meta}$), 101.3 (C-1'), 99.7 (C-1''), 98.6 (C-1), 78.5 (<u>C</u>≡CH), 76.2 (C-4), 75.4 (C≡<u>C</u>H), 74.5 (C-4'), 72.2 (C-3), 71.9 (C-3'), 71.3 (C-2'), 70.9 (C-2), 70.5 (C-3''), 70.2 (C-2''), 68.2 (C-4''), 62.8 (C-5), 62.3 (C-5'), 61.3 (C-5''), 55.8 (<u>C</u>H$_2$C≡CH), 20.8 ($CH3_{OAc\ at\ C\text{-}4''}$), 20.7 ($CH3_{OAc\ at\ C\text{-}2''}$, $CH3_{OAc\ at\ C\text{-}3''}$) ppm.

**ESI-HRMS:** *m/z* = 1012.33511 $[M+NH_4]^+$ (calculated *m/z* = 1012.32337 for $[M+NH_4]^+$).

##### 2.35. Propargyl 2,3,4-tri-*O*-acetyl-β-D-xylopyranosyl-(1→4)-2,3-di-*O*-benzoyl-β-D-xylopyranosyl-(1→4)-2,3-di-*O*-benzoyl-β-D-xylopyranosyl-(1→4)-2,3-di-*O*-benzoyl-β-D-xylopyranoside (S24)

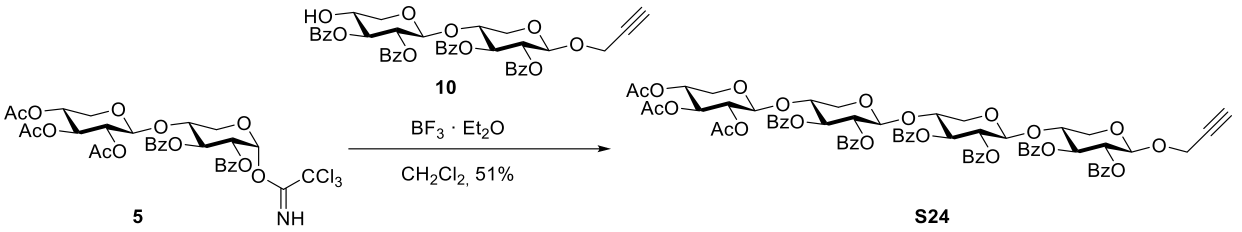

The xylosyl donor **5** (101 mg, 133 µmol) and acceptor disaccharide **10** (70.0 mg, 95.0 µmol) were dissolved in dry $CH_2Cl_2$ (3.0 mL) under a nitrogen atmosphere, freshly activated powdered molecular sieves 4Å was added and the reaction mixture was stirred at room temperature for 15 min. At 0°C boron trifluoride diethyl etherate (15 µL, 119 µmol) was added. The reaction mixture was stirred at 0 °C for 2 h followed by 11 h at room temperature. The reaction mixture was quenched with a satd. aq. $NaHCO_3$ solution (1 mL) and diluted with $CH_2Cl_2$. After filtration over celite® the organic phase (100 mL) was washed with a satd. aq. $NaHCO_3$ solution (15 mL) and the aq. phase was extracted with $CH_2Cl_2$ (15 mL). All organic phases were combined and dried over $MgSO_4$, filtrated and the solvent was removed under reduced pressure. The crude product was purified on silica gel via automated flash chromatography (toluene:ethyl acetate, 9:1→8:2) followed by purification on silica gel via automated flash chromatography (cyclohexane:ethyl acetate, 8:2→6:4) to obtain the product as a colorless amorphous solid (64.8 mg, 48.5 µmol, 51%).

***R*$_F$** (toluene:ethyl acetate, 4:1) = 0.25.

$[\alpha]_D^{20}$ = -22.4 (*c* = 0.30, $CHCl_3$).

**IR (ATR):** $\tilde{\nu}$ = 2945, 2867, 1724, 1249, 1068, 706 cm$^{-1}$.

**$^1$H NMR** (500 MHz, $CDCl_3$, 298 K): δ = 7.99-7.87 (m, 12H, 6 x $OBz_{ortho}$), 7.59-7.48 (m, 5H, 5 x $OBz_{para}$), 7.45-7.33 (m, 11H, 5 x $OBz_{meta}$, $OBz_{para}$), 7.25-7.20 (m, 2H, $OBz_{meta}$), 5.59 (t, $^3J_{2,3}$ = $^3J_{3,4}$ = 8.0 Hz, 1H, H-3), 5.49 (t, $^3J_{2,3}$ = $^3J_{3,4}$ =8.1 Hz, 1H, H-3'), 5.40 (t, $^3J_{2,3}$ = $^3J_{3,4}$ = 8.1 Hz, 1H, H-3''), 5.23 (dd, $^3J_{2,3}$ = 8.5 Hz, $^3J_{1,2}$ = 6.6 Hz, 1H, H-2), 5.15 (dd, $^3J_{2,3}$ = 8.3 Hz, $^3J_{1,2}$ = 6.4 Hz, 1H, H-2'), 5.07 (dd, $^3J_{2,3}$ = 8.3 Hz, $^3J_{1,2}$ = 6.3 Hz, 1H, H-2''), 4.94 (t, $^3J_{2,3}$ = $^3J_{3,4}$ = 7.7 Hz, 1H, H-3'''), 4.86 (d,

$^3J_{1,2}$ = 6.6 Hz, 1H, H-1), 4.71-4.67 (m, 2H, H-1', H-2'''), 4.64-4.58 (m, 1H, H-4'''), 4.52 (d, $^3J_{1,2}$ = 6.3 Hz, 1H, H-1''), 4.36 (d, $^3J_{1,2}$ = 6.0 Hz, 1H, H-1'''), 4.33-4.25 (m, 2H, C<u>H</u>$_2$C≡CH), 4.00-3.92 (m, 2H, H-4, H-5$_{eq}$), 3.79-3.68 (m, 2H, H-4', H-4''), 3.62 (dd, $^2J_{5ax,5eq}$ = 12.1 Hz, $^3J_{4,5eq}$ = 4.6 Hz, 1H, H-5$_{eq}$'''), 3.51 (dd, $^2J_{5ax,5eq}$ = 12.2 Hz, $^3J_{4,5eq}$ = 4.8 Hz, 1H, H-5$_{eq}$''), 3.47-3.38 (m, 2H, H-5$_{eq}$', H-5$_{ax}$), 3.10 (dd, $^2J_{5ax,5eq}$ = 12.2 Hz, $^3J_{4,5ax}$ = 8.5 Hz, 2H, H-5$_{ax}$', H-5$_{ax}$''), 3.03 (dd, $^2J_{5ax,5eq}$ = 12.1 Hz, $^3J_{4,5ax}$ = 7.7 Hz, 1H, H-5$_{ax}$'''), 2.35 (t, $^4J_{CH2,C≡CH}$ = 2.4 Hz, 1H, C≡C<u>H</u>), 1.97 (s, 3H, CH3$_{OAc\ at\ C-4'''}$), 1.96 (s, 3H, CH3$_{OAc\ at\ C-3'''}$), 1.92 (s, 3H, CH3$_{OAc\ at\ C-2'''}$) ppm.

**$^{13}$C NMR** (126 MHz, CDCl$_3$, 298 K): δ = 170.0 (C=O$_{OAc\ at\ C-3'''}$), 169.6 (C=O$_{OAc\ at\ C-4'''}$), 169.0 (C=O$_{OAc\ at\ C-2'''}$), 165.4 (C=O$_{OBz\ at\ C-3}$), 165.3 (3 x C=O$_{OBz\ at\ C-3',\ OBz\ at\ C-2,\ OBz\ at\ C-3''}$), 165.0 (C=O$_{OBz\ at\ C-2'}$), 164.9 (C=O$_{OBz\ at\ C-2''}$), 133.4 (OBz$_{para}$), 133.3 (OBz$_{para}$), 133.2 (OBz$_{para}$), 133.10 (2 x OBz$_{para}$), 133.05 (OBz$_{para}$), 130.0 (OBz$_{ortho}$), 129.8 (OBz$_{ortho}$), 129.7 (2 x OBz$_{ortho}$), 129.64 (2 x OBz$_{ortho}$), 129.60 (2 x OBz$_{ipso}$), 129.5 (OBz$_{ipso}$), 129.4 (OBz$_{ipso}$), 129.3 (OBz$_{ipso}$), 129.2 (OBz$_{ipso}$), 128.5 (2 x OBz$_{meta}$), 128.40 (OBz$_{meta}$), 128.35 (OBz$_{meta}$), 128.3 (OBz$_{meta}$), 128.2 (OBz$_{meta}$), 101.0 (C-1'), 100.8 (C-1''), 99.5 (C-1'''), 98.5 (C-1), 78.3 (<u>C</u>≡CH), 75.9 (C-4), 75.5 (C-4'), 75.2 (C≡<u>C</u>H), 74.4 (C-4''), 72.1 (C-3, C-3'), 71.7 (C-3''), 71.4 (C-2'), 71.0 (C-2''), 70.8 (C-2), 70.3 (C-3'''), 70.0 (C-2'''), 68.1 (C-4'''), 62.7 (C-5), 62.3 (C-5'), 62.1 (C-5''), 61.1 (C-5'''), 55.7 (<u>C</u>H$_2$C≡CH), 20.7 (CH3$_{OAc\ at\ C-4'''}$), 20.6 (CH3$_{OAc\ at\ C-2'''}$, CH3$_{OAc\ at\ C-3'''}$) ppm.

**ESI-HRMS:** *m/z* = 1352.41706 [M+NH$_4$]$^+$ (calculated *m/z* = 1352.41806 for [M+NH$_4$]$^+$).

#### 2.36. Propargyl 2,3,4,6-tetra-*O*-acetyl-α-D-mannopyranosyl-(1→4)-[2,3,4,6-tetra-*O*-acetyl-β-D-glucopyranosyl-(1→6)]-2,3-di-*O*-acetyl-β-D-glucopyranoside (S25)

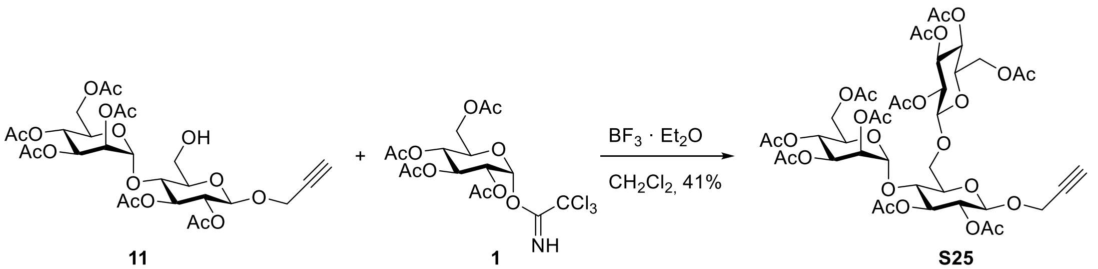

**11** **1** **S25**

The glucosyl donor **1**[1] (535 mg, 1.09 mmol, 1.2 equiv) and acceptor disaccharide **11** (572 mg, 904 µmol, 1 equiv) were dissolved in dry $CH_2Cl_2$ (22 mL) under nitrogen atmosphere, freshly activated molecular sieves (4Å, ~200 mg) was added and the reaction mixture stirred at room temperature for 30 min. At 0 °C boron trifluoride diethyl etherate (126 µL, 995 µmol, 1.1 equiv) was added and the reaction mixture was stirred at 0 °C for 2 h, followed by stirring at room temperature for 17 h. The reaction mixture was diluted with $CH_2Cl_2$ (20 mL) and satd. aq. $NaHCO_3$ solution (20 mL) and strongly stirred for 10 min. After filtration over celite® the phases were separated and the aq. phase was extracted with $CH_2Cl_2$ (2 x 20 mL). All organic phases were combined and dried over $MgSO_4$, filtrated and the solvent was removed under reduced pressure. The crude was purified on silica gel via automated flash chromatography (toluene:ethyl acetate, 60:40 → 50:50) to yield a mixture of acceptor **11** and product (505 mg). The crude was dissolved in pyridine (5 mL) and acetic anhydride (2.5 mL) was added at room temperature. After stirring for 3 h the reaction mixture was diluted with ethyl acetate (50 mL), washed with 1 M HCl solution (50 mL), satd. aq. $NaHCO_3$ solution (50 mL) and brine (25 mL). The organic phase was dried over $MgSO_4$, filtrated and the solvent was removed under reduced pressure. The crude was purified on silica gel via automated flash chromatography (cyclohexane:ethyl acetate, 50:50) to yield the product as a colorless foam (360 mg, 374 µmol, 41%).

***R*<sub></sub>**$\boldsymbol{R_F}$ (cyclohexane:ethyl acetate, 1:1) = 0.23.

$[\alpha]_D^{20}$ = -26.0 (*c* = 0.45 in $CHCl_3$).

**IR (ATR):** $\tilde{\nu}$ = 3278 (w), 2942 (w), 1742 (s), 1432 (w), 1367 (m), 1212 (s), 1138 (m), 1035 (s), 902 (m), 600 (m) $cm^{-1}$.

**$^1$H NMR** (500 MHz, $CDCl_3$, 298 K) δ = 5.28 – 5.17 (m, 4H, H-$3_{Glc}$, H-$3_{Glc'}$, H-$3_{Man}$, H-$4_{Man}$), 5.10 (dd, $^3J$ = 10.0 Hz, $^3J$ = 9.3 Hz, 1H, H-$4_{Glc'}$), 5.04 – 4.99 (m, 2H, H-$2_{Glc'}$, H-$2_{Man}$), 4.91 (d, $^3J_{1,2}$ = 2.1 Hz, 1H, H-$1_{Man}$), 4.86 (dd, $^3J_{2,3}$ = 9.5 Hz, $^3J_{1,2}$ = 7.9 Hz, 1H, H-$2_{Glc}$), 4.74 (d, $^3J_{1,2}$ = 8.0 Hz, 1H, H-$1_{Glc}$), 4.74 (d, $^3J_{1,2}$ = 7.9 Hz, 1H, H-$1_{Glc'}$), 4.42 – 4.34 (m, 2H, $C\underline{H}_2C{\equiv}CH$), 4.29 – 4.20 (m, 2H, H-$6a_{Glc'}$, H-$6a_{Man}$), 4.18 – 4.11 (m, 3H, H-$6a_{Glc}$, H-$6b_{Glc'}$, H-$6b_{Man}$), 4.03 (ddd, $^3J_{4,5}$ = 8.3 Hz, $^3J_{5,6a}$ = 5.5 Hz, $^3J_{5,6b}$ = 2.5 Hz, 1H, H-$5_{Man}$), 3.81 (dd, $^2J$ = 11.8 Hz, $^3J_{5,6b}$ = 6.1 Hz, 1H, H-$6b_{Glc}$), 3.76 – 3.68 (m, 2H, H-$4_{Glc}$, H-$5_{Glc'}$), 3.64 (ddd, $^3J_{4,5}$ = 9.8 Hz, $^3J_{5,6b}$ = 6.1 Hz, $^3J_{5,6a}$ = 1.8 Hz, 1H, H-$5_{Glc}$), 2.49 (t, $^4J_{C{\equiv}CH,CH2}$ = 2.4 Hz, 1H, $C{\equiv}C\underline{H}$), 2.13 (s, 3H, OAc), 2.11 (s, 3H, OAc), 2.09 (s, 3H, OAc), 2.09 (s, 3H, OAc), 2.07 (s, 3H, OAc), 2.06 (s, 3H, OAc), 2.05 (s, 3H, OAc), 2.02 (s, 3H, OAc), 2.00 (s, 3H, OAc), 1.99 (s, 3H, OAc) ppm.

**$^{13}$C NMR** (126 MHz, $CDCl_3$, 298 K) δ = 170.6 ($CH_3\underline{C}O$), 170.5 ($CH_3\underline{C}O$), 170.3 ($CH_3\underline{C}O$), 170.2 ($CH_3\underline{C}O$), 169.8 ($CH_3\underline{C}O$), 169.7 (2 x $CH_3\underline{C}O$), 169.6 ($CH_3\underline{C}O$), 169.4 ($CH_3\underline{C}O$), 169.2 ($CH_3\underline{C}O$), 100.7 (C-$1_{Glc'}$), 99.8 (C-$1_{Man}$), 97.9 (C-$1_{Glc}$), 78.3 ($CH_2\underline{C}{\equiv}CH$), 78.0 (C-$4_{Glc}$/C-$5_{Glc'}$), 75.6 ($CH_2C{\equiv}\underline{C}H$), 74.6 (C-$5_{Glc}$), 74.1 (C-$3_{Glc}$), 72.8 (C-$3_{Glc'}$), 71.9 (C-$4_{Glc}$/C-$5_{Glc'}$), 71.4 (C-$2_{Glc}$), 71.2 (C-$2_{Glc'}$), 69.7 (C-$5_{Man}$), 69.6 (C-$2_{Man}$), 68.4 (C-$3_{Man}$/C-$4_{Man}$), 68.2 (C-$4_{Glc'}$), 67.9 (C-$6_{Glc}$), 65.9 (C-$3_{Man}$/C-$4_{Man}$), 62.3 (C-$6_{Man}$), 61.7 (C-$6_{Glc'}$), 55.9 ($\underline{C}H_2C{\equiv}CH$), 20.75 ($CH_3$), 20.71 (2 x $CH_3$), 20.68 ($CH_3$), 20.66 (3 x $CH_3$), 20.60 ($CH_3$), 20.58 ($CH_3$), 20.58 ($CH_3$) ppm.

**ESI-HRMS:** *m/z* = 980.32172 $[M+NH_4]^+$ (calculated *m/z* = 980.31962).

##### 2.37. Propargyl 2,3,4,6-tetra-*O*-benzoyl-β-D-glucopyranosyl-(1→4)-[2,3,4,6-tetra-*O*-acetyl-α-D-mannopyranosyl-(1→6)]-2,3-di-*O*-benzoyl-β-D-glucopyranoside (S26)

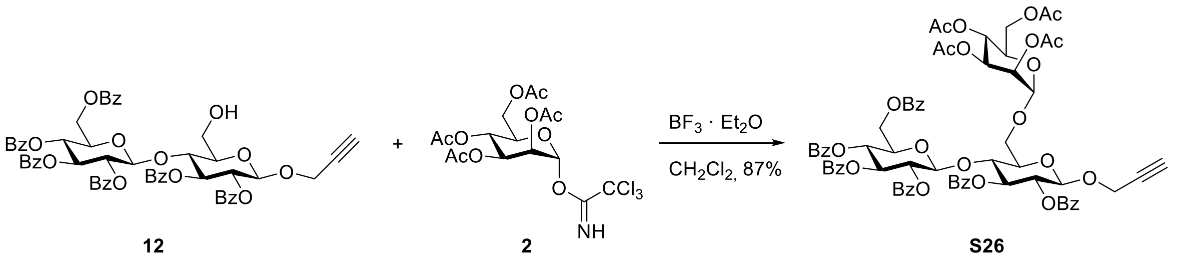

12 2 S26

The mannosyl donor **2**[2] (289 mg, 587 µmol, 1.2 equiv) and acceptor disaccharide **12** (491 mg, 489 µmol, 1 equiv) were dissolved in dry $CH_2Cl_2$ (14 mL) under nitrogen atmosphere, freshly activated molecular sieve (3Å, 850 mg) was added and the reaction mixture was stirred at room temperature for 30 min. At 0 °C boron trifluoride diethyl etherate (68 µL, 537 µmol, 1.1 equiv) was added and the reaction mixture was stirred at 0 °C for 2 h, followed by stirring at room temperature for 17 h. The reaction mixture was diluted with satd. aq. $NaHCO_3$ solution (20 mL) and strongly stirred for 30 min. After filtration over celite® ethyl acetate (200 mL) was added and the mixture was washed with water (2 x 20 mL) and brine (40 mL). The organic phase was dried over $MgSO_4$, filtrated and the solvent was removed under reduced pressure. The crude was purified on silica gel via automated flash chromatography (toluene:ethyl acetate, 100:0 → 80:20) to yield the product as a colorless foam (568 mg, 425 µmol, 87%).

***R*$_F$** (cyclohexane:ethyl acetate, 1:1) = 0.66.

$[\alpha]_D^{20}$ = +31.8 (*c* = 1.19 in $CHCl_3$).

**IR (ATR):** $\tilde{\nu}$ = 2957 (w), 1725 (s), 1602 (w), 1451 (m), 1368 (m), 1247 (s). 1220 (s), 1089 (s), 1067 (s), 1026 (s), 976 (m) $cm^{-1}$.

**$^1$H NMR** (600 MHz, $CDCl_3$, 298 K) δ = 7.98 – 7.87 (m, 8H, OBz), 7.80 – 7.72 (m, 4H, OBz), 7.57 – 7.10 (m, 18H, OBz), 5.86 (t, $^3J_{2,3}$ = $^3J_{3,4}$ = 9.7 Hz, 1H, H-3$_{Glc'}$), 5.77 (t, $^3J_{2,3}$ = $^3J_{3,4}$ = 9.5 Hz, 1H, H-3$_{Glc}$), 5.51 (dd, $^3J_{2,3}$ = 9.9 Hz, $^3J_{1,2}$ = 8.0 Hz, 1H, H-2$_{Glc'}$), 5.47 (t, $^3J_{3,4}$ = $^3J_{4,5}$ = 9.8 Hz, 1H, H-4$_{Glc'}$), 5.41 – 5.33 (m, 3H, H-2$_{Glc}$, H-3$_{Man}$, H-4$_{Man}$), 5.27 (dd, $^3J_{2,3}$ = 3.1 Hz, $^3J_{1,2}$ = 1.9 Hz, 1H, H-2$_{Man}$), 5.02 (d, $^3J_{1,2}$ = 8.0 Hz, 1H, H-1$_{Glc'}$), 4.90 – 4.86 (m, 2H, H-1$_{Glc}$, H-1$_{Man}$), 4.33 (dd, $^2J$ = 12.3 Hz, $^3J_{5,6a}$ = 4.5 Hz, 1H, H-6a$_{Man}$), 4.31 – 4.29 (m, 2H, C<u>H</u>$_2$C≡CH), 4.20 – 4.08 (m, 3H, H-4$_{Glc}$, H-5$_{Glc'}$, H-6b$_{Man}$), 4.01 (ddd, $^3J_{4,5}$ = 9.3 Hz, $^3J_{5,6a}$ = 4.4 Hz, $^3J_{5,6b}$ = 2.5 Hz, 1H, H-5$_{Man}$), 3.99 – 3.90 (m, 2H, H-6a$_{Glc'}$, H-6b$_{Glc'}$), 3.82 – 3.77 (m, 2H, H-6a$_{Glc}$, H-6b$_{Glc}$), 3.67 (dt, $^3J_{4,5}$ = 9.9 Hz, $^3J_{5,6a}$ = $^3J_{5,6b}$ =3.1 Hz, 1H, H-5$_{Glc}$), 2.37 (t, $^4J_{C≡CH,CH2}$ = 2.4 Hz, 1H, C≡C<u>H</u>), 2.19 (s, 3H, OAc), 2.12 (s, 3H, OAc), 2.11 (s, 3H, OAc), 1.96 (s, 3H, OAc) ppm.

**$^{13}$C NMR** (151 MHz, $CDCl_3$, 298 K) δ = 170.7 ($CH_3$<u>C</u>O), 169.9 ($CH_3$<u>C</u>O), 169.8 ($CH_3$<u>C</u>O), 169.7 ($CH_3$<u>C</u>O), 165.7 (Ph<u>C</u>O), 165.7 (Ph<u>C</u>O), 165.4 (Ph<u>C</u>O), 165.2 (Ph<u>C</u>O), 164.9 (Ph<u>C</u>O), 164.7 (Ph<u>C</u>O), 133.5 (OBz$_{para}$), 133.3 (OBz$_{para}$), 133.2 (OBz$_{para}$), 133.1 (OBz$_{para}$), 133.1 (OBz$_{para}$), 133.0 (OBz$_{para}$), 129.9 (OBz), 129.8 (OBz), 129.7 (OBz), 129.7 (OBz), 129.6 (OBz), 129.3 (OBz), 129.3 (OBz), 129.0 (OBz), 128.8 (OBz), 128.7 (OBz), 128.6 (OBz), 128.6 (OBz), 128.4 (OBz), 128.3 (OBz), 128.3 (OBz), 128.2 (OBz), 128.2 (OBz), 128.2 (OBz), 101.2 (C-1$_{Glc'}$), 98.5 (C-1$_{Glc}$), 97.5 (C-1$_{Man}$), 78.0 ($CH_2$<u>C</u>≡CH), 76.7 (C-4$_{Glc}$), 75.6 ($CH_2$C≡<u>C</u>H), 74.7 (C-5$_{Glc}$), 73.0 (C-3$_{Glc'}$), 72.6 (C-3$_{Glc}$), 72.2 (C-2$_{Glc'}$, C-5$_{Glc'}$), 71.6 (C-3$_{Man}$), 69.12 (C-4$_{Glc'}$), 69.07 (C-2$_{Glc}$), 69.0 (C-2$_{Man}$, C-5$_{Man}$), 65.9 (C-4$_{Man}$), 64.5 (C-6$_{Glc}$), 62.5 (C-6$_{Glc'}$), 62.3 (C-6$_{Man}$), 56.0 (<u>C</u>H$_2$C≡CH), 20.9 ($CH_3$), 20.8 ($CH_3$), 20.8 ($CH_3$), 20.6 ($CH_3$) ppm.

**ESI-HRMS:** *m/z* = 1352.41760 $[M+NH_4]^+$ (calculated *m/z* = 1352.41806).

#### 3. NMR Spectra of Synthesized Saccharides

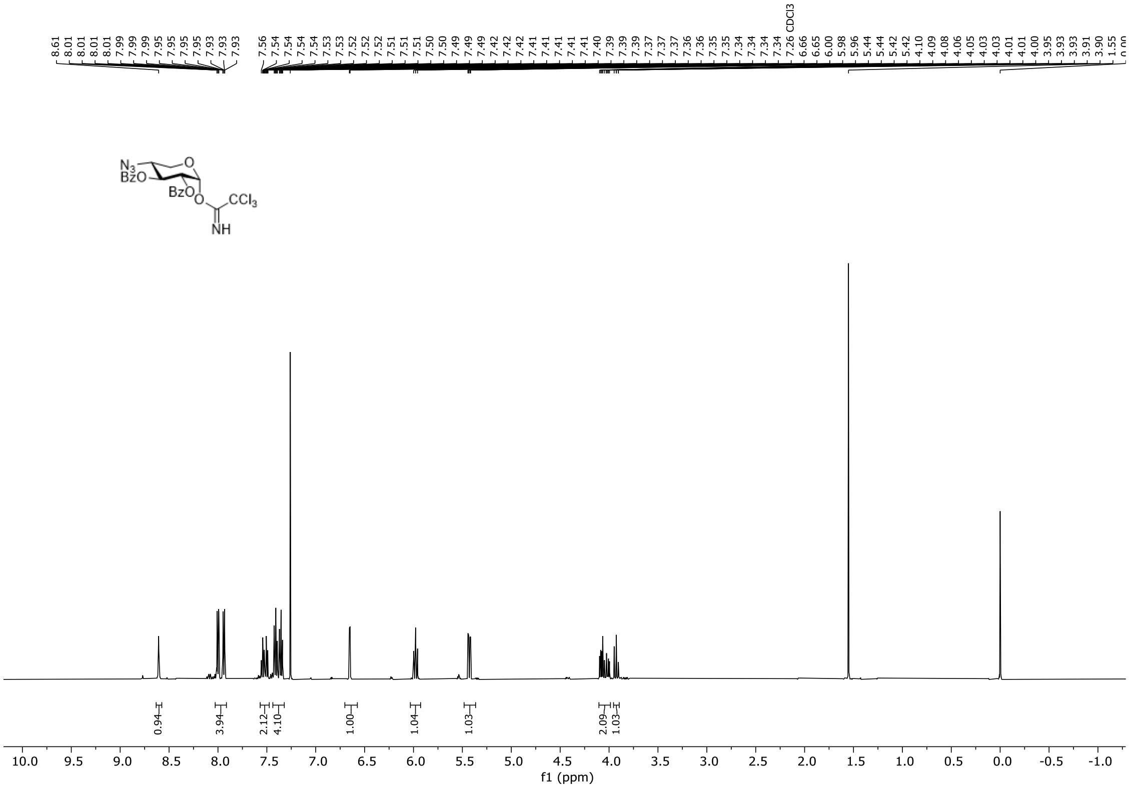

**Figure S2.** $^{1}H$ NMR spectrum of compound **4** (500 MHz, $CDCl_3$, 298 K).

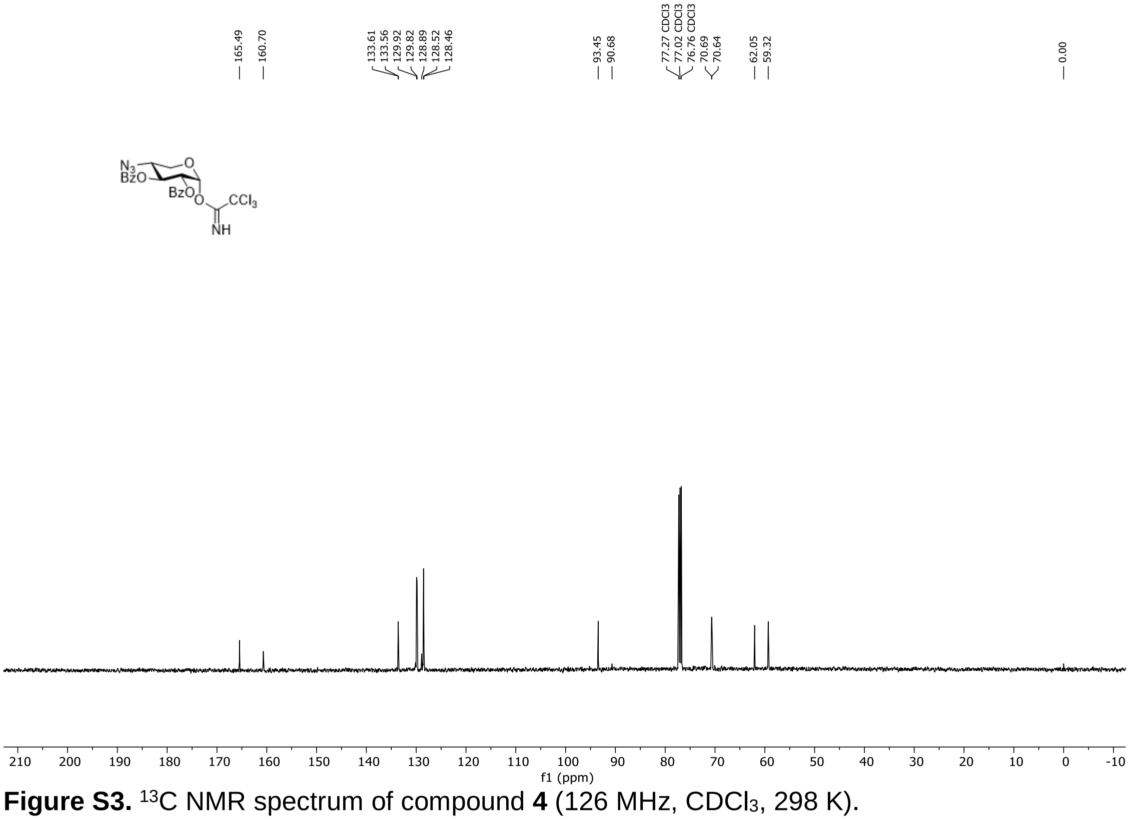

**Figure S3.** $^{13}C$ NMR spectrum of compound **4** (126 MHz, $CDCl_3$, 298 K).

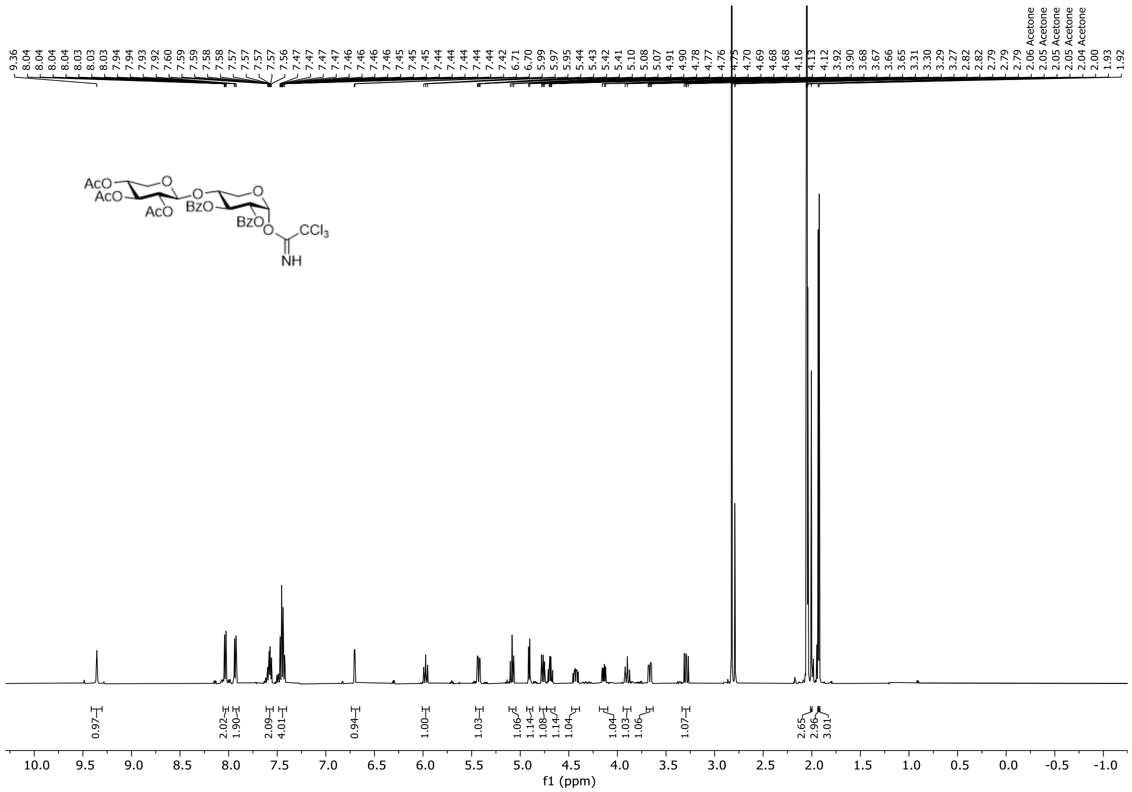

**Figure S4.** $^{1}H$ NMR spectrum of compound **5** (500 MHz, acetone-$d_6$, 298 K).

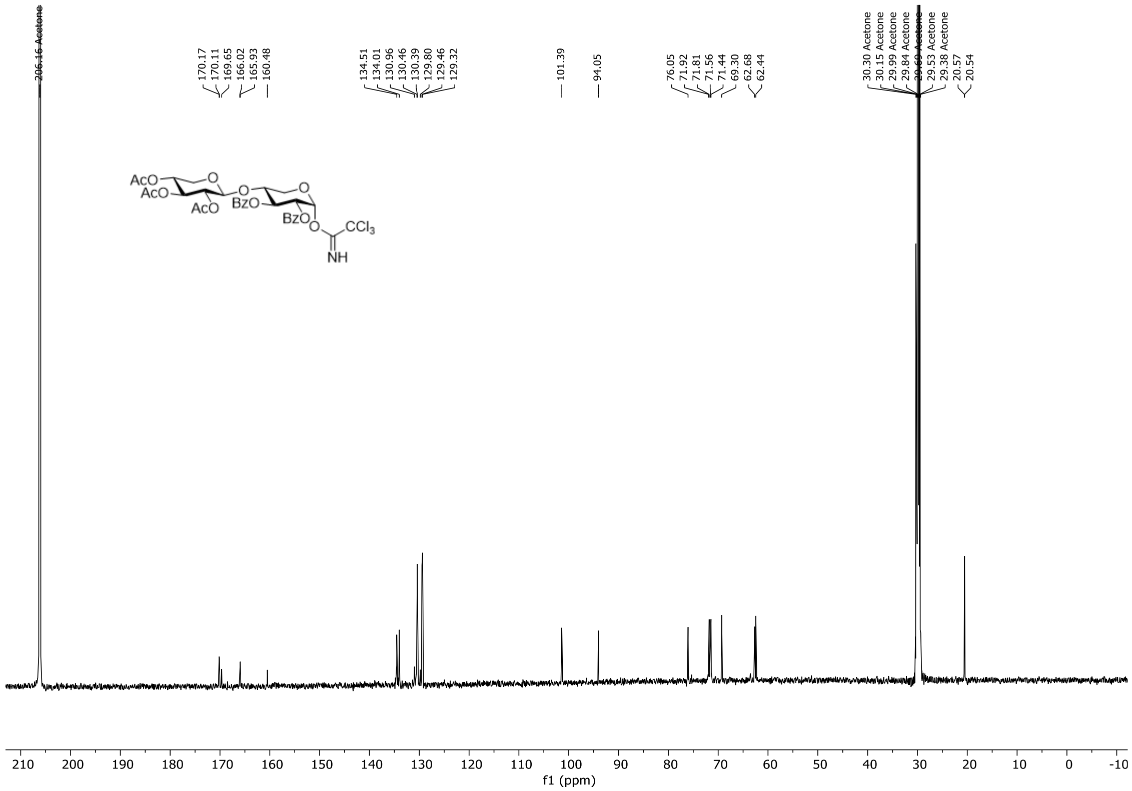

**Figure S5.** $^{13}C$ NMR spectrum of compound **5** (126 MHz, acetone-$d_6$, 298 K).

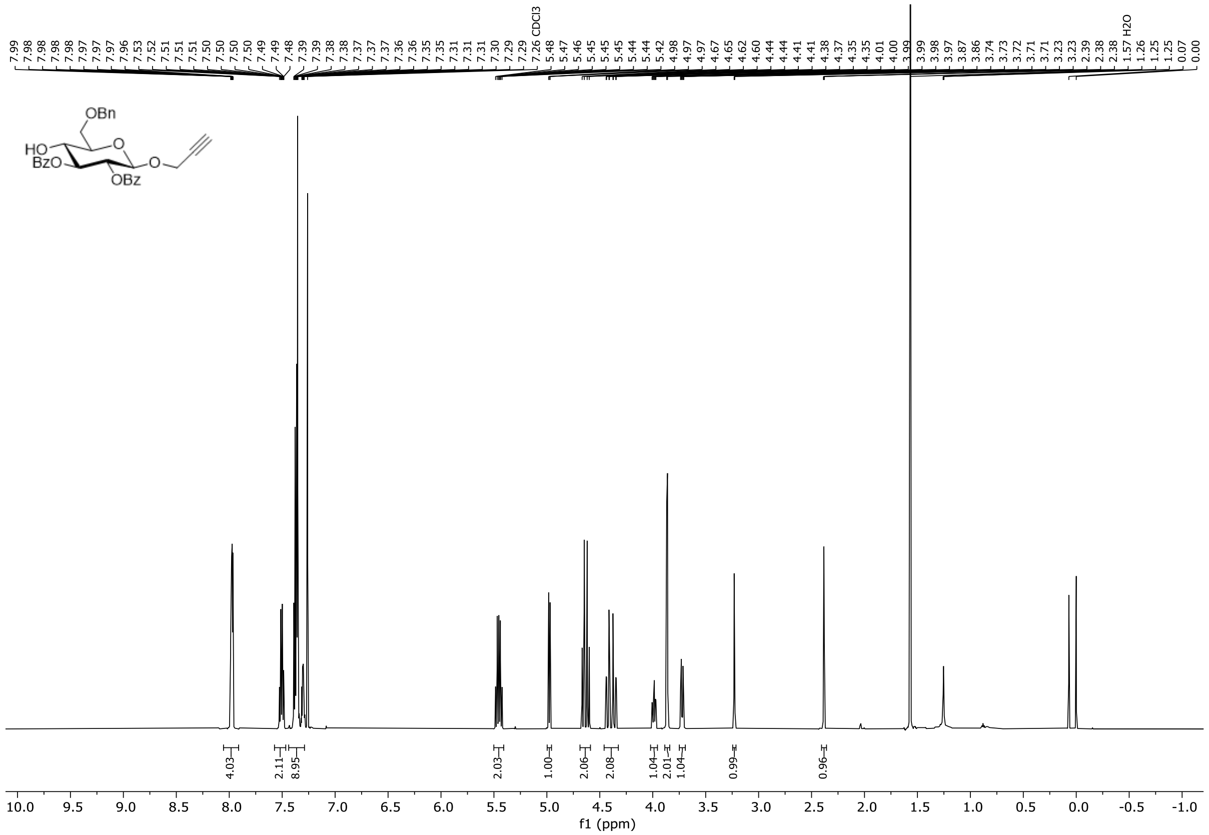

**Figure S6.** $^{1}H$ NMR spectrum of compound **8** (600 MHz, $CDCl_3$, 298 K).

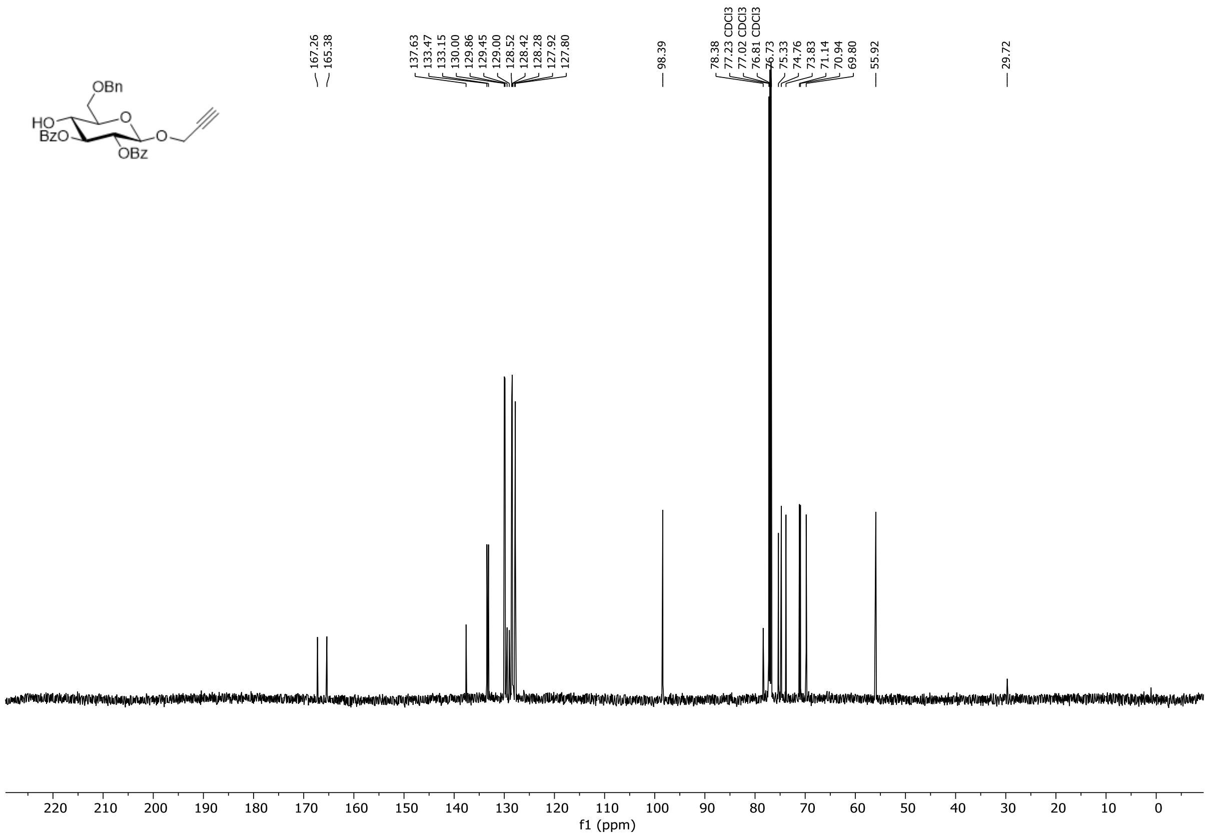

**Figure S7.** $^{13}C$ NMR spectrum of compound **8** (151 MHz, $CDCl_3$, 298 K).

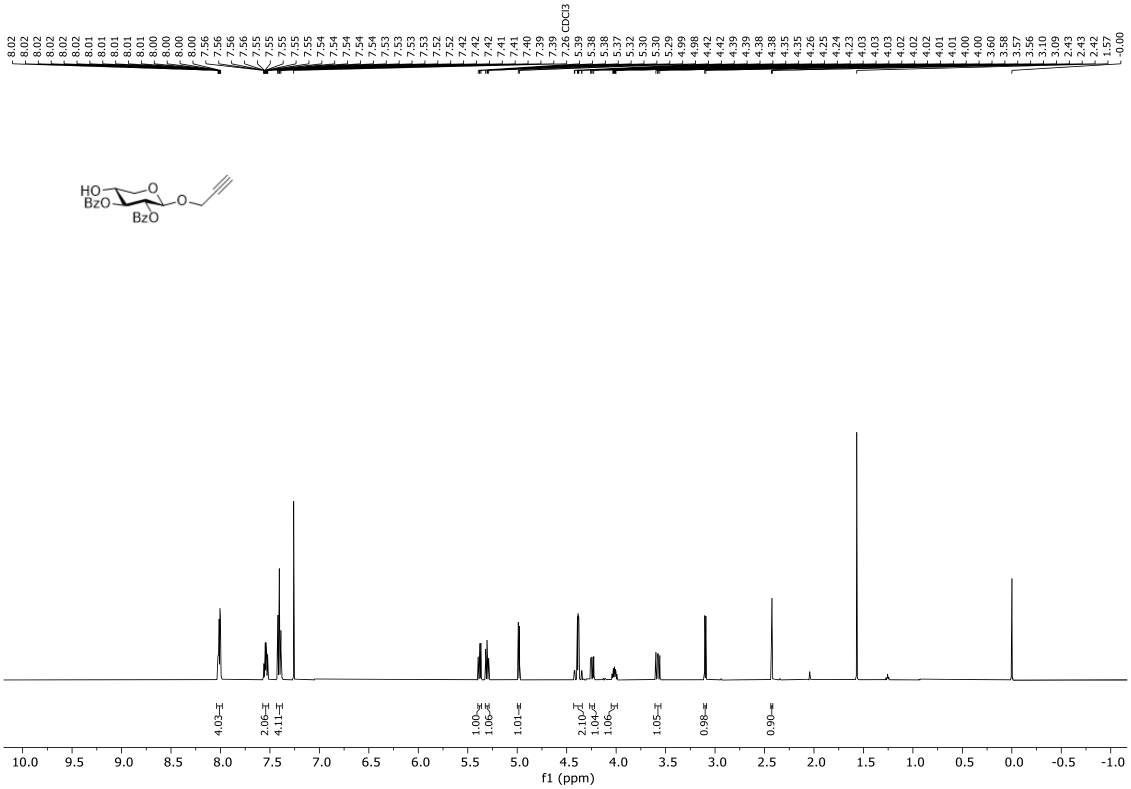

**Figure S8.** $^1$H NMR spectrum of compound **9** (500 MHz, $CDCl_3$, 298 K).

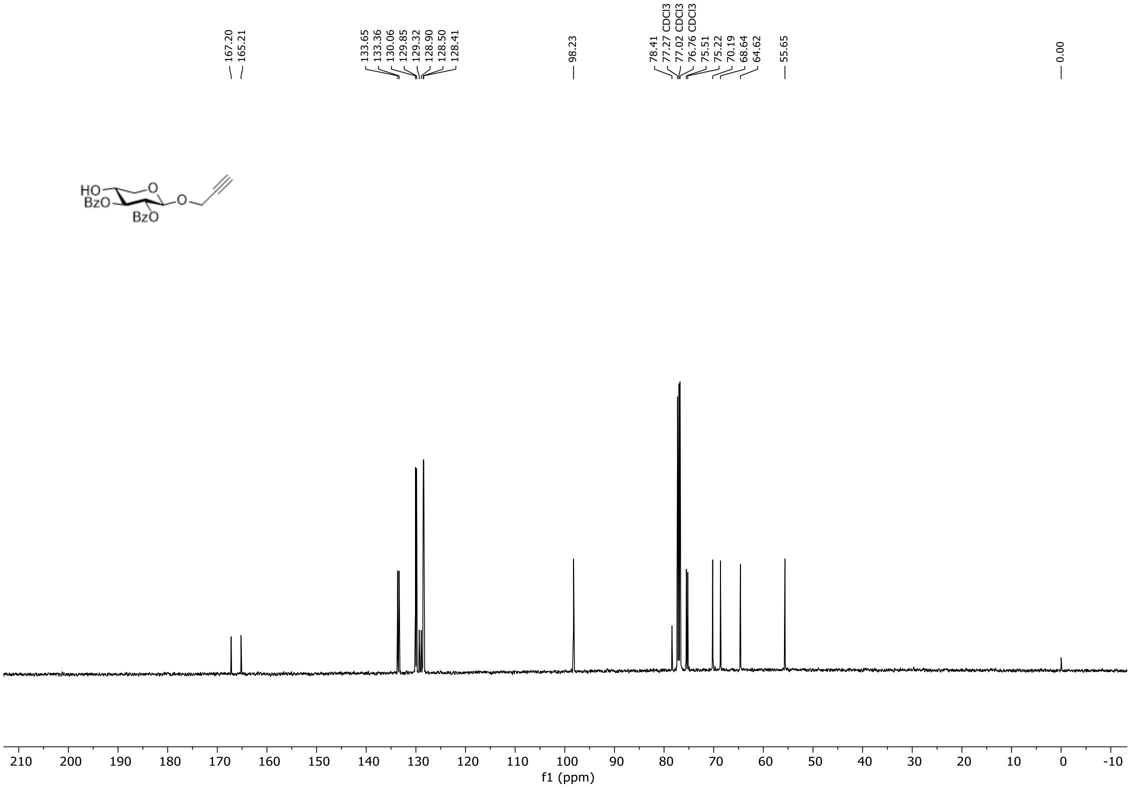

**Figure S9.** $^{13}$C NMR spectrum of compound **9** (126 MHz, $CDCl_3$, 298 K).

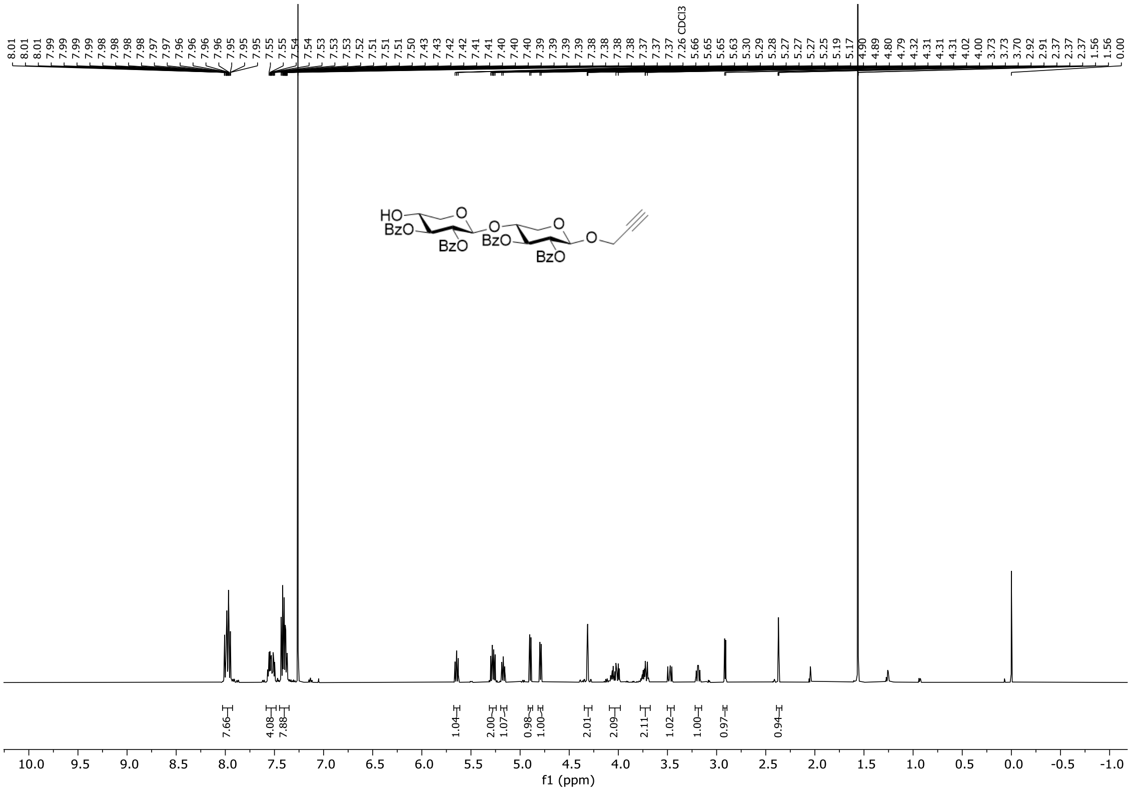

**Figure S10.** $^{1}$H NMR spectrum of compound **10** (500 MHz, $CDCl_3$, 298 K).

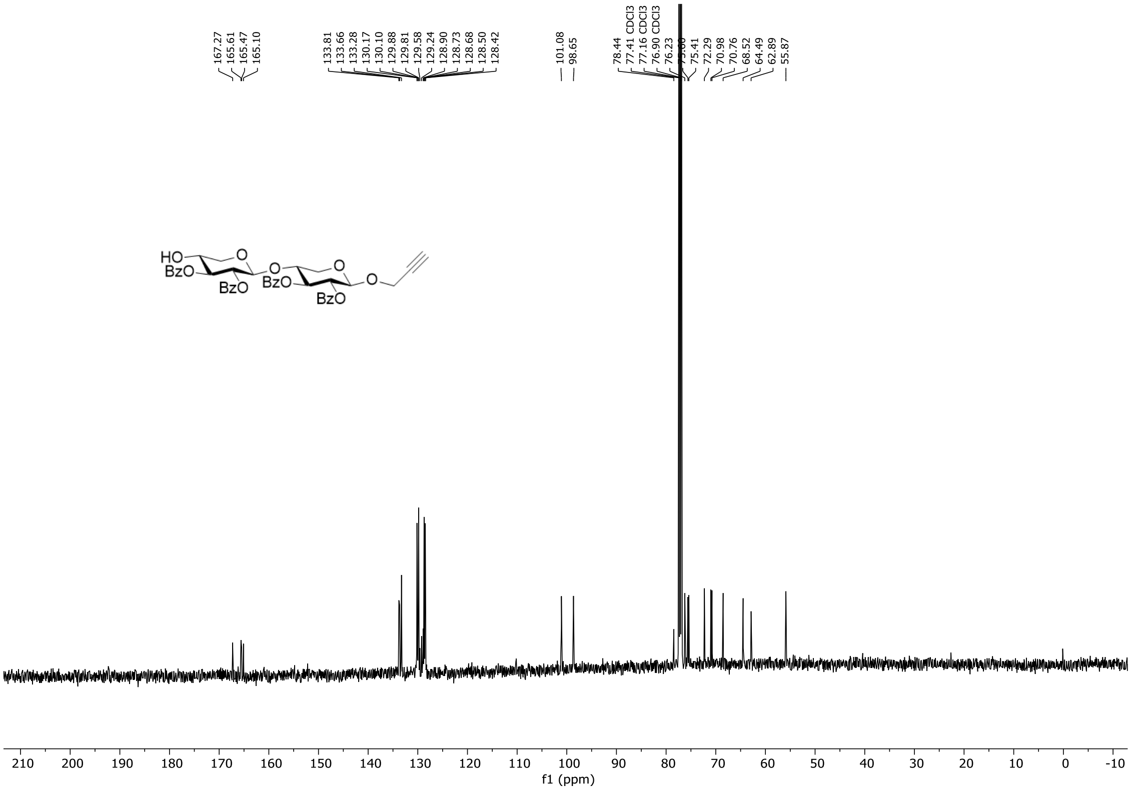

**Figure S11.** $^{13}$C NMR spectrum of compound **10** (126 MHz, $CDCl_3$, 298 K).

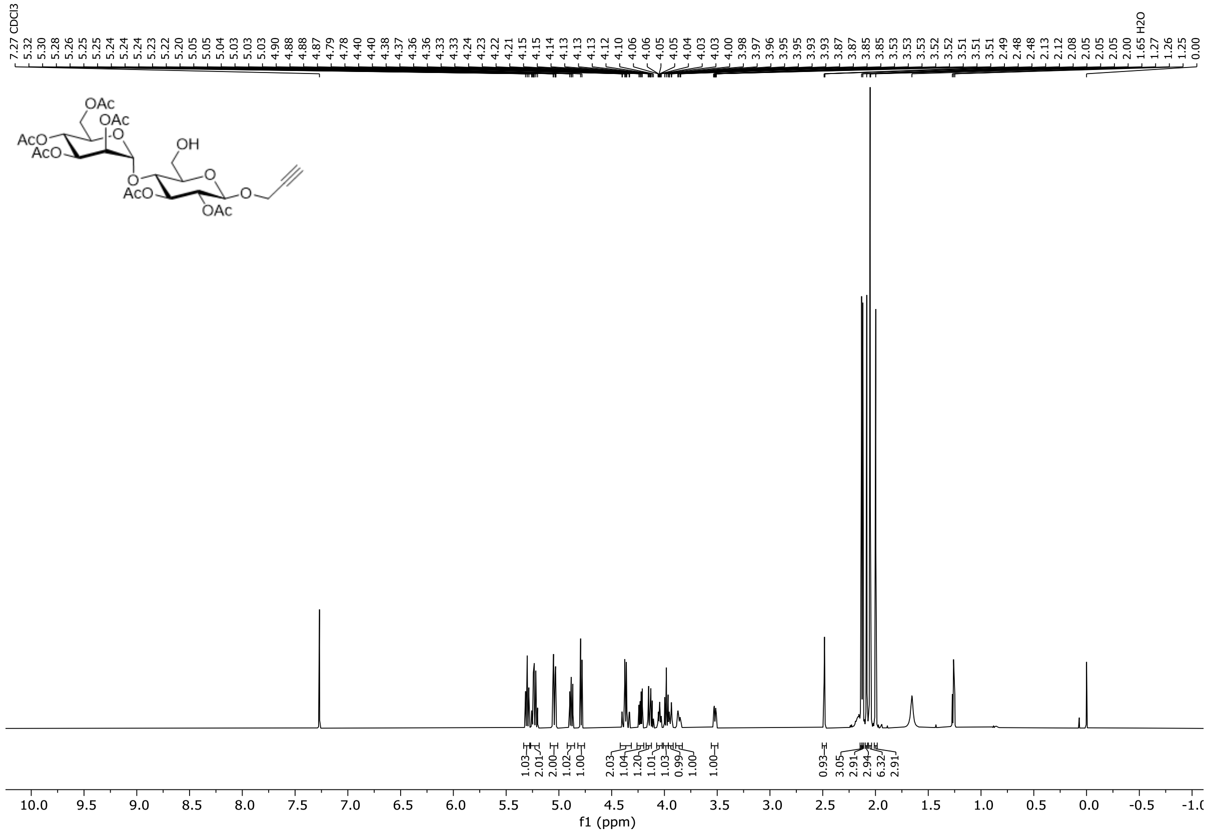

**Figure S12.** $^{1}$H NMR spectrum of compound **11** (600 MHz, $CDCl_3$, 298 K).

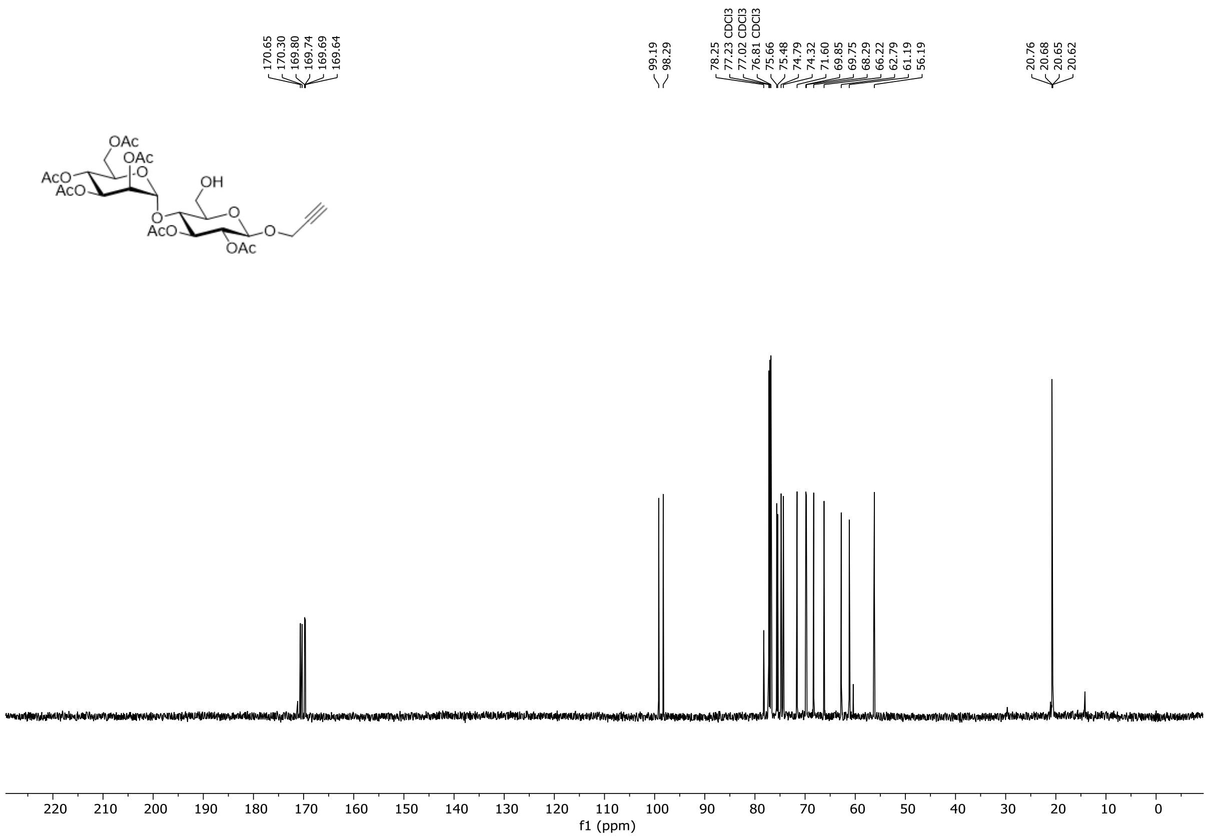

**Figure S13.** $^{13}$C NMR spectrum of compound **11** (151 MHz, $CDCl_3$, 298 K).

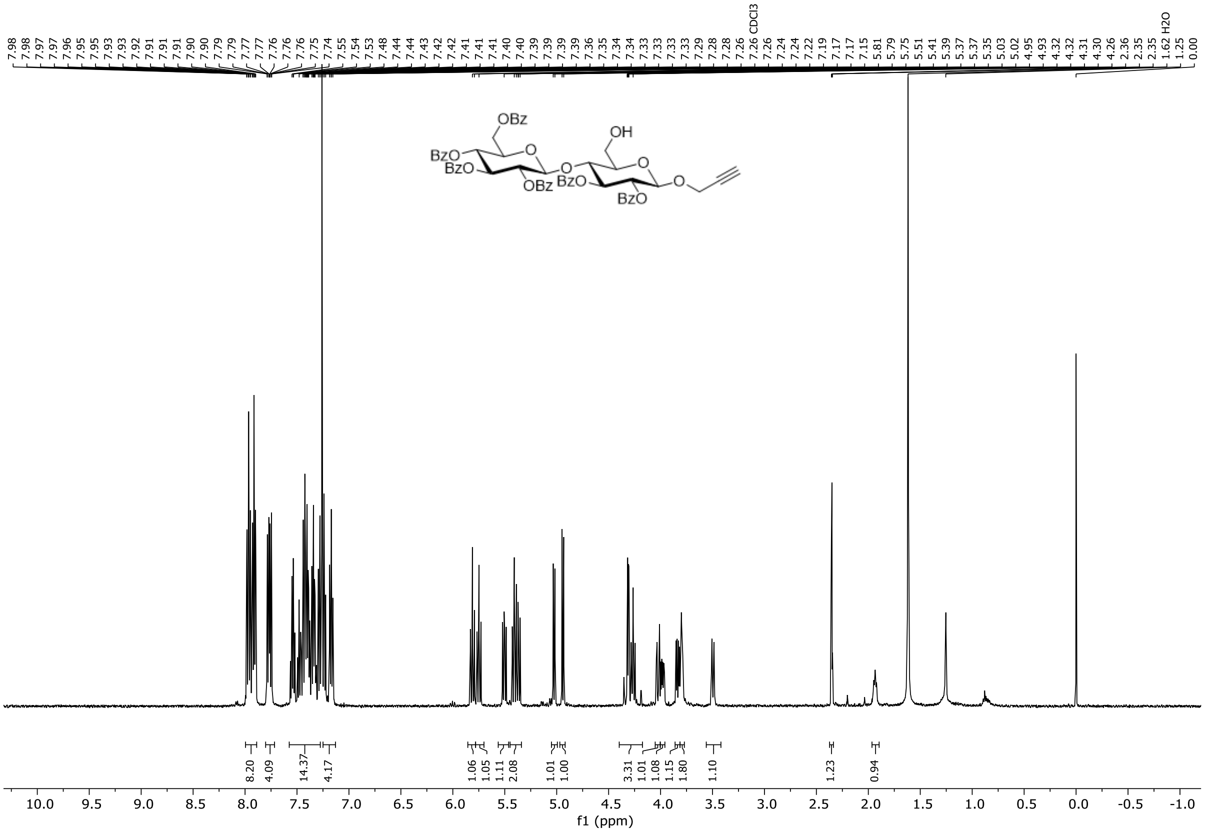

**Figure S14.** $^{1}H$ NMR spectrum of compound **12** (500 MHz, $CDCl_3$, 298 K).

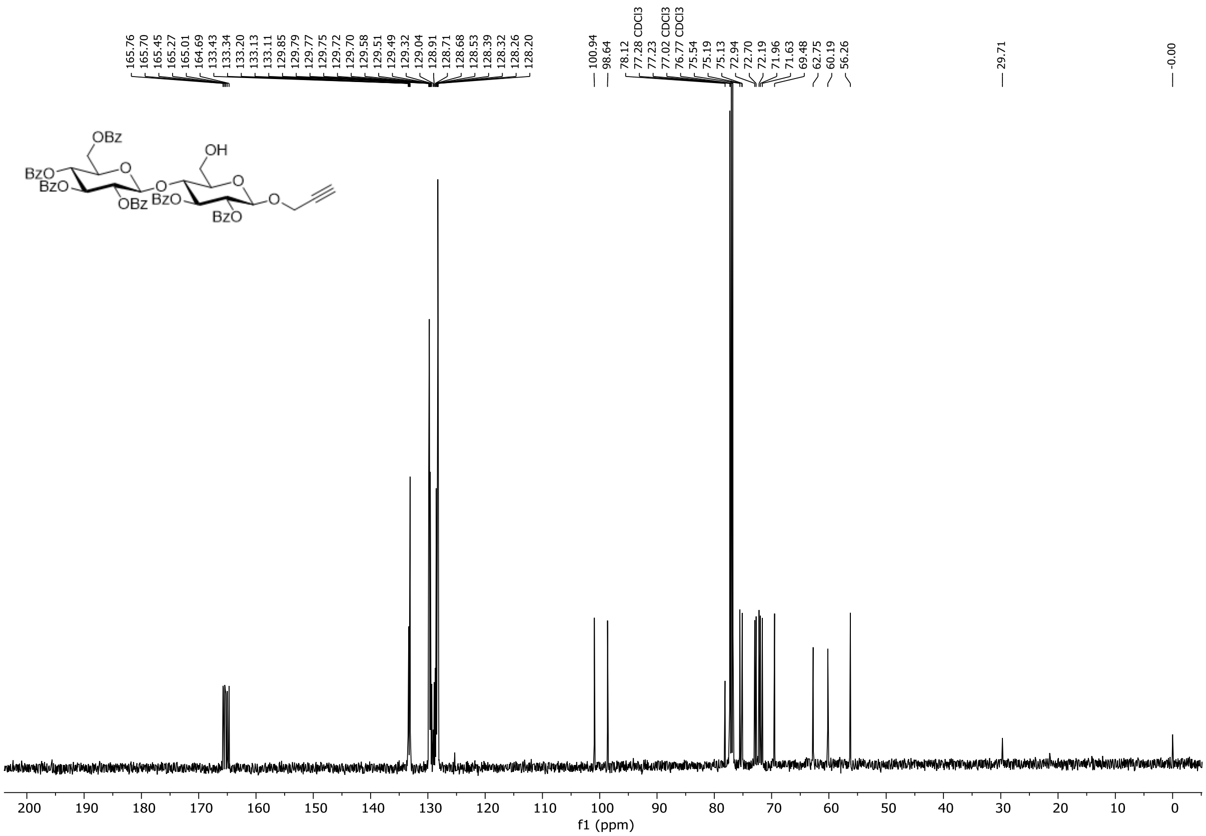

**Figure S15.** $^{13}C$ NMR spectrum of compound **12** (126 MHz, $CDCl_3$, 298 K).

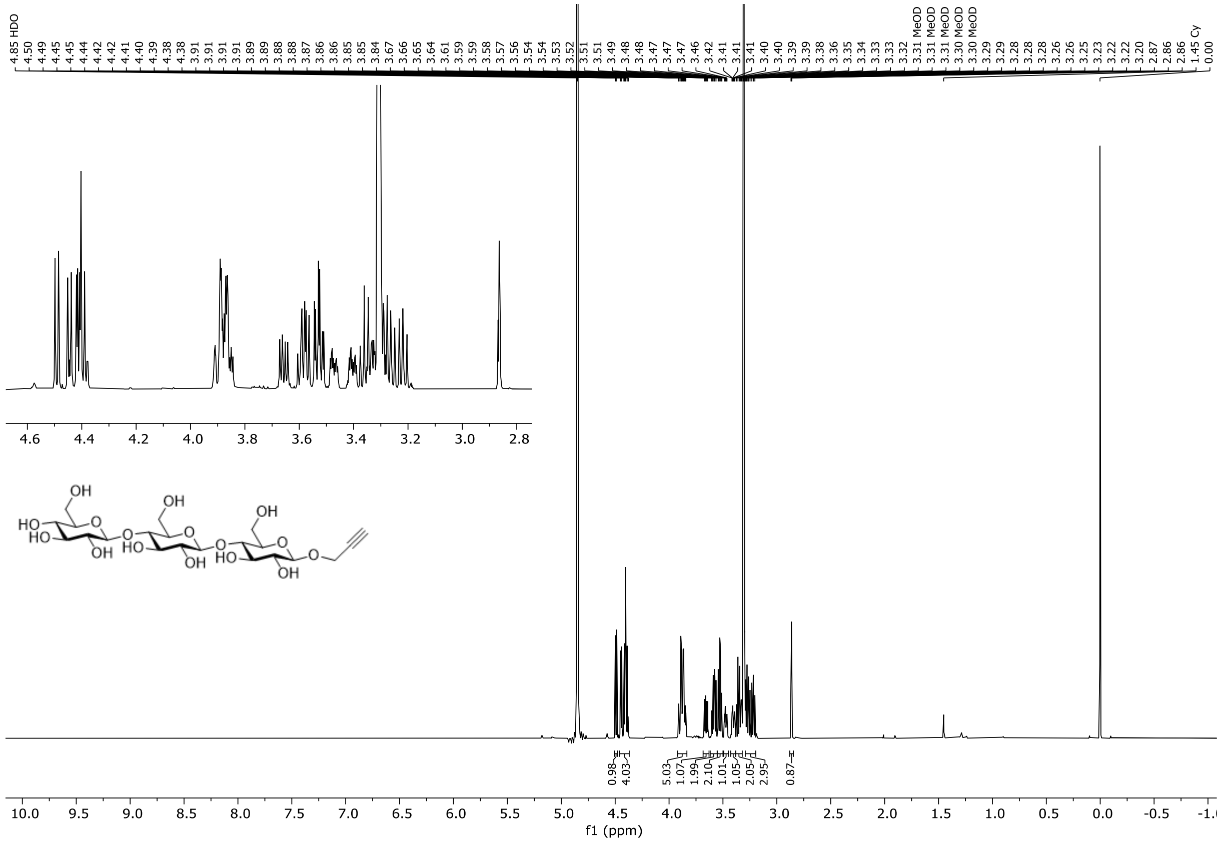

**Figure S16.** $^{1}$H NMR spectrum of compound **17** (600 MHz, methanol-d$_4$, 298 K).

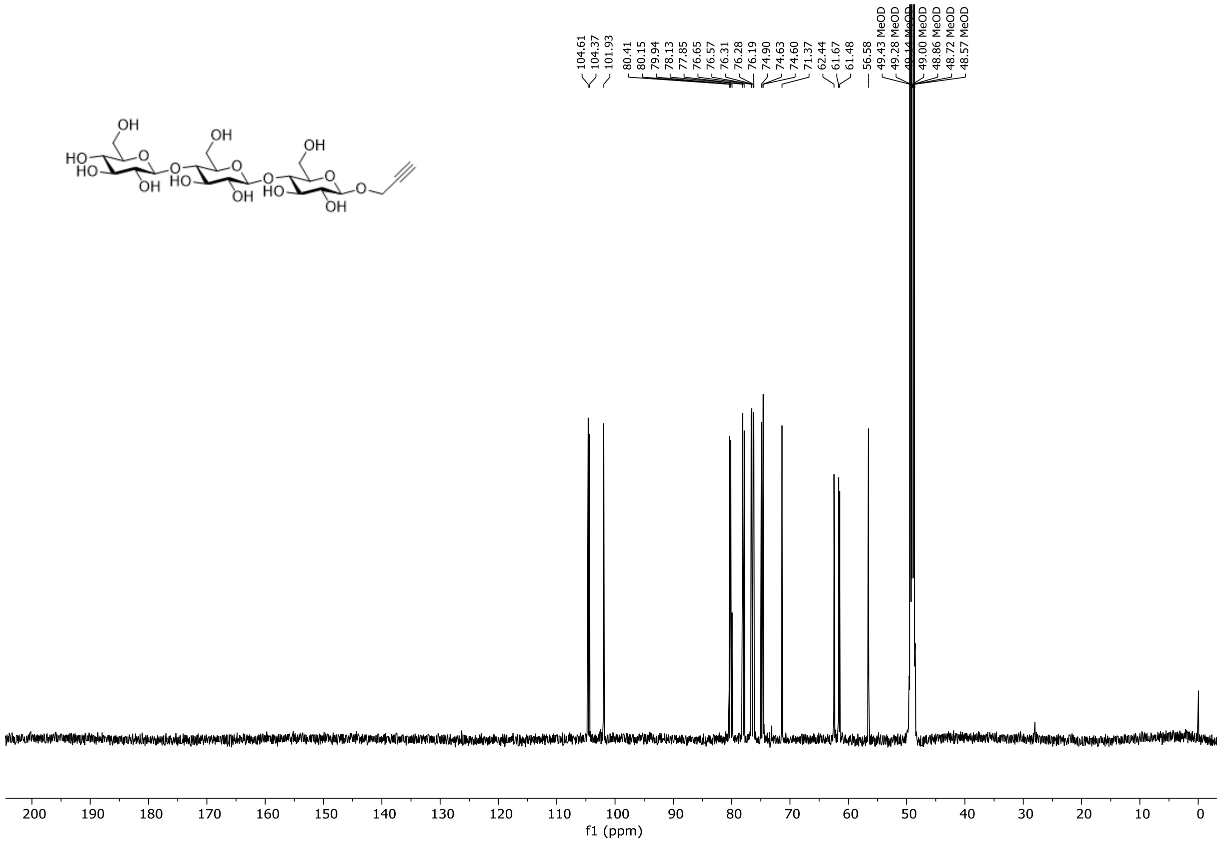

**Figure S17.** $^{13}$C NMR spectrum of compound **17** (151 MHz, methanol-d$_4$, 298 K).

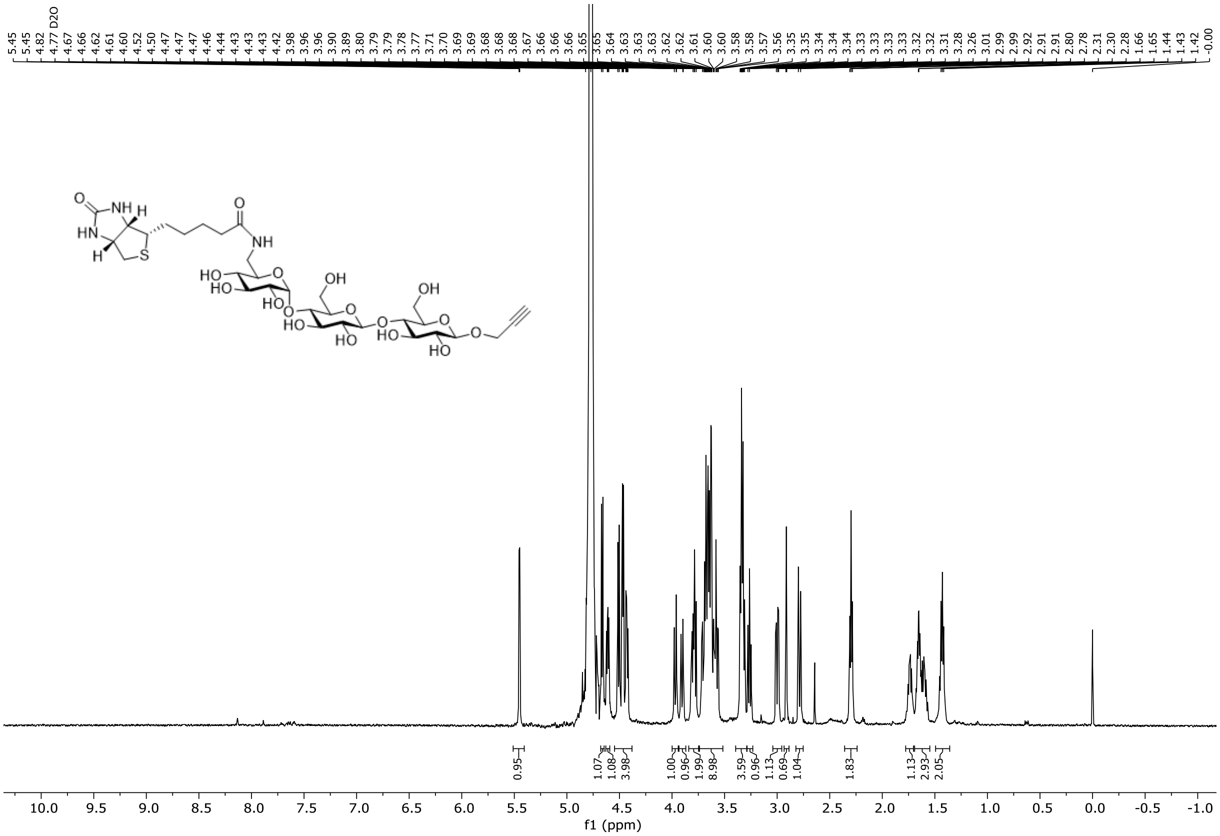

**Figure S18.** $^{1}$H NMR spectrum of compound **18** (600 MHz, $D_2O$, 298 K).

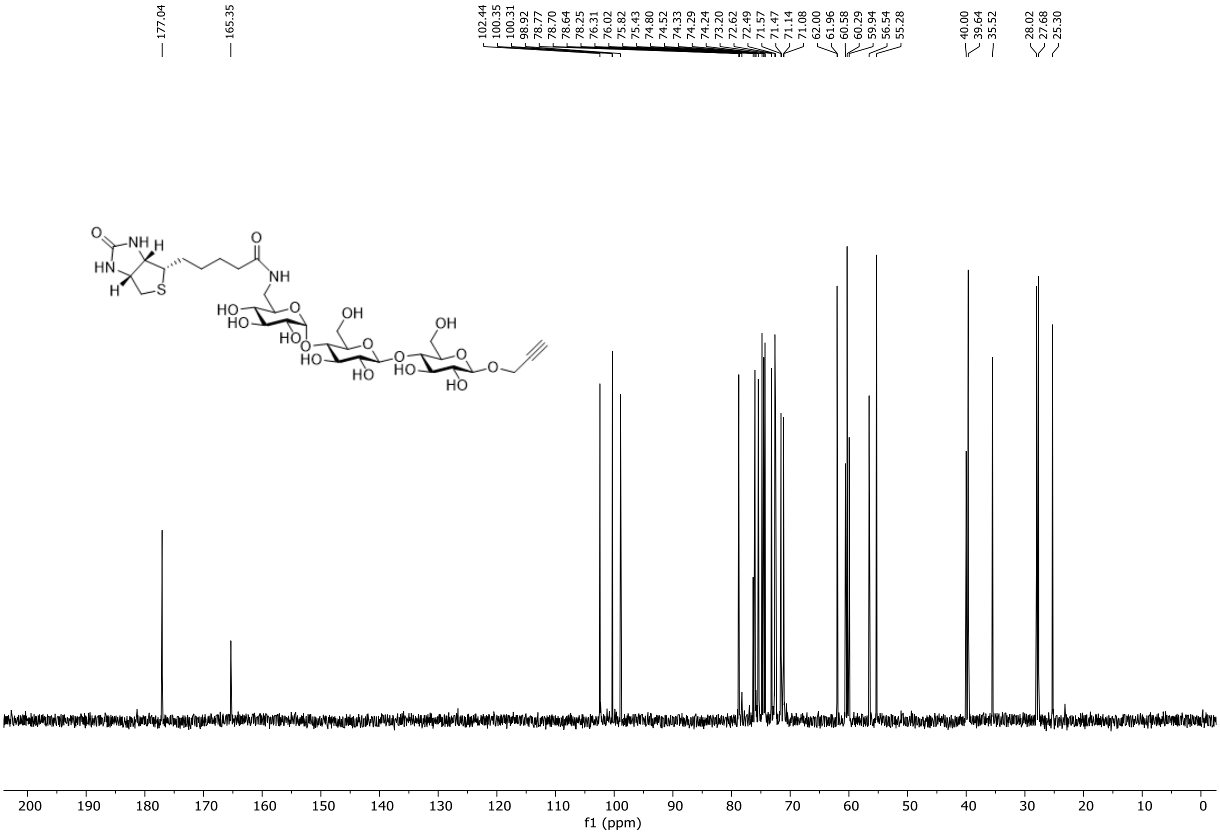

**Figure S19.** $^{13}$C NMR spectrum of compound **18** (151 MHz, $D_2O$, 298 K).

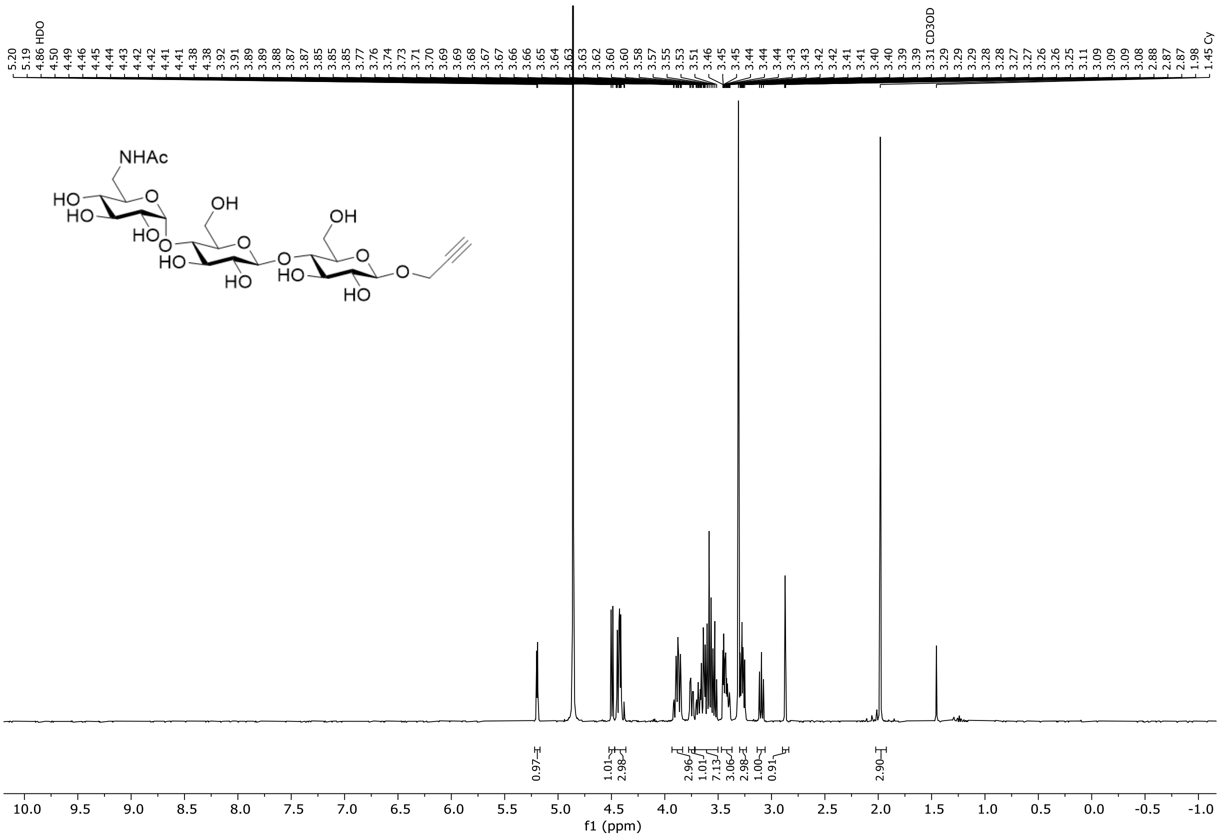

**Figure S20.** $^{1}H$ NMR spectrum of compound **19** (500 MHz, methanol-$d_4$, 298 K).

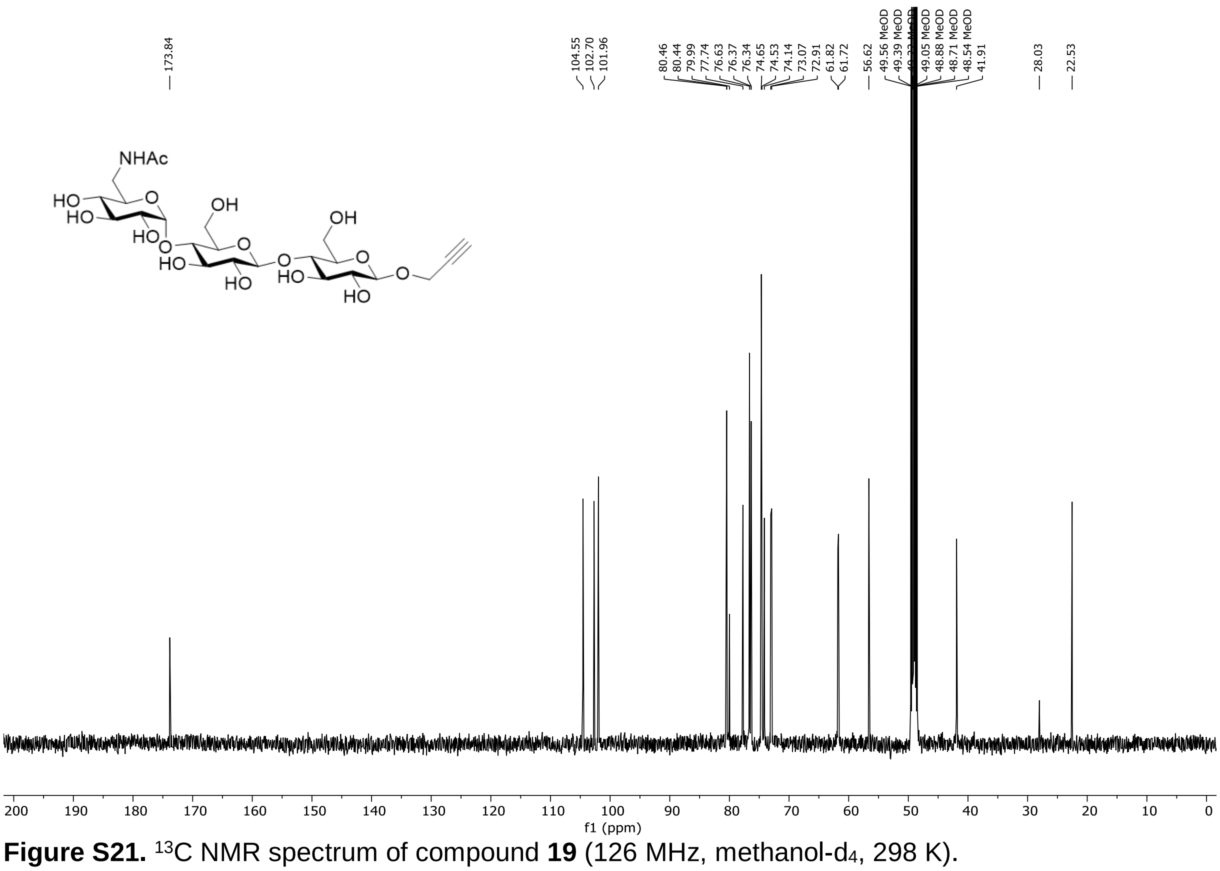

**Figure S21.** $^{13}C$ NMR spectrum of compound **19** (126 MHz, methanol-$d_4$, 298 K).

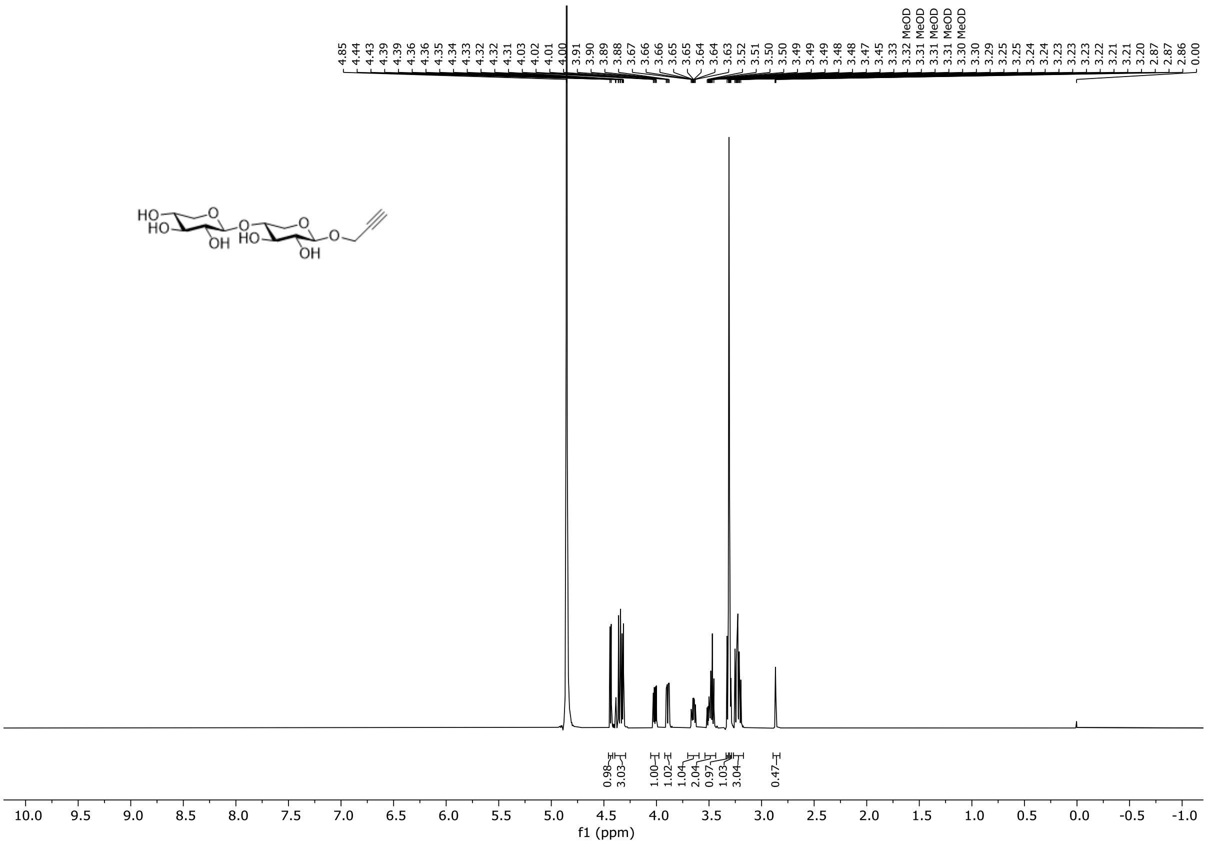

**Figure S22.** $^{1}H$ NMR spectrum of compound **21** (600 MHz, methanol-d$_4$, 298 K).

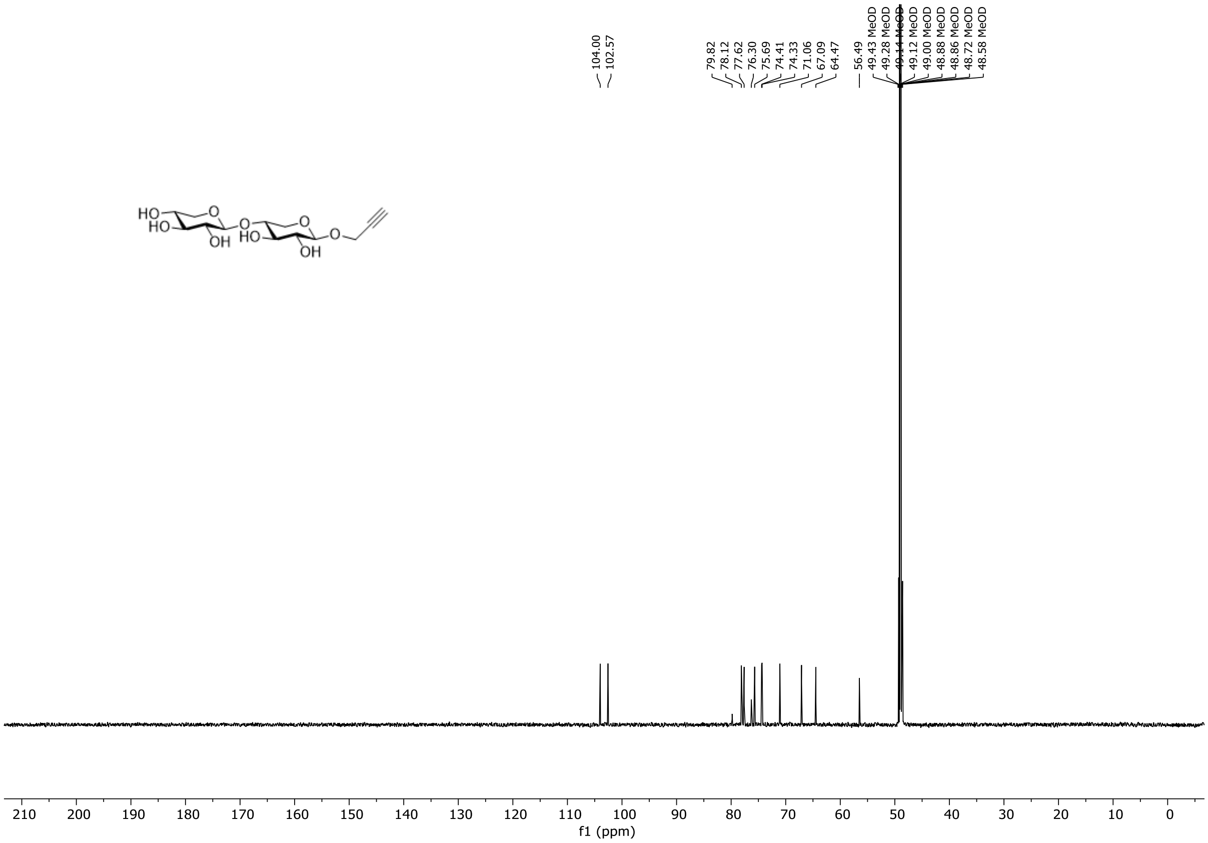

**Figure S23.** $^{13}C$ NMR spectrum of compound **21** (151 MHz, methanol-d$_4$, 298 K).

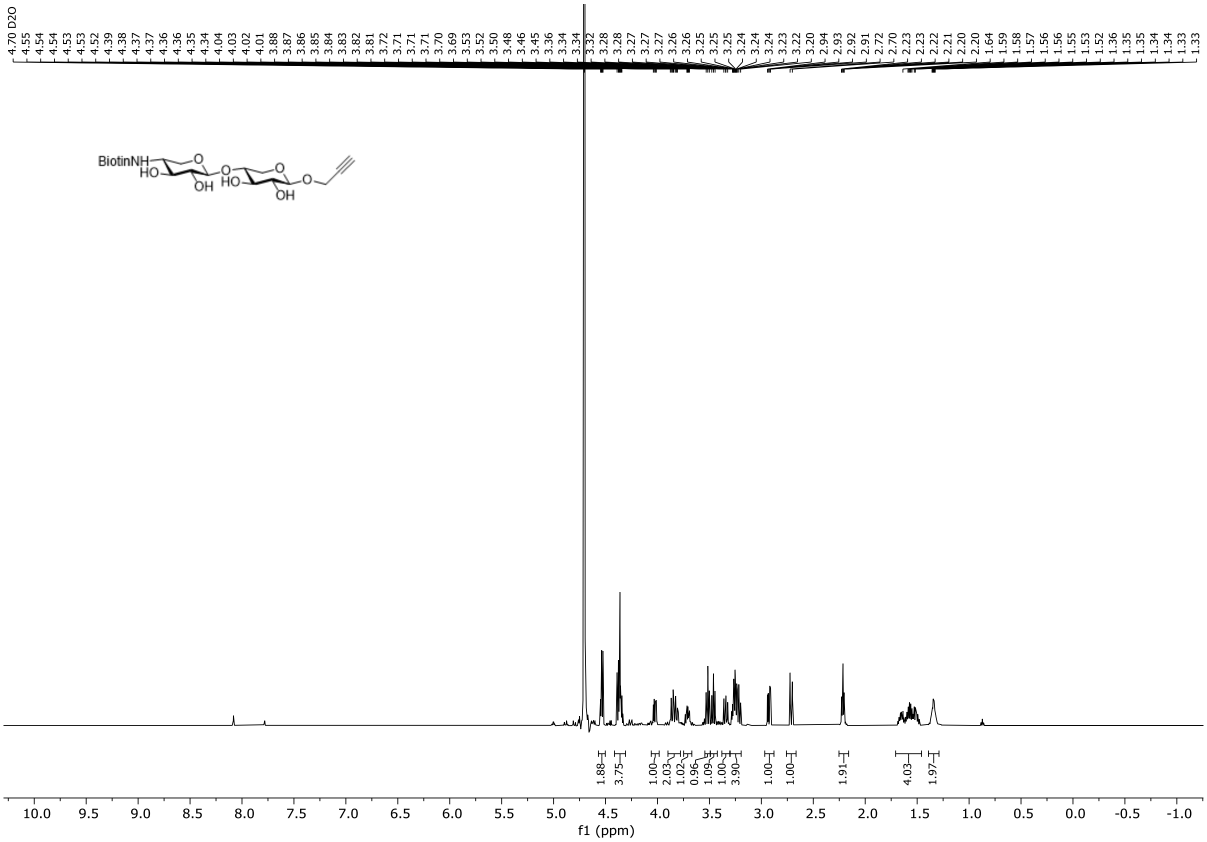

**Figure S24.** $^{1}H$ NMR spectrum of compound **22** (600 MHz, $D_2O$, 298 K).

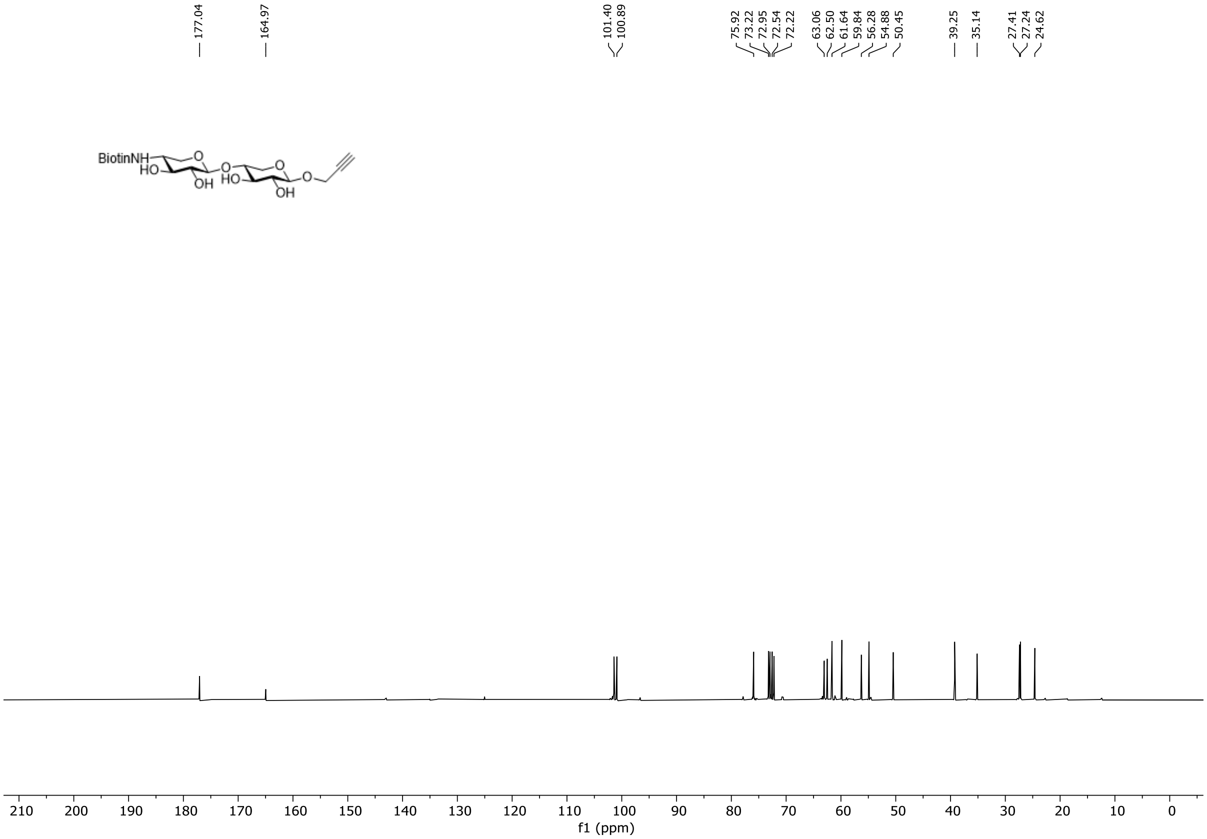

**Figure S25.** $^{13}C$ NMR spectrum of compound **22** (151 MHz, $D_2O$, 298 K).

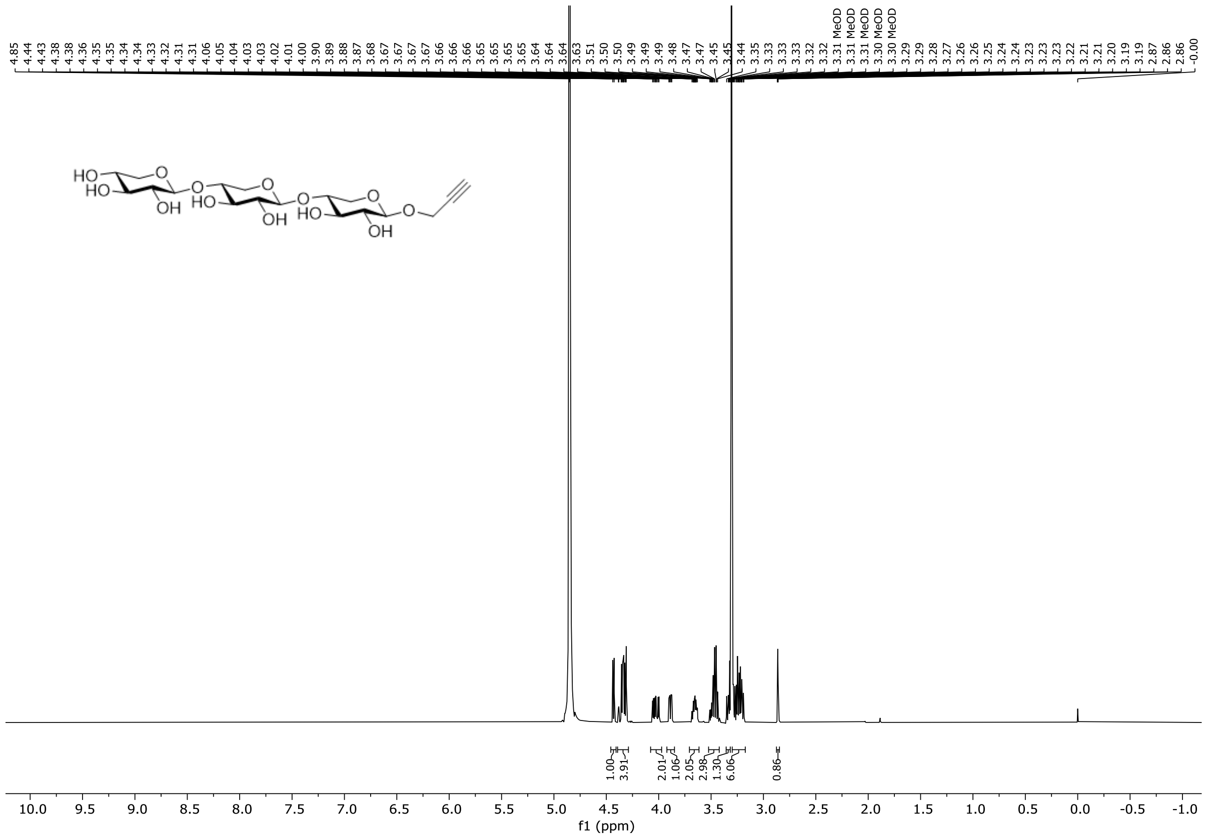

**Figure S26.** $^{1}H$ NMR spectrum of compound **23** (600 MHz, methanol-$d_4$, 298 K).

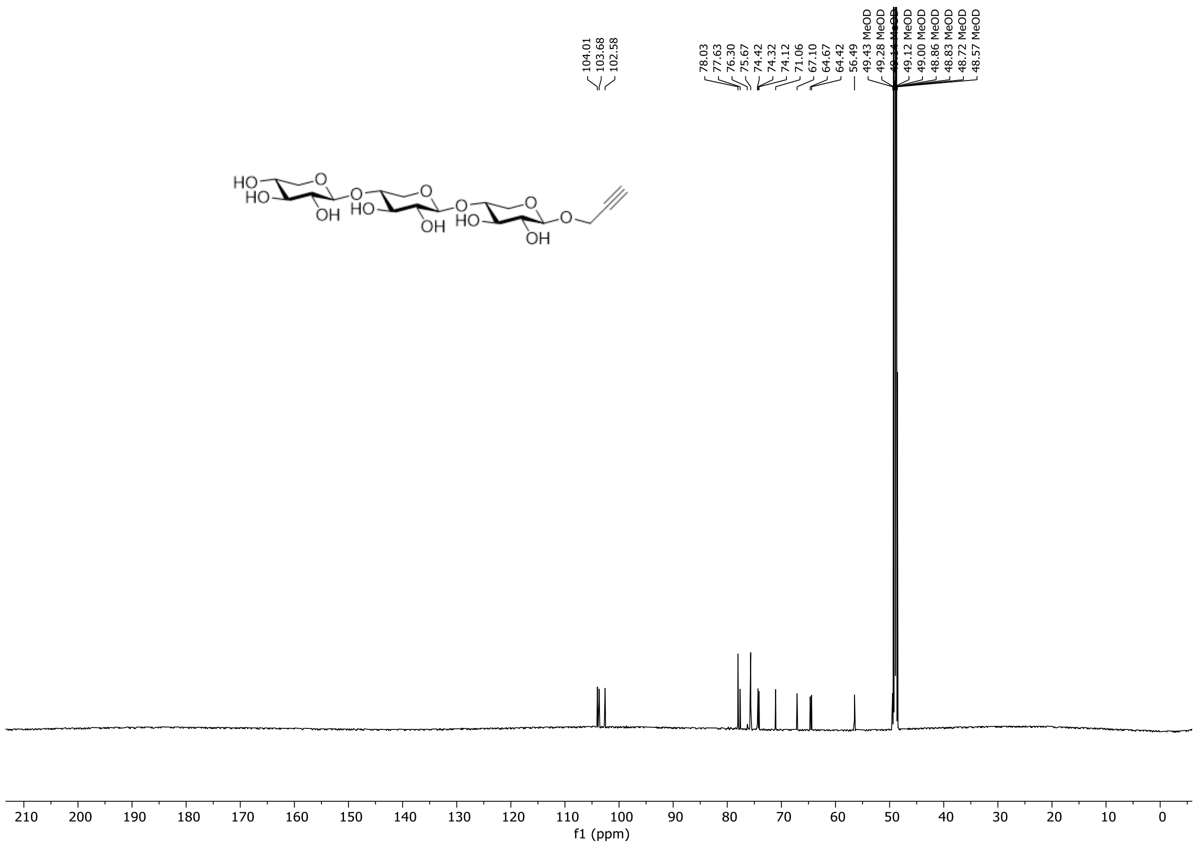

**Figure S27.** $^{13}C$ NMR spectrum of compound **23** (151 MHz, methanol-$d_4$, 298 K).

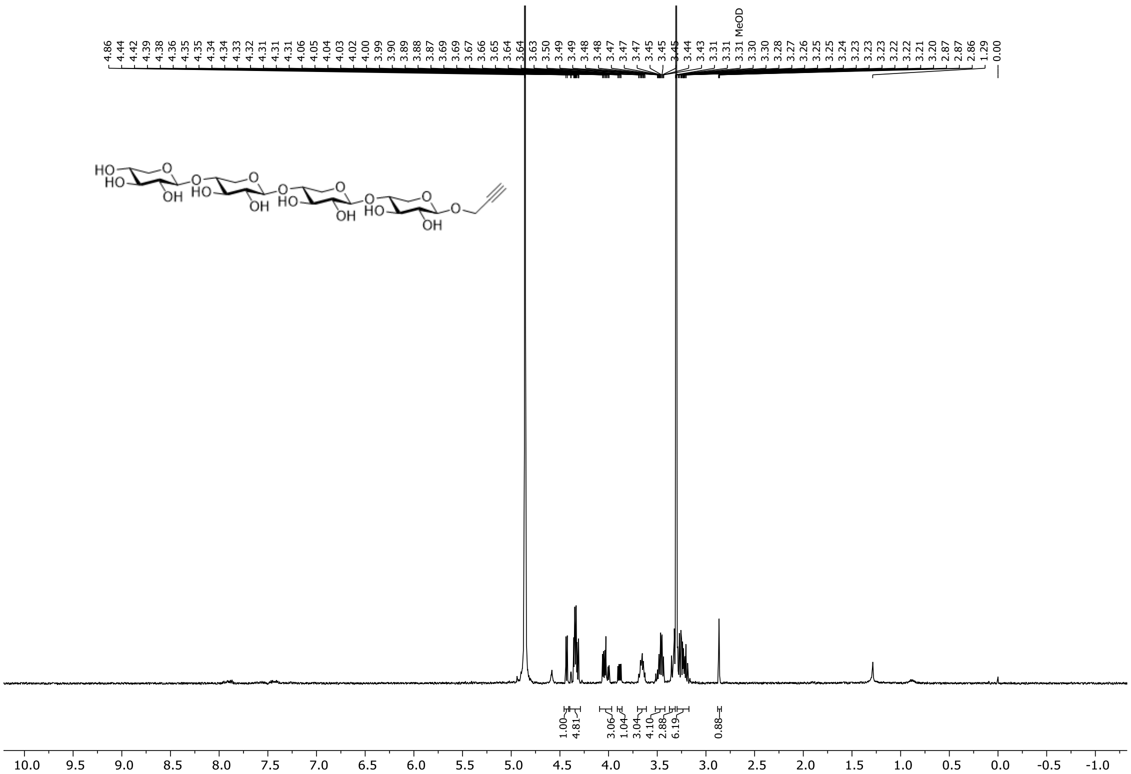

**Figure S28.** $^{1}$H NMR spectrum of compound **24** (500 MHz, methanol-$d_4$, 298 K).

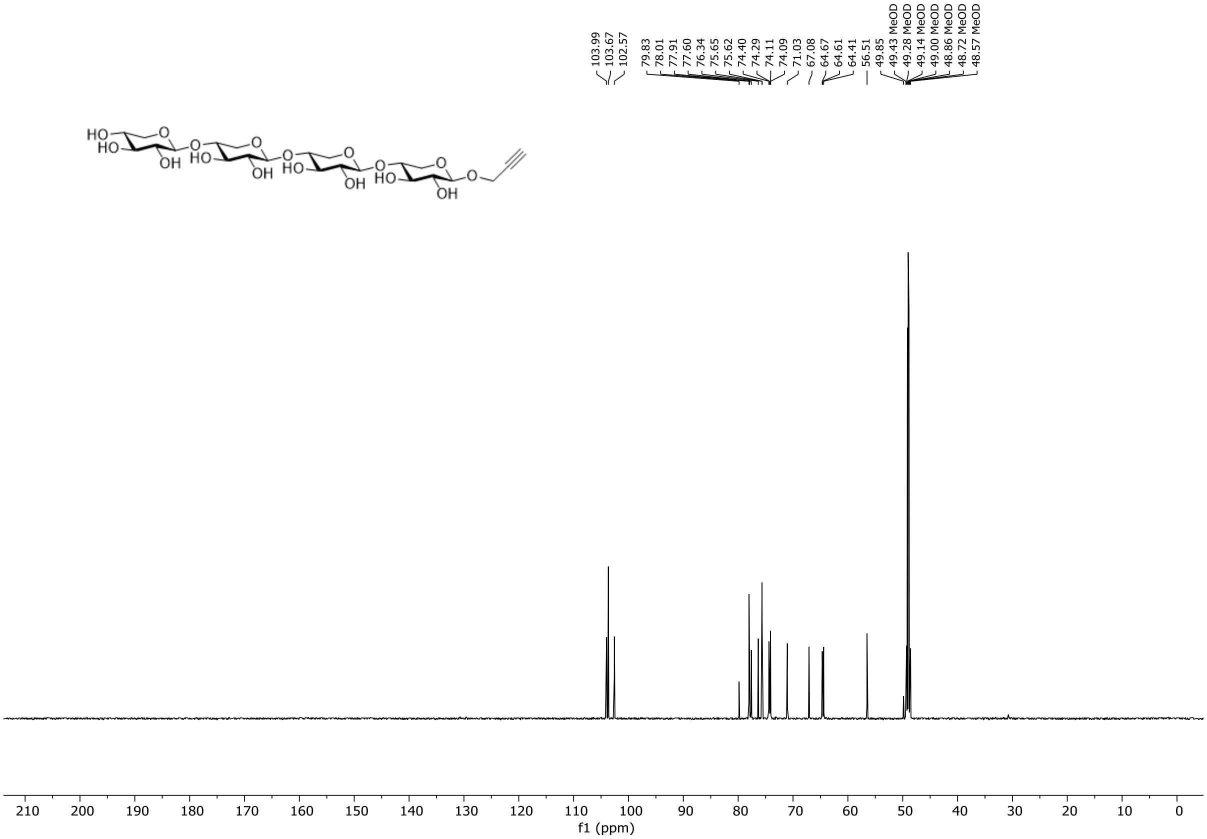

**Figure S29.** $^{13}$C NMR spectrum of compound **24** (151 MHz, methanol-$d_4$, 298 K).

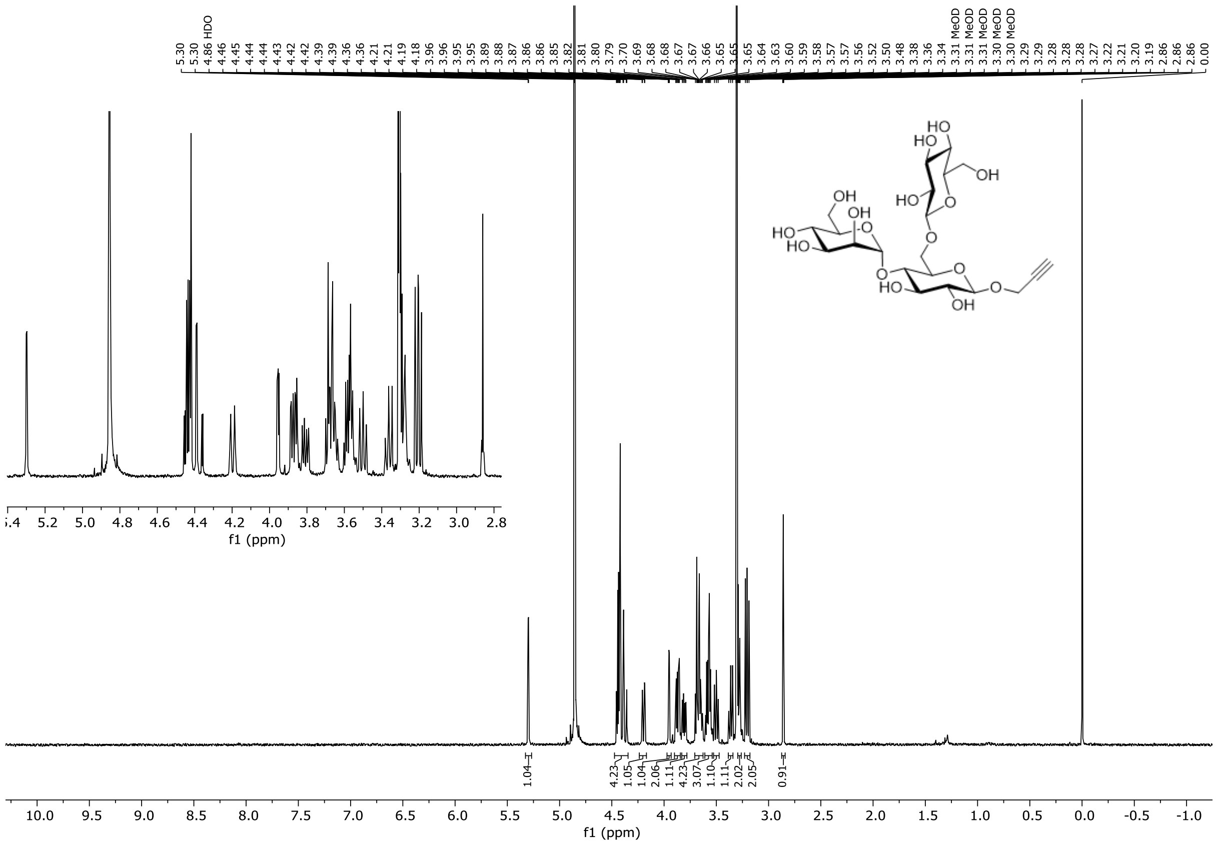

**Figure S30.** $^{1}$H NMR spectrum of compound **25** (500 MHz, methanol-d$_4$, 298 K).

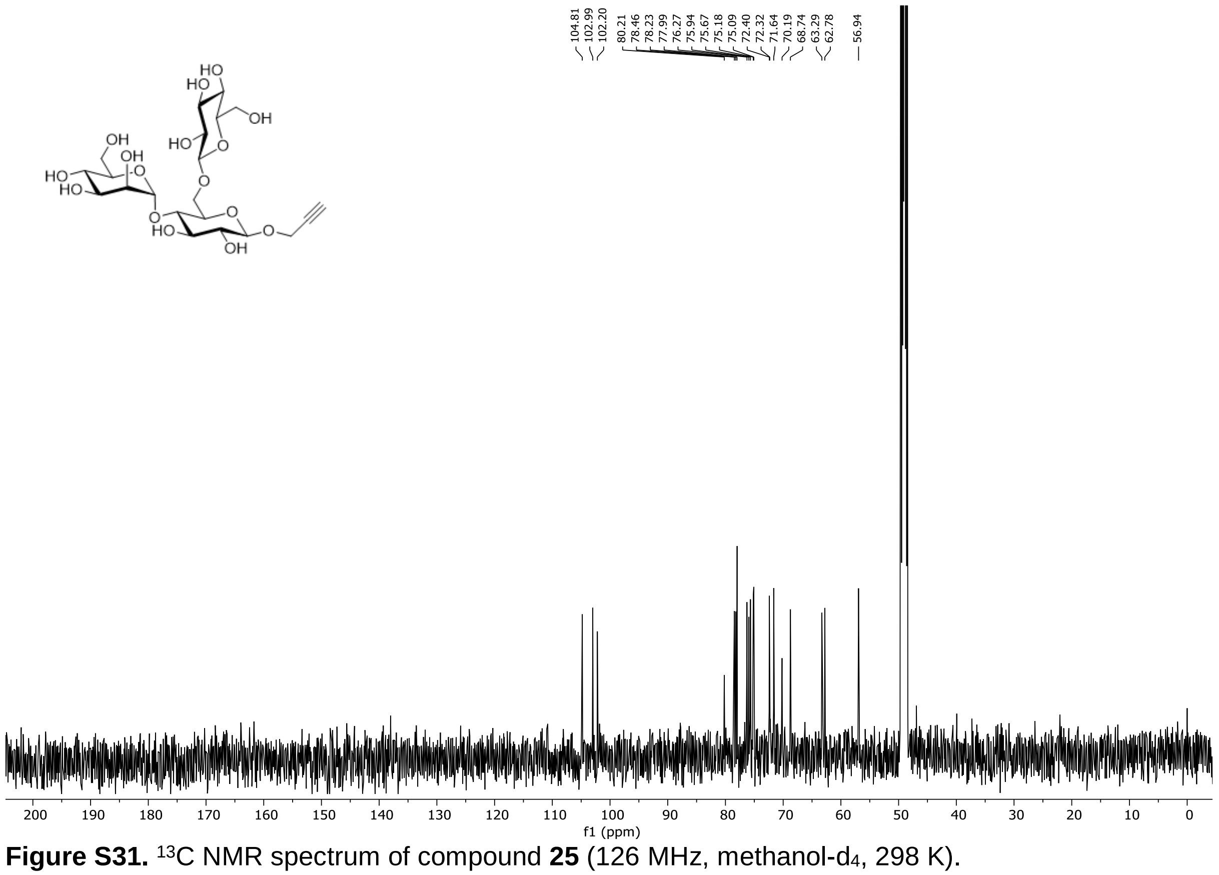

**Figure S31.** $^{13}$C NMR spectrum of compound **25** (126 MHz, methanol-d$_4$, 298 K).

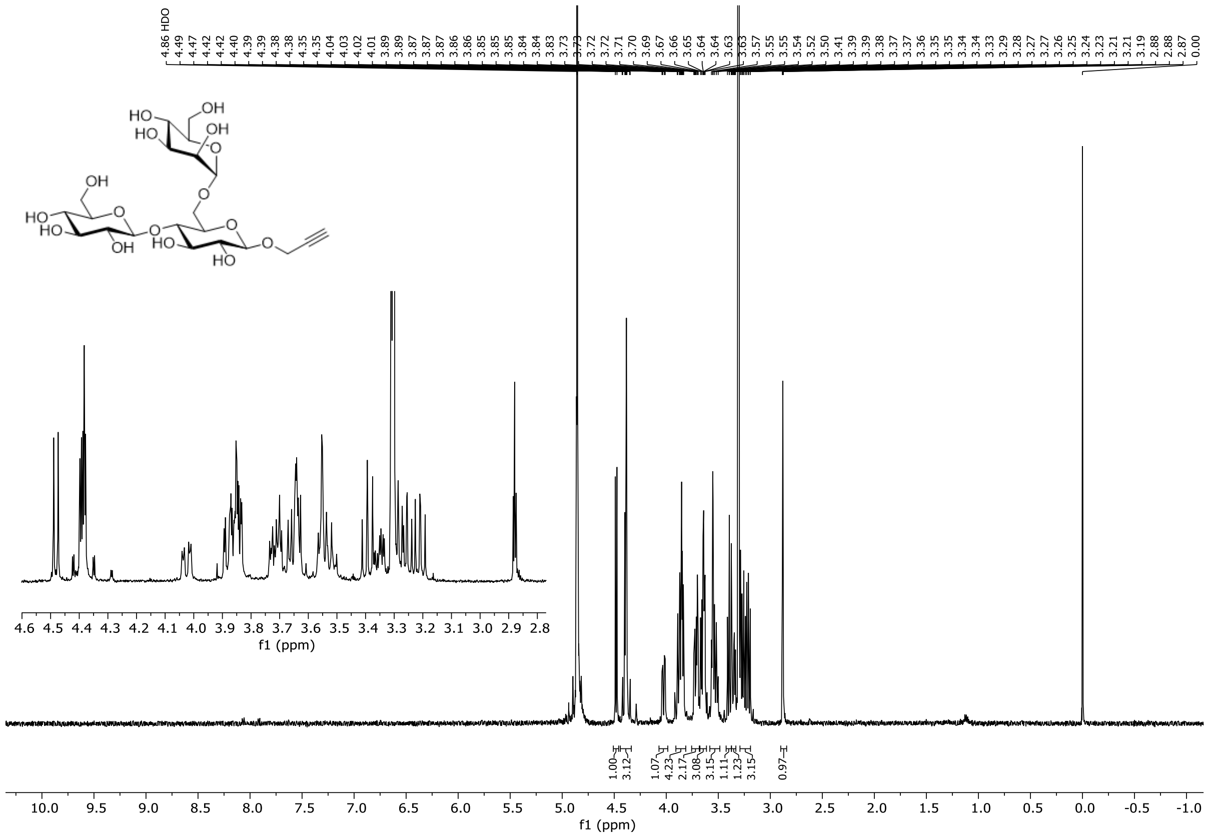

**Figure S32.** $^{1}H$ NMR spectrum of compound **26** (500 MHz, methanol-$d_4$, 298 K).

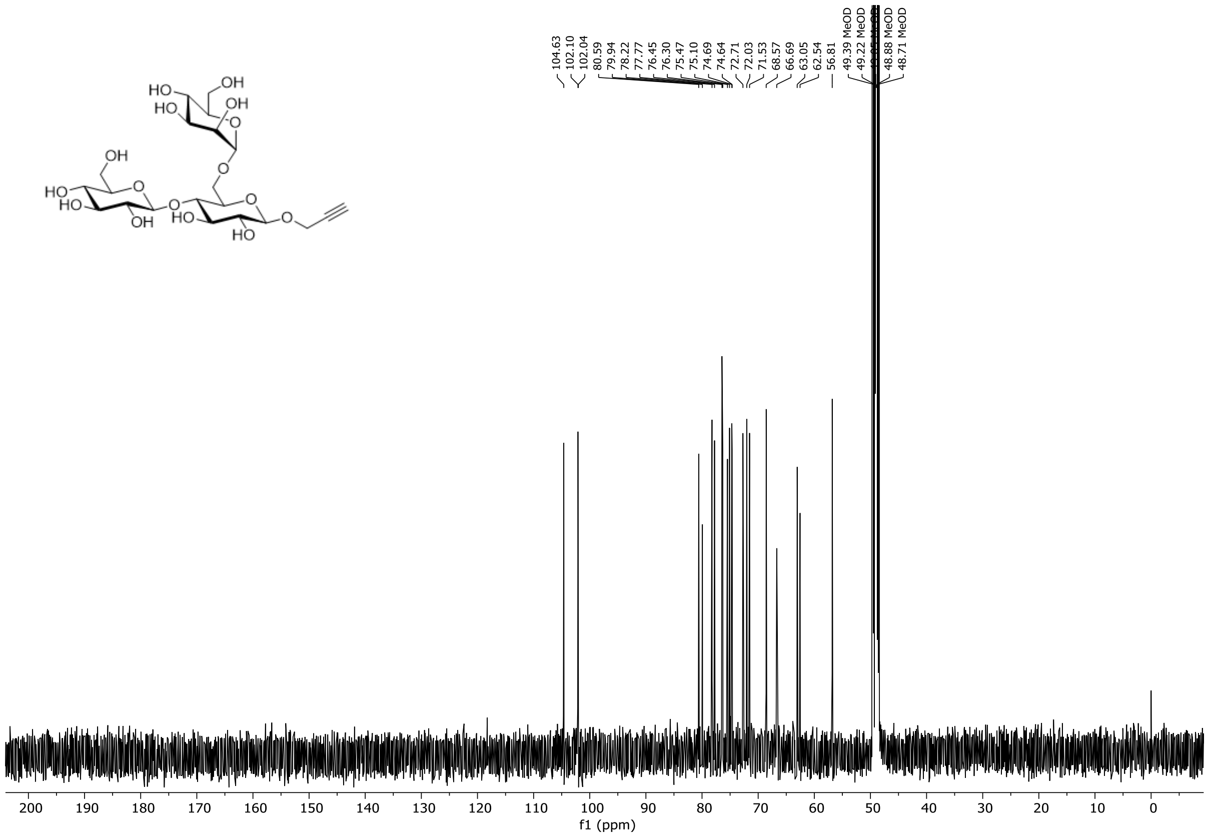

**Figure S33.** $^{13}C$ NMR spectrum of compound **26** (126 MHz, methanol-$d_4$, 298 K).

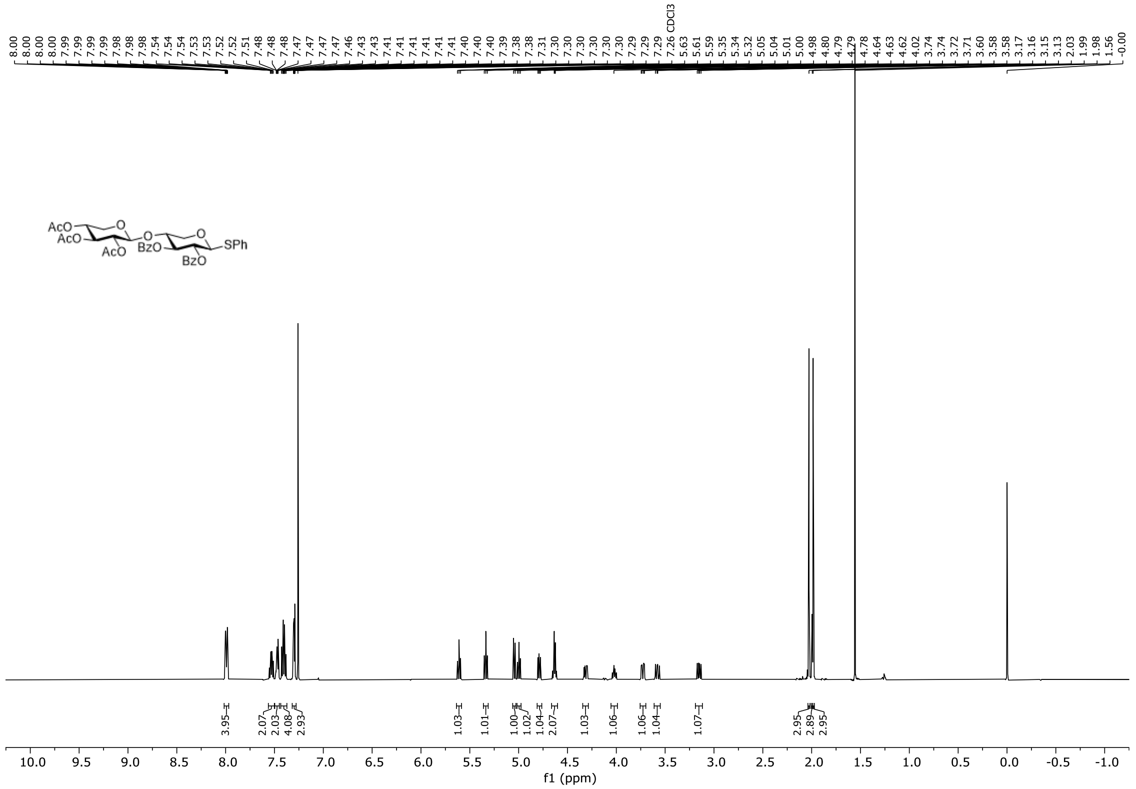

**Figure S34.** $^{1}$H NMR spectrum of compound **S6** (500 MHz, $CDCl_3$, 298 K).

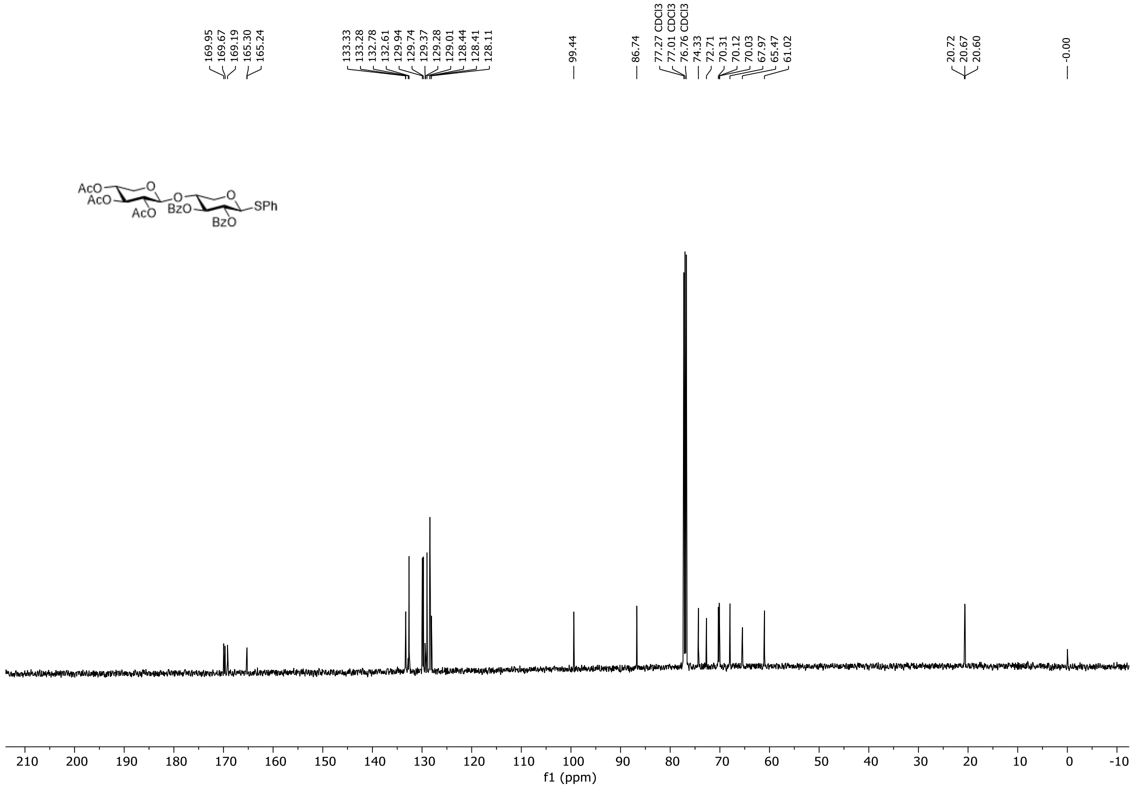

**Figure S35.** $^{13}$C NMR spectrum of compound **S6** (126 MHz, $CDCl_3$, 298 K).

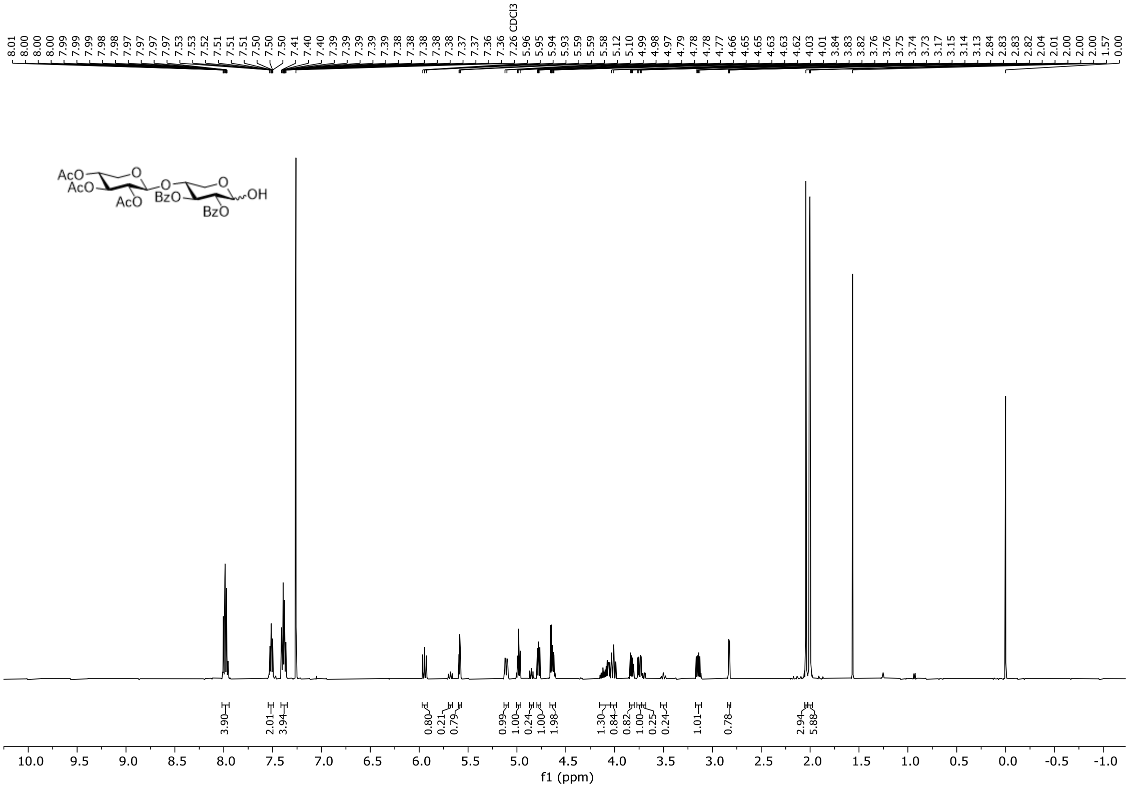

**Figure S36.** $^{1}$H NMR spectrum of compound **S7** (500 MHz, $CDCl_3$, 298 K).

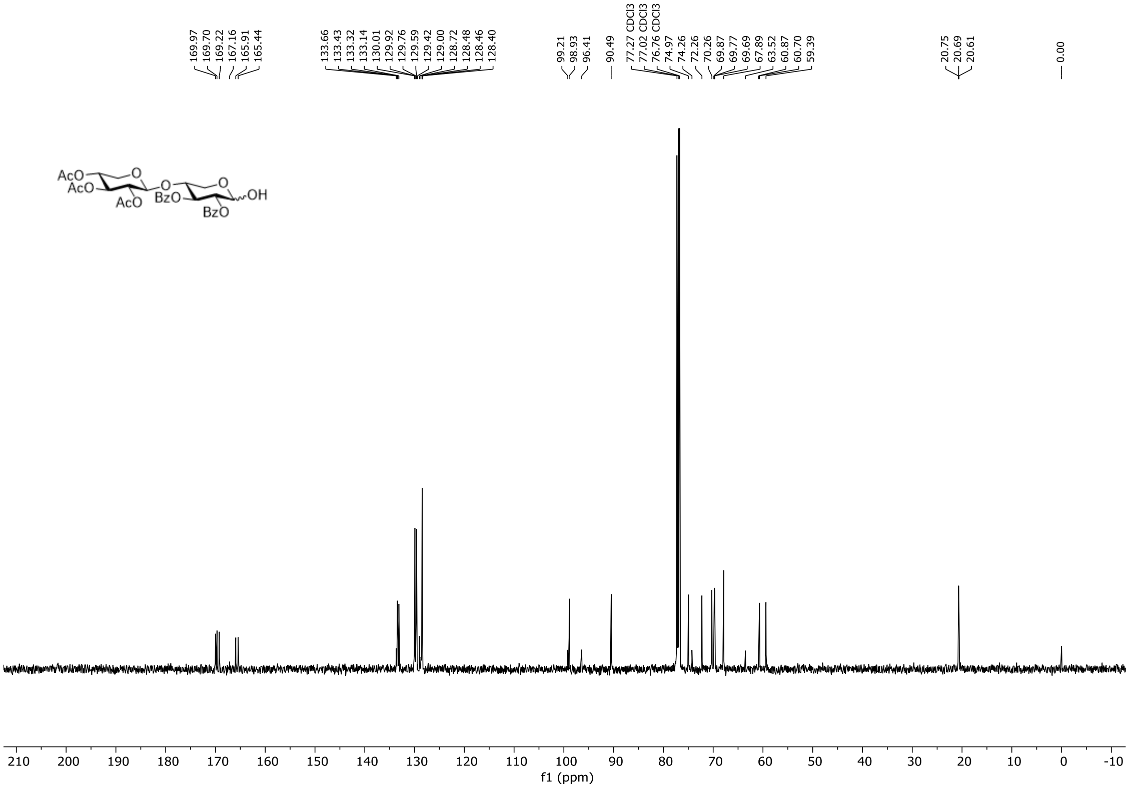

**Figure S37.** $^{13}$C NMR spectrum of compound **S7** (126 MHz, $CDCl_3$, 298 K).

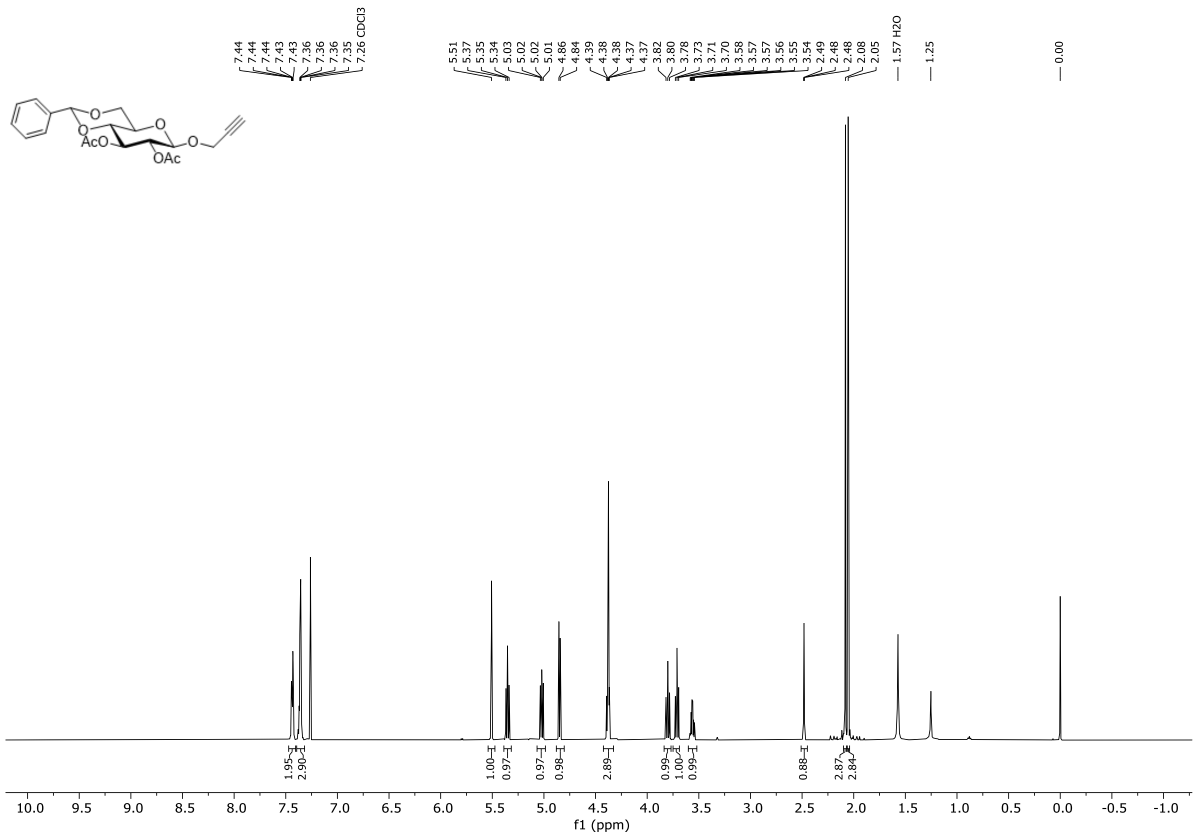

Figure S38. $^{1}H$ NMR spectrum of compound **S8** (600 MHz, $CDCl_3$, 298 K).

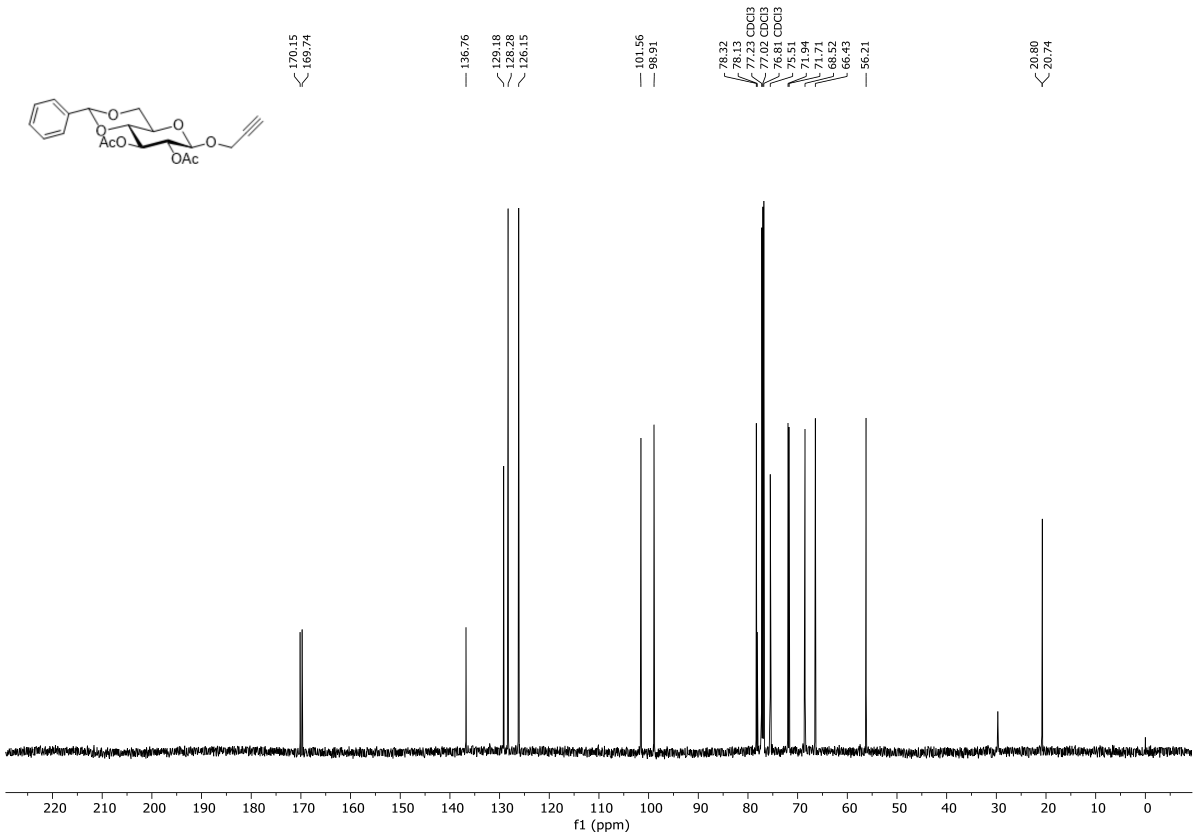

Figure S39. $^{13}C$ NMR spectrum of compound **S8** (151 MHz, $CDCl_3$, 298 K).

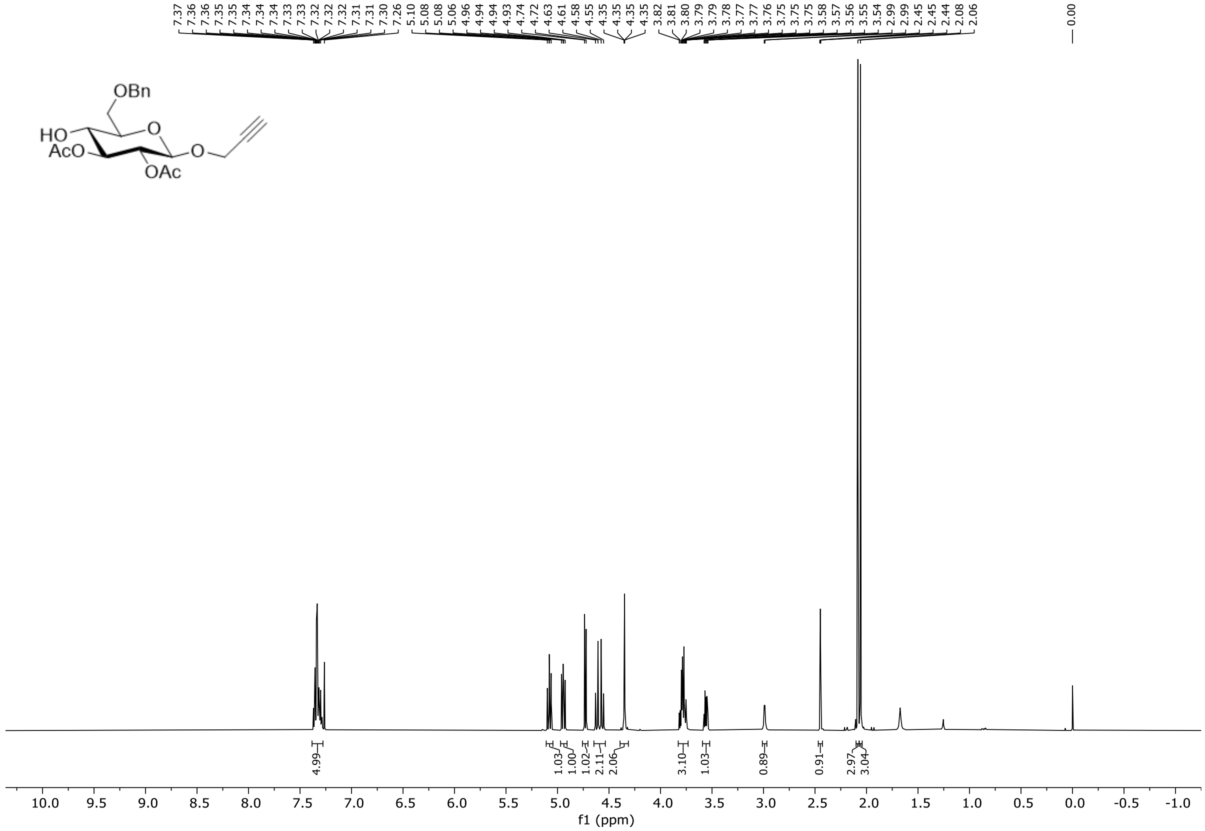

**Figure S40.** $^{1}$H NMR spectrum of compound **S9** (500 MHz, $CDCl_3$, 298 K).

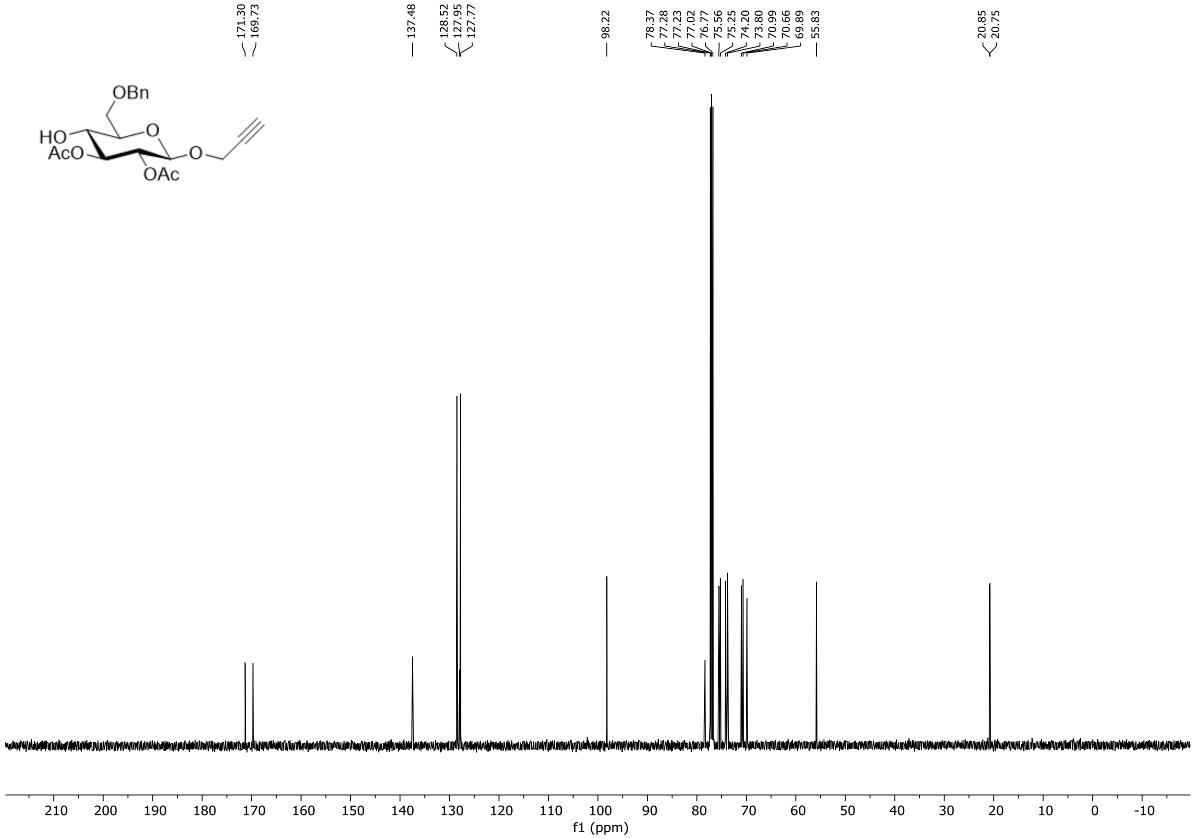

**Figure S41.** $^{13}$C NMR spectrum of compound **S9** (126 MHz, $CDCl_3$, 298 K).

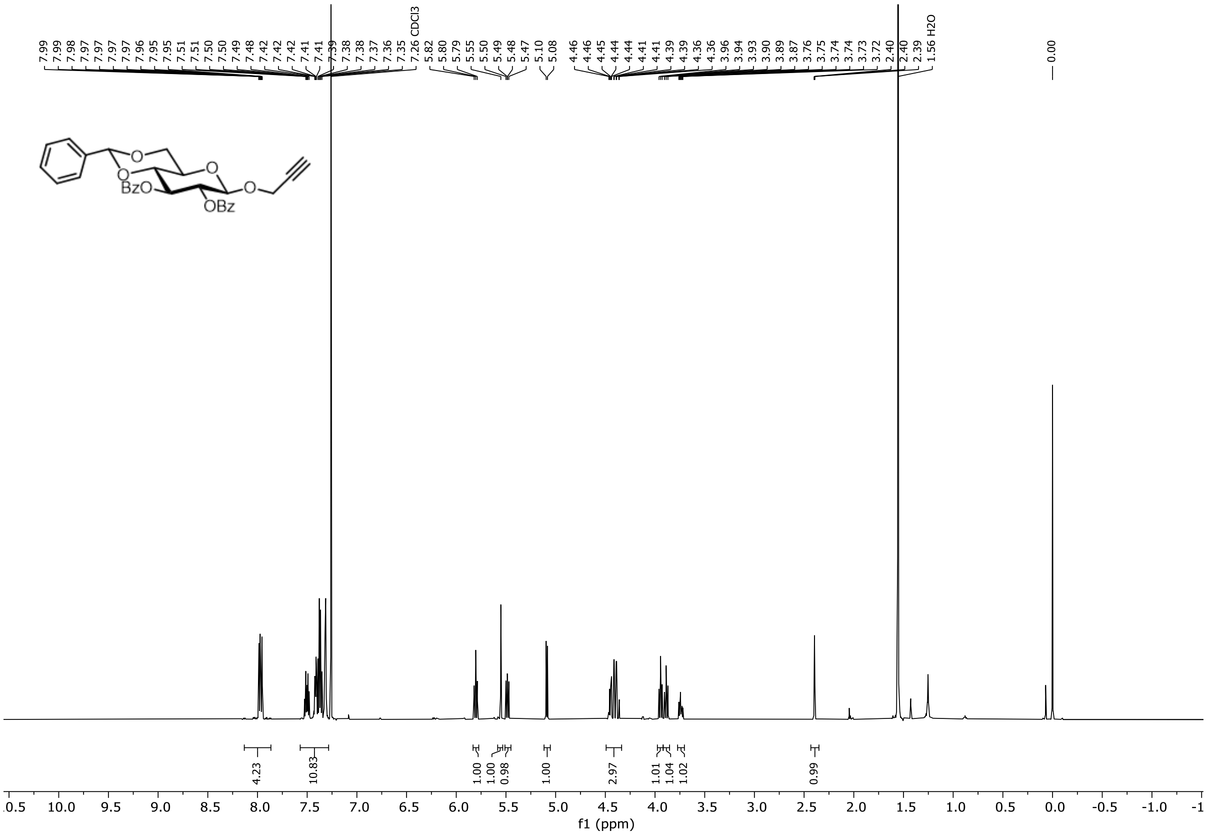

Figure S42. $^{1}H$ NMR spectrum of compound **S10** (600 MHz, $CDCl_3$, 298 K).

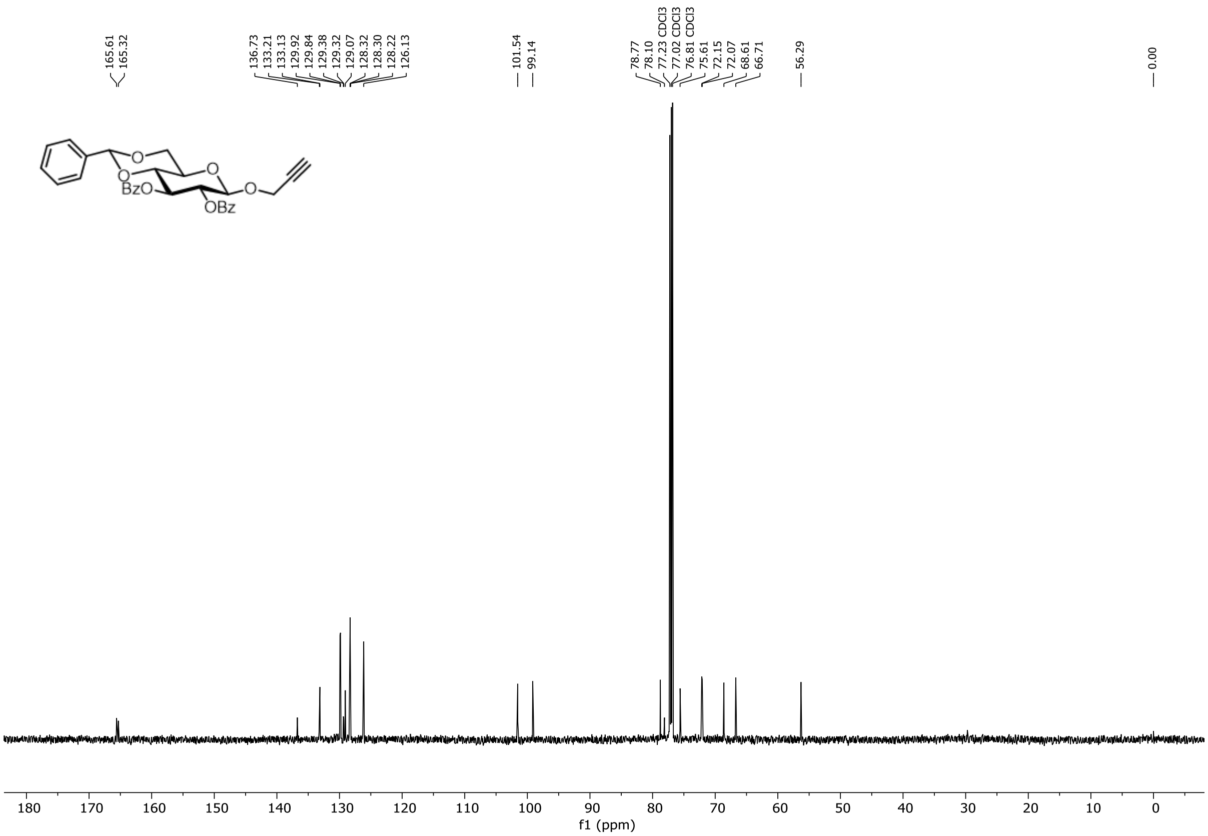

Figure S43. $^{13}C$ NMR spectrum of compound **S10** (151 MHz, $CDCl_3$, 298 K).

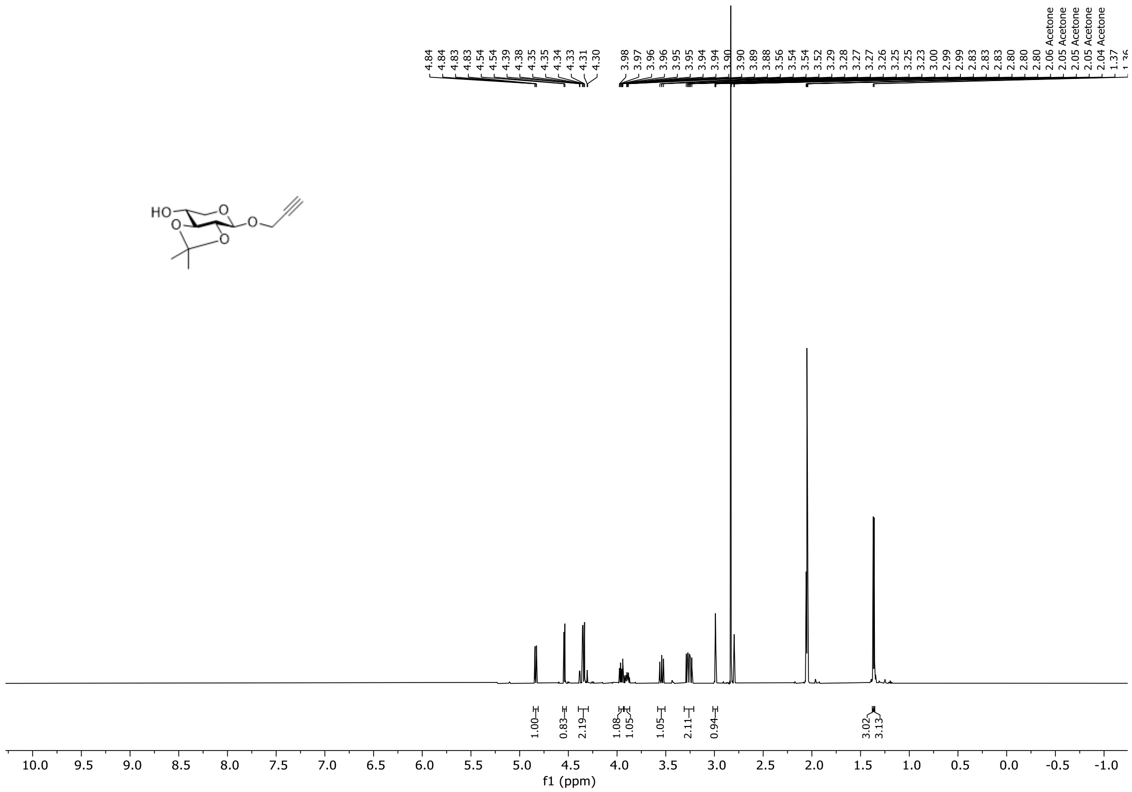

**Figure S44.** $^{1}H$ NMR spectrum of compound **S11** (500 MHz, acetone-$d_6$, 298 K).

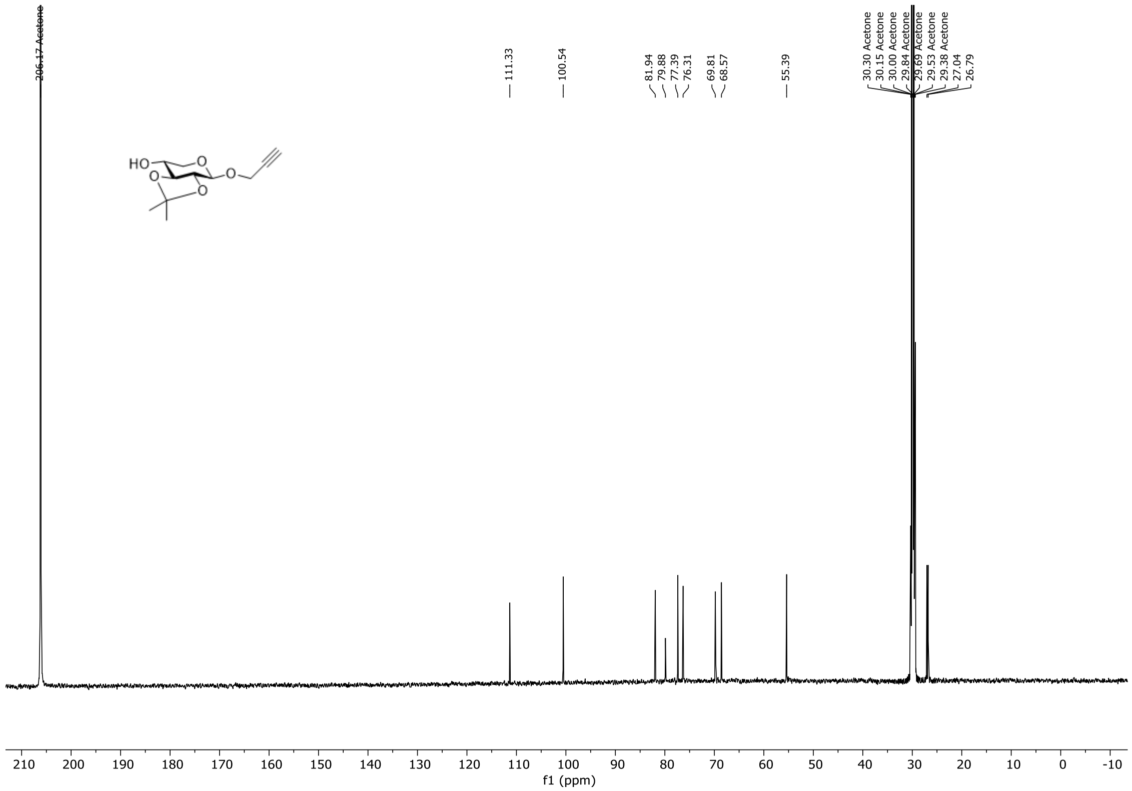

**Figure S45.** $^{13}C$ NMR spectrum of compound **S11** (126 MHz, acetone-$d_6$, 298 K).

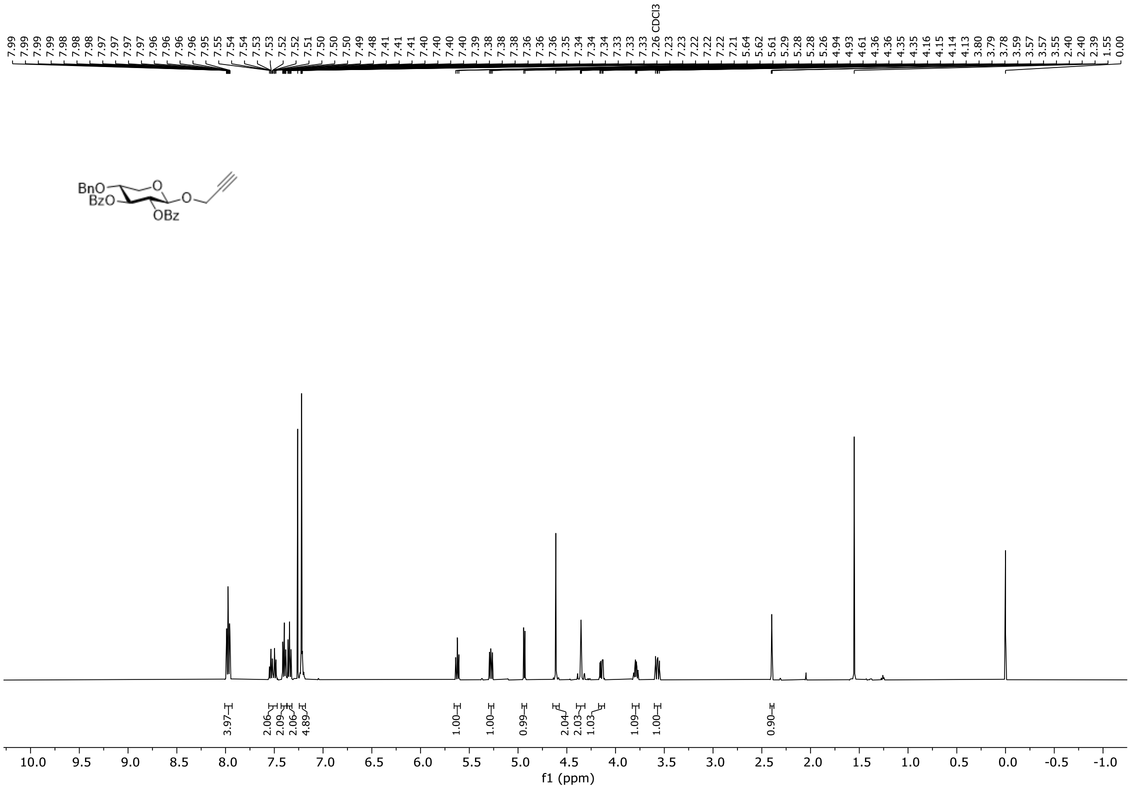

**Figure S46.** $^{1}$H NMR spectrum of compound **S12** (500 MHz, $CDCl_3$, 298 K).

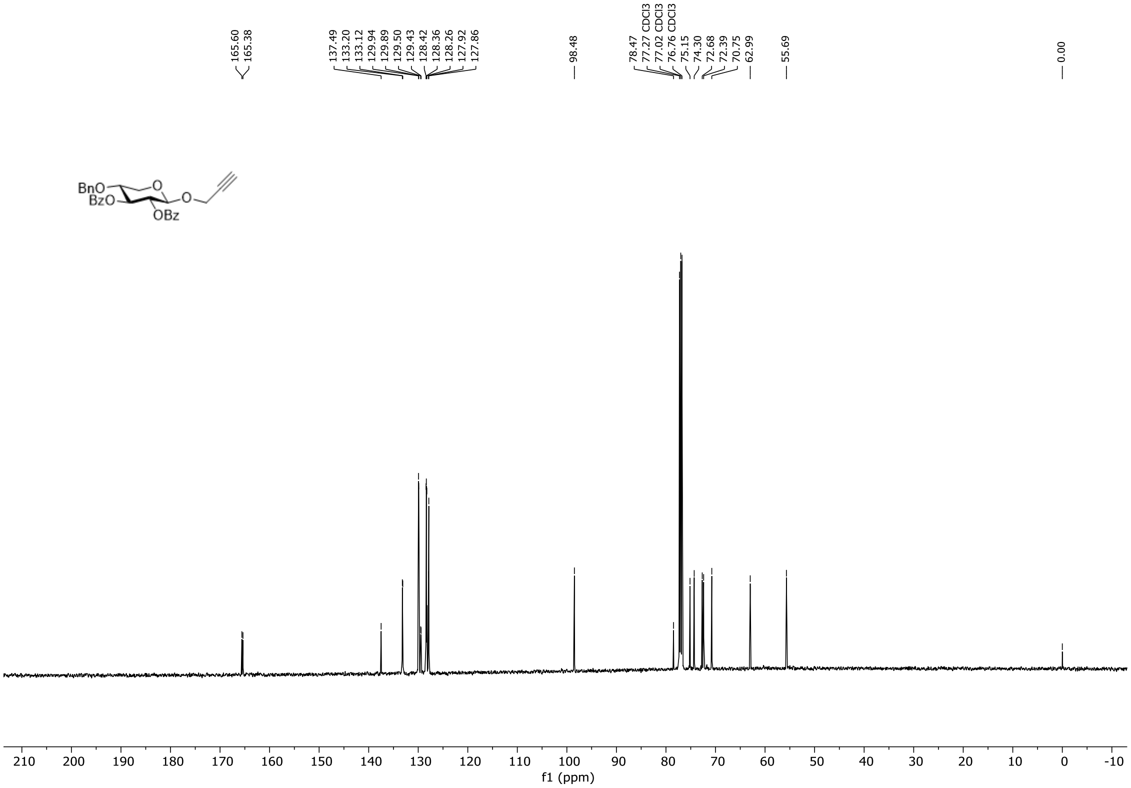

**Figure S47.** $^{13}$C NMR spectrum of compound **S12** (126 MHz, $CDCl_3$, 298 K).

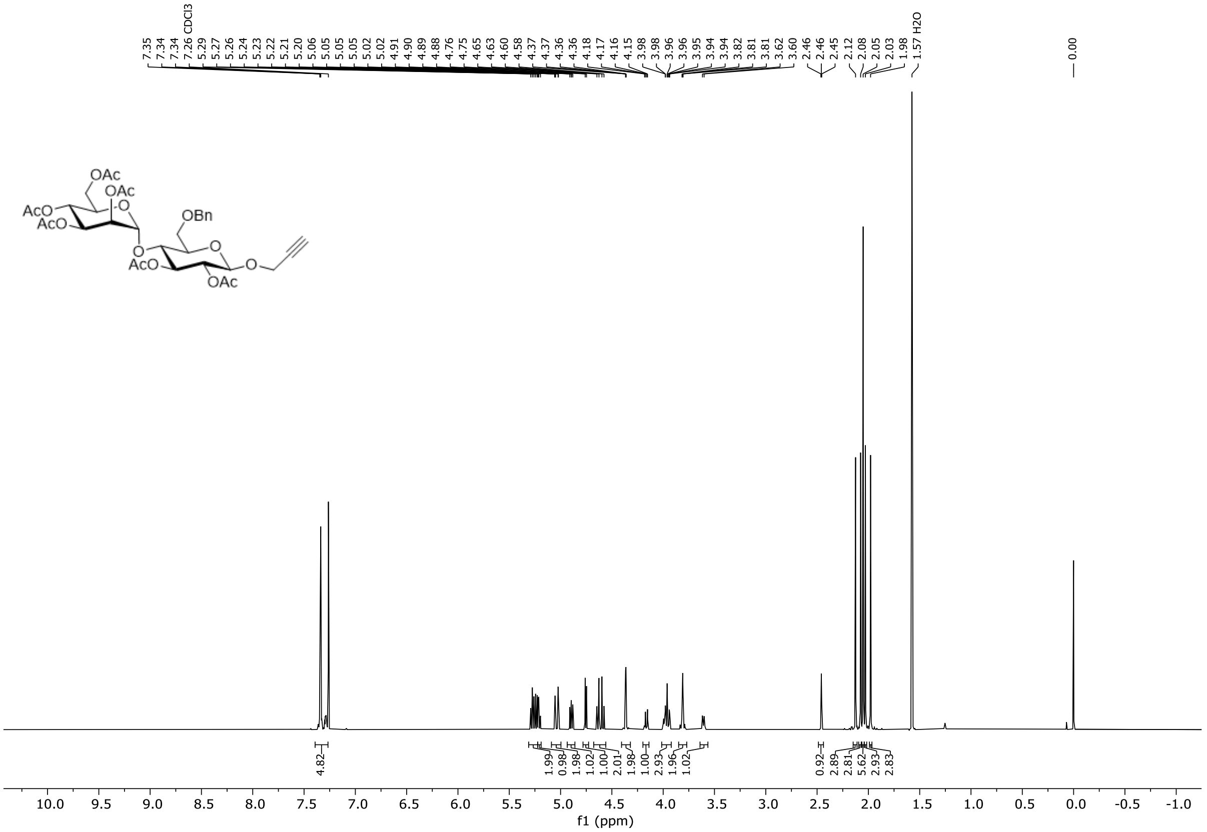

**Figure S48.** $^{1}H$ NMR spectrum of compound **S13** (600 MHz, $CDCl_3$, 298 K).

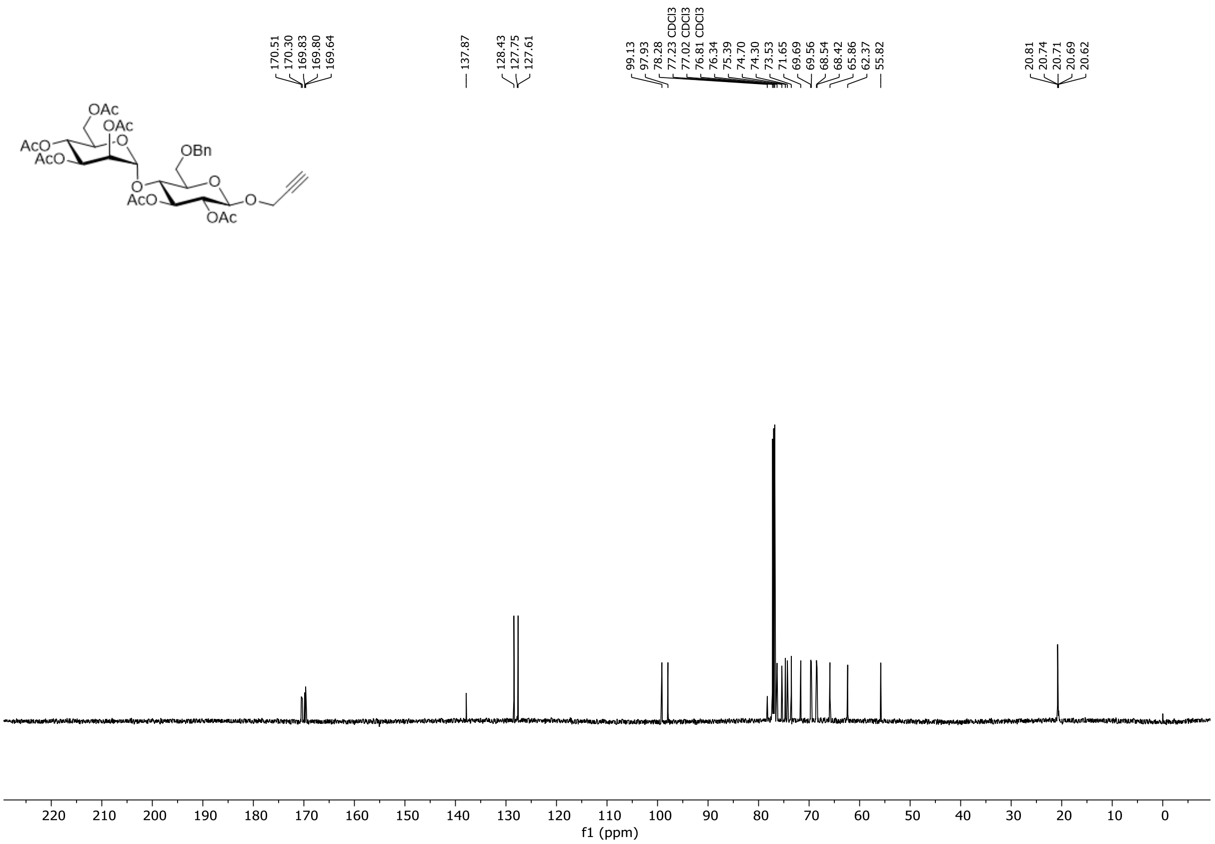

**Figure S49.** $^{13}C$ NMR spectrum of compound **S13** (151 MHz, $CDCl_3$, 298 K).

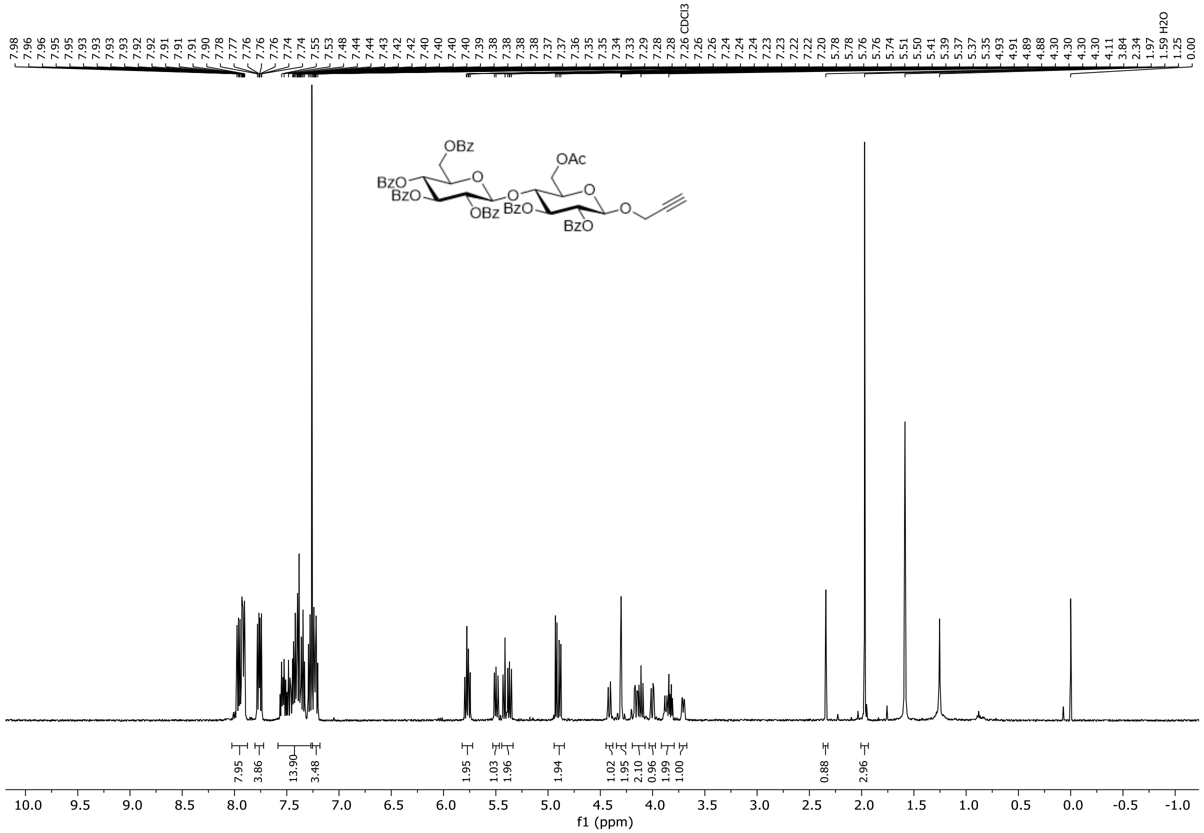

**Figure S50.** $^{1}H$ NMR spectrum of compound **S14** (600 MHz, $CDCl_3$, 298 K).

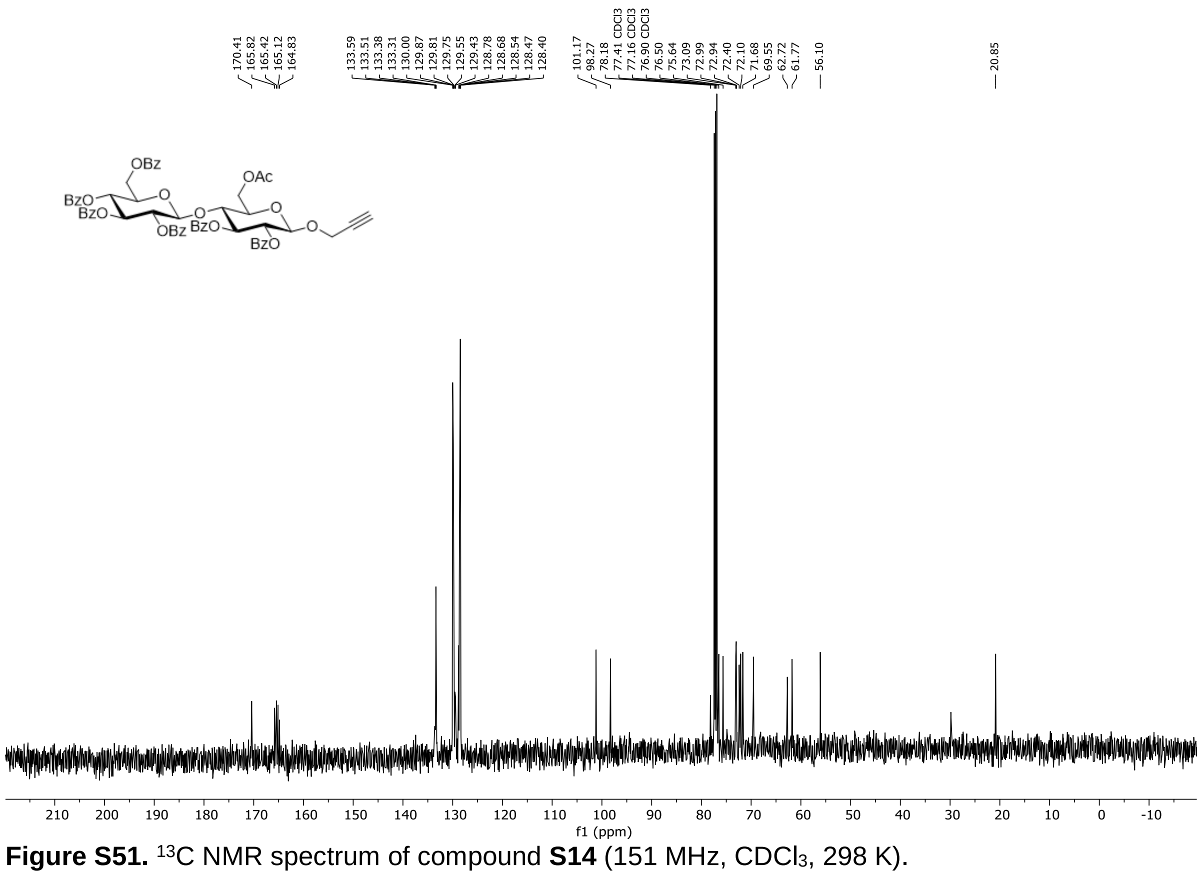

**Figure S51.** $^{13}C$ NMR spectrum of compound **S14** (151 MHz, $CDCl_3$, 298 K).

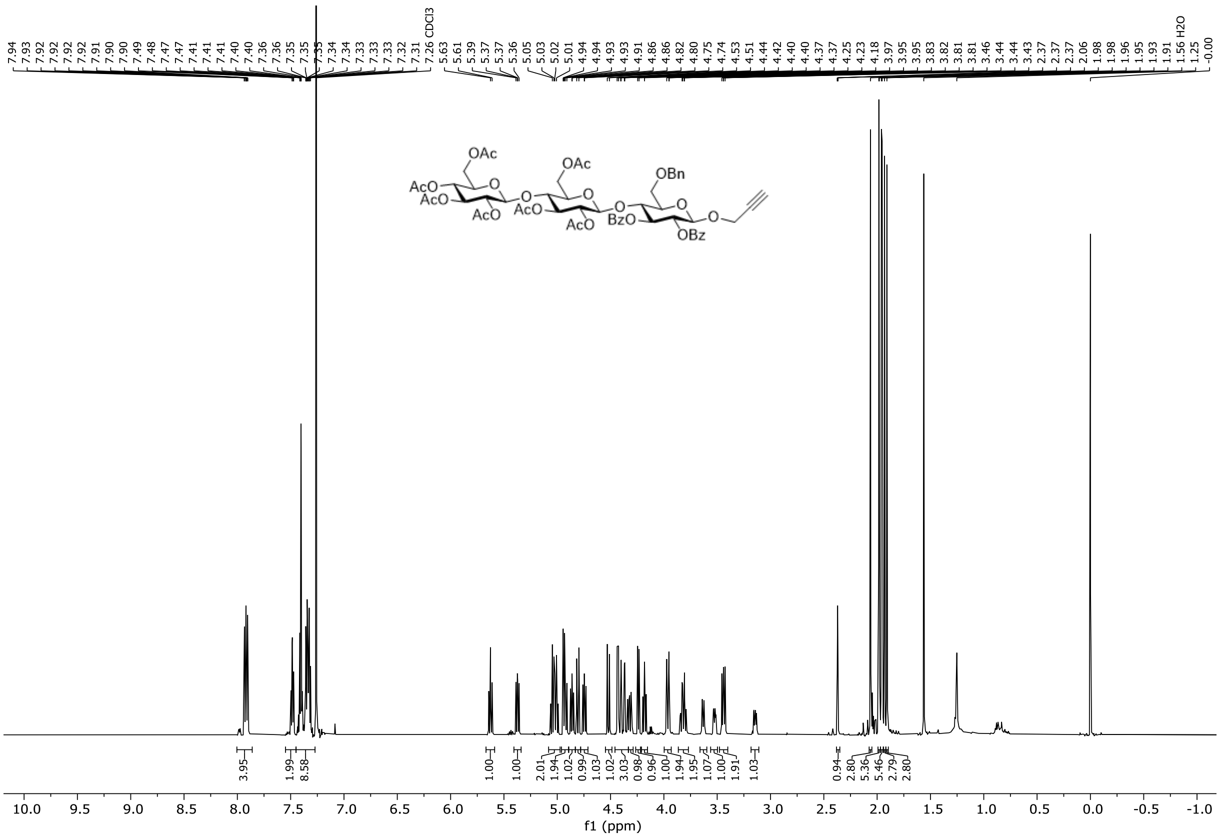

**Figure S52.** $^{1}$H NMR spectrum of compound **S15** (600 MHz, $CDCl_3$, 298 K).

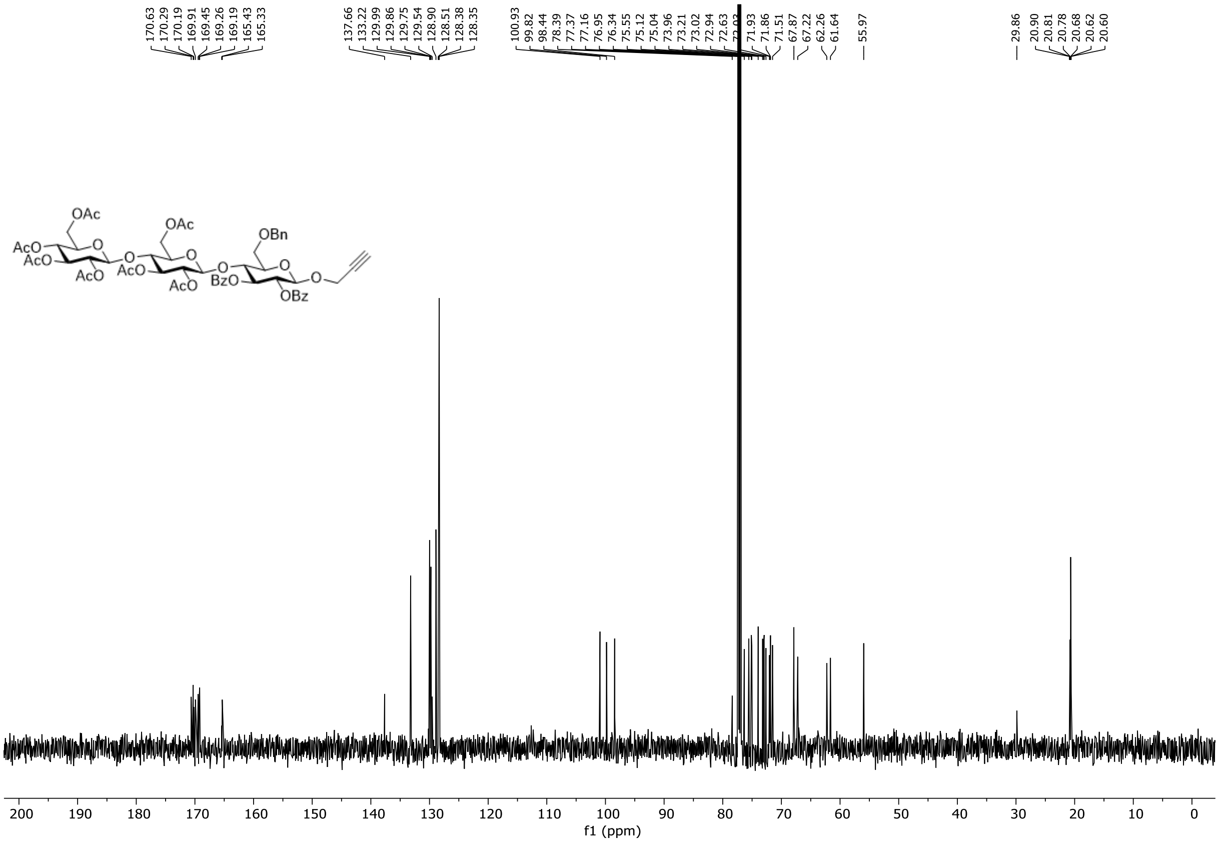

**Figure S53.** $^{13}$C NMR spectrum of compound **S15** (151 MHz, $CDCl_3$, 298 K).

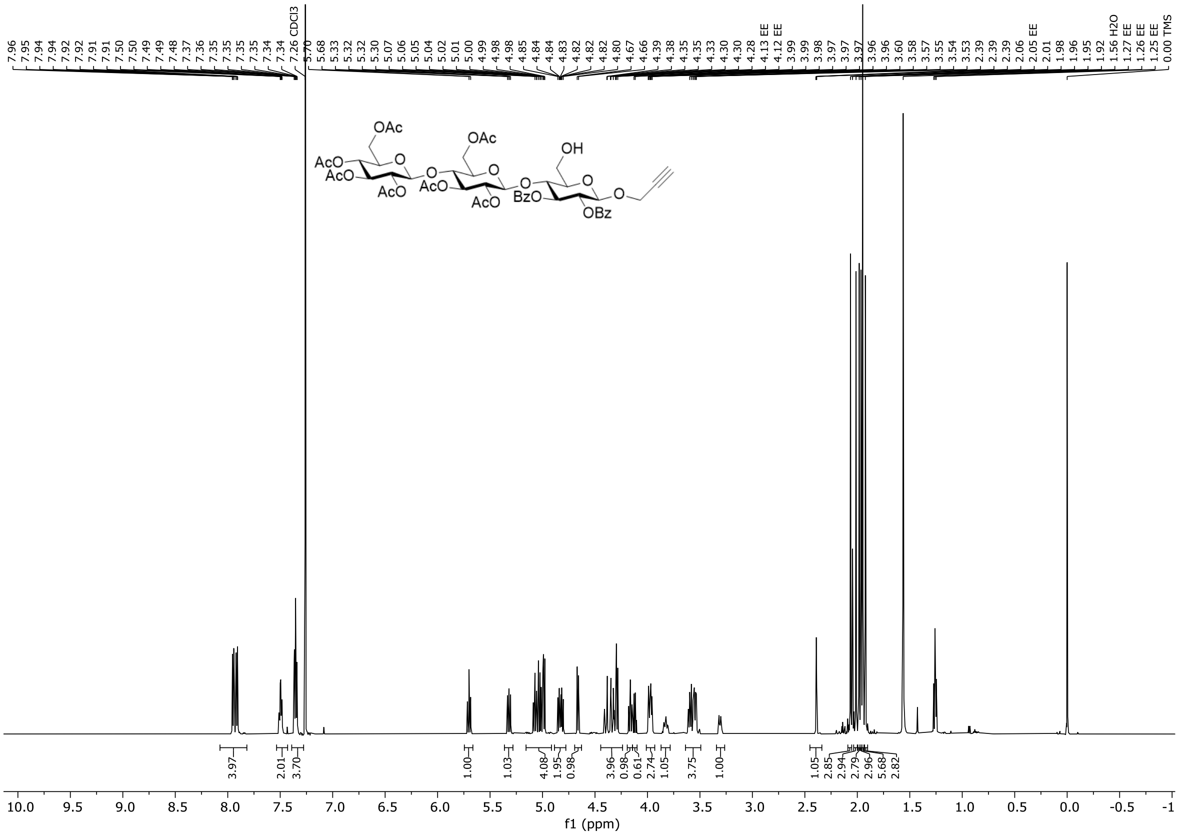

**Figure S54.** $^1$H NMR spectrum of compound **S16** (600 MHz, $CDCl_3$, 298 K).

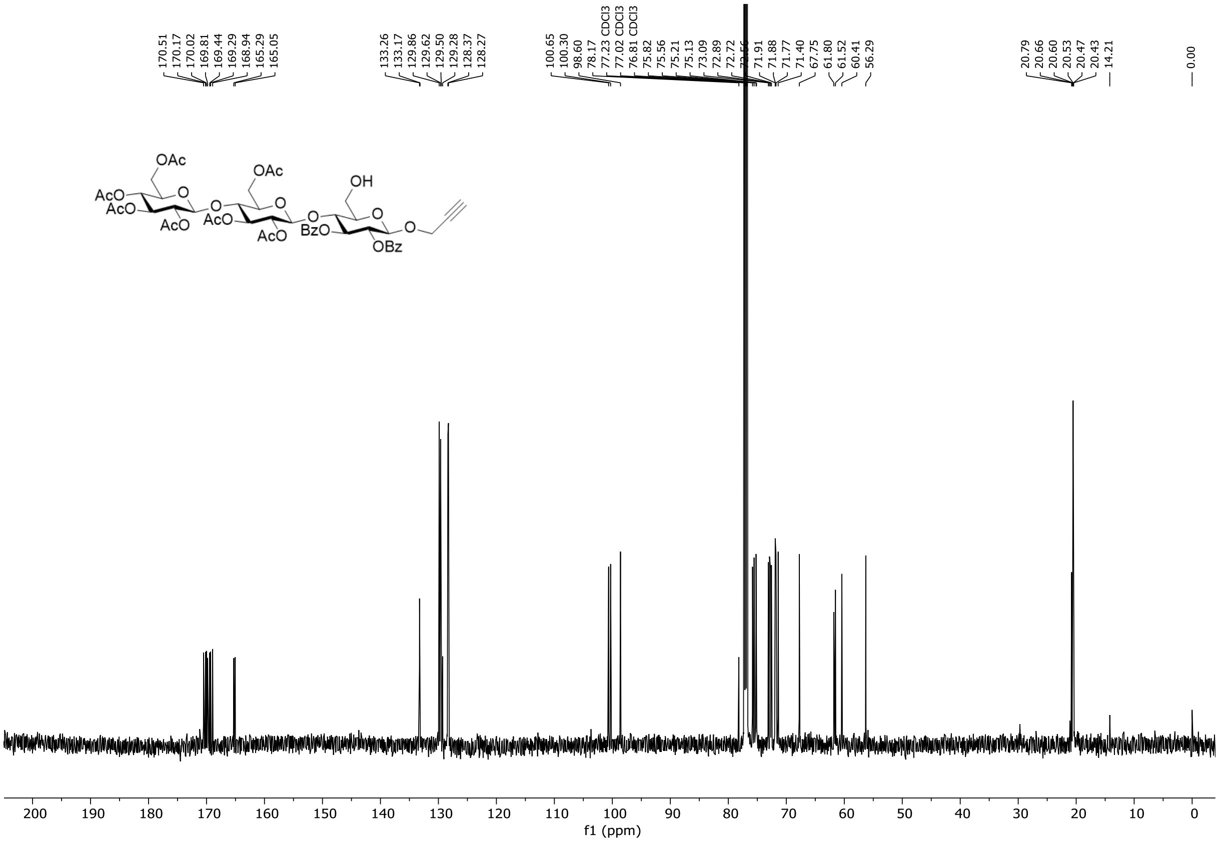

**Figure S55.** $^{13}$C NMR spectrum of compound **S16** (151 MHz, $CDCl_3$, 298 K).

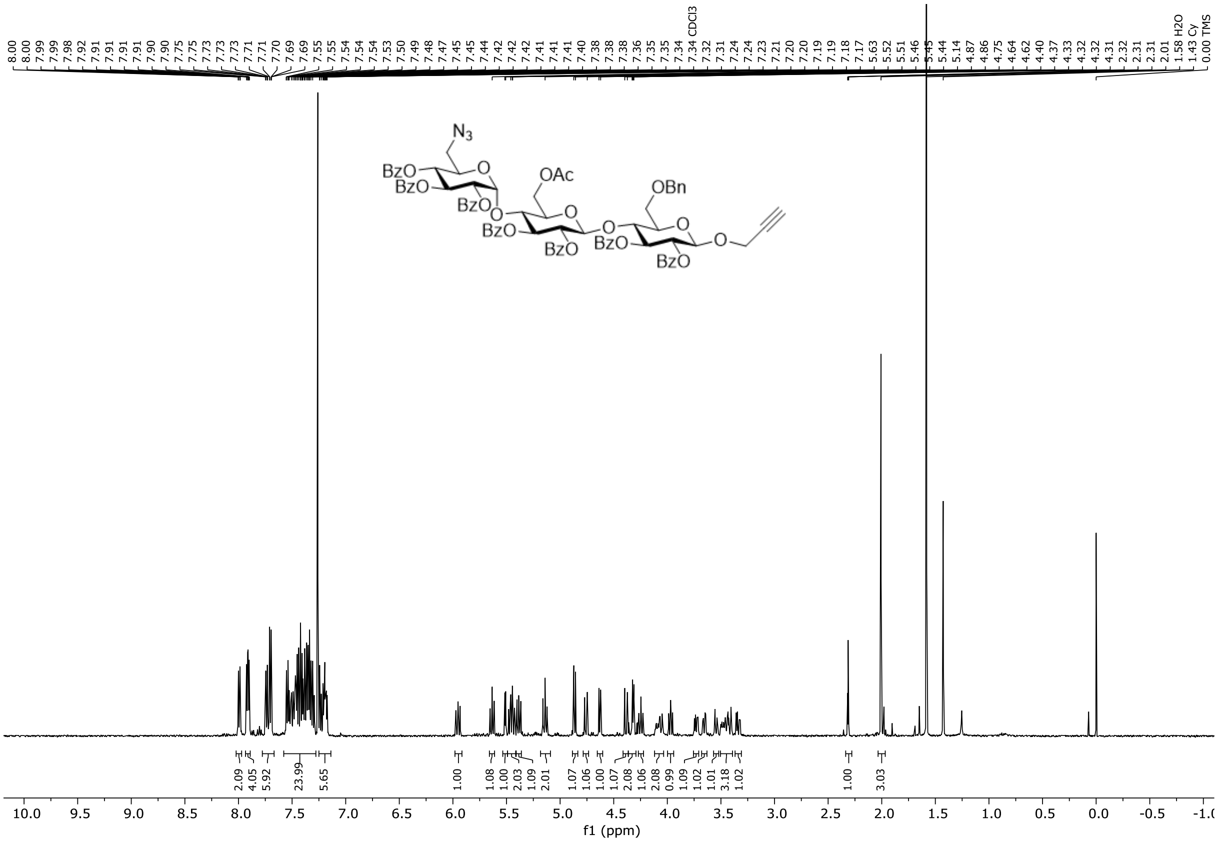

**Figure S56.** $^{1}$H NMR spectrum of compound **S17** (500 MHz, $CDCl_3$, 298 K).

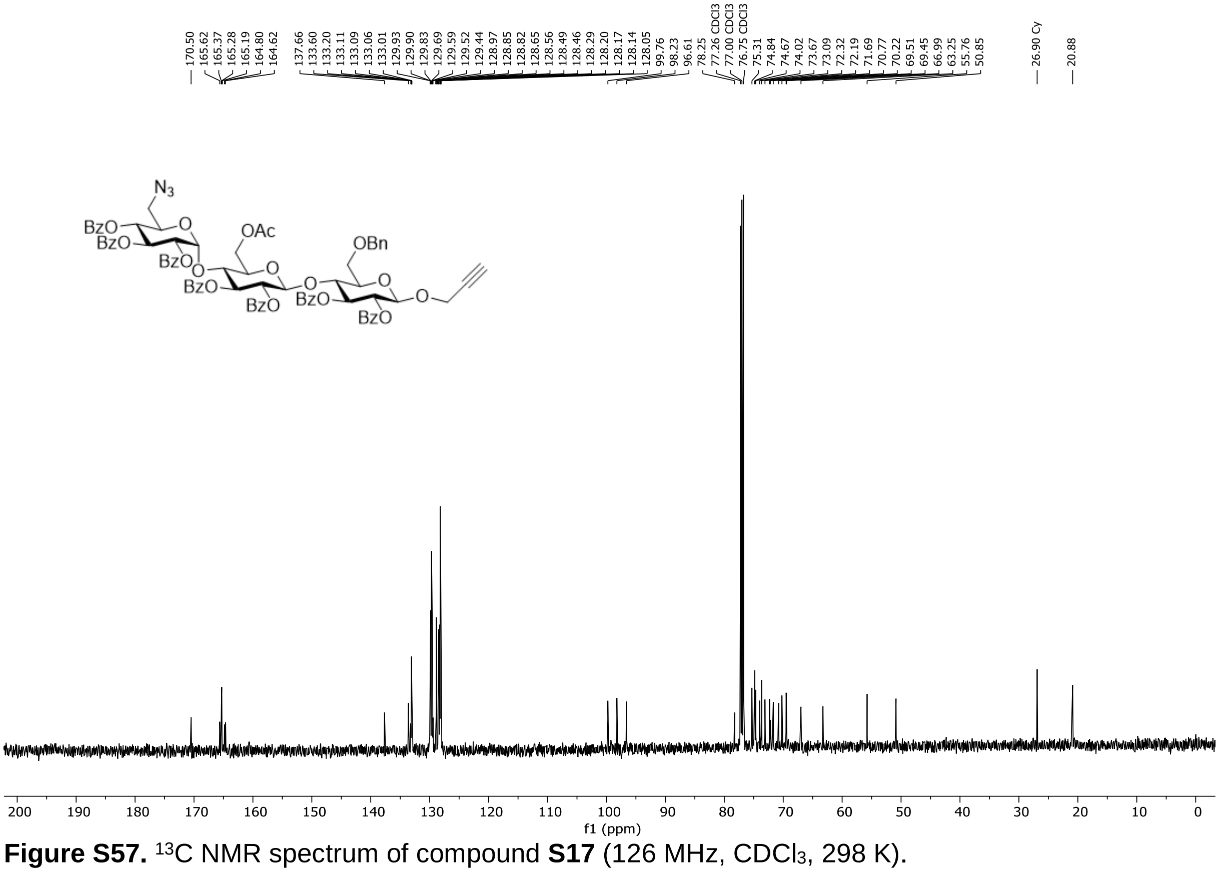

**Figure S57.** $^{13}$C NMR spectrum of compound **S17** (126 MHz, $CDCl_3$, 298 K).

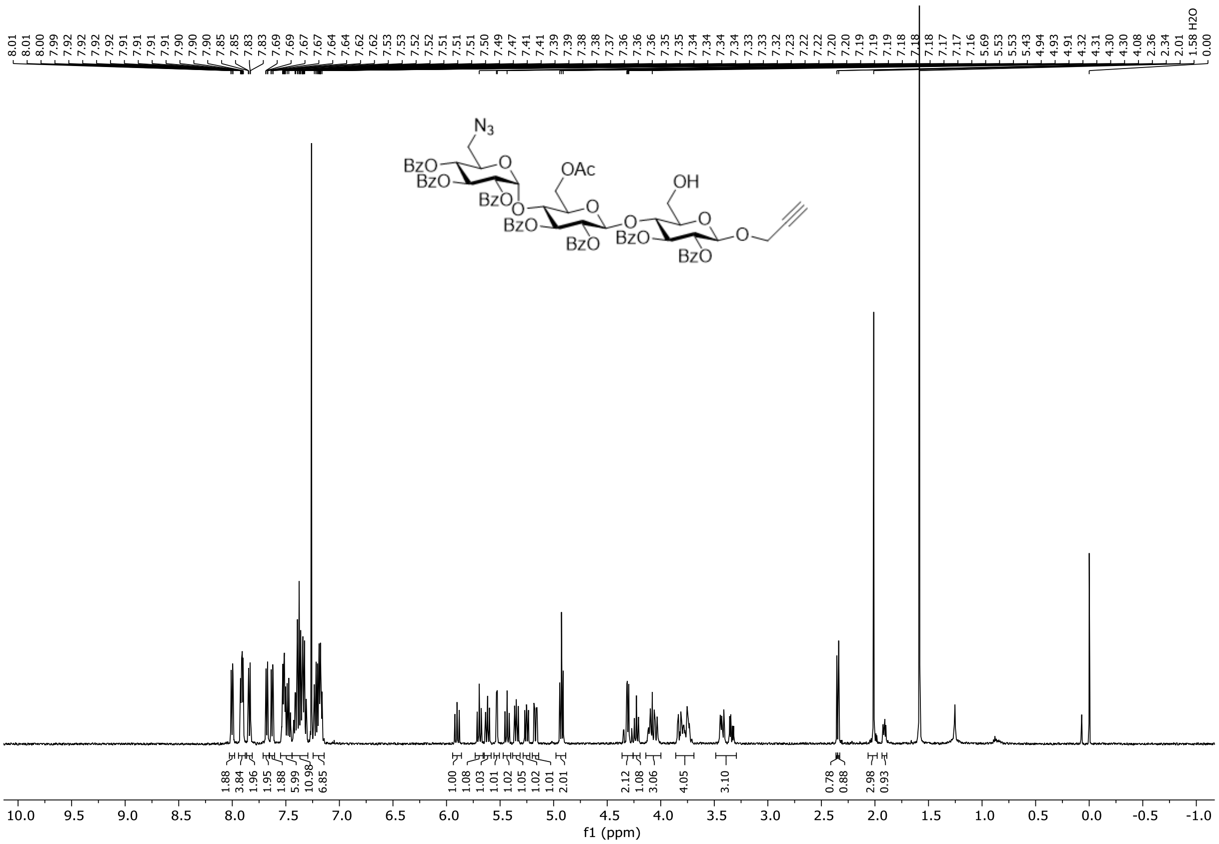

**Figure S58.** $^{1}H$ NMR spectrum of compound **S18** (500 MHz, $CDCl_3$, 298 K).

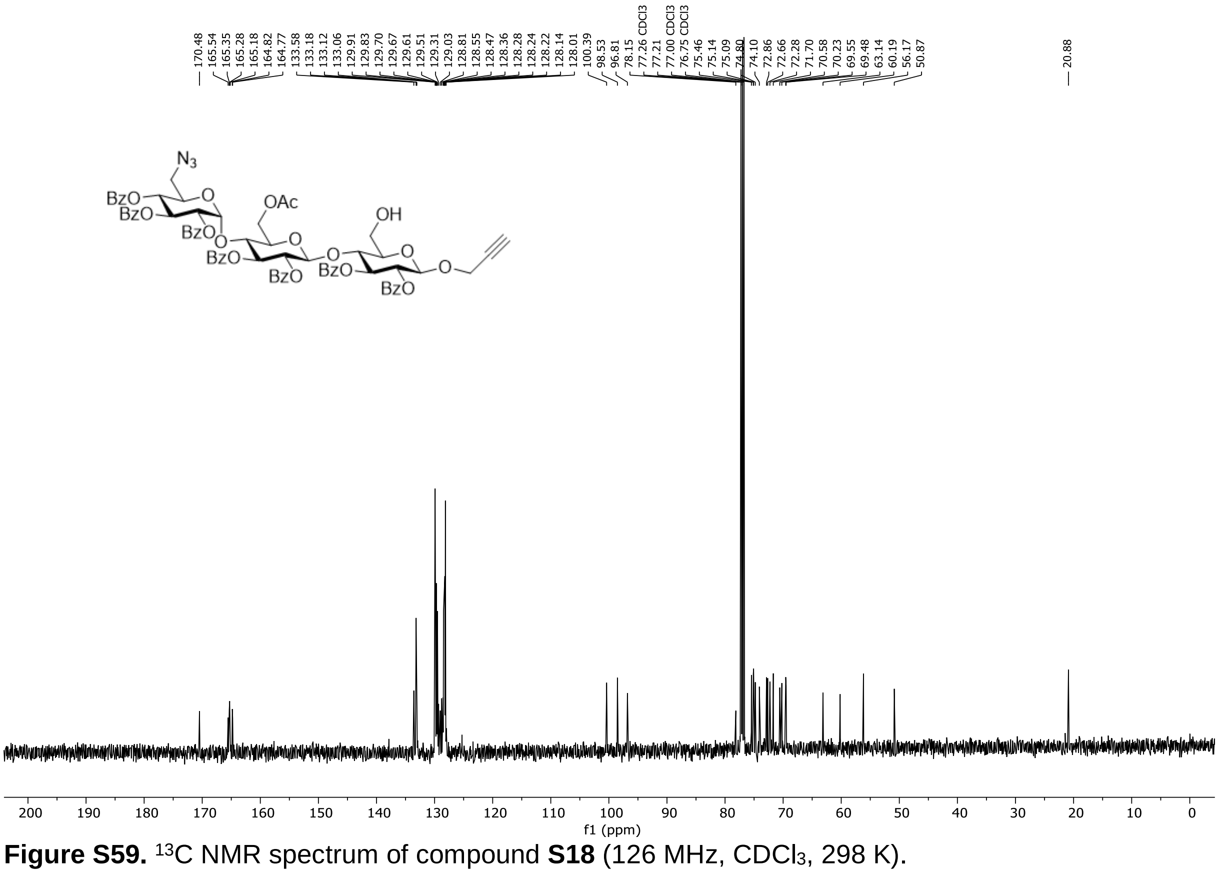

**Figure S59.** $^{13}C$ NMR spectrum of compound **S18** (126 MHz, $CDCl_3$, 298 K).

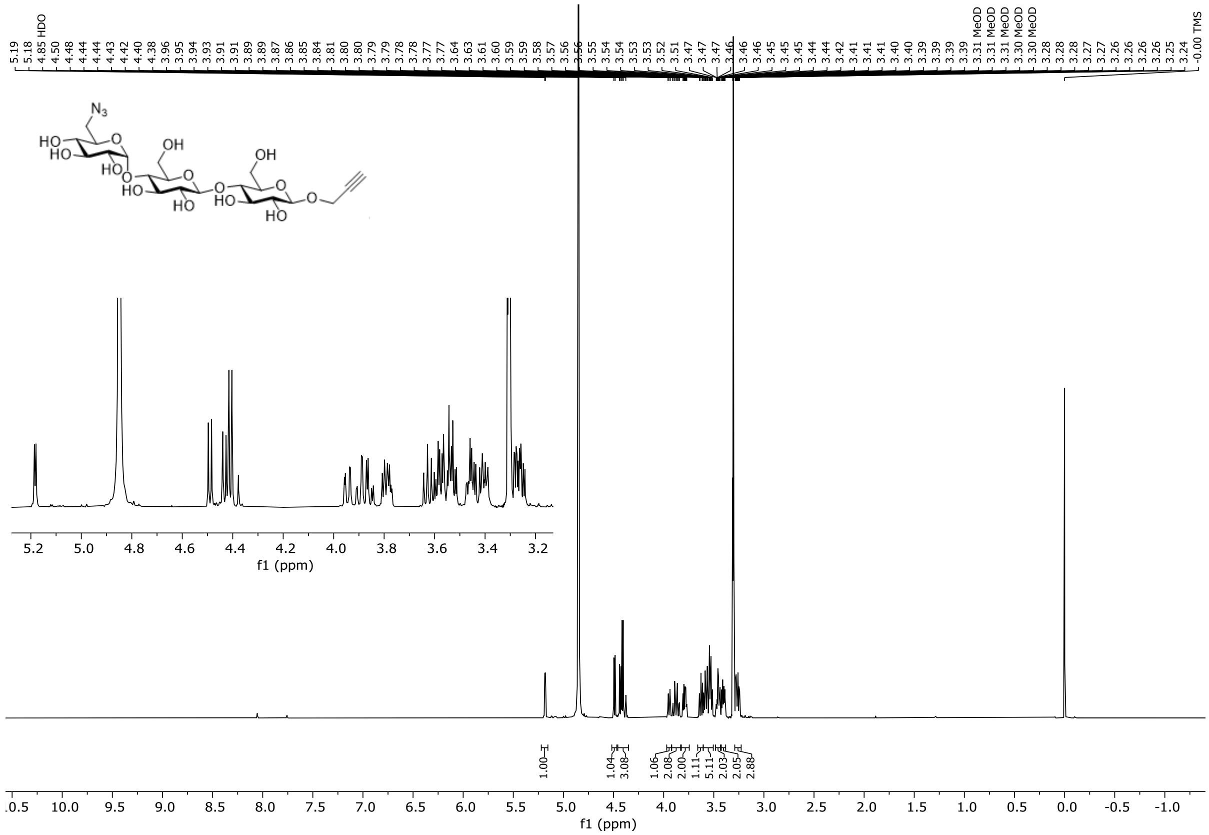

**Figure S60.** $^{1}H$ NMR spectrum of compound **S19** (600 MHz, methanol-$d_4$, 298 K).

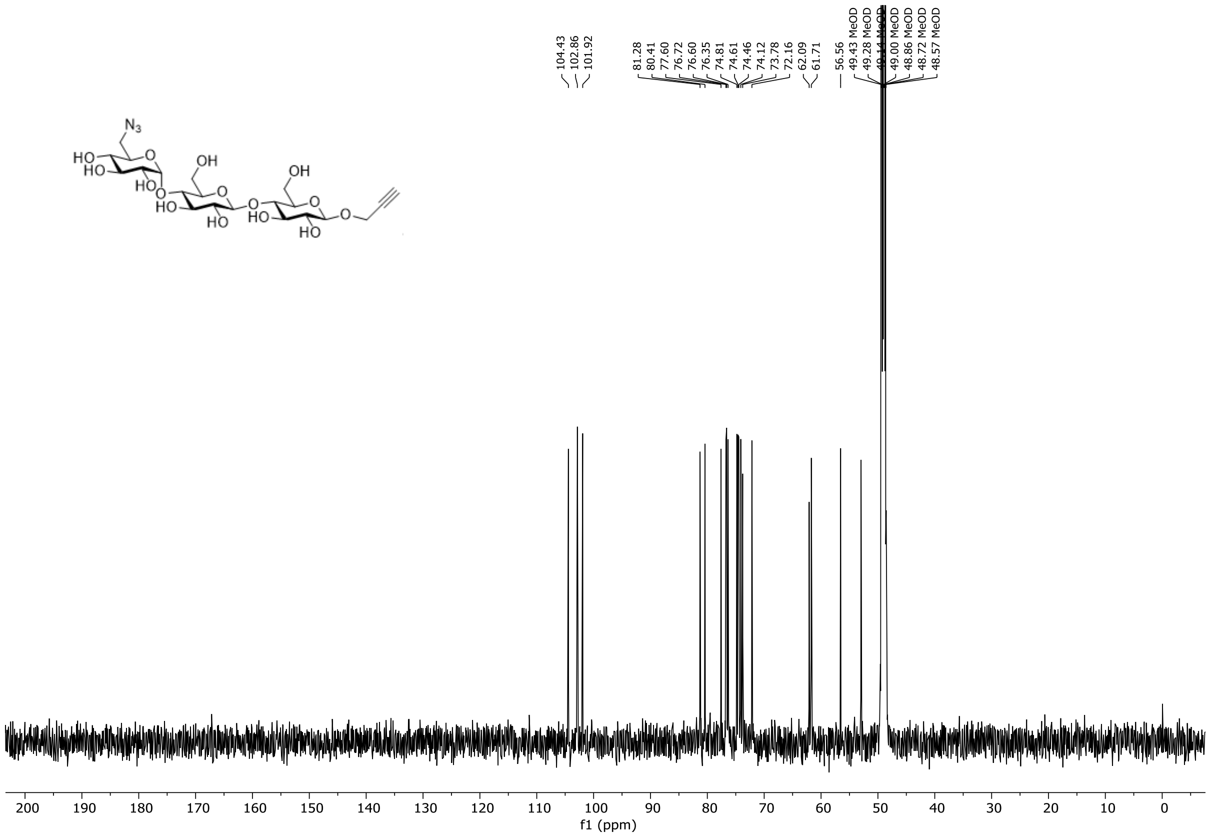

**Figure S61.** $^{13}C$ NMR spectrum of compound **S19** (151 MHz, methanol-$d_4$, 298 K).

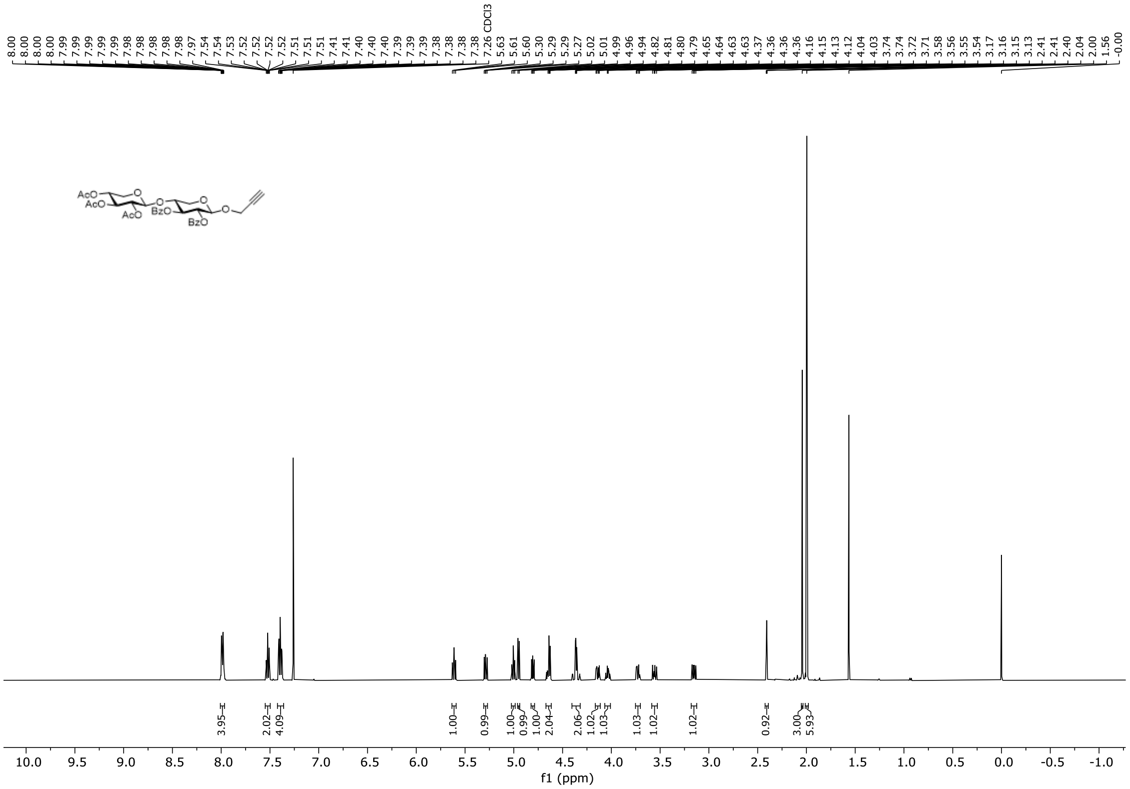

**Figure S62.** $^{1}H$ NMR spectrum of compound **S20** (500 MHz, $CDCl_3$, 298 K).

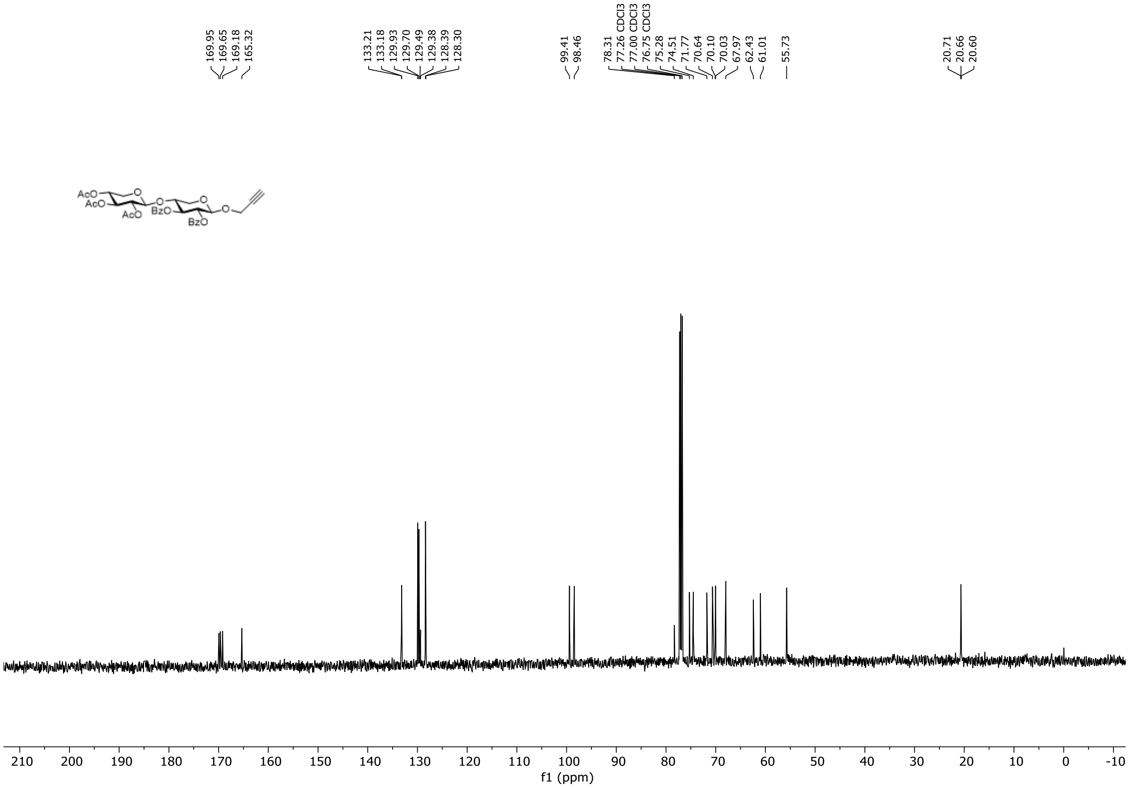

**Figure S63.** $^{13}C$ NMR spectrum of compound **S20** (126 MHz, $CDCl_3$, 298 K).

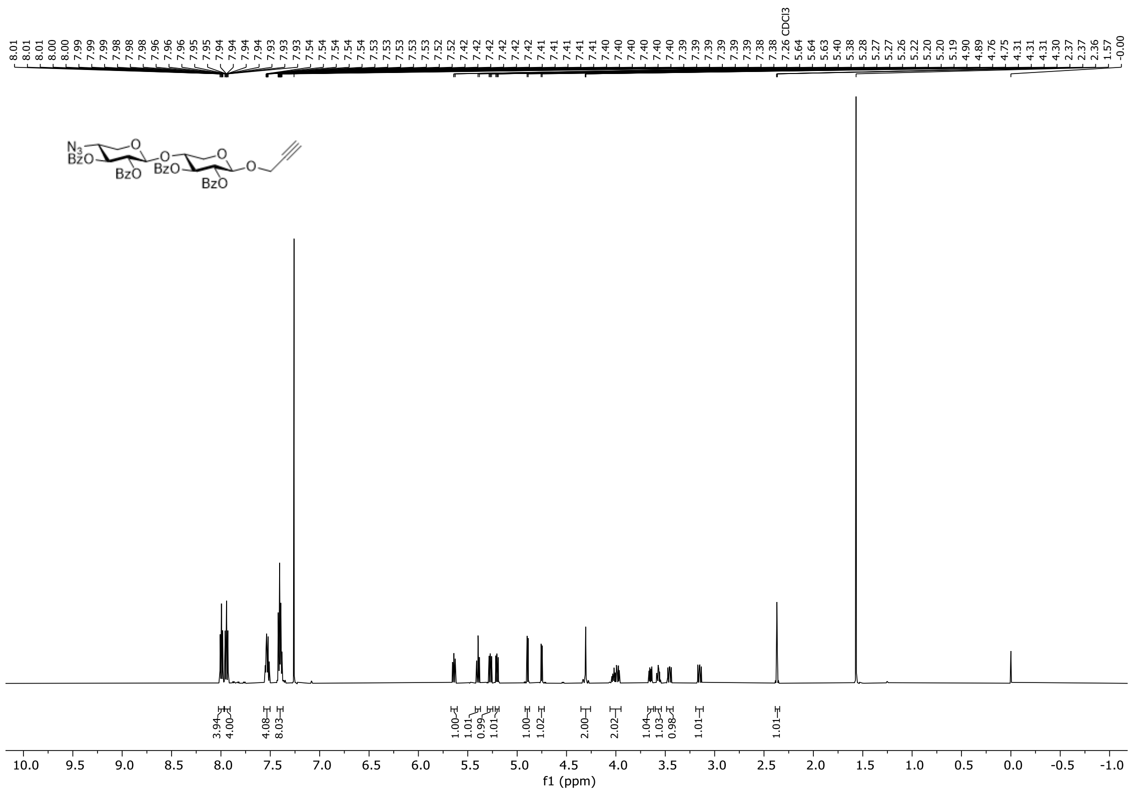

**Figure S64.** $^{1}$H NMR spectrum of compound **S21** (600 MHz, $CDCl_3$, 298 K).

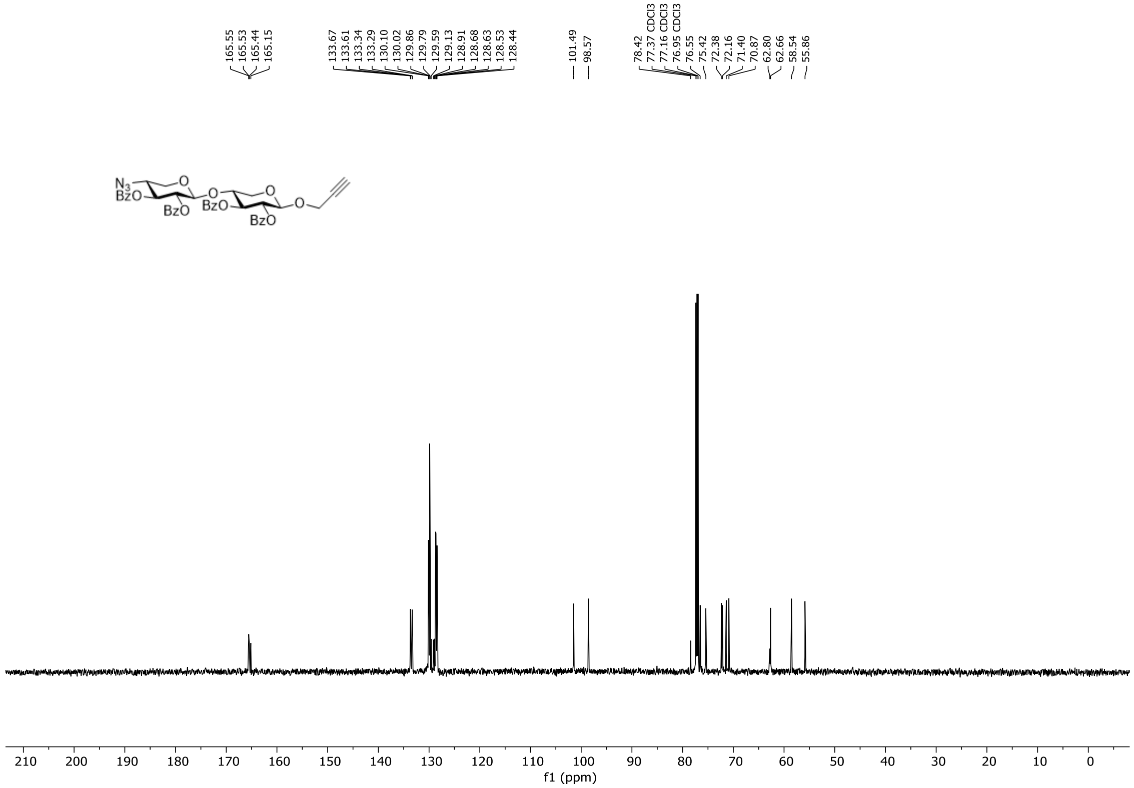

**Figure S65.** $^{13}$C NMR spectrum of compound **S21** (151 MHz, $CDCl_3$, 298 K).

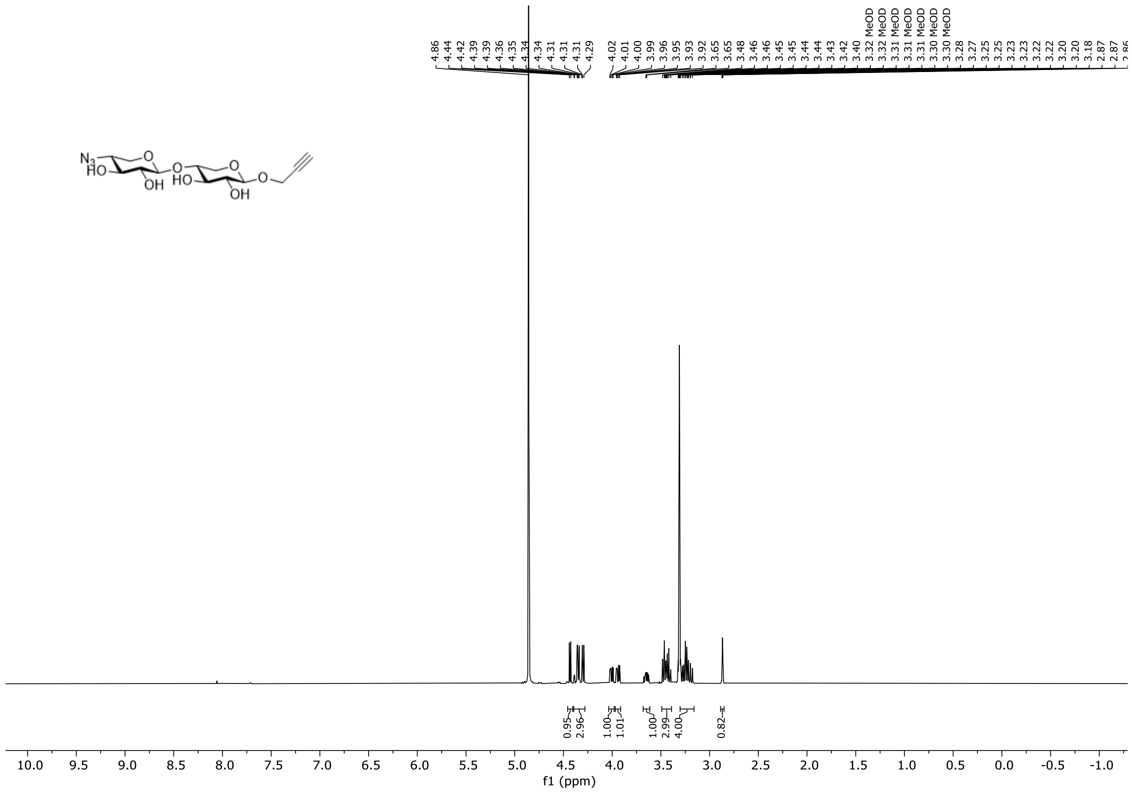

**Figure S66.** $^{1}$H NMR spectrum of compound **S22** (500 MHz, methanol-$d_4$, 298 K).

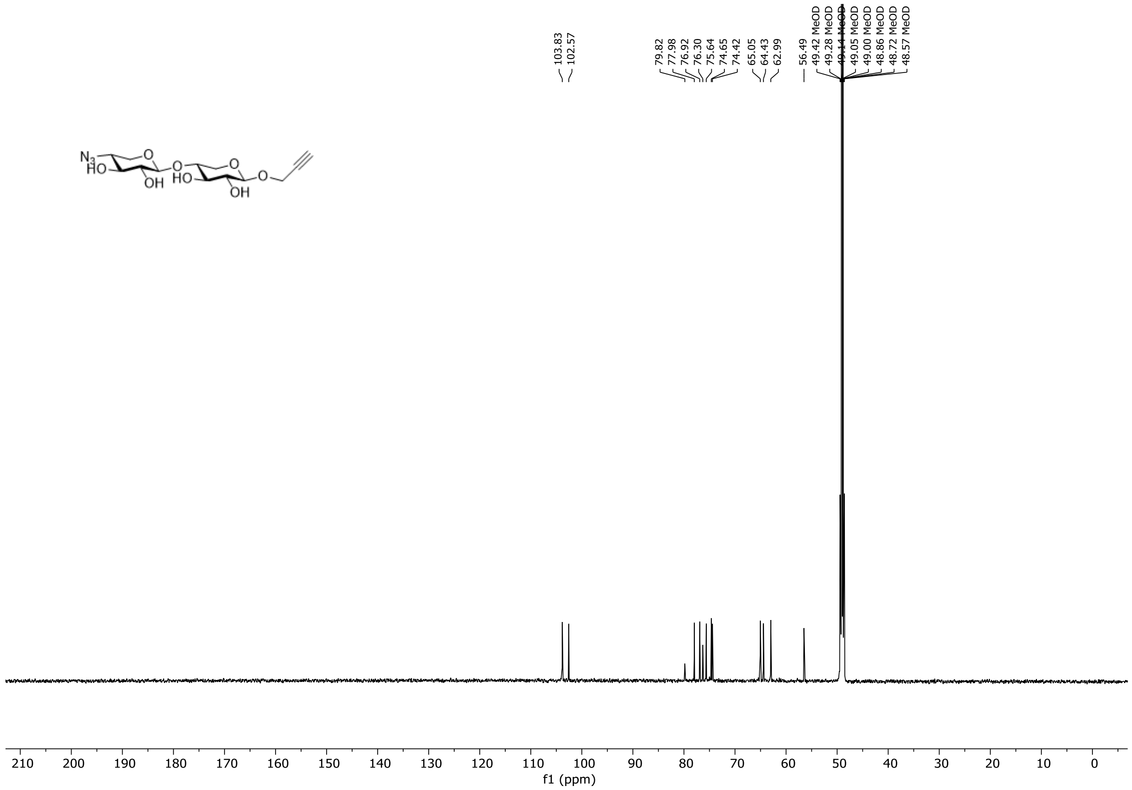

**Figure S67.** $^{13}$C NMR spectrum of compound **S22** (151 MHz, methanol-$d_4$, 298 K).

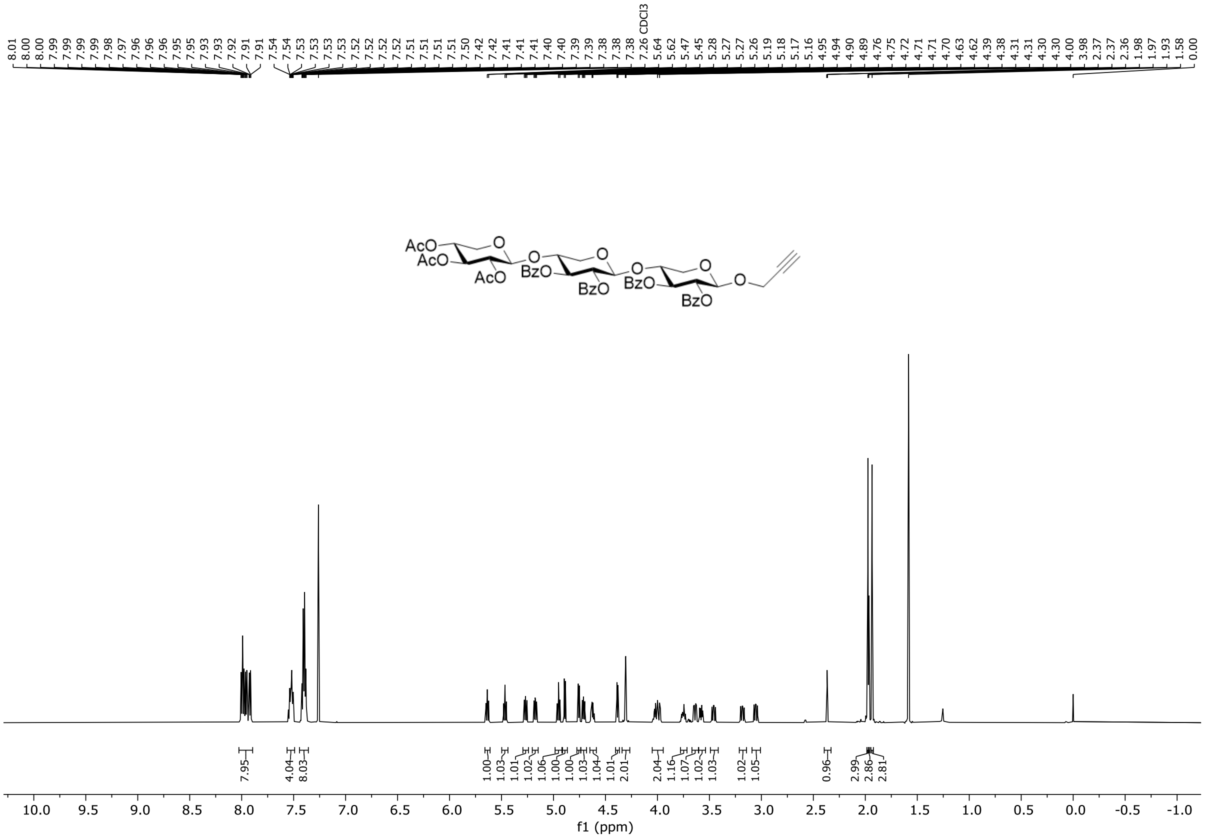

**Figure S68.** $^{1}$H NMR spectrum of compound **S23** (600 MHz, $CDCl_3$, 298 K).

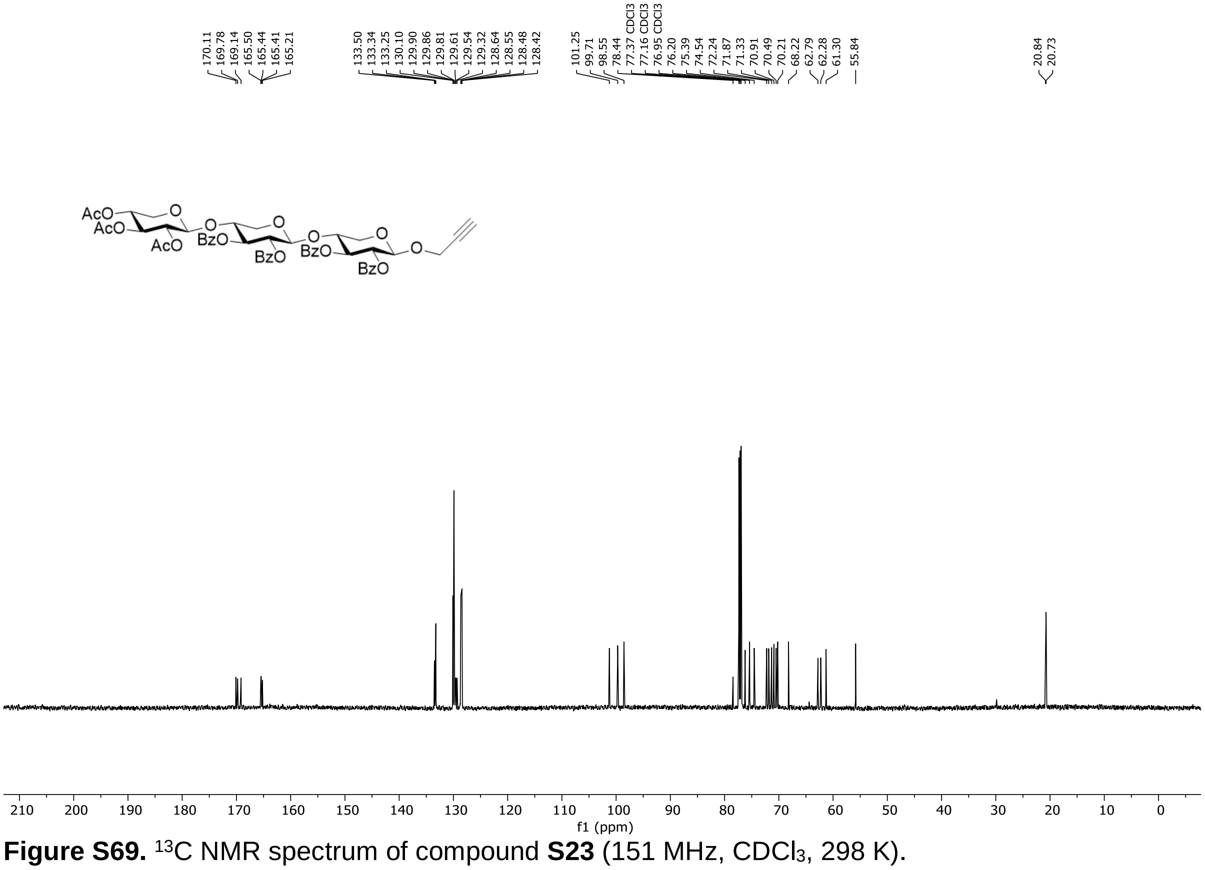

**Figure S69.** $^{13}$C NMR spectrum of compound **S23** (151 MHz, $CDCl_3$, 298 K).

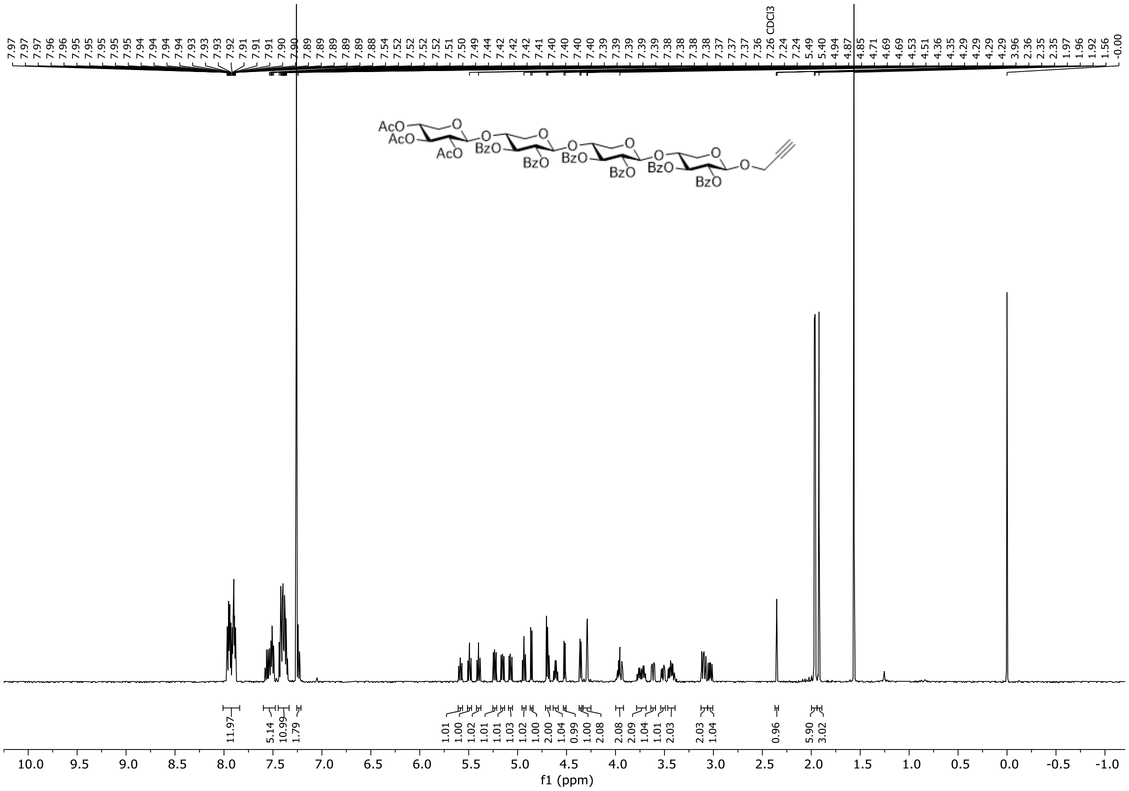

**Figure S70.** $^{1}$H NMR spectrum of compound **S24** (500 MHz, $CDCl_3$, 298 K).

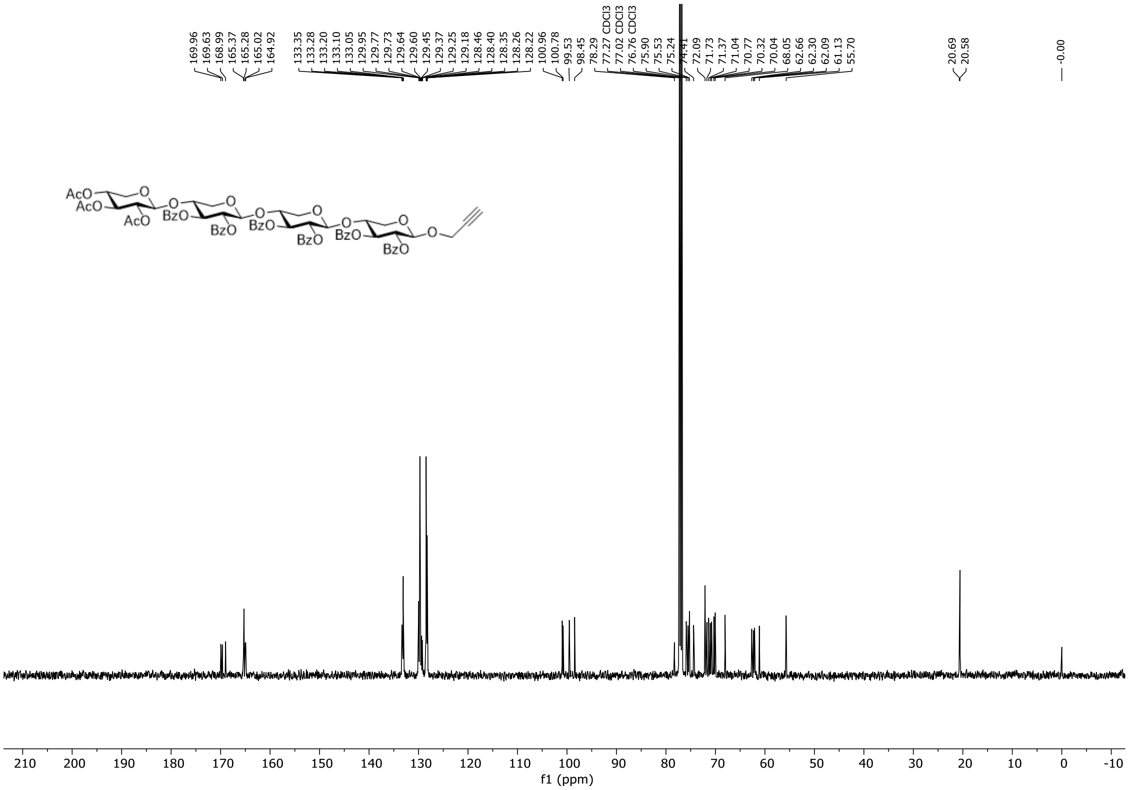

**Figure S71.** $^{13}$C NMR spectrum of compound **S24** (126 MHz, $CDCl_3$, 298 K).

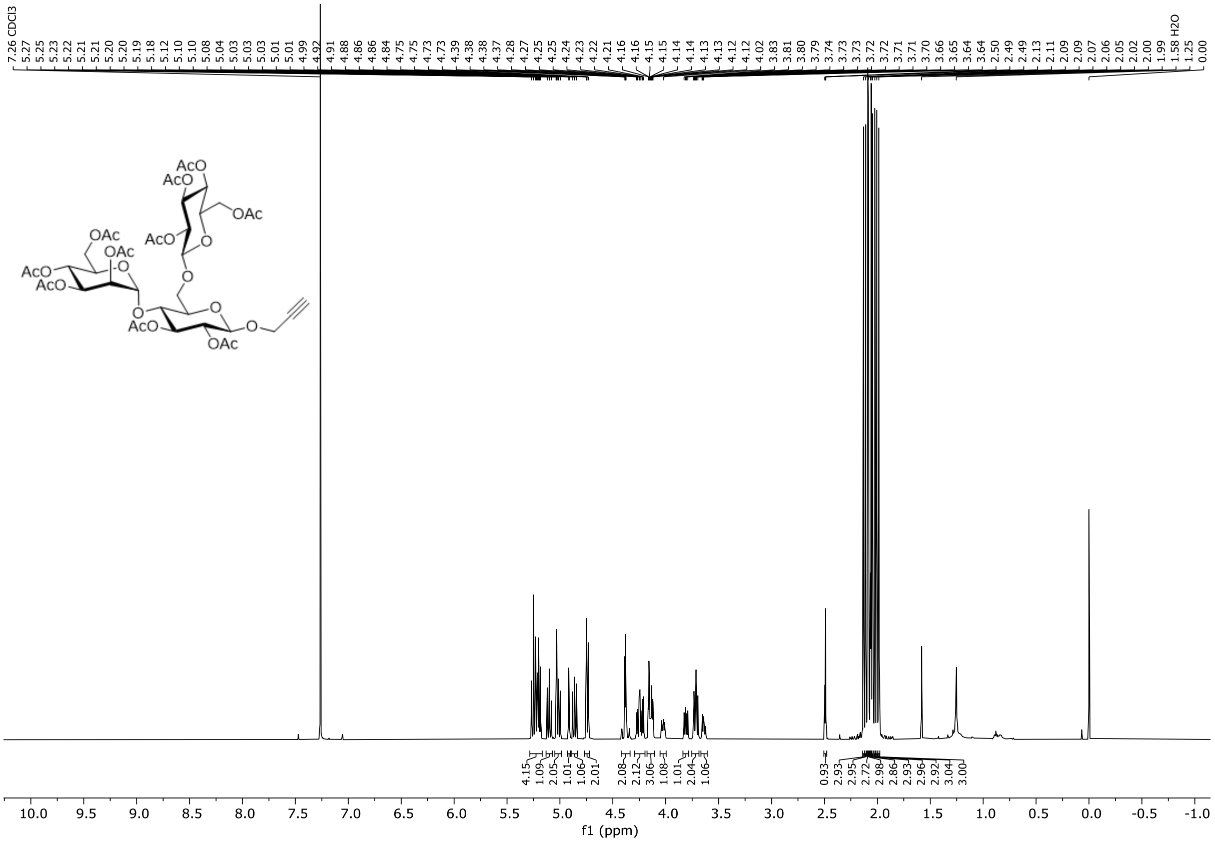

**Figure S72.** $^{1}$H NMR spectrum of compound **S25** (500 MHz, $CDCl_3$, 298 K).

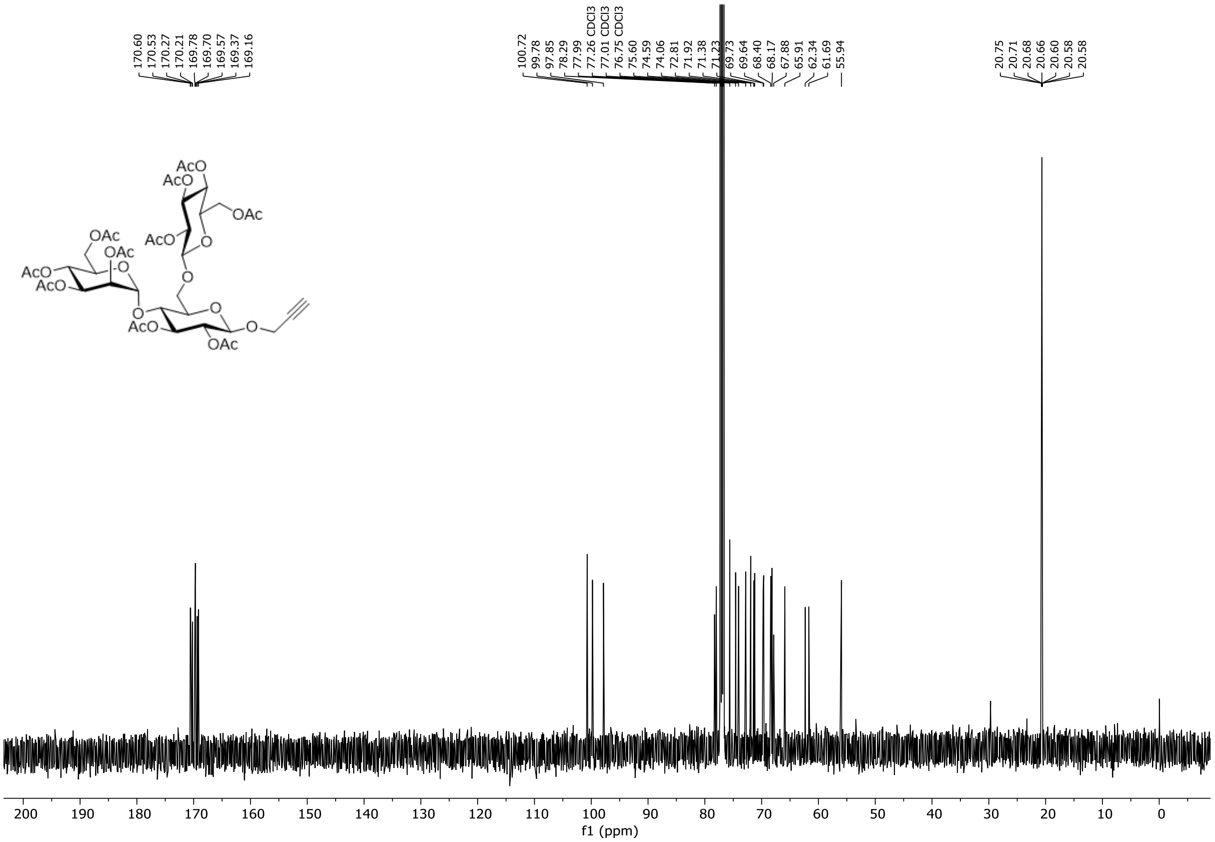

**Figure S73.** $^{13}$C NMR spectrum of compound **S25** (126 MHz, $CDCl_3$, 298 K).

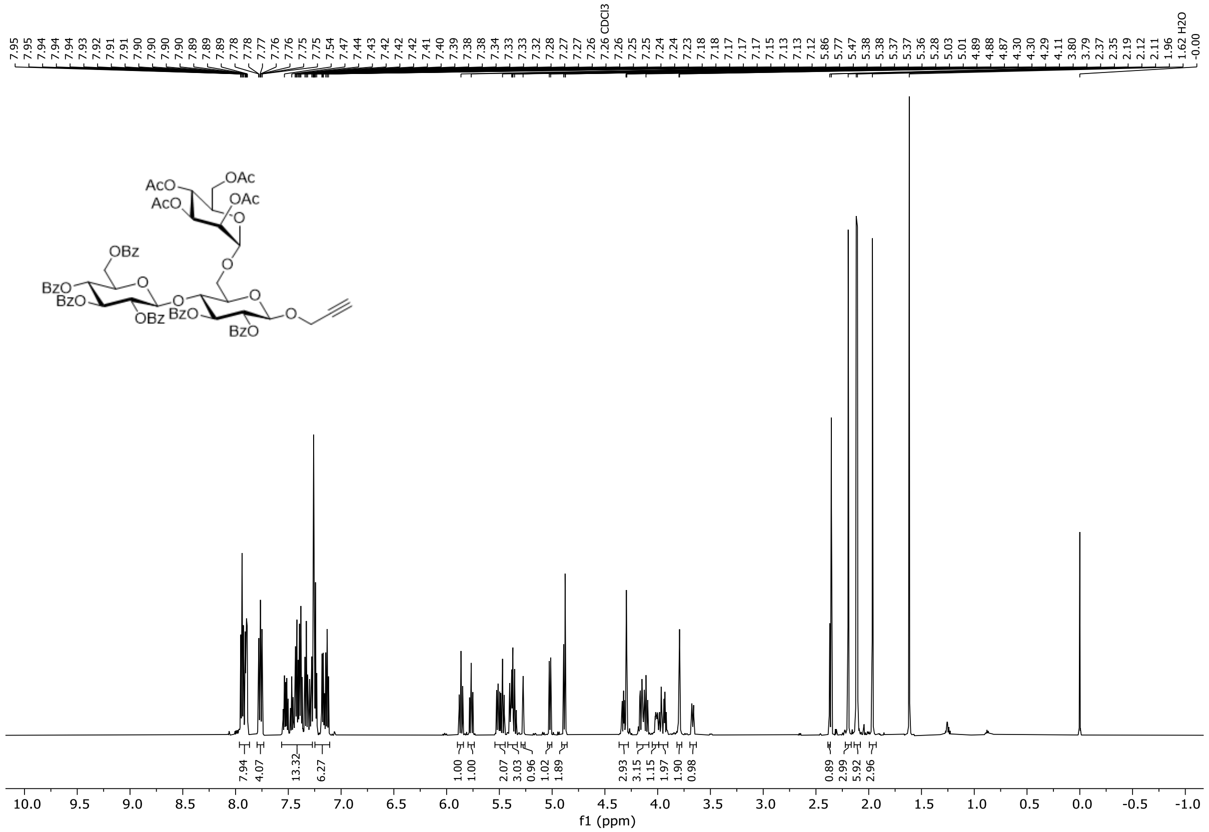

**Figure S74.** $^{1}H$ NMR spectrum of compound **S26** (600 MHz, $CDCl_3$, 298 K).

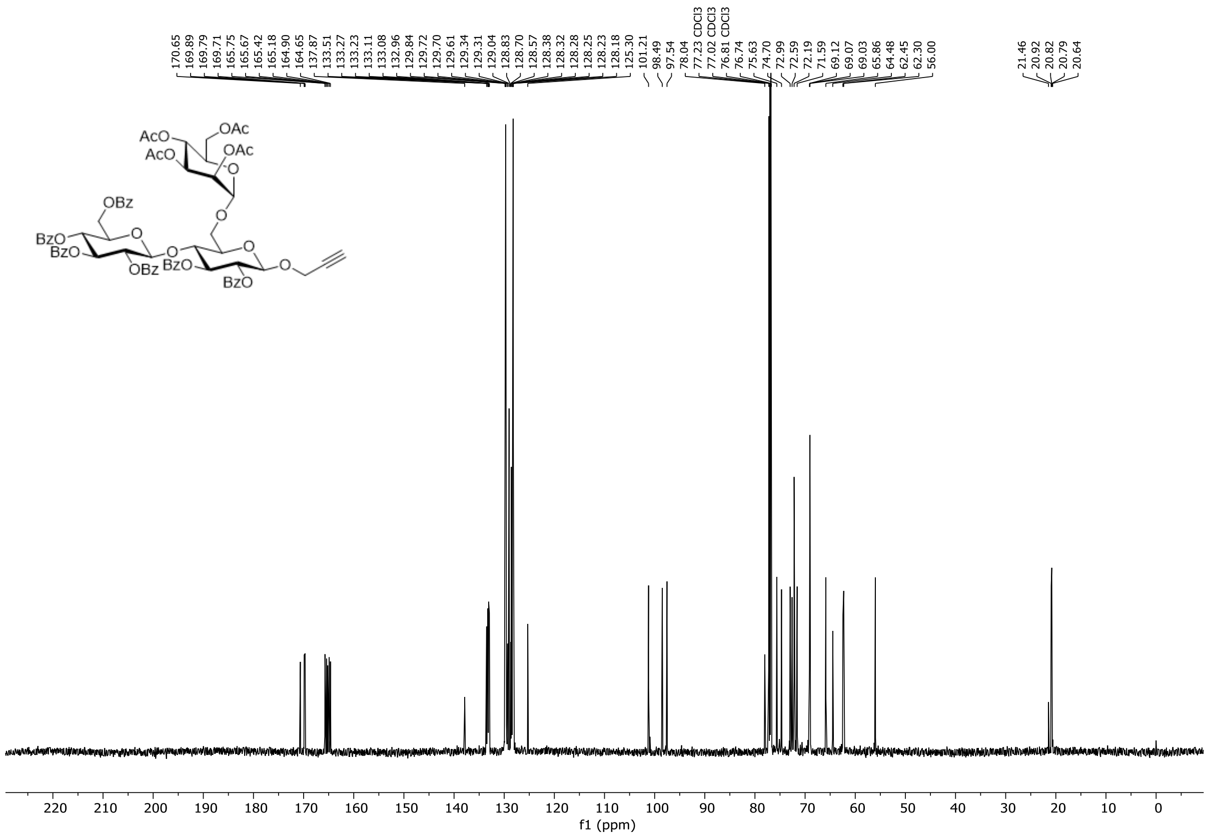

**Figure S75.** $^{13}C$ NMR spectrum of compound **S26** (151 MHz, $CDCl_3$, 298 K).

#### 4. Preparation of Supported Lipid Bilayers (SLBs)

The coverslips (VWR, Ø25mm, thickness No. 1.5, borosilicate glass) were sonicated (Fisherbrand S-line) in 2% Hellmanex III solution for 10 min. Afterwards they were rinsed with type I water and sonicated again in type I water for 20 min, followed by short immersions in ethanol and isopropanol respectively. The solvent was blow-dried off and the cleaned coverslips were stored in a sealed container until use. The gaskets (Grace Bio-Labs FastWells™ reagent barriers, 1-20 mm diam. X 1.0 mm depth) were sonicated in 2% Hellmanex III solution for 10 min. Afterwards they were rinsed with type I water and sonicated again in type I water for 20 min, followed by sonication in ethanol and isopropanol for 10 minutes respectively. The solvent was blow-dried off and the cleaned gaskets were stored in a sealed container until use.

The mixture of lipids was prepared by dissolving the lipids 1,2-dioleoyl-sn-glycero-3-phosphocholine (DOPC, Avanti®), 1,2-dioleoyl-*sn*-glycero-3-phospho-L-serine sodium salt (DOPS, Avanti®), 1,2-dioleoyl-*sn*-glycero-3-phosphoethanolamine-*N*-(cap biotinyl) sodium salt (Biotinyl-CAP-PE, Avanti®), 1,2-dioleoyl-*sn*-glycero-3-phospho-(2-azido-*N*-(2-hydroxyethyl)-acetamide) (DOPE-$N_3$, Nanocs®) and 1,2-dioleoyl-*sn*-glycero-3-phosphoethanolamine (DOPE, Avanti®) in a ratio of DOPC:DOPS:Biotinyl-CAP-PE:DOPE-$N_3$:DOPE (69%:5%:1%:25-n%:n%) in 2 mL $CHCl_3$ in a 5 mL vial, resulting in a final concentration of 1 mg/mL. The solvent was evaporated under a gentle flow of nitrogen, while turning the vial to generate a thin lipid film. Remaining solvent was removed under high vacuum for 15 minutes. To the vial 2 mL of type I water were added, thoroughly vortexed and transferred into a 15 mL falcon tube. The mixture was cooled to 0 °C and sonicated (QSonica Q700) for 25 minutes with an amplitude of 25% (on/off: 1 sec). Afterwards, the suspension was transferred into two 1.5 mL reaction vials and centrifuged for 20 min @16.000 × g at 4 °C (Eppendorf centrifuge 5417R). The small unilamellar vesicles (SUVs) containing supernatant was transferred to a separate vial and stored at 4 °C for a maximum of 2 weeks.

Directly before SLB formation the coverslips were plasma cleaned (10 min, oxygen, maximum power, Diener®). The gasket was attached and 45 µL type I water, then 15 µL of the corresponding SUV solution was added. To help SLB formation 1 µL of 50 mM aq. $MgCl_2$ solution was added. After 15 minutes at room temperature the SLBs were washed a total of 5 times. Each washing step consists of adding 50 µL of type I water first, followed by gently pipetting up and down five times, then discarding 50 µL of water.

To the formed SLB 50 µL of glycan click mixture (50 µM $CuSO_4$, 2.5 mM sodium ascorbate, 1 mM aminoguanidine hydrochloride, 250 µM tris(3-hydroxypropyltriazolylmethyl)amine and 100 µM of propargyl glycan in type I water) were added, followed by gently pipetting up and down five times, then discarding 50 µL. Further 50 µL of glycan click mixture were added followed by gently pipetting up and down five times and incubated for 30 min at room temperature. Afterwards, the SLBs were washed a total of 5 times. Each washing step consists of adding 50 µL of type I water first, followed by gently pipetting up and down five times, then discarding 50 µL of water. Finally, blocked AuNPs were prepared by addition of 450 µL of a 5 nM biotin solution in type I water to 50 µL of streptavidin functionalized gold nanoparticles (abcam®, 40nm, 10 OD). From this solution 15 µL were added to the SLBs, which were then directly investigated by iSCAT microscopy.

#### 5. iSCAT Data Analysis

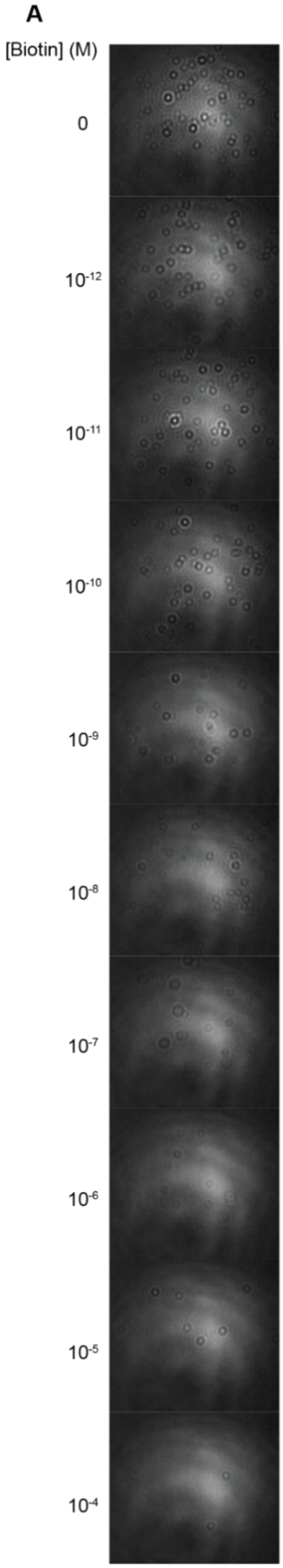

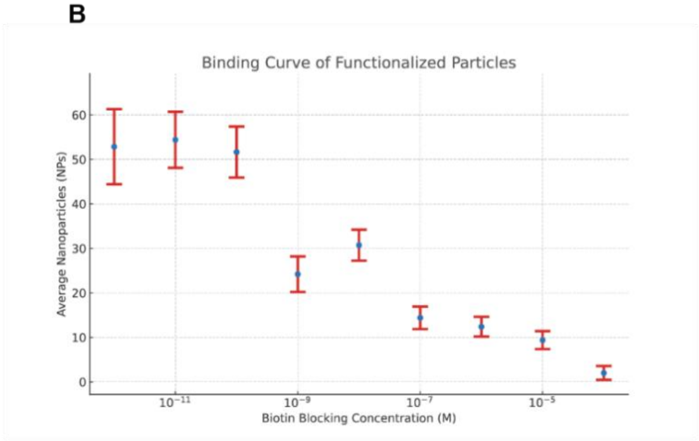

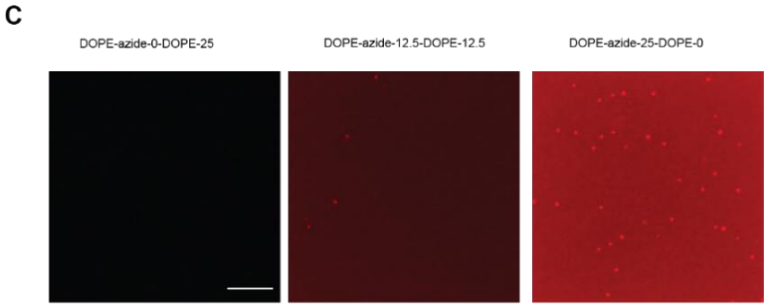

**Figure S76. Characterization of AuNP valency.** Streptavidin-functionalized AuNPs (abcam®, 40nm, 10 OD) were blocked in different concentrations of biotin in PBS. At low concentrations, few binding sites are blocked. At higher concentrations, the fraction of free binding sites is significantly reduced, yielding mainly monovalent AuNPs. In the following, we chose a biotin concentration of 5 nM for blocking, thus achieving an optimal trade-off between sufficient binding probability and reduction of multivalency. A: Imaging AuNPs on SLBs on coverslips within a fixed FoV at different concentrations. B: Number of AuNPs identified within the FoV vs biotin concentration. C: Imaging the effect of substituting DOPE with DOPE-azide (0%, 50%, 100% substitution) during SLB formation and subsequent copper-catalysed azide–alkyne cycloaddition with alexa fluor 647 alkyne (Thermo Fisher®) to validate the presence of free azido functional groups on the SLB.

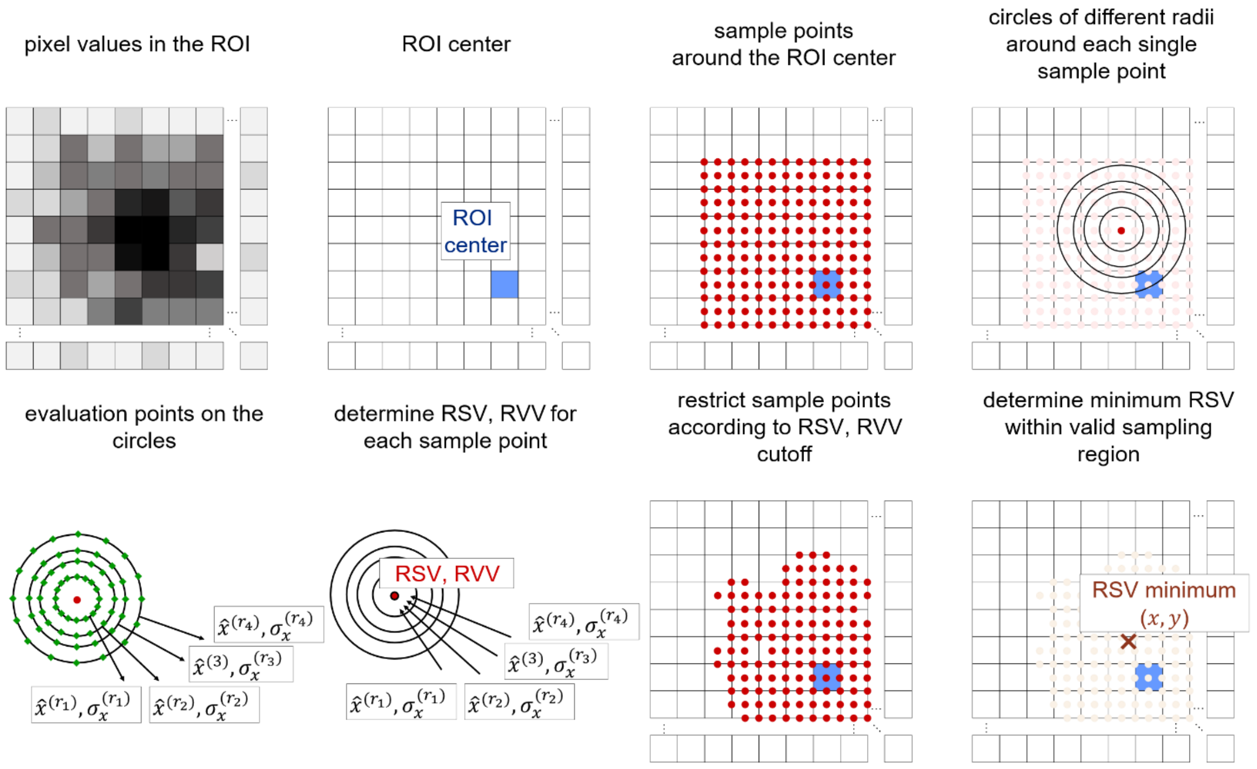

**Figure S77. Tracking.** For each recorded video file, we manually select the estimated locations of all AuNP PSFs in the first frame. The subsequent analysis steps are fully automated and thus do not require additional user input. In each single frame, we continue by restricting the frame to a square-shaped ROI around a central pixel, specified either by the manually selected location (in the first frame) or the pixel closest to the determined PSF center in the previous frame (for all subsequent frames). The pixel values in the ROI are normalized to the interval [0,1]. The ROI size was set to 31px, corresponding to ≈ 1.2 µm. Within the ROI, we determine a radial symmetry center. This is implemented via the steps described below.

1. A set of sample points around the ROI center is chosen. These are arranged in a square, spaced by 0.5 px and extend to a certain fraction of the ROI (0.8 for the first frame, 0.4 for all subsequent frames).
2. We interpolate the normalized pixel values on the integer pixel coordinates.[19]
3. On concentric circles around each sample point, a set of interpolation evaluation points is defined. Here, we choose circles of radii 5, 7, 10 and 13 px and 50 evaluation points on each circle. On these points, we evaluate the interpolation function from 2.
4. On each of the 4 circles, we determine the standard deviation and the mean value of the interpolation function values.
5. We assign (a) a 'radial symmetry value' (RSV) to the sample point defined by the mean value of the standard deviations obtained from the four circles and (b) a 'radial variance value' (RVV) defined by the variance of the mean values corresponding to the four circles.
6. We remove all points from the sample set (step 1.) for which the RVV is below the 20$^{th}$ percentile of all RVVs or for which the RSV is above the 20$^{th}$ percentile of all RSVs.
7. On the remaining sample points, we apply an RBF interpolator[20] to interpolate on their RVVs.
8. We find the location of optimal RSV by minimization of the function from step 7.[21] This location we assume to be the best estimate for the PSF center.

9. Finally, collecting all PSF locations across 27560 frames, we save the particle trajectory as a sequence of x- and y- coordinates along with the time stamps.

For the subsequent global analysis steps, if not stated otherwise, we split the full trajectories of duration 1.1 s and define individual particle trajectories of 110 ms duration. We discard any trajectories where the particle moved a distance greater than 190 nm between two consecutive frames to rule out tracking errors. In total, we obtain 42035 trajectories across the 19 conditions. For any 110 ms trajectory we calculate the mean squared displacements (MSDs) for time delays [$\tau = 0.04 - 27.6$ ms (corresponding to 1-690 frames)]. Assuming perfectly Brownian diffusion, we estimate a 2D diffusion constant for each particle by fitting a linear function to this curve[22], according to:

$$MSD(\tau) = 4D\tau + 2\sigma^2.$$

Here, $\tau$ is the time delay, $D$ is the diffusion constant and $\sigma$ denotes the localization precision. Only in case of Brownian motion, in the absence of superdiffusive or subdiffusive behavior, does the MSD scale linearly with the time delay. Therefore, we have focused on MSD values and in the following a Brownian diffusion coefficient serves as an approximate measure that allows for comparison with previous results.

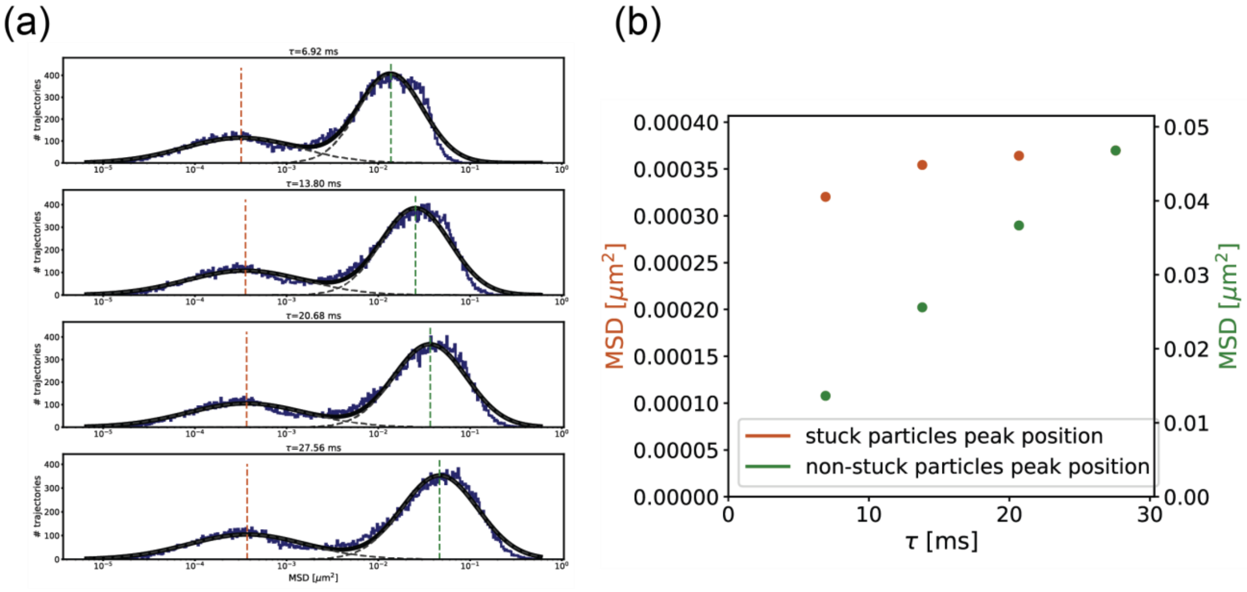

**Figure S78. Definition of particle mobility.** Further examining the distribution of trajectories with different MSDs at several time delays, we observe that this distribution shows two separate peaks. This agrees with the observation that a significant number of particles were immobile on the SLBs. Since we want to concentrate on the diffusing particles in subsequent analysis steps, we remove immobile particles' trajectories from the initial set of trajectories. (a) Fitting two Gaussians to the distribution of MSDs in logarithmic scale at several time delays $\tau$, we obtain the MSD cutoff as the intersection of the two Gaussians at $\tau$ = 27.56 ms, which is 0.0049773 $\mu m^2$. (b) Note that the peak positions truly correspond to two separate particle populations, i.e. immobile and mobile particles: The right peak position shows a linear increase with time delay $\tau$ while the immobile population peak position does not change significantly.

(a)

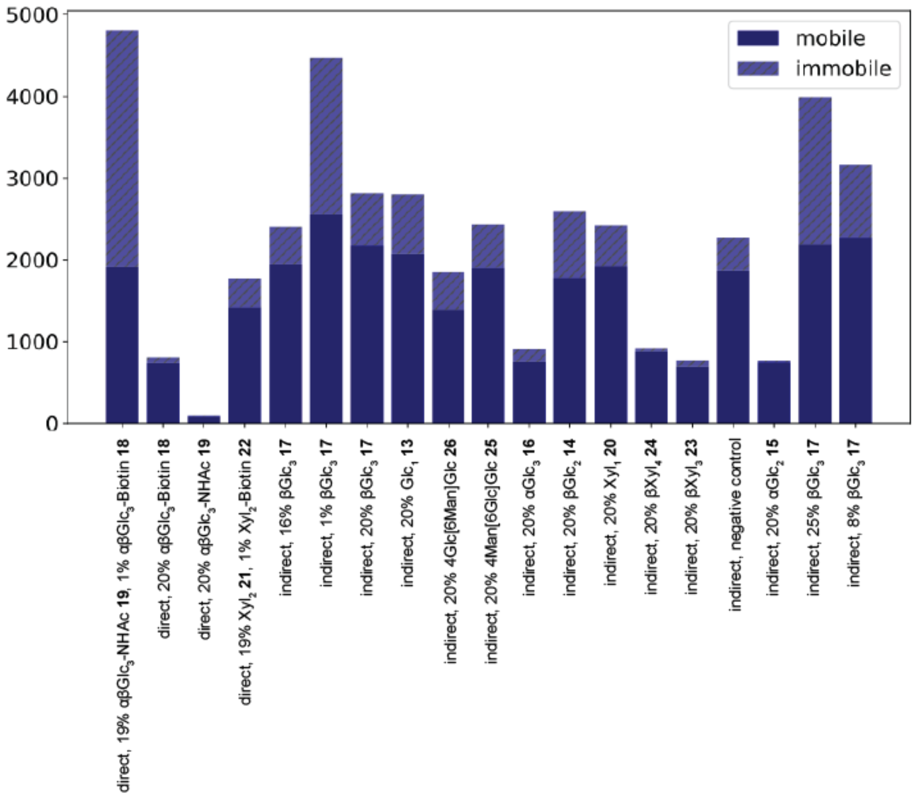

(b)

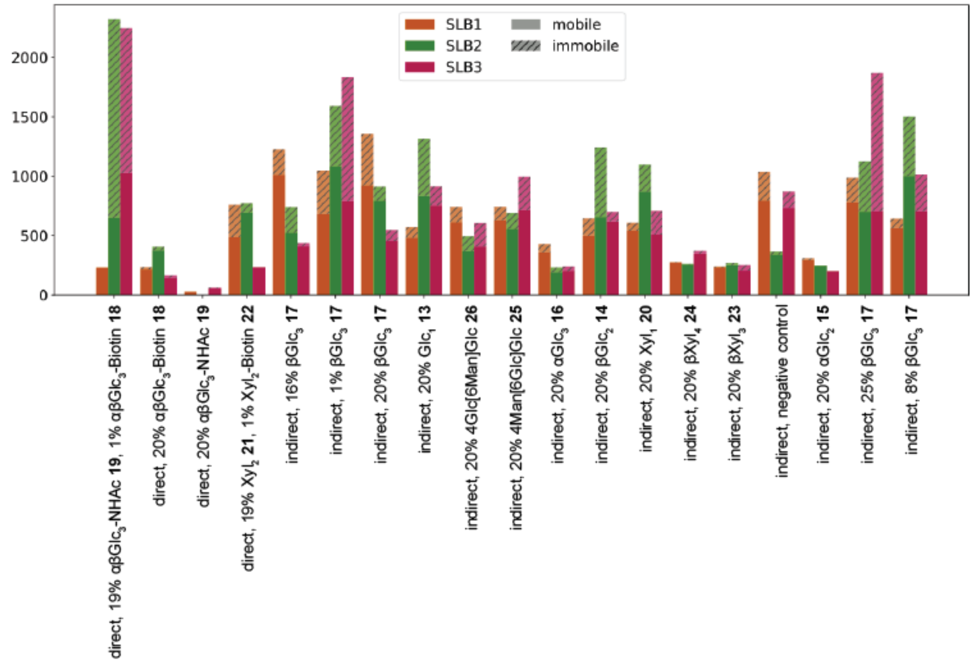

**Figure S79. Mobile and immobile particles.** The share of immobile particles varies among different conditions and SLBs prepared under the same conditions. (a) Number of trajectories of mobile and immobile particles per condition. (b) Number of trajectories of mobile and immobile particles per SLB per condition.

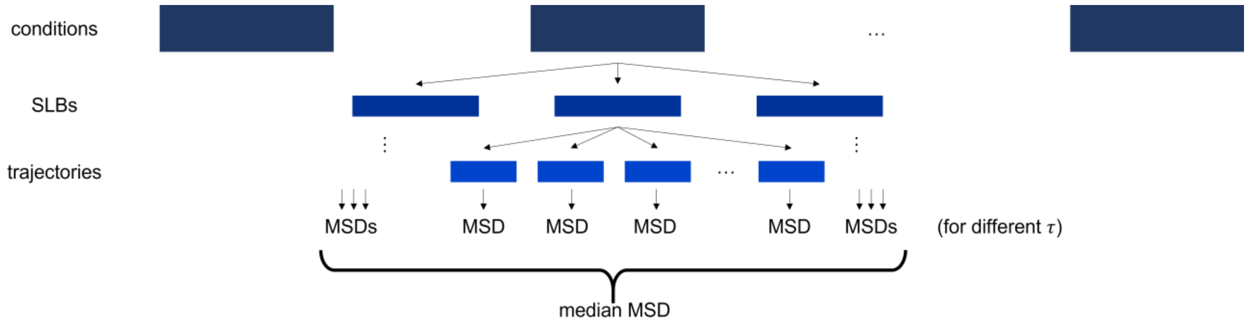

**Figure S80. Global MSD calculation.** For all of the considered conditions and at each time delay, we determine the median MSD across all the mobile trajectories belonging to the condition.

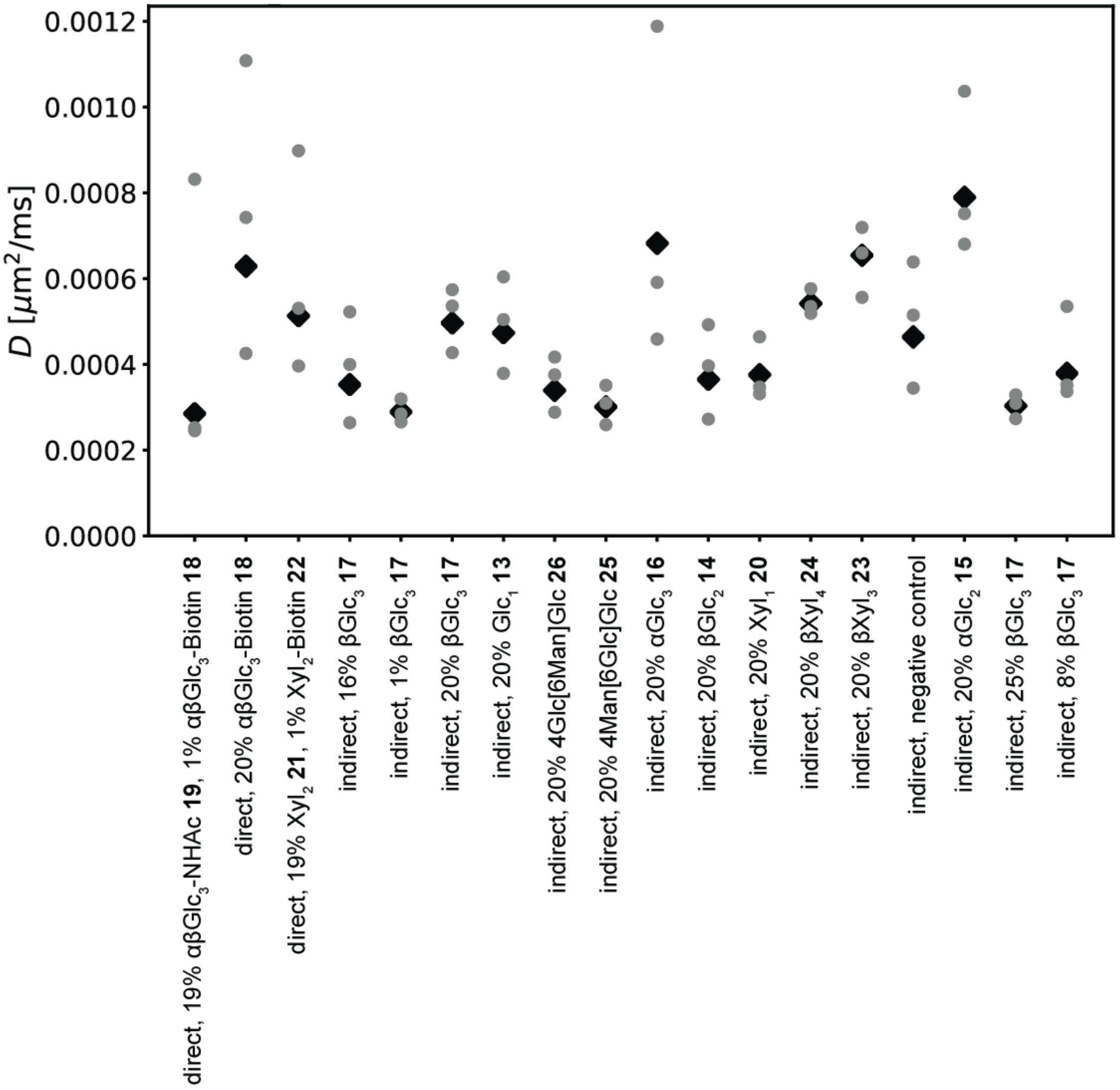

**Figure S81. Diffusion constants.** To each condition, and to each SLB separately, we assign a diffusion constant via linear fit of the median MSD vs. time delay $\tau$ (see above). The results are summarized here. The black diamonds represent the value obtained from fitting to the median across all SLBs, the grey points denote the values obtained from fitting to the median of a single SLB.

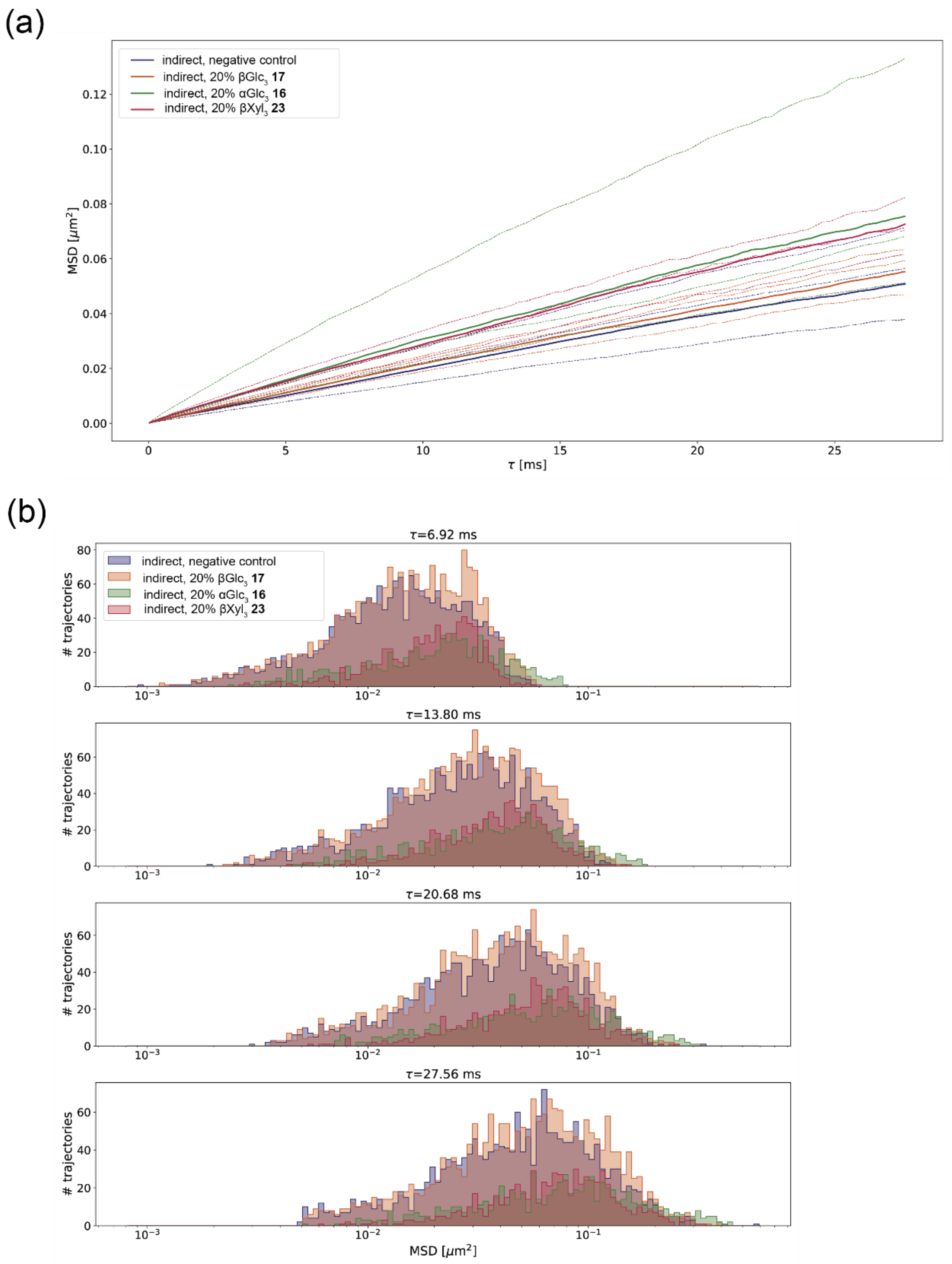

**Figure S82. Influence of glycan stereochemistry on AuNP diffusion.** We investigate the influence of glycan stereochemistry on the obtained MSDs. Larger MSDs signify larger displacements after a fixed time delay, i.e. faster diffusion. (a) The solid lines show the median MSDs across a condition, the dashed lines the median MSDs across an SLB belonging to a certain condition versus time delay. There is a tendency towards faster diffusion on SLBs glycosylated with αGlc$_3$ **16** and βXyl$_3$ **23** than in the condition with βGlc$_3$ **17** (or for the negative control). (b) Histograms showing the MSD distributions for the indicated time delays for the negative control and βGlc$_3$ **17**, αGlc$_3$ **16**, βXyl$_3$ **23**. For the latter two, the peaks of the distribution are at slightly higher MSD values.

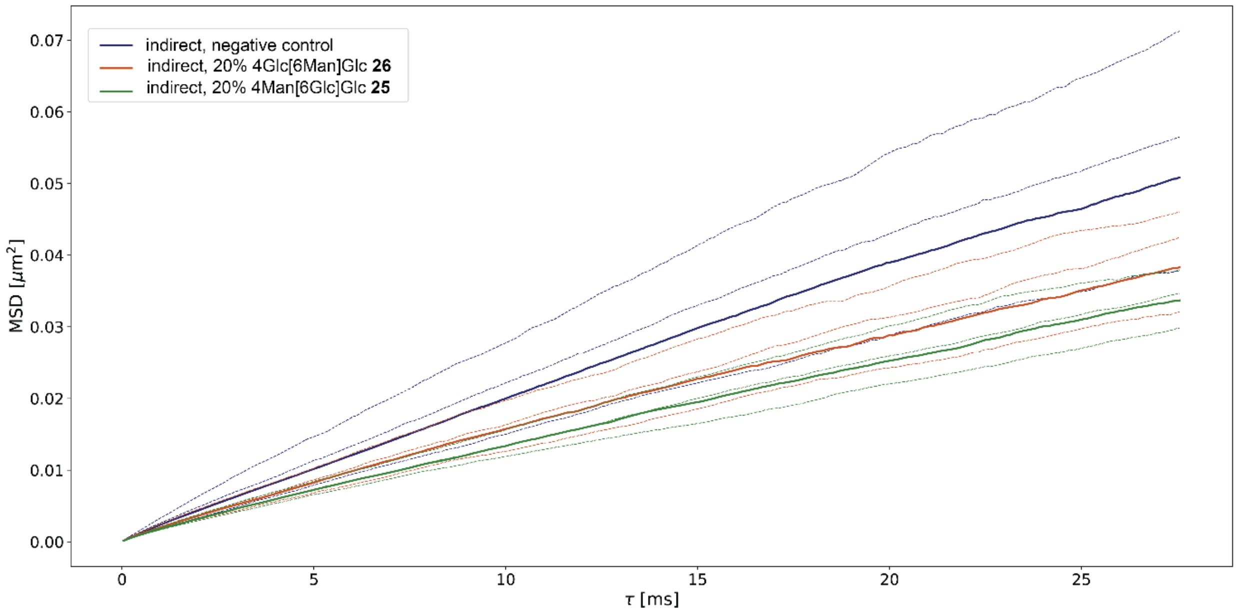

**Figure S83. Influence of glycan regiochemistry on AuNP diffusion.** We investigate the influence of glycan regiochemistry on the obtained MSDs. Larger MSDs signify larger displacements after a fixed time delay, i.e. faster diffusion. The solid lines show the median MSDs across a condition, the dashed lines the median MSDs across an SLB belonging to a certain condition versus time delay. The diffusion on SLBs glycosylated with 4Glc[6Man]Glc **26** and 4Man[6Glc]Glc **25** is slower than in the negative control condition. However, the difference is not as significant as for the other comparisons. For these specific conditions, the glycan branching appears to have only little effect on diffusion behaviour. However, we can observe a significant difference compared to the negative control.

(a)

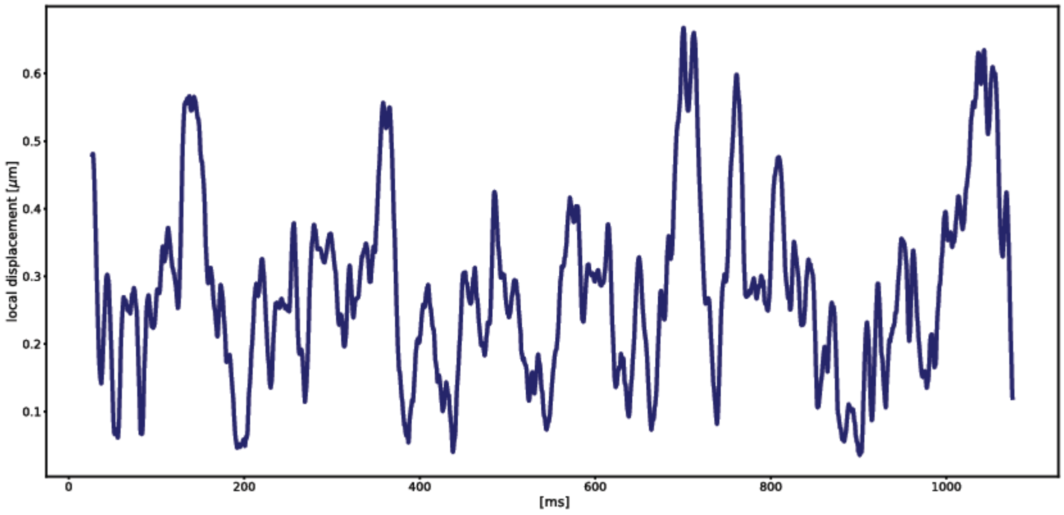

(b)

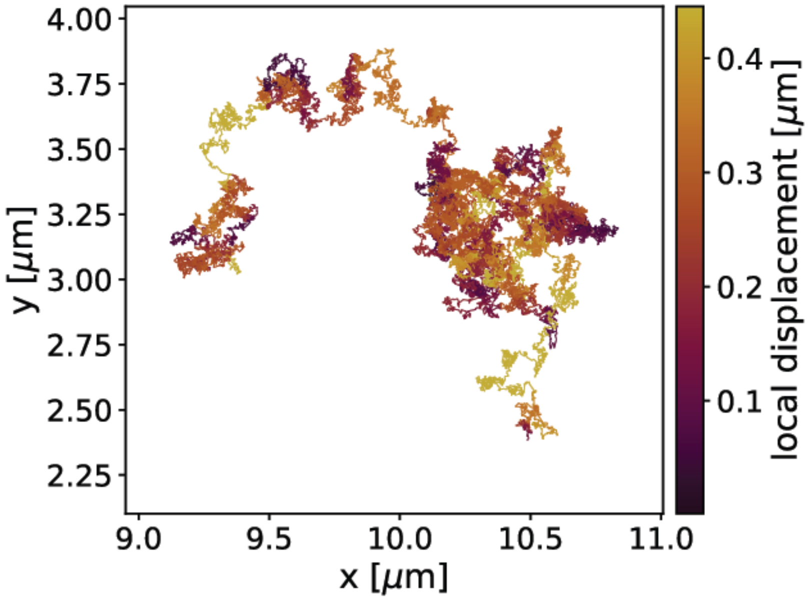

**Figure S84. Local displacement for Glc$_1$ 13.** Averaging over a time window of 4.0 ms duration, we determine the local displacement at each point in time, here exemplarily for one representative trajectory from the Glc$_1$ **13** condition: $\mathrm{MSD}(\tau) = \langle(\Delta\vec{r}\,(t, t+\tau))\rangle_t$. (a) The averaged local displacement versus time. Short periods of high local displacement values are alternating with periods of lower local displacement values. (b) The representative trajectory, colored according to the calculated local displacement values.

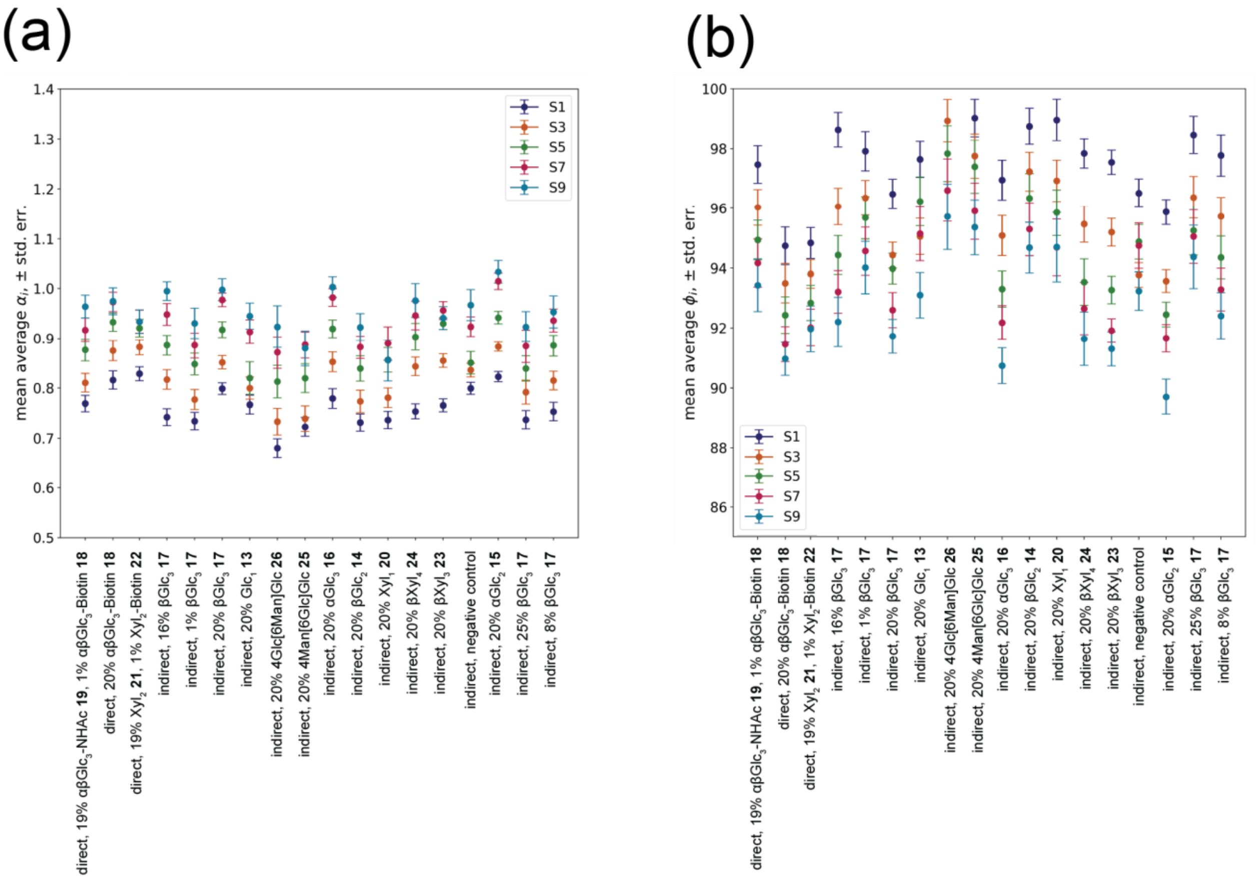

**Figure S85. Confinement analysis for different time scales.** We determine the mean local anomalous diffusion exponents $\alpha_i$ and deflection angle $\phi_i$ as described in the main text for all representative trajectories of all 19 conditions for different time scales S1-S10 (Table S1). For all conditions, the local anomalous diffusion exponent increases for longer time scales while the deflection angle decreases. Moreover, on all time scales, the trends signifying differences between conditions are similar.

**Table S1.** Different time scales for local anomalous diffusion exponents $\alpha_i$ and deflection angle $\phi_i$ calculations in Figure S85.

| Setting | S1 | S2 | S3 | S4 | S5 | S6 | S7 | S8 | S9 | S10 |
|---|---|---|---|---|---|---|---|---|---|---|
| $\boldsymbol{\tau_1}$ $(\boldsymbol{\alpha_i})$ [ms] | 0.24 | 0.4 | 0.8 | 1.6 | 2.4 | 4.0 | 6.0 | 8.0 | 12.0 | 20.0 |
| $\boldsymbol{\tau_2}$ $(\boldsymbol{\alpha_i})$ [ms] | 0.48 | 0.8 | 1.6 | 3.2 | 4.8 | 8.0 | 12.0 | 16.0 | 24.0 | 40.0 |
| $\boldsymbol{\tau}$ $(\boldsymbol{C})$ [ms] | 0.12 | 0.2 | 0.4 | 0.8 | 1.2 | 2.0 | 3.0 | 4.0 | 6.0 | 10.0 |

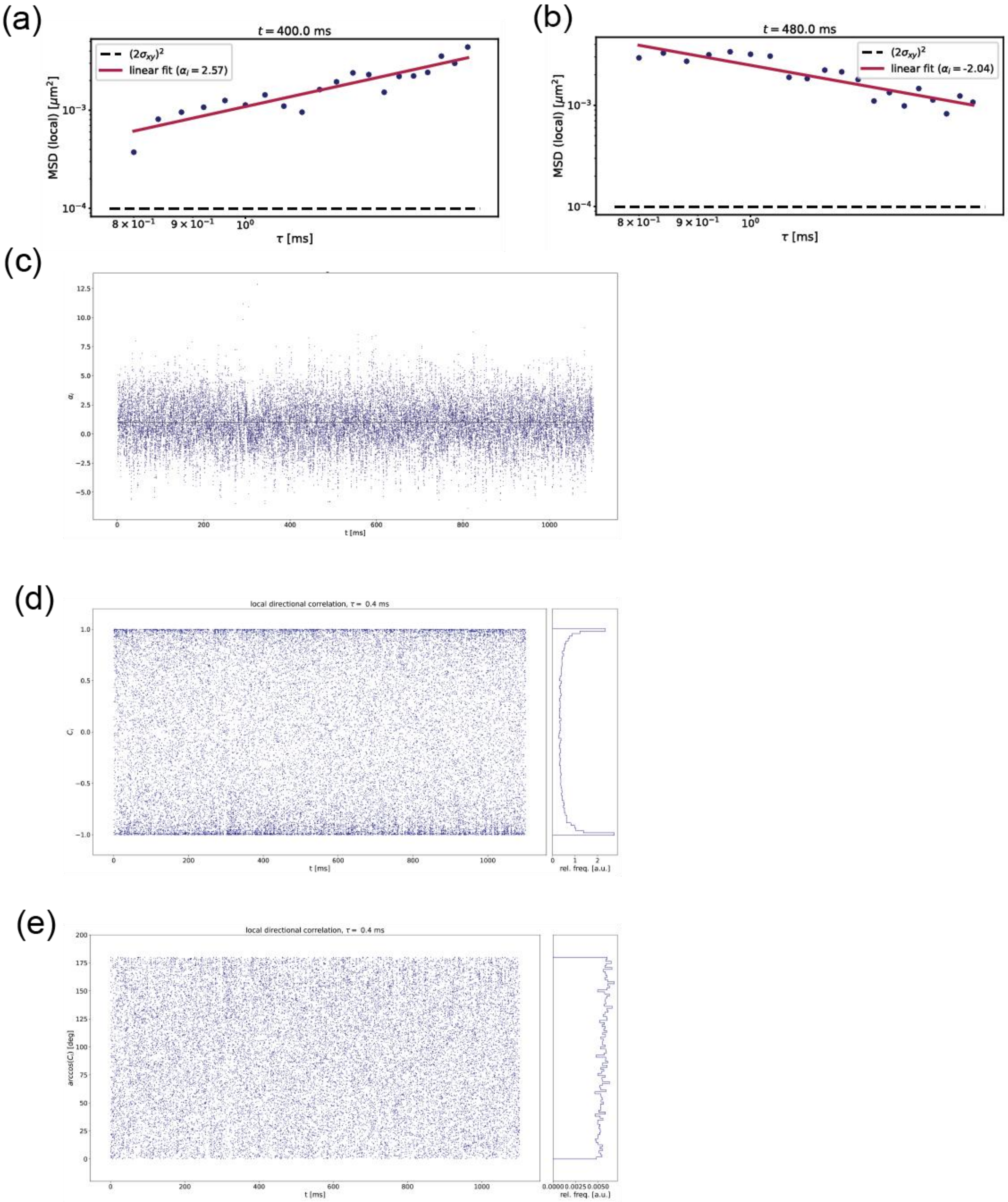

**Figure S86. Confinement analysis for Glc$_1$ 13.** Exemplarily for one representative trajectory of the Glc$_1$ **13** condition and setting S3, we show how the local metrics are determined. (a) Fitting of the local anomalous diffusion exponent $\alpha_i$ at $t = 400$ ms. Local squared displacement values below $10^{-4}\ \mu m^2$ (black dashed line, value derived from $(2\sigma)^2$ for an estimated localization precision of $\sigma = 5$ nm) are discarded from the data set for fitting. If the linear fit does not converge, no local $\alpha_i$ is determined and the time point will not contribute to the average. (b) Fitting of the local anomalous diffusion exponent $\alpha_i$ at $t = 480$ ms. The particle moves closer to the position at $t = 480$ ms between $t + \tau_1$ and $t + \tau_2$, thus yielding a negative $\alpha_i$. (c) Scatter plot of all fitted $\alpha_i$ at every point in time. The black dashed line at $\alpha_i = 1$ represents the limit between sub- and super-diffusive behavior. (d) Scatter plot of all calculated $C_i$. On the right, a histogram shows the relative frequencies of occurrence of the $C_i$ over time. (e) Scatter plot of all calculated $\phi_i$. On the right, a histogram shows the relative frequencies of occurrence of the $\phi_i$ over time.

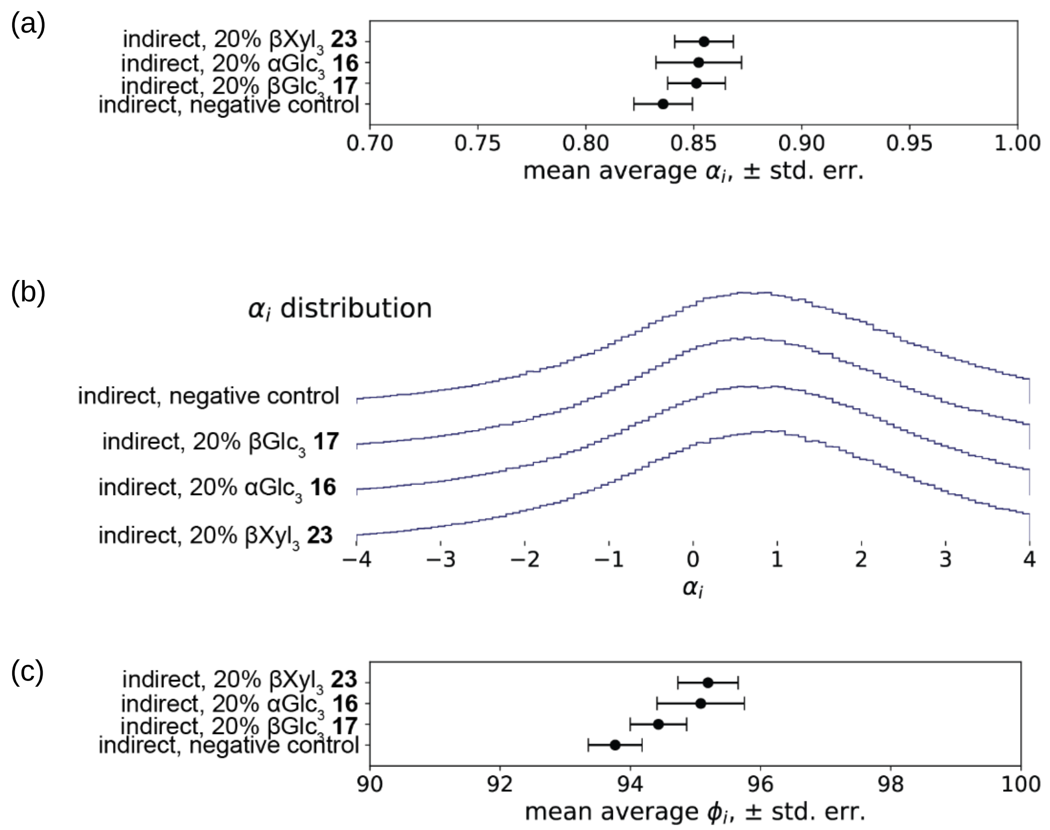

**Figure S87. Influence of glycan stereochemistry on confinement.** We investigate the influence of glycan stereochemistry on the average anomalous diffusion coefficient $\alpha_i$ and average deflection angle $\phi_i$. (a) There are no significant differences between the obtained mean average $\alpha_i$ between αGlc$_3$ **16**, βXyl$_3$ **23,** βGlc$_3$ **17** and they are only slightly higher than for the negative control. (b) Histograms of the distribution of the $\alpha_i$ values for all time points and all 36 representative trajectories for αGlc$_3$ **16**, βXyl$_3$ **23,** βGlc$_3$ **17** and the negative control. (c) The mean average deflection angle $\phi_i$ is slightly larger for αGlc$_3$ **16**, βXyl$_3$ **23,** βGlc$_3$ **17** than for the negative control. We observe a tendency towards smaller deflection angles $\phi_i$ for βGlc$_3$ **17** compared to αGlc$_3$ **16**, βXyl$_3$ **23**.

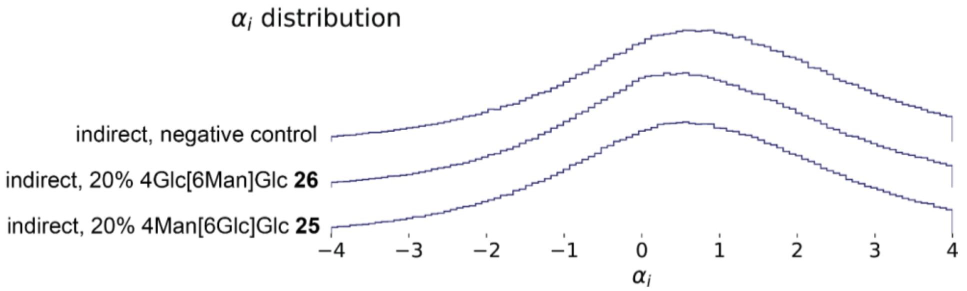

**Figure S88. Influence of glycan regiochemistry on confinement.** We investigate the influence of glycan regiochemistry on the average anomalous diffusion coefficient $\alpha_i$ and average deflection angle $\phi_i$. Histograms of the distribution of the $\alpha_i$ values for all time points and all 36 representative trajectories for 4Glc[6Man]Glc **26**, 4Man[6Glc]Glc **25** and the negative control. There are no significant differences between the distributions obtained for 4Glc[6Man]Glc **26** and 4Man[6Glc]Glc **25** but they are shifted to the left compared to the negative control, signifying stronger confinement.

#### 6. Computational Modeling

##### 6.1. Linker Parameterization

Molecular mechanics (MM) parameter for sugars[23,24] and lipids[25] were taken from the CHARMM36 force field family (version July 2022). However, parameters for the triazole linker between the phospholipid and the carbohydrate headgroup of the azido-functionalized lipids were not available in the canonical CHARMM36 force field. For the simulations in this work, parameters for the triazole linker were therefore generated based on the CHARMM General Force Field (CGenFF, v4.6).[26] Due to the size of the linker region, no bonded potentials (i.e., bond, angle, or dihedral) needed to be defined between atom types of the carbohydrate and the lipid force field. For each of the two force field combinations (**A** CGenFF-Carbohydrate, **B** CGenFF-Lipid), separate molecules were created using the Avogadro software suite[27,28] (v1.2.0) to determine partial charges and identify missing bonded potentials. Both small molecules contained substructures already described by the CHARMM force field, so their partial charges were directly adopted from the larger molecule. Upon connecting two molecules, the partial charges of the removed hydrogen atoms were summed into those of the corresponding heavy atom (e.g., carbon atom).[26] Bonded parameters for combinations of atom types not yet included in the force field were derived via analogy from already existing parameter sets.[23–26,29–31] Parameters with poor analogies were optimized using *lsfitpar* (v0.9.1 beta)[32] via the FFParam package (v1.2.0).[33] It should be noted that such poor analogies were only identified for proper dihedrals. Accordingly, for the refinement of proper dihedral parameters, we employed the following procedure:

1. An initial stream input file for the molecule was automatically generated via the CGenFF webserver (v2.5),[34] based on the algorithms developed by Vanommeslaeghe et al.[35,36] Atom types and partial charges were subsequently edited according to the sub-structures found in the respective force fields.
2. A quantum mechanical (QM) optimization of the initial structure was performed at the MP2/6-31G(d) level of theory, as recommended for the CHARMM force field.[26]
3. The QM optimized structure served as starting point for potential energy surface (PES) scans again at the MP2/6-31G(d) level of theory. Unless otherwise specified, full 360° scans with a 10° step size were performed. The scanned dihedral was fixed, while the remaining structure was allowed to relax.
4. The relaxed QM structures were subsequently minimized using classical mechanics constraining all dihedrals of interest with a force constant of 9,999 kcal $mol^{-1}$ $rad^{-2}$.[32] First, the steepest descent algorithm was applied with a maximum number of 100 steps and a gradient tolerance of 0.2 kcal $mol^{-1}$ $Å^{-1}$, followed by conjugate gradient minimization with a maximum of 300 steps and a gradient tolerance of 0.0001 kcal $mol^{-1}$ $Å^{-1}$. If the root mean square gradient would have been greater than 0.0001 kcal $mol^{-1}$ $Å^{-1}$ after the two minimization steps, an additional Newton-Raphson minimization was performed with a step limit of 100 and a gradient tolerance of 0.0001 kcal $mol^{-1}$ $Å^{-1}$.
5. Both relaxed 1-dimensional dihedral scans (QM & MM) from all selected dihedrals were used as input for lsfitpar[32] to refine the MM parameters of all dihedrals simultaneously. Notably, the current version of FFParam[33] always sets the initial force constant for the selected dihedrals to 0.

6. To ensure the self-consistency of the force field, steps 3-5 were repeated. As input structure for the second QM PES scan, the QM-optimized structure was energetically minimized using the fitted MM parameters. This minimization was performed in two stages:[33] first, a conjugated gradient minimization with a maximum of 1000 steps, followed by an Adopted Basis Newton-Raphson minimization with a maximum of 500 steps and a gradient tolerance of 0.00001 kcal $mol^{-1}$ $Å^{-1}$.

The CHARMM software package (developmental version 47b1)[37] was used to perform all MM calculations, while the open-source software Psi4[38] was used to perform all QM calculations. Input files for all types of performed calculations were generated with the FFParam suite.[33]

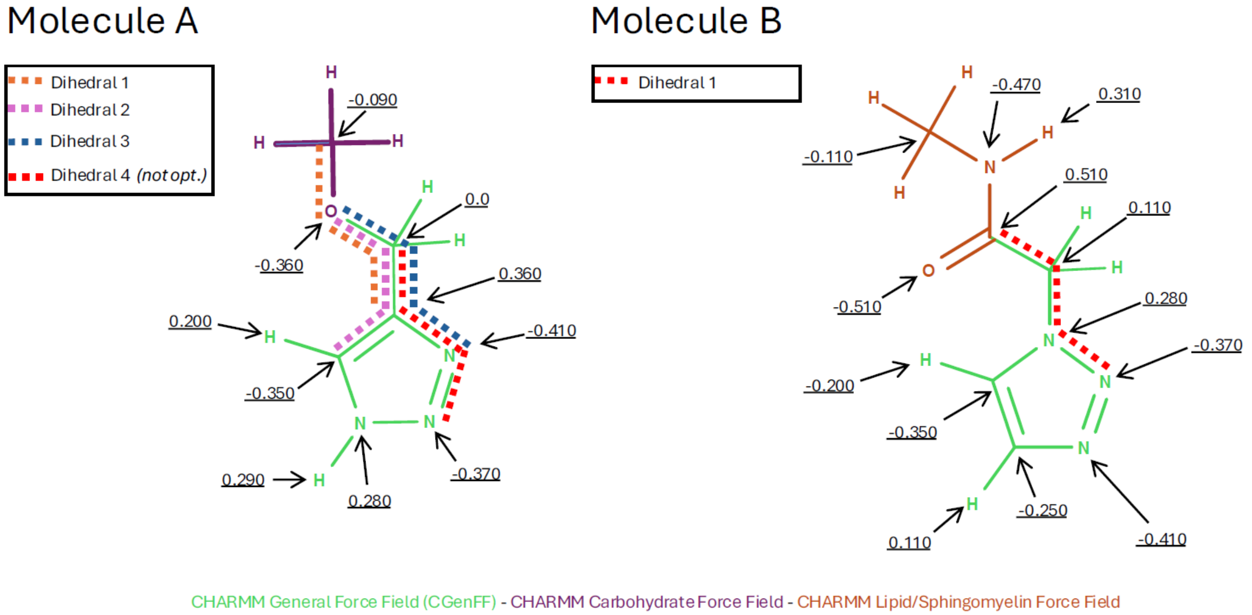

**Figure S89.** Structures of molecule **A** and molecule **B** used for the parameterization of the triazole linker connecting the lipid anchor to the carbohydrate headgroup. Colors indicate atom types from different CHARMM force fields, while underlined values are the partial charges in units of the elementary charge. Hydrogens without an explicit charge value were assigned a value of 0.09e. Dihedrals selected for further optimization are shown as dashed lines.

For molecule **A**, the small molecule dimethylether was added to the fourth position of 4,5-Dihydro-1H-1,2,3-triazole. Their partial charges were taken from the glycolipid force field[29] (resname: C12DEG) and CGenFF[26] (resname: TRZ3) (see Fig. S89, molecule **A**). Good analogies for parameters of missing bond and angle combinations were found in the force fields. However, four dihedrals were identified with only poor analogies (see Fig. S89, molecule **A**). One of these dihedrals (see Fig. S89, molecule **A**, dihedral 1) was declared as non-rotatable since it included three ring-atoms, and was therefore not considered for further optimization. The remaining three dihedrals were optimized using the aforementioned workflow.

For molecule **B**, the ethyl chain in 1-Ethyl-1,2,3-triazole (resname: ETRZ) was replaced with a methyl acetamide group. MM parameters for ETRZ were directly copied from CGenFF (see Fig. S89, molecule **B**). For the amide motif, atom types were taken from the CHARMM force field for sphingomyelin lipids, while partial charges were adopted from the CHARMM protein force field. In this structure, only one dihedral was identified for further optimization (see Fig. S89, molecule **B**).

Optimization of the selected dihedral parameters through the lsfitpar routine[32] resulted in an excellent approximation of the QM PES by the corresponding MM energies for both molecules (see Fig. S90). The optimization started always with the force constants set to 0.

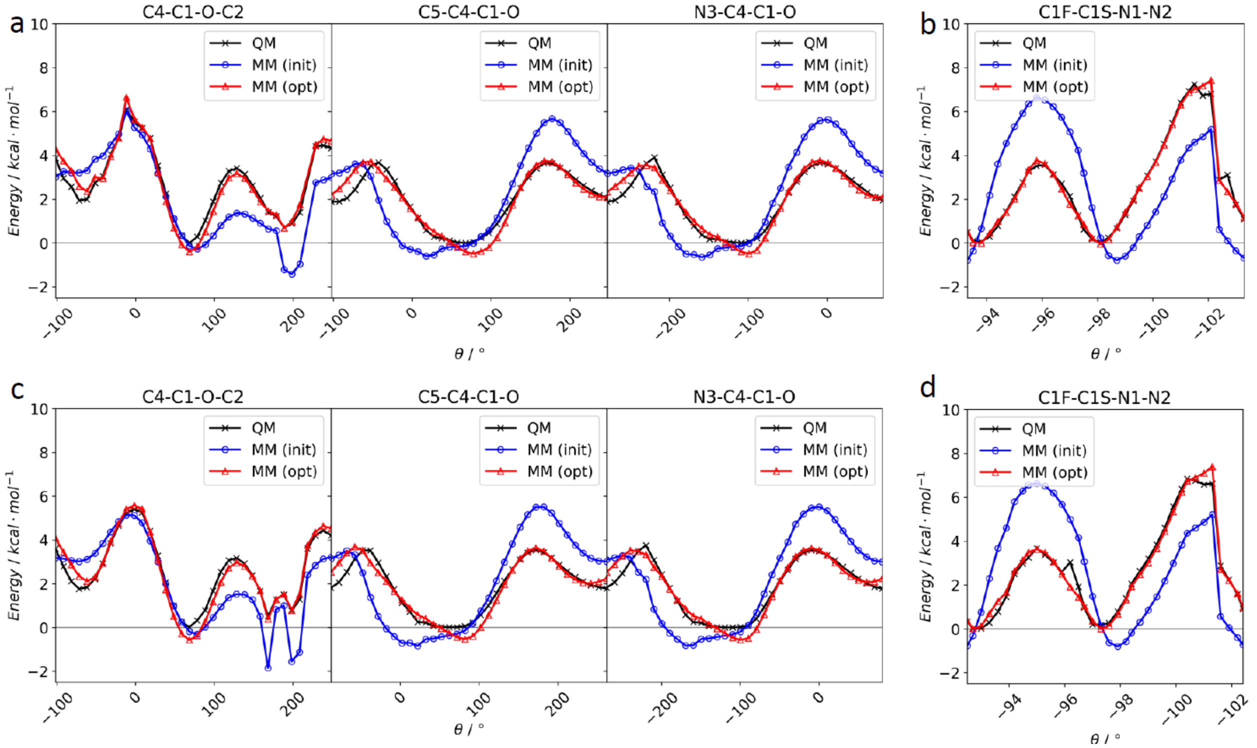

**Figure S90.** Results of the *lsfitpar* fitting routine[32] for the first (top panel) and second (bottom panel) parameterization rounds of molecule **A** (a, c) and molecule **B** (b, d). In the QM PES scan (black), only the scanned dihedral is restrained, while in the MM PES scans (red, blue), all fitted dihedrals are restrained. The blue curve represents the MM energies with the initial parameters (force constants set to 0), and the red curve shows the MM energies after optimization of force constants and periodicities.

To assess the quality of the refined parameters in the context of the full force field, the MM PES were re-calculated restraining only the scanned dihedral and allowing the remaining degrees of freedom to relax. Fig. S91 compares the performance of the refined parameters and the initial parameters. For both molecules **A** (Fig. S91 a) and **B** (Fig. S91 b) a large improvement of the MM PES is observed after the first round of optimization, and only minor changes after the second round of optimization (Fig. S91 c, d). The MM energies still approximate the QM PES very well, but some details seen in the QM calculations were lost through the structure optimization with the MM parameters (e.g., in Fig. S91 c between 150° and 200°).

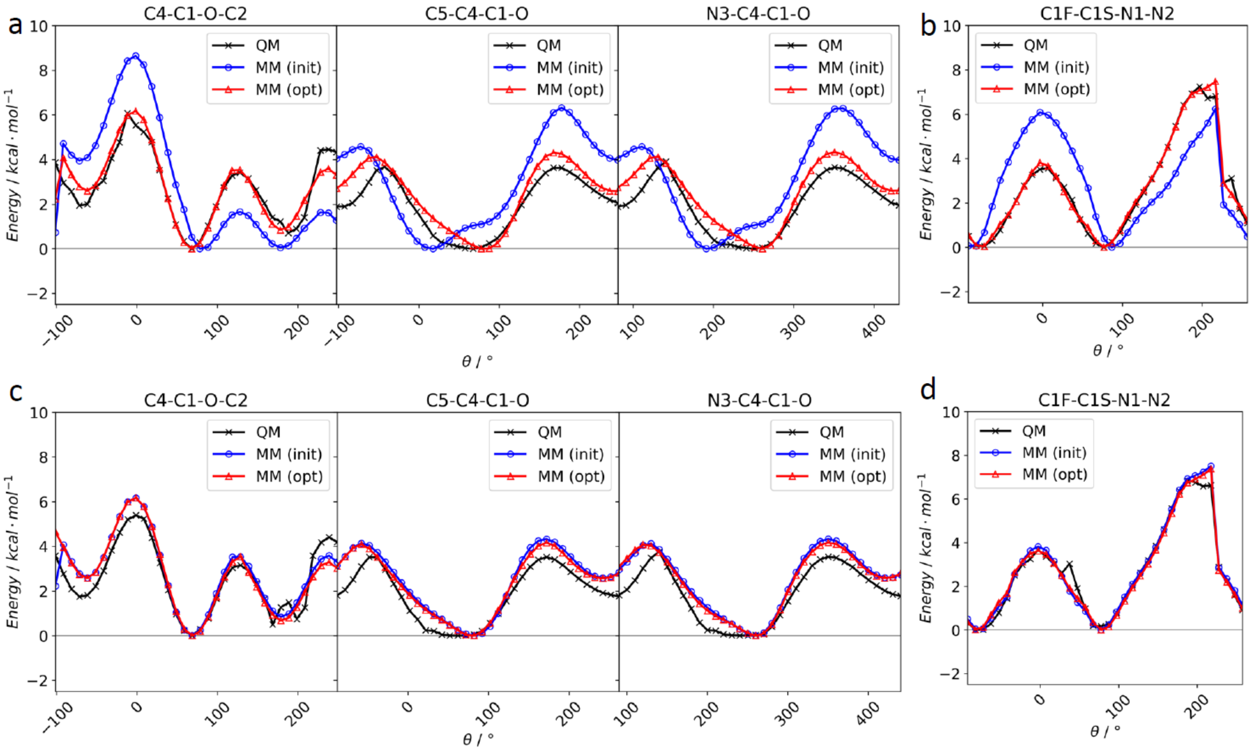

**Figure S91.** Comparison of QM and MM potential energy surfaces (PES) before/after the first (top panel) and the second (bottom panel) parameterization rounds of molecules **A** (a, c) and **B** (b, d). In all PES scans, only the scanned dihedral angle is restrained. The blue curve represents the MM energies using parameters found by analogy (a, b) or parameters after the first round of fitting (c, d). The red curve shows the optimized MM energies after refinement of force constants and periodicities.

Fig. S92 shows structures of molecule **A** and **B** before and after an energy minimization using either quantum mechanics or classical mechanics with different sets of parameters. For molecule **A**, the initial set of MM parameters led to a significant deviation from the structure obtained at the MP2/6-31G(d) level of theory. The subsequent parameter optimization improved the structural alignment, despite some minor differences in the angle between the ring plane and the ether side chain. For molecule **B**, the parameter optimization of one dihedral improved the alignment between the QM and MM structure only slightly.

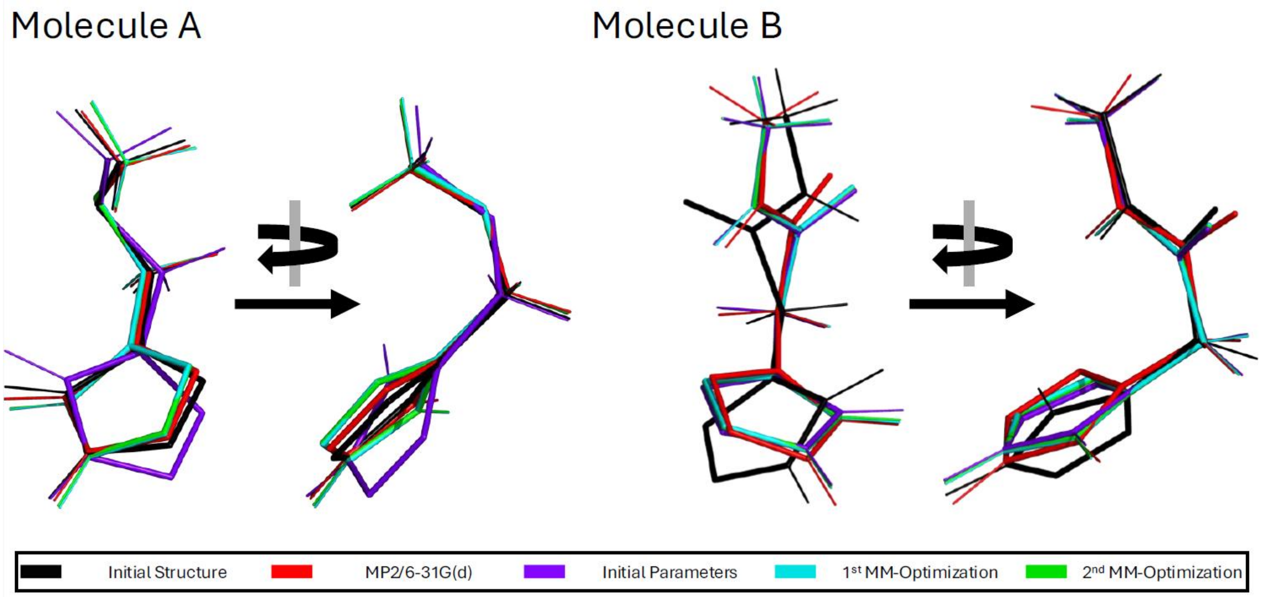

**Figure S92.** Minimum energy structures of molecules **A** and **B** at different optimization stages: initial structure (black), QM optimization at MP2/6-31G(d) (red), MM optimization with initial parameters (purple), MM optimization with parameters from first fitting (cyan), and MM optimization with parameters from second fitting (green). Optimization was performed in two stages:[33] first, a conjugated gradient minimization with a maximum of 1,000 steps, followed by an Adopted Basis Newton-Raphson minimization with a maximum of 500 steps and a gradient tolerance of 0.00001 kcal $mol^{-1}$ $Å^{-1}$. All structures were aligned towards the QM optimized structure using the *align* tool implemented in PyMOL.[39] Images were rendered with PyMOL.[39]

After the parameters for each separate molecule were defined, the CHARMM-readable files were converted via the *cgenff_charmm2gmx_py3_nx2.py* script, downloaded from the website of the MacKerell group,[40] and the *charmm2gmx* tool from Wacha & Lemkul[41] to a GROMACS readable format. Structures of the full azido-functionalized lipid with different glyco-headgroups were built with the Avogadro software suite (v1.2.0).[27,28] Final missing parameters that appeared while generating the topology of the whole lipids could be deduced via analogy from the existing force field. Energy minimization of the lipids with the steep descent algorithm (maximum of 20,000 steps and a gradient tolerance of 0.01 kJ $mol^{-1}$ $nm^{-1}$) implemented in the GROMACS MD engine[42] using the refined MM parameters converged to chemically reasonable structures.

##### 6.2. Atomistic Simulations

To study the dynamics of model glycocalyx systems, µs-long all-atom molecular dynamics (MD) simulations of a complex, symmetric lipid bilayer were performed using the CHARMM36 force field and GROMACS version 2023.3[42–44] in the isothermal-isobaric ensemble (NPT, 298K). The bilayer was composed of DOPC (1,2-dioleoylphosphatidylcholine, 69%), DOPE (1,2-dioleoylphosphatidylethanolamine, 6%), DOPS (1,2-dioleoylphosphatidylserine, 5%) and glycolipids (20%), amounting to a total of 500 lipids (250 per bilayer leaflet). The bilayer was solvated with water (TIP3P,[45,46] at least 59 water molecules per lipid) and (only) counter ions were added (either sodium or magnesium ions) to neutralize the negative charges of DOPS and the glycolipids. In total, 7 all-atom systems were prepared (see Table S2 for details).

1. The solvated systems were minimized using the steepest descent algorithm for 500 steps with an initial step size of 0.0001 nm, to resolve unnatural steric clashes.

2. A 5 ns gentle equilibration was then performed with a time step of $\Delta t$ = 1fs at 298K. Pair-lists were generated using the Verlet cutoff-scheme, with the Verlet buffer tolerance set to -1 to maintain a constant neighbor list update frequency of 20. Both temperature and pressure coupling update frequencies are also set to 20, following the recommendations by Kim et al.[47] The cut-off distance for the short-range neighbor list was set to 1.35 nm. Van der Waals forces were calculated using a plain cut-off scheme with a force-switch modifier, smoothly transitioning the force to zero for $r_{vdw} > r_{vdw\text{-}switch}$, with $r_{vdw}$ = 1.2 nm and $r_{vdw\text{-}switch}$ = 1.0 nm. Long-range electrostatics were handled using the Particle Mesh Ewald (PME) algorithm[48] with an initial $r_{coulomb}$ set to 1.2 nm. Temperature coupling was managed using the v-rescale with a time constant $\tau_T$ = 1.0 ps, and the membrane and solvent were coupled separately for all systems. Pressure was set to 1 bar and handled semi-isotropically with the c-rescale method, with the pressure, normal to the membrane plane (*z*) and the pressure in the *xy*-plane coupled separately. The compressibility for both groups was set to 4.5 $10^{-5}$ $bar^{-1}$ with a pressure coupling constant $\tau_p$ = 5.0 ps. Constraints were applied to hydrogen bonds and handled using the Linear Constraint Solver (LINCS) algorithm.[49] Center of mass motion was removed every 100 steps for the membrane and solvent separately.
3. After the short equilibration, the time step was increased to $\Delta t$ = 2 fs, keeping the rest of the molecular dynamics parameters the same as in the previous step, and MD simulations were run for 25 µs. The final 20 µs of each simulation were used for analysis.

**Table S2.** The composition of the eight all-atom glycocalyx model systems. $N_{Glc}$ stands for the number of glycolipids. The glycolipids are named after the sugar in their head group.

| System | Glycolipid | $N_{Glc}$ | $N_{DOPC}$ | $N_{DOPE}$ | $N_{DOPS}$ | $N_{ion}$ |
|---|---|---|---|---|---|---|
| 1 | βGlc **13** | 100 | 344 | 30 | 26 | 126 $Na^+$ |
| 2 | βGlc **13** | 100 | 344 | 30 | 26 | 63 $Mg^{2+}$ |
| 3 | α$Glc_2$ **15** | 100 | 344 | 30 | 26 | 126 $Na^+$ |
| 4 | α$Glc_2$ **15** | 100 | 344 | 30 | 26 | 63 $Mg^{2+}$ |
| 5 | α$Glc_3$ **16** | 100 | 344 | 30 | 26 | 126 $Na^+$ |
| 6 | α$Glc_3$ **16** | 100 | 344 | 30 | 26 | 63 $Mg^{2+}$ |
| 7 | - | 0 | 430 | 38 | 32 | 16 $Mg^{2+}$ |

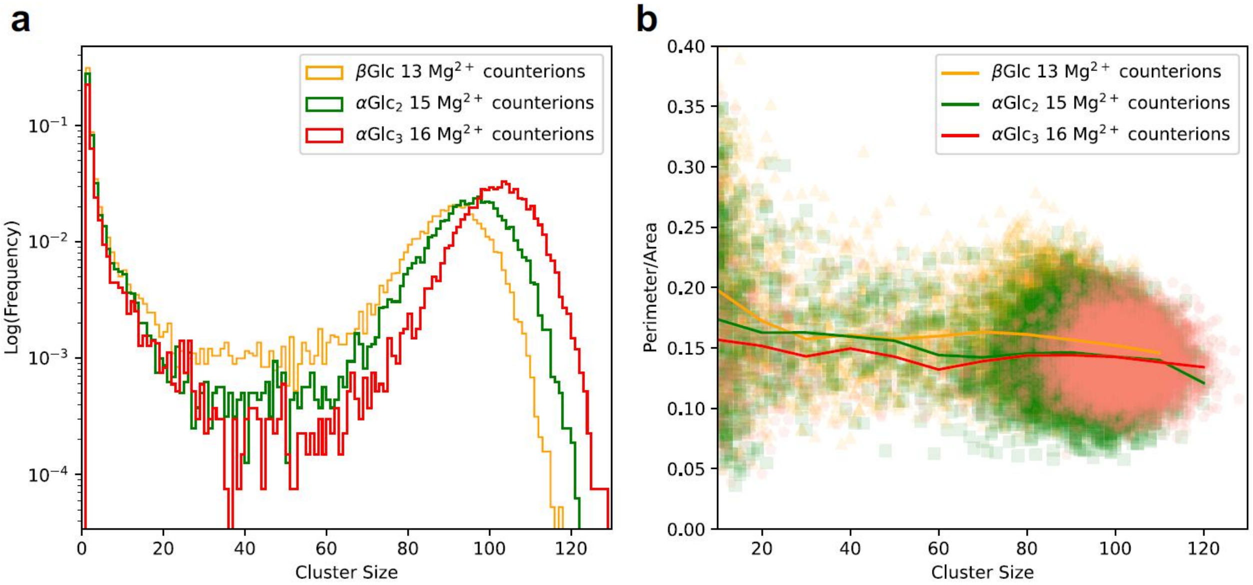

**Figure S93.** (a) Distribution of glycolipid-enriched cluster sizes in systems βGlc **13**, $\alpha$Glc$_2$ **15** and $\alpha$Glc$_3$ **16** glycolipids (systems with $Mg^{2+}$ counterions). Lipids are part of a cluster, if the glycolipid enrichment index $E_A = N^A_{glycolipid} / < N_{glycolipid} > > 1$. $N^A_{glycolipid}$ is the number of glycolipid neighbours within 0.35 nm of lipid A, and $N_{glycolipid}$ is the average across all lipids.[50] (b) Scatter plot for each cluster's perimeter-to-area ratio, $P_i/A_i$, versus cluster size. The solid curve shows the mean $P\ /\ A$ in size of bins of width 10: $< P/A >_{bin} = 1/L \cdot \sum_{i\epsilon[N_{bin}, N_{bin+1}]} P_i/A_i$, where $L$ is the number of clusters in that bin. The area of a cluster $A_i$ was calculated as the sum of the Voronoi cell areas of the lipids comprising the cluster, while the perimeter, $P_i$, was determined by summing the lengths of the Voronoi edges forming the cluster boundary. Voronoi tessellation was performed using the C2 atoms of the lipids' headgroups as reference points. Data was sampled every 10 frames (5 ns), after discarding the first 5 µs for equilibration.

##### 6.3. Brownian Dynamics

The experimental temporal resolution of 40 µs hampers a direct comparison of lipid mean squared displacements analyzed from all-atom biomolecular dynamics simulations. However, by employing a simplified and computationally efficient Brownian dynamics (BD) approach (see Fig. S94) to simulate the two-dimensional motion of lipids, much longer time scales on the order of milliseconds can be reached. In this model, lipids are represented as soft spheres interacting via a Lennard-Jones potential. The algorithm for the evolution of the systems incorporates both a force- and a stochastic contribution:[51,52]

$$\vec{r}(t + \Delta t) = \vec{r}(t) + \frac{D \cdot \Delta t}{RT} \cdot \overrightarrow{F^{LJ}}(\vec{r}) + \sqrt{2 \cdot D \cdot \Delta t} \cdot \vec{G} \qquad (1)$$

$D$ is the diffusion coefficient; $\Delta t$ the BD time step, $R$ the molar gas constant, and $T$ the temperature. The diffusion coefficient $D$ of any particle with radius $R$ in the BD simulation was calculated via the Saffman-Delbrück model,

$$D = \frac{k_B T}{4\pi\eta_m} \cdot \left( \ln\left(\frac{\eta_m}{\eta_f R}\right) - \gamma \right) \qquad (2)$$

$\eta_m$is the membrane surface viscosity ($\approx 10^{-8}$ Pa·s·m), $\eta_f$ is the bulk viscosity of the surrounding solvent ($\approx 2.47^{-3}$Pa·s),[53] $k_B$ the Boltzmann constant, $T$ is the temperature (298 K), and $\gamma$ is Euler's constant (≈ 0.5772). If not otherwise stated a diffusion coefficient of $257.8 \times 10^{-6}$nm$^2$·ns$^{-1}$ was used for the diffusion of the lipids ($R \approx 0.865$ nm). The force term $\overrightarrow{F_{LJ}}(\vec{r})$ emerges from a standard Lennard-Jones potential to model the interaction between the lipids:

$$V_{ij}^{LJ}(r_{ij}) = 4 \cdot \epsilon \cdot \left(\frac{\sigma^{12}}{r_{ij}^{12}} - \frac{\sigma^6}{r_{ij}^6}\right) \tag{3}$$

The parameters $\sigma = 0.7706$ nm and $\epsilon = 0.3221$ kJ · mol$^{-1}$ were fitted to reproduce the radial distribution factor (RDF) of lipids in an equilibrated all-atom membrane patch without glycolipids (System 7, Tab. S2). From the same simulation the value for the average area per lipid (0.66 nm$^2$) was estimated.

To enhance computational efficiency, a pairlist scheme was implemented with a buffer radius of 3 nm, a Lennard-Jones interaction cutoff of 1.2 nm, and an update frequency of 10 steps. Periodic boundary conditions were employed.

$\vec{G}$ is the random displacements of particles, sampled from a standard normal distribution. The scaling factor $\sqrt{2 \cdot D \cdot \Delta t}$ ensures that the random displacement in each spatial dimension has a variance of $2 \cdot D \cdot \Delta t$.

To account for glycolipid clustering in the lipid bilayer due to sugar group interactions, "domains" were introduced in the BD system by defining regions inaccessible to other lipids. For simplicity, the domains were assumed to be circular with a fixed position within the simulation box. The overall domain size was chosen to occupy 30% of the total membrane area. Interaction between domains were modeled using the same $\sigma$ parameter as those governing forces between lipids, and a scaled $\epsilon = 0.88$ accounts for stronger interactions between the sugar headgroups. The Lorentz-Berthelot rules were applied to derive parameters between lipids and domains. This reflects also the behavior observed in the all-atom MD simulations that lipids close to glyco-lipids display a decreased velocity. An offset was subtracted from the distance calculation to account for the domain radius, ensuring that the energy minimum of $V_{ij}^{LJ}(r_{ij})$ coincides with the domain boundary. The number of lipids in the different simulations was chosen to yield similar lipid bulk densities. The mean-square displacement (MSD, Fig. S95) was calculated via a Fourier-based method.[54] The code was written in Python, and will be released as GitHub repository upon acceptance of the paper.

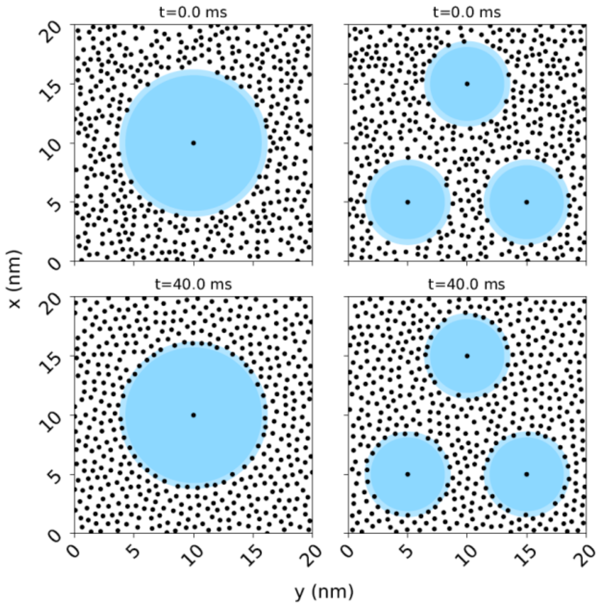

**Figure S94.** Brownian dynamics simulation systems with one large (left panel) and three smaller spherical domains (right panel) with the same total area.

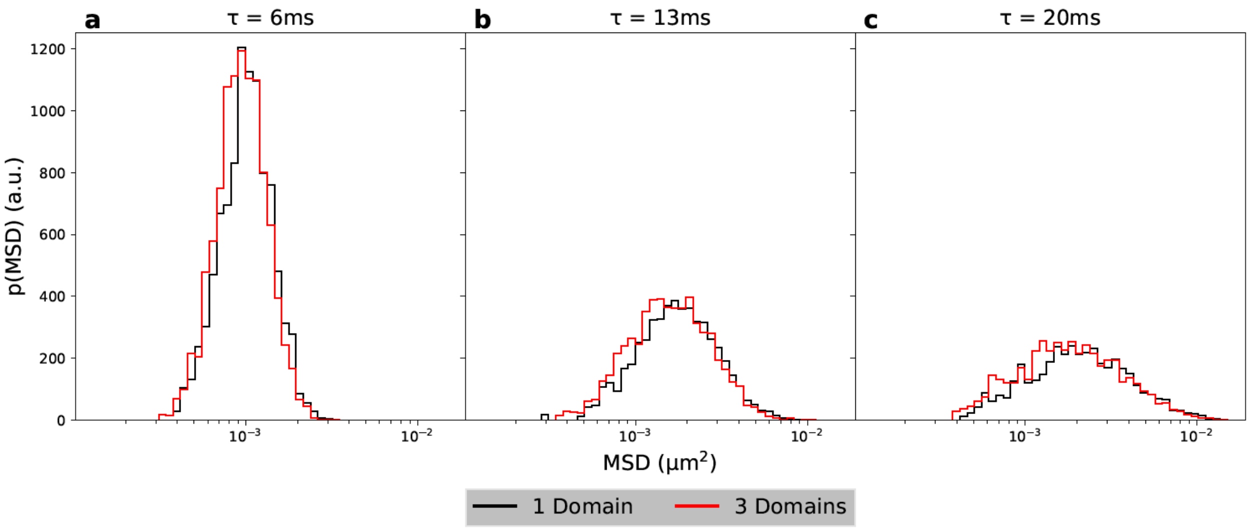

**Figure S95.** Effect of domain size on the lipid dynamics. Histograms of MSD distributions from BD simulations are shown for systems with one large domain (black line) versus three smaller domains of equal total area (red line) at time lags $\tau$ = 6 ms, 13 ms and 20 ms. At each $\tau$, a two-sided, non-parametric Kolmogorov-Smirnov test[55] (*ks_2samp* from the SciPy 1.15.1 package[56]) was performed to test if the samples were drawn from the same distribution ($H_0$) or from different distributions ($H_1$). The p-values of $1.44 \times 10^{-6}$ ($\tau$=6ms), $8.37 \times 10^{-4}$ ($\tau$=13ms), and $10.63 \times 10^{-4}$ ($\tau$=20ms) were considered small enough to reject $H_0$ and assume that the MSD distributions differ significantly between the two domain configurations.

**Table S3.** Brownian Dynamics simulation systems and employed parameters.

| System | $N^{Lipids}$ | $N^{Domains}$ | $r^{Domains}$ | $D^{Lipids}$ ($10^{-6}nm^2 \cdot ns^{-1}$) | $L^{x/y}$(nm) | Δt(ns) | t(μs) |
|---|---|---|---|---|---|---|---|
| 1 | 452 | 1 | 6.18 | 257.8 | 20 | 0.2 | 40 |
| 2 | 472 | 3 | 3.57 | 257.8 | 20 | 0.2 | 40 |
| 3 | 452 | 1 | 6.18 | 257.8 | 20 | 0.2 | 40 |
| 4 | 472 | 3 | 3.57 | 257.8 | 20 | 0.2 | 40 |
| 5 | 452 | 1 | 6.18 | 257.8 | 20 | 0.2 | 40 |
| 6 | 472 | 3 | 3.57 | 257.8 | 20 | 0.2 | 40 |
| 7 | 452 | 1 | 6.18 | 257.8 | 20 | 0.2 | 39.8 |
| 8 | 472 | 3 | 3.57 | 257.8 | 20 | 0.2 | 39.8 |

(56) Virtanen, P.; Gommers, R.; Oliphant, T. E.; Haberland, M.; Reddy, T.; Cournapeau, D.; Burovski, E.; Peterson, P.; Weckesser, W.; Bright, J.; van der Walt, S. J.; Brett, M.; Wilson, J.; Millman, K. J.; Mayorov, N.; Nelson, A. R. J.; Jones, E.; Kern, R.; Larson, E.; Carey, C. J.; Polat, İ.; Feng, Y.; Moore, E. W.; VanderPlas, J.; Laxalde, D.; Perktold, J.; Cimrman, R.; Henriksen, I.; Quintero, E. A.; Harris, C. R.; Archibald, A. M.; Ribeiro, A. H.; Pedregosa, F.; van Mulbregt, P.; Vijaykumar, A.; Bardelli, A. P.; Rothberg, A.; Hilboll, A.; Kloeckner, A.; Scopatz, A.; Lee, A.; Rokem, A.; Woods, C. N.; Fulton, C.; Masson, C.; Häggström, C.; Fitzgerald, C.; Nicholson, D. A.; Hagen, D. R.; Pasechnik, D. V.; Olivetti, E.; Martin, E.; Wieser, E.; Silva, F.; Lenders, F.; Wilhelm, F.; Young, G.; Price, G. A.; Ingold, G.; Allen, G. E.; Lee, G. R.; Audren, H.; Probst, I.; Dietrich, J. P.; Silterra, J.; Webber, J. T.; Slavič, J.; Nothman, J.; Buchner, J.; Kulick, J.; Schönberger, J. L.; Cardoso, J. V. d. M.; Reimer, J.; Harrington, J.; Rodríguez, J. L. C.; Nunez-Iglesias, J.; Kuczynski, J.; Tritz, K.; Thoma, M.; Newville, M.; Kümmerer, M.; Bolingbroke, M.; Tartre, M.; Pak, M.; Smith, N. J.; Nowaczyk, N.; Shebanov, N.; Pavlyk, O.; Brodtkorb, P. A.; Lee, P.; McGibbon, R. T.; Feldbauer, R.; Lewis, S.; Tygier, S.; Sievert, S.; Vigna, S.; Peterson, S.; More, S.; Pudlik, T.; Oshima, T.; Pingel, T. J.; Robitaille, T. P.; Spura, T.; Jones, T. R.; Cera, T.; Leslie, T.; Zito, T.; Krauss, T.; Upadhyay, U.; Halchenko, Y. O.; Vázquez-Baeza, Y. SciPy 1.0: Fundamental Algorithms for Scientific Computing in Python. *Nat. Methods* **2020**, *17* (3), 261–272. https://doi.org/10.1038/s41592-019-0686-2.
